# The global population of SARS-CoV-2 is composed of six major subtypes

**DOI:** 10.1101/2020.04.14.040782

**Authors:** Ivair José Morais Júnior, Richard Costa Polveiro, Gabriel Medeiros Souza, Daniel Inserra Bortolin, Flávio Tetsuo Sassaki, Alison Talis Martins Lima

## Abstract

The World Health Organization characterized the COVID-19 as a pandemic in March 2020, the second pandemic of the 21^st^ century. Severe acute respiratory syndrome coronavirus 2 (SARS-CoV-2) is a positive-stranded RNA betacoronavirus of the family *Coronaviridae*. Expanding virus populations, as that of SARS-CoV-2, accumulate a number of narrowly shared polymorphisms imposing a confounding effect on traditional clustering methods. In this context, approaches that reduce the complexity of the sequence space occupied by the SARS-CoV-2 population are necessary for a robust clustering. Here, we proposed the subdivision of the global SARS-CoV-2 population into sixteen well-defined subtypes by focusing on the widely shared polymorphisms in nonstructural (*nsp*3, *nsp*4, *nsp*6, *nsp*12, *nsp*13 and *nsp*14) cistrons, structural (*spike* and *nucleocapsid*) and accessory (*ORF8*) genes. Six virus subtypes were predominant in the population, but all sixteen showed amino acid replacements which might have phenotypic implications. We hypothesize that the virus subtypes detected in this study are records of the early stages of the SARS-CoV-2 diversification that were randomly sampled to compose the virus populations around the world, a typical founder effect. The genetic structure determined for the SARS-CoV-2 population provides substantial guidelines for maximizing the effectiveness of trials for testing the candidate vaccines or drugs.

## Main

In December 2019, a local pneumonia outbreak of initially unknown etiology was detected in Wuhan (Hubei, China) and quickly determined to be caused by a novel coronavirus^1^, named Severe acute respiratory syndrome coronavirus 2 (SARS-CoV-2)^2^ and the disease as COVID-19^3^. SARS-CoV-2 is classified in the family *Coronaviridae*, genus *Betacoronavirus*, which comprises enveloped, positive stranded RNA viruses of vertebrates^2^. Two-thirds of SARS-CoVs genome is covered by the ORF1ab, that encodes a large polypeptide which is cleaved into 16 nonstructural proteins (NSPs) involved in replication-transcription in vesicles from endoplasmic reticulum (ER)-derived membranes^4,5^. The last third of the virus genome encodes four essential structural proteins: spike (S), envelope (E), membrane (M), nucleocapsid (N) and several accessory proteins that interfere with the host innate immune response^6^.

Populations of RNA viruses evolve rapidly due to their large population sizes, short generation times, and high mutation rates, this latter being a consequence of the RNA-dependent RNA polymerase (RdRP) which lacks the proofreading activity^7^. In fact, virus populations are composed of a broad spectrum of closely related genetic variants resembling one or more master sequences^8–10^. Mutation rates inferred for SARS-CoVs are considered moderate^11,12^ due to the independent proofreading activity^13^. However, the large SARS-CoV genomes (from 27 to 31 kb)^14^ provide to them the ability to explore the sequence space^15^. In order to better understand the diversification of SARS-CoV-2 genomes during the pandemics (from December 2019 to March 25, 2020), we applied a simple, but robust approach to reduce the complexity of the sequence space occupied by the virus population by detecting its widely shared polymorphisms.

The 767 SARS-CoV-2 genomes with high sequencing coverage obtained from GISAID (https://www.gisaid.org/) and GenBank were clustered into 593 haplotypes (Table S1). We conducted a fine-scale sequence variation analysis on the 593 genomes-containing alignment by calculating the nucleotide diversity (π) using a sliding window and step size of 300 and 20 nucleotides, respectively (multiple sequence alignments generated in this study are available from the authors upon request). Such an approach allows to identify genomic regions of increased genetic variation from polymorphic sites harboring two or more distinct nucleotide bases. Noticeably, one or more large clusters of closely related sequences, when analyzed by this approach, show locally increased nucleotide diversity. We observed a contrasting distribution of the genetic variation across the full-length genomes of SARS-CoV-2 (Fig. 1) with eight segments showing increased genetic variation, arbitrarily defined as nucleotide (nt) segments with π ≥ 0.001. Seven out of eight segments (S) had about 280 nucleotides in length, corresponding approximately to the size of a single sliding window, except the S10 whose length was equivalent to two sliding windows (600 nt). To further investigate the diversification of segments with contrasting content of genetic variability, we constructed maximum likelihood (ML) phylogenetic trees and analyzed the diversification patterns of eight segments with higher (S2, 4, 6, 8, 10, 12, 14 and 16), and nine with lower (S1, 3, 5, 7, 9, 11, 13, 15 and 17) content of genetic variation, respectively.

**Fig. 1.**
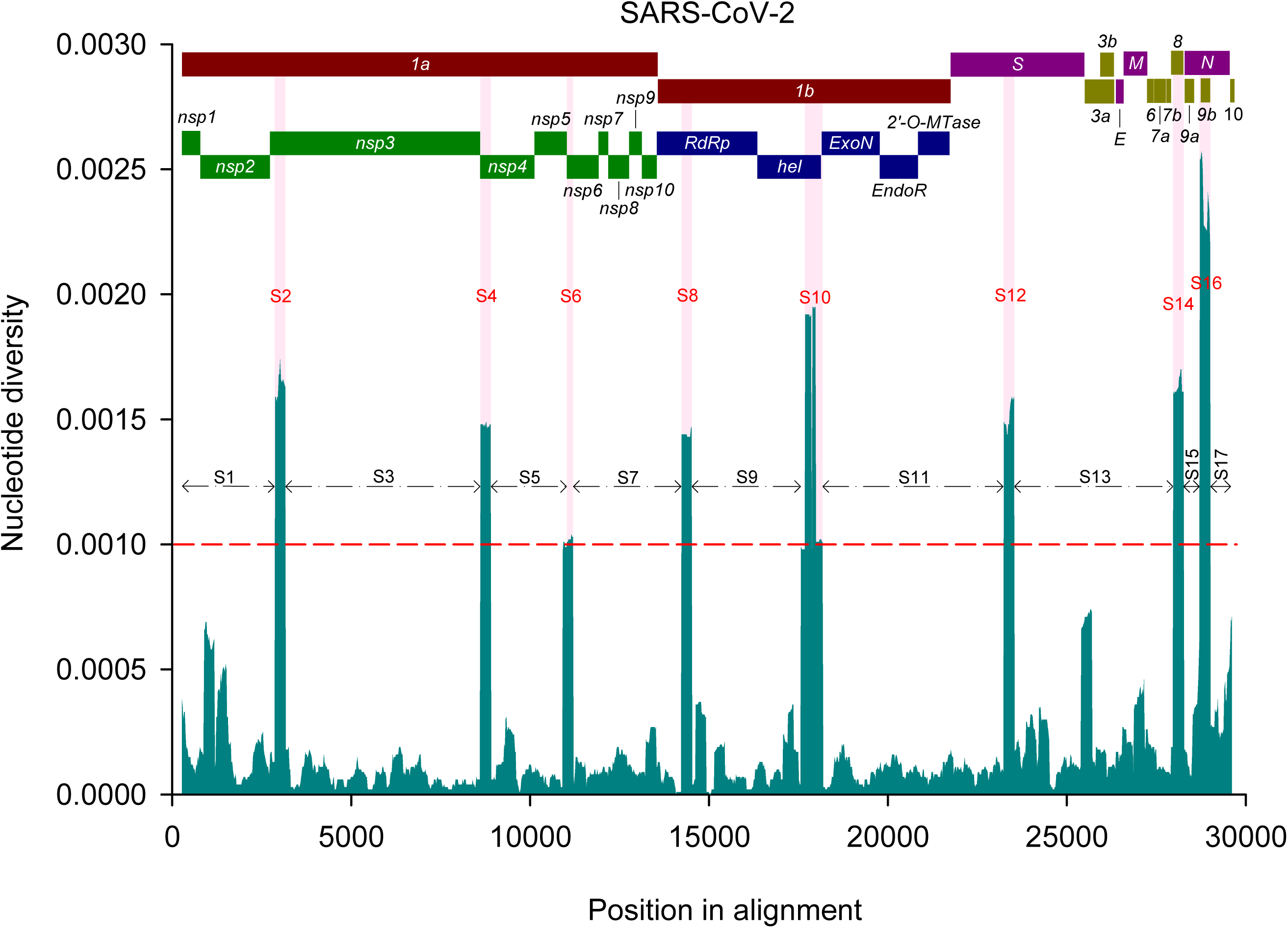
Mean pairwise number of nucleotide differences per site (nucleotide diversity, π) calculated using a sliding window of 300 nucleotides across the multiple sequence alignment for full-length genomes of SARS-CoV-2. The red dashed line at π = 0.001 represents an arbitrary threshold to subdivide the segments (S) with increased (S2, 4, 6, 8, 10, 12, 14 and 16) and lower (S1, 3, 5, 7, 9, 11, 13,15 and 17) content of genetic variation. The SARS-CoV-2 genome organization is represented on top of the plot.

Although the data set was composed of hundreds of SARS-CoV-2 genomes sampled from around the world, in the S2-based tree we observed two clusters (Fig. S1a). Markedly, each cluster was composed of very closely related, if not identical, sequences. Therefore, the increased content of genetic variation at the S2 was a result of the inter-cluster sequence comparisons. Similar results were obtained for the other seven ML-trees based on segments with increased genetic variation (Fig. S1b - h). In contrast, the ML-trees based on segments with lower content of genetic variation did not show a consistent number of well-defined clusters (Fig. S2).

We mapped the polymorphic sites in segments with increased content of genetic variation responsible for the segregation of ML-trees into two well-defined clusters (Table 1). Only a few (from one to three) nt positions with polymorphisms shared by a number of SARS-CoV-2 genomes could be identified within each segment with increased genetic variation. These polymorphisms were henceforth referred to as ‘widely shared polymorphisms’ (WSPs), while the remaining nt positions in virus genomes were designed as ‘non widely shared polymorphisms’ (nWSPs).

**Table 1.** Characterization of the WSPs detected in genomes of SARS-CoV-2

| Segment ID | Segment position* (begin - end) | WSPs† | nt mutation (# isolates) | Position in the codon | #codon | Amino acid |
| --- | --- | --- | --- | --- | --- | --- |
| **S2** | 2,872 - 3,152 | *nsp3*-[318] | **U** (184) / **C** (409) | Third | 106 | Phenylalanine/Phenylalanine |
| **S4** | 8,612 - 8,892 | *nsp4*-[228] | **U** (183) / **C** (410) | Third | 76 | Serine/Serine |
| **S6** | 10,932 - 11,192 | *nsp6*-[111] | **C** (1) / **U** (99) / **G** (493) | Third | 37 | Phenylalanine/Phenylalanine/Leucine |
| **S8** | 14,232 - 14,512 | *nsp12*-[967] | **U** (184) / **C** (409) | Second | 323 | Leucine/Proline |
| **S10** | 17,573 - 18,173 | *nsp13*-[1,511] | **U** (101) / **C** (492) | Second | 504 | Leucine/Proline |
|  |  | *nsp13*-[1,622] | **G** (101) / **A** (492) | Second | 541 | Cysteine/Tyrosine |
|  |  | *nsp14*-[21] | **U** (105) / **C** (488) | Third | 7 | Leucine/Leucine |
| **S12** | 23,243 - 23,523 | *S*-[1,841] | **G** (185) / **A** (408) | Second | 614 | Glycine/Aspartate |
| **S14** | 27,977 - 28,258 | *ORF8*-[251] | **C** (184) / **U** (409) | Second | 84 | Serine/Leucine |
| **S16** | 28,718 - 28,998 | *N*-[608] | **A** (60) / **G** (533) | Second | 203 | Lysine/Arginine |
|  |  | *N*-[609] | **A** (60) / **G** (533) | Third |  |  |
|  |  | *N*-[610] | **C** (60) / **G** (533) | First | 204 | Glycine/Arginine |

\*Relative to the multiple sequence alignment constructed for full-length genomes
†Widely Shared Polymorphisms (WSPs) are conventionally indicated by *cistron/gene*-[nt position within the cistron or gene]

We compared the topologies of the seventeen ML-trees by computing their pairwise distances followed by a multivariate analysis to group similar trees (Fig. 2). The seventeen trees were subdivided into seven groups, the largest one containing those nWSPs-containing segments-based trees (S1, 3, 5, 7, 9, 11, 13, 15 and 17; Fig. 2, Group 7). Given the low content of genetic variation in these segments, the resulting trees were poorly resolved suggesting that such regions represent a wide mutant spectrum of narrowly shared polymorphisms. It is important to note that there are minor clusters in nWSPs-containing segments-based ML-trees, *e.g.*, in those for S1, S13 and S17. This is a consequence of our conservative threshold in which we focused on segments with π ≥ 0.001. S1, S13 and S17 also show locally increased genetic variation with π higher than 0.0005 but lower than 0.001. For example, stretches 889 - 1,169; 1,409 - 1,509 (within the S1); 25,403 - 25,693 (S13); 29,538 - 29,610 (S17) (Fig. 1).

**Fig. 2.**
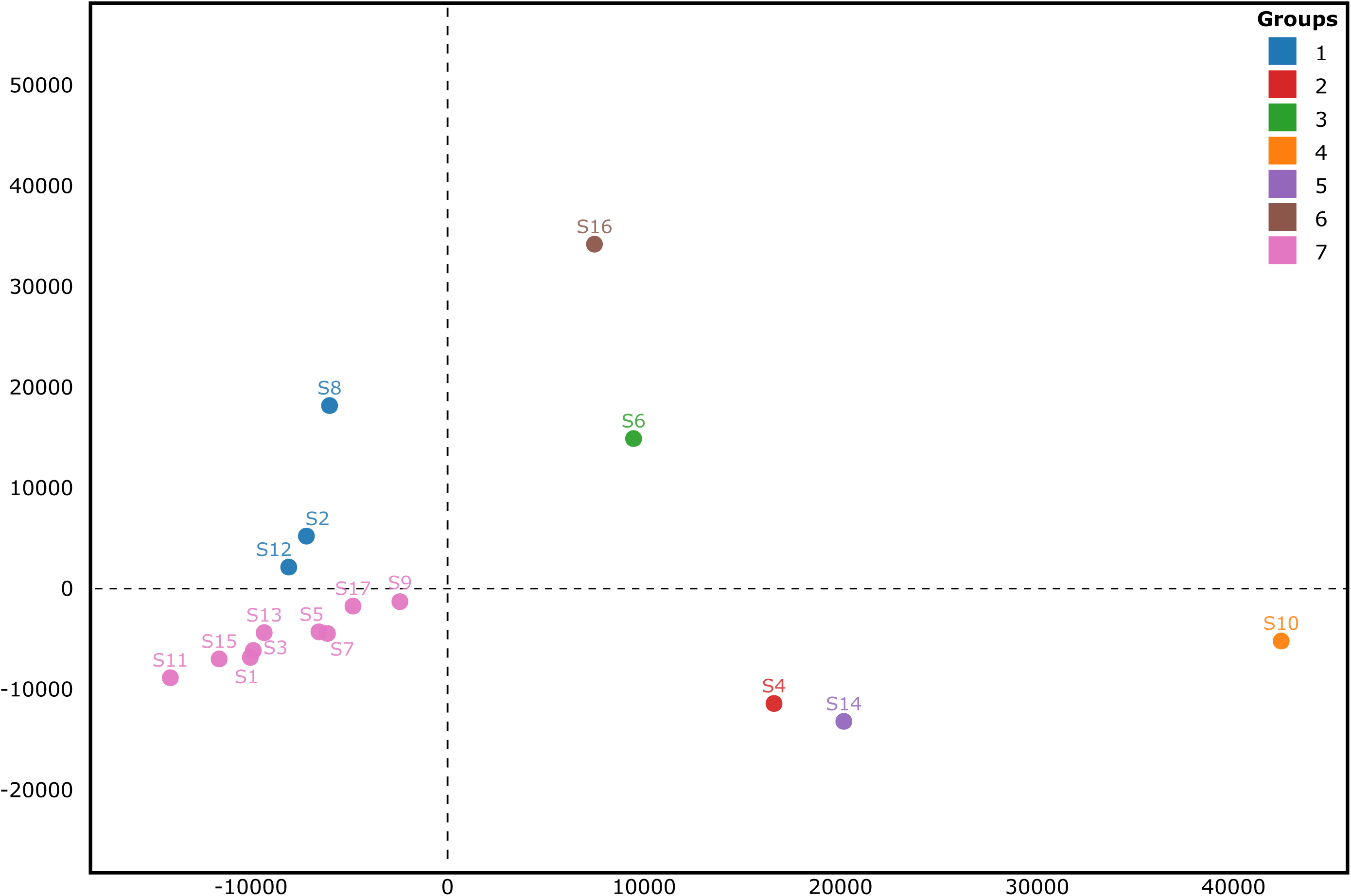
Multidimensional Scaling (MDS) visualization of tree distances based on Kendall Colijn metrics (λ = 0). The seventeen ML-trees (each one with 593 tips) are represented as dots. Seven groups of trees are represented with different colors. The WSP-containing segments-based trees (S2, S8 and S12) formed six groups (1 - 6 are indicated in blue, red, green, orange, purple and brown, respectively). All nWSPs-containing segments-based ML-trees formed a single group indicated in pink.

The S2, S8 and S12-based ML-trees (Fig. 2, Group 1) were considerably congruent and the nucleotides at their WSPs tended to co-segregate (UUG or CCA, Table 1), which results into two major subtypes of SARS-CoV-2. Reciprocally, the incongruency among the S4, 6, 10, 14 and 16-based trees (Fig. 2, Groups 2-6) suggests the segregation of nucleotides at their WSPs, which increases the possible combinations of virus genotypes.

Therefore, our approach reduced the complexity of the sequence space occupied by the SARS-CoV-2 genomes and provided a robust clustering solution based on the combination of 12 WSPs (Table 1) to identify the major viral genotypes spread worldwide (Table 2 and Table S2). The global population of SARS-CoV-2 is structured into six major subtypes (I - VI), comprising 578 out of 593 (about 97.5%) isolates analyzed in this study. The Subtype I (N=132) was represented by the combination of the most frequent nucleotides at all WSPs, *i.g*., the canonical genotype: CCGCCACAUGGG. The SARS-CoV-2 reference isolate (GISAID accession ID: EPI_ISL_402124, GenBank accession: MN908947) is a representative member of this subtype. Subtype IV (N=91) was represented by the combination of the most frequent nucleotides at eleven out of 12 WSPs (**<u>CC</u>**U**<u>CCACAUGGG</u>**; the most frequent nucleotides at each WSP are highlighted in bold and underlined). Subtypes V (N=74, **<u>C</u>**U**<u>GCCACA</u>**C**<u>GGG</u>**), II (N=122, U**<u>CG</u>**U**<u>CAC</u>**G**<u>UGGG</u>**), III (N=101, **<u>C</u>**U**<u>GC</u>**UGU**<u>A</u>**C**<u>GGG</u>**) and VI (N=58, U**<u>CG</u>**U**<u>CAC</u>**G**<u>U</u>**AAC) were represented by the combination of the most frequent nucleotides at ten, nine, seven and six out of 12 WSPs, respectively. It is important to emphasize that the contrasting sample sizes (Subtypes I - VI *vs.* VII - XVI) are not necessarily associated with fitness variation and might be due to a sampling bias. For example, the three isolates composing the Subtype VIII showed a genotype (**<u>C</u>**U**<u>GCCA</u>**U**<u>ACGGG</u>**) very similar to that of the canonical reference isolate. In addition, even though our data set was composed exclusively by genomes with high sequencing coverage, we cannot rule out that the virus subtypes X to XVI, which were represented by a single genome might be a consequence of poor sampling or sequencing errors.

**Table 2.** Unique genotypes of SARS-CoV-2 based on 12 WSPs and their associated amino acid replacements

| SUBTYPES | N* | S2<sup>‡</sup> | S4 | S6 | S8 | S10 |  |  | S12 | S14 | S16 |  |  |
| --- | --- | --- | --- | --- | --- | --- | --- | --- | --- | --- | --- | --- | --- |
|  |  | *nsp3* | *nsp4* | *nsp6* | *nsp12* | *nsp13* |  | *nsp14* | *S* | *ORF8* | *N* |  |  |
|  |  | #318 | #228 | #111 | #967 | #1511 | #1622 | #21 | #1,841 | #251 | #608 | #609 | #610 |
| I | 132 | C [Phe]<sup>†</sup> | C [Ser] | G [Leu] | C [Pro] | C [Pro] | A [Tyr] | C [Leu] | A [Asp] | U [Leu] | G [Arg] | G [Arg] | G [Arg] |
| II | 122 | U [Phe] | C [Ser] | G [Leu] | U [Leu] | C [Pro] | A [Tyr] | C [Leu] | G [Gly] | U [Leu] | G [Arg] | G [Arg] | G [Arg] |
| III | 101 | C [Phe] | U [Ser] | G [Leu] | C [Pro] | U [Leu] | G [Cys] | U [Leu] | A [Asp] | C [Ser] | G [Arg] | G [Arg] | G [Arg] |
| IV | 91 | C [Phe] | C [Ser] | U [Phe] | C [Pro] | C [Pro] | A [Tyr] | C [Leu] | A [Asp] | U [Leu] | G [Arg] | G [Arg] | G [Arg] |
| V | 74 | C [Phe] | U [Ser] | G [Leu] | C [Pro] | C [Pro] | A [Tyr] | C [Leu] | A [Asp] | C [Ser] | G [Arg] | G [Arg] | G [Arg] |
| VI | 58 | U [Phe] | C [Ser] | G [Leu] | U [Leu] | C [Pro] | A [Tyr] | C [Leu] | G [Gly] | U [Leu] | A [Lys] | A [Lys] | C [Gly] |
| VII | 3 | C [Phe] | U [Ser] | U [Phe] | C [Pro] | C [Pro] | A [Tyr] | C [Leu] | A [Asp] | C [Ser] | G [Arg] | G [Arg] | G [Arg] |
| VIII | 3 | C [Phe] | U [Ser] | G [Leu] | C [Pro] | C [Pro] | A [Tyr] | U [Leu] | A [Asp] | C [Ser] | G [Arg] | G [Arg] | G [Arg] |
| IX | 2 | U [Phe] | C [Ser] | U [Phe] | U [Leu] | C [Pro] | A [Tyr] | C [Leu] | G [Gly] | U [Leu] | A [Lys] | A [Lys] | C [Gly] |
| X | 1 | U [Phe] | C [Ser] | U [Phe] | U [Leu] | C [Pro] | A [Tyr] | C [Leu] | G [Gly] | U [Leu] | G [Arg] | G [Arg] | G [Arg] |
| XI | 1 | U [Phe] | C [Ser] | G [Leu] | C [Pro] | C [Pro] | A [Tyr] | C [Leu] | G [Gly] | U [Leu] | G [Arg] | G [Arg] | G [Arg] |
| XII | 1 | C [Phe] | U [Ser] | C [Phe] | C [Pro] | C [Pro] | A [Tyr] | C [Leu] | A [Asp] | C [Ser] | G [Arg] | G [Arg] | G [Arg] |
| XIII | 1 | C [Phe] | C [Ser] | G [Leu] | C [Pro] | C [Pro] | A [Tyr] | C [Leu] | A [Asp] | C [Ser] | G [Arg] | G [Arg] | G [Arg] |
| XIV | 1 | C [Phe] | U [Ser] | G [Leu] | C [Pro] | C [Pro] | A [Tyr] | C [Leu] | G [Gly] | C [Ser] | G [Arg] | G [Arg] | G [Arg] |
| XV | 1 | C [Phe] | C [Ser] | U [Phe] | U [Leu] | C [Pro] | A [Tyr] | C [Leu] | A [Asp] | U [Leu] | G [Arg] | G [Arg] | G [Arg] |
| XVI | 1 | C [Phe] | C [Ser] | U [Phe] | C [Pro] | C [Pro] | A [Tyr] | U [Leu] | A [Asp] | U [Leu] | G [Arg] | G [Arg] | G [Arg] |

\*Sample size
<sup>‡</sup>Segment containing the WSP
<sup>#</sup>Nucleotide position
<sup>†</sup>Nucleotide base and the encoded amino acid residue

The phylogenetic tree depicting all 593 SARS-CoV-2 haplotypes (Fig. 3) showed some geographical structure with two clusters: a smaller one comprised of isolates mostly sampled from Western hemisphere (Subtypes II, VI, IX, X and XI) and a larger one whose isolates were sampled from Western and Eastern hemispheres (I, III, IV, V, VII, VIII, XII, XIII, XIV, XV and XVI).

**Fig. 3.**
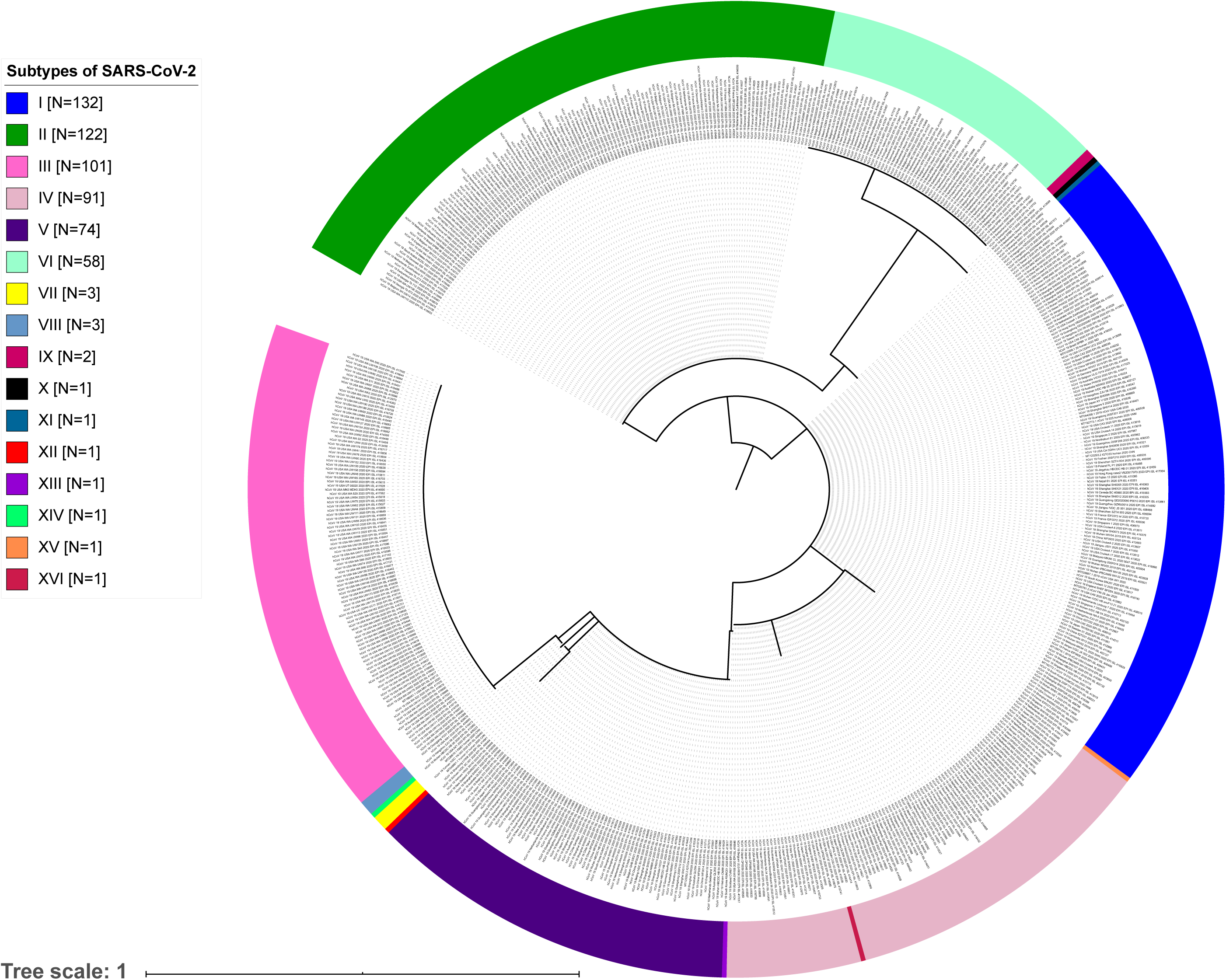
Maximum likelihood phylogenetic tree based on 12 WSPs detected across the SARS-CoV-2 genomes.

We hypothesize that our clustering method for the SARS-CoV-2 population could involve at some extent a biological context. Nine out of 12 WSPs led to amino acid replacements (Table 2), *e.g.*, the WSP *nsp*6-[111] in nine SARS-CoV-2 subtypes led to a leucine at the aa residue#37 of the protein and a phenylalanine in seven other subtypes. NSP6 is an integral membrane protein that interferes in the autophagosome formation during the SARS-CoV infection. Additionally, in yeast two-hybrid experiments, NSP6 has been shown to interact with NSP3^16^. Some evidence demonstrates that NSP6 protein limits the expansion of autophagosomes or, alternatively, might remove host proteins involved in inhibition of viral replication by activating autophagy from the ER^17^.

The WSP *nsp*12-[967] resulted in a proline in eleven subtypes of SARS-CoV-2 and a leucine in others five subtypes at the aa residue #323 of the NSP12 (RNA-dependent RNA polymerase, RdRP) protein. It is located at the Interface domain of RdRP of SARS-CoV-2, which is responsible for the connection between the nidovirus RdRP-associated nucleotidyltransferase domain (NiRAN) and the “Right hand” polymerase domain^18^. The S protein mediates viral entry into host cells by first binding to a receptor, angiotensin-converting enzyme 2 (ACE2), through the receptor-binding domain (RBD) in the S1 subunit and then fusing the viral and host membranes through the S2 subunit^19–22^. Sites of glycosylation are important for S protein folding^23^, affecting priming by host proteases^24^ and might modulate antibody recognition^25,26^. The WSP *S*-[1,841] resulted in a glycine and an aspartate at the aa residue#614 of the S protein in six and ten subtypes of SAR-CoV-2, respectively. The replacement was mapped in the intermediate region between the S1 and S2 subunits. This WSP is near a glycosylation site (N616CT)^27^.

The WSP *ORF8*-[251] involved a non-synonymous mutation at the codon#84 encoding for leucine and serine in nine and seven subtypes, respectively. The SARS-CoV *ORF8* encodes for an ER-associated protein that induces the activation of ATF6, and this latter is an ER stress-regulated transcription factor that stimulates the production of chaperones^28^. In addition, the ORF8 protein has been demonstrated to induce apoptosis^29^. In SARS epidemics, the *ORF8* from different coronaviruses was targeted by a number of mutations and recombination events during transmission from animals to humans^30^.

Three WSPs mapped in the *N* gene led to two amino acid replacements at residues#203 and #204. The multifunctional N protein is composed of three domains^31^, two of which are structurally independent: the N-terminal domain (NTD) and the C-terminal domain (CTD). Both amino acid replacements were mapped in an intermediary domain referred to as the linker region (LKR), a positively charged serine-arginine-rich region. As an intrinsically disordered region (IDR) it allows the independent folding of the NTD and CTD^32^ and is also functionally implicated in RNA binding activity^31^. Key determinants of the interaction between the N and NSP3 proteins were also mapped at the LKR^33^. The SARS-CoV N protein is also responsible for an antigenic response in humans predominantly involving the immunoglobulin G^34^. Although the host biological factors involved in the response to SARS-CoV-2 infection are still poorly known, the existence of distinct virus subtypes, all of them exhibiting amino acid replacements, could alter important aspects of COVID-19.

We hypothesized that in the early stages of the SARS-CoV-2 epidemics, due to the rapid virus population expansion, a number of genetic variants might have arisen followed by a spread of non-representative sampling of variants to other countries and continents, *i.g.*, a founder effect. We argue that the virus subtypes and their associated WSPs detected in this study would be records of diversification in these early stages of the epidemics after transmission from animal to humans. After the virus introduction in a given geographic region, a number of unique or narrowly shared mutations is accumulated, however, most of them reduce the fitness and are removed by purifying selection in a medium to long term evolutionary scale, tending to a decreasing genetic variability^8^.

We propose a classification into at least sixteen distinct subtypes of SARS-CoV-2, six of them accounting for more than 97% of the sampled isolates from around the world. Such classification might guide the validation of candidate vaccines or drugs for the widest range of virus subtypes. In this context, our clustering solution provides a robust approach to effectively reduce the complexity of the mutant spectrum composed of closely related SARS-CoV-2 genomes focusing on WSPs. Additionally, through the exhaustive sequencing, it would be possible to identify novel virus subtypes and follow the evolutionary dynamics of the SARS-CoV-2 population during the adaptive process imposed by the human host.

## Methods

A total of 1,137 full-length genomes of SARS-CoV-2 sampled from December 2019 to March 25, 2020 (at 2:30 pm) were obtained from Genbank^35^ and GISAID^36^ (Table S1). Only genomes with high sequencing coverage, intact ORFs (no frameshifts, except that of *nsp*12 cistron) and without any indeterminate nucleotide base (indicated by ‘N’s or ambiguous codes) totalizing 767 high quality full-length sequences were effectively analyzed in this study. We wish to acknowledge all researchers that deposited the SARS-CoV-2 genomes in GISAID and/or GenBank databases.

The genomic data set was aligned using MAFFT-FFT-NS-2^37^. The calculation of the average number of nucleotide differences per site (nucleotide diversity, π) was conducted in DnaSP v.6^38^ using a sliding window and step size of 300 and 20 nucleotides, respectively. Sites with gaps alignment were not considered in analysis.

Maximum likelihood (ML) phylogenetic trees were constructed using RAxML^39^ under the nucleotide substitution model General Time-Reversible with gamma distribution (GTRGAMMA). The branch support for ML-trees based on 300-nucleotides and larger segments was assessed with 1,000 and 5,000 bootstrap replications, respectively. ML-trees were used in this study essentially as a clustering method due to the weak phylogenetic signal in the data set. All phylogenetic trees were edited using iTOL^40^. In order to assess the similarity among ML-tree topologies, we computed all possible pairwise distances using the Kendall–Colijn metric^41^ followed by Principal Coordinates Analysis (PCoA) using the package TREESPACE^42^ in R^43^.

The detection of polymorphic sites was conducted using PAUP* v. 4.0^44^ and MEGA X^45^. Those sites responsible for the segregation of the isolates into two clusters in the ML-trees were referred to as “widely shared polymorphisms” (WSPs), while the remaining nt positions in the virus genomes were designed as “non widely shared polymorphisms” (nWSPs). The WSPs were conventionally indicated by *cistron/gene* name-[nt position within the cistron or gene]. We opted by indicating the nt position within the cistron or gene due to their highly conserved lengths (no gap was introduced during the construction of sequence alignments), in contrast to the full-genomes whose 5’- and 3’-untranslated regions (UTRs) were highly variable in terms of length.

## Data Availability

multiple sequence alignments and ML-phylogenetic trees generated in this study are available from the authors upon request.

## Acknowledgments

This study was financed in part by the Coordenação de Aperfeiçoamento de Pessoal de Nível Superior - Brasil (CAPES) - Finance Code 001. IJM and RCP were recipients of CNPq and CAPES doctoral fellowships, respectively. DIB was the recipient of a FAPEMIG master fellowship. We wish to acknowledge all researchers that deposited the SARS-CoV-2 genomes in GISAID and/or GenBank databases.

## Author Contributions

ATML designed the bioinformatics analyses. IJM, ATML, RCP, GMS, DIB and FTS conducted the analyses. ATML, IJM, RCP and FTS analyzed data and results. IJM, RCP, FTS and ATML wrote the manuscript. All authors contributed to the content and writing of the Supplementary Information.

## Additional information

Supplementary information is available for this manuscript.

