## Supplementary Information for "The global population of SARS-CoV-2 is composed of six major subtypes"

**Supplementary Fig. S1 |** Maximum likelihood (ML) phylogenetic trees based on nucleotide segments with increased genetic variation (**a**) S2, (**b**) S4, (**c**) S6, (**d**) S8, (**e**) S10, (**f**) S12, (**g**) S14, (**h**) S16.

a)

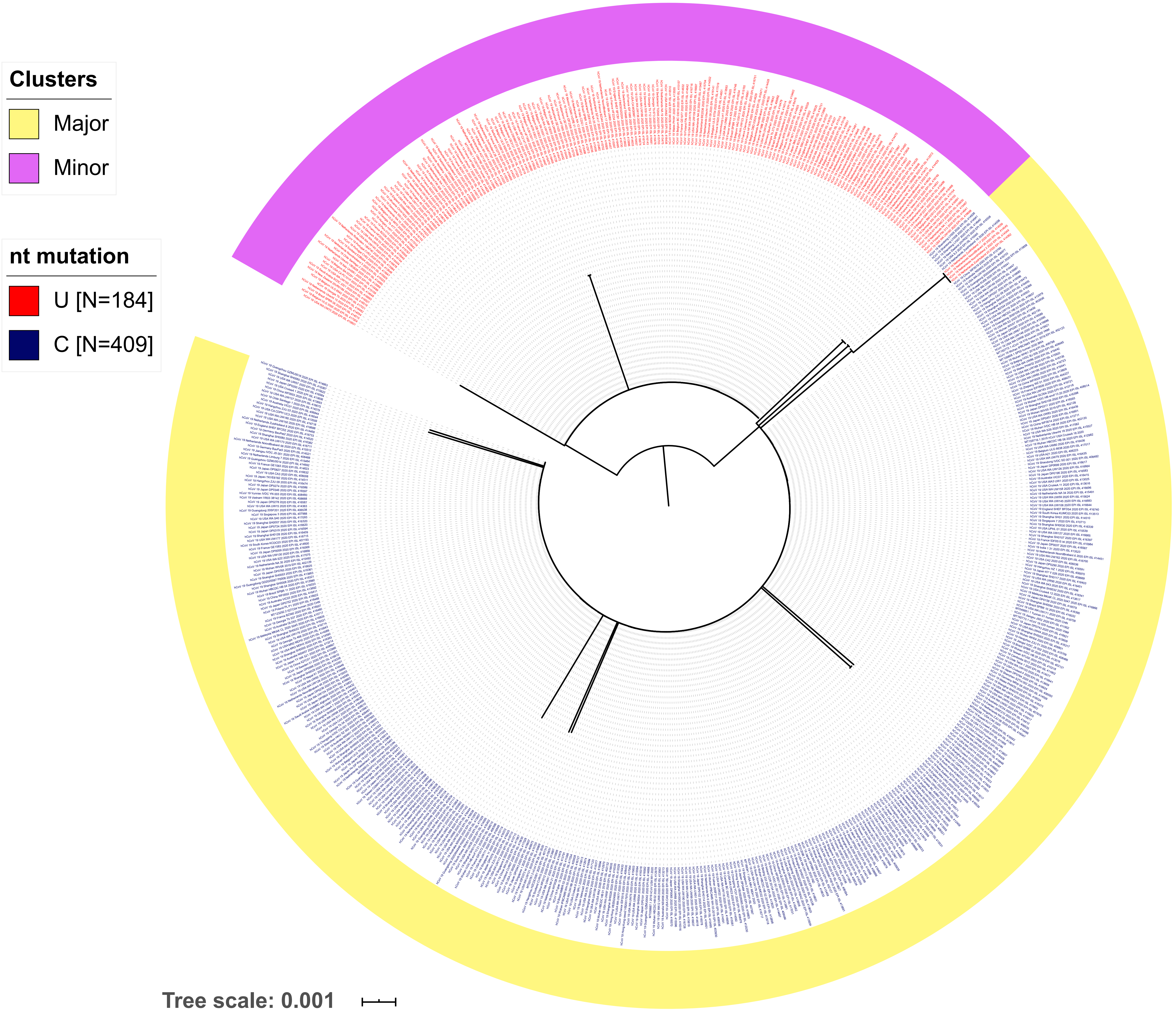
Clusters
Major
Minor
nt mutation
U [N=184]
C [N=409]
Tree scale: 0.001

b)

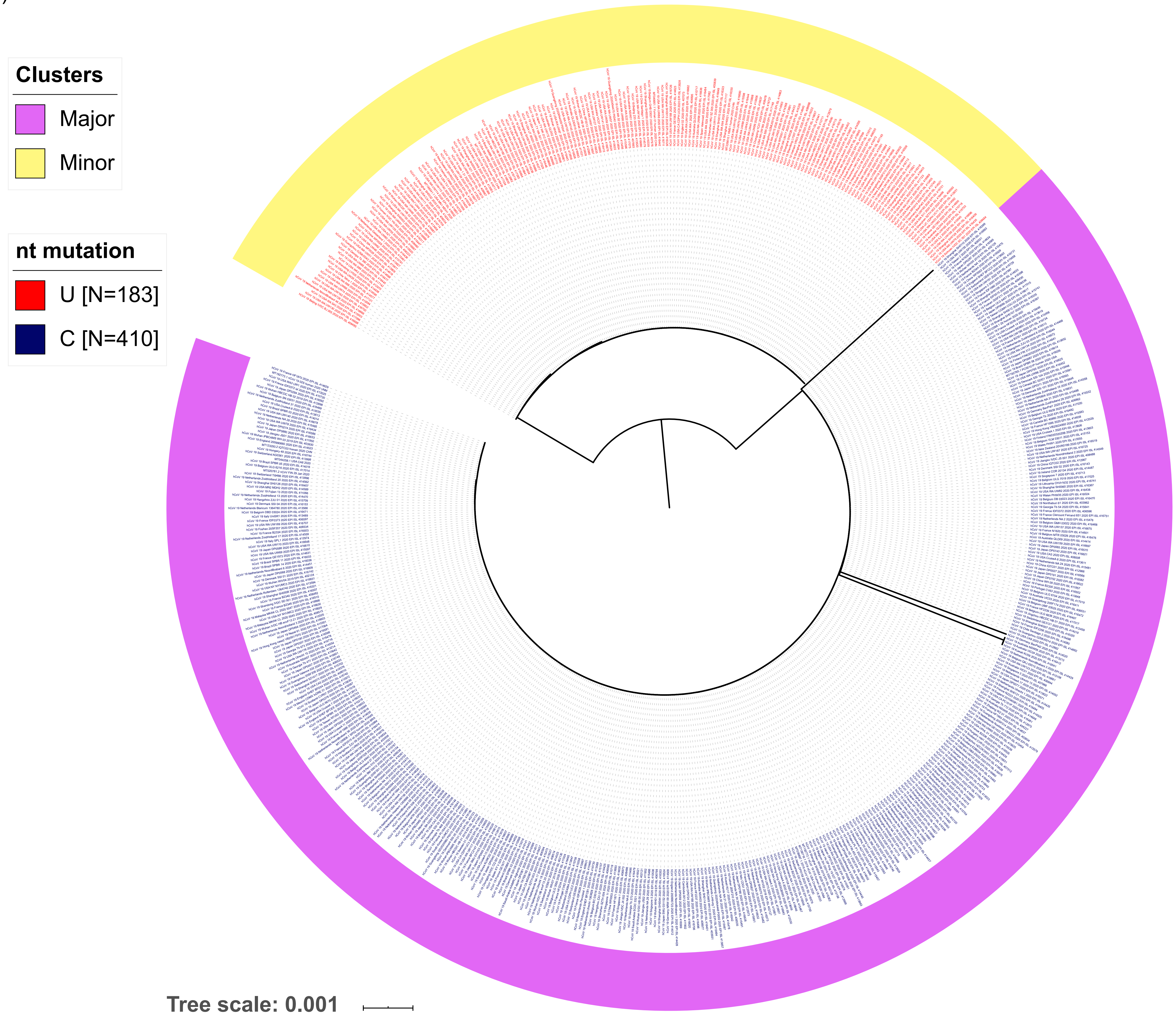
Clusters
Major
Minor
nt mutation
U [N=183]
C [N=410]
Tree scale: 0.001

c)

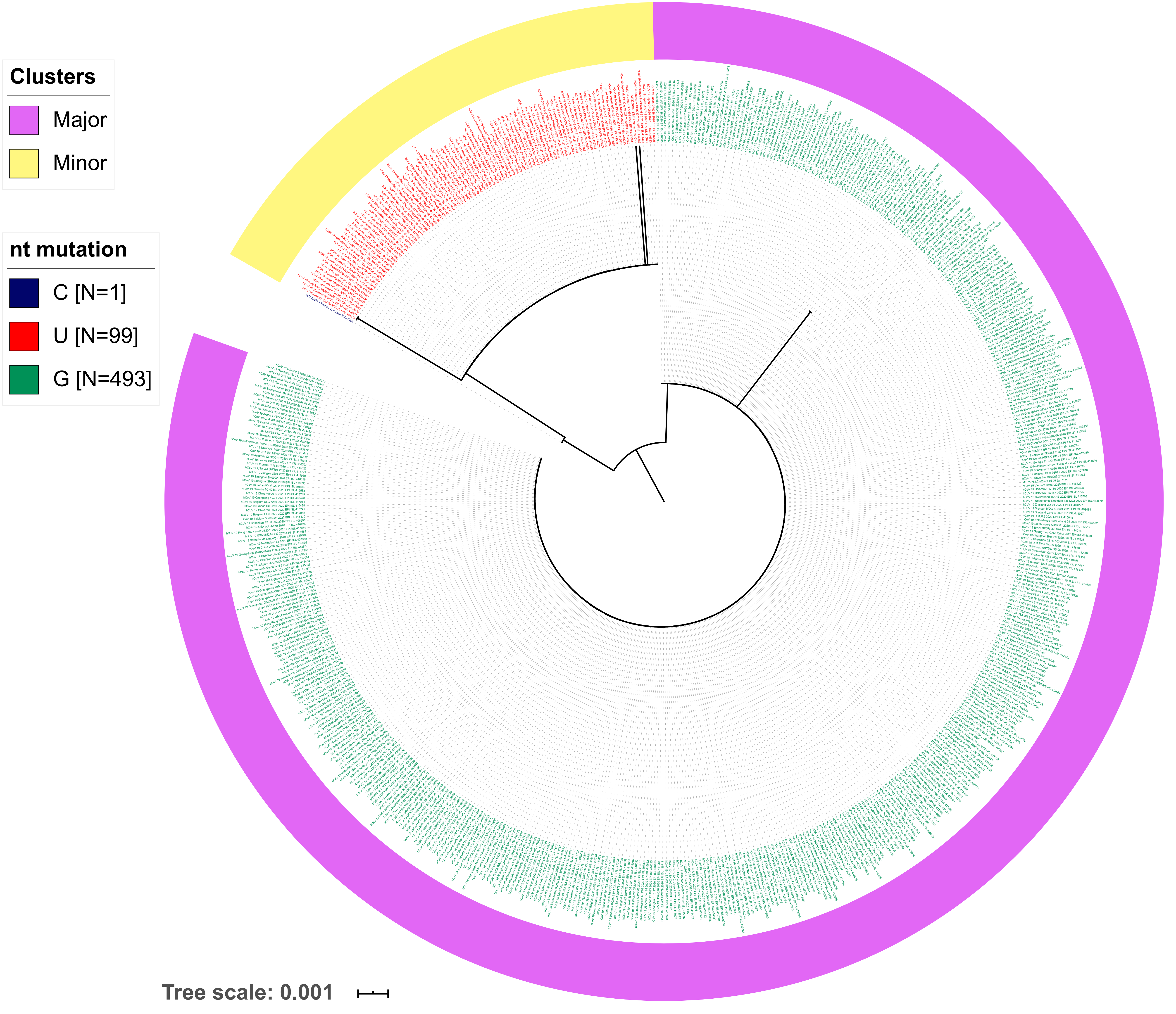
Clusters
Major
Minor
nt mutation
C [N=1]
U [N=99]
G [N=493]
Tree scale: 0.001

d)

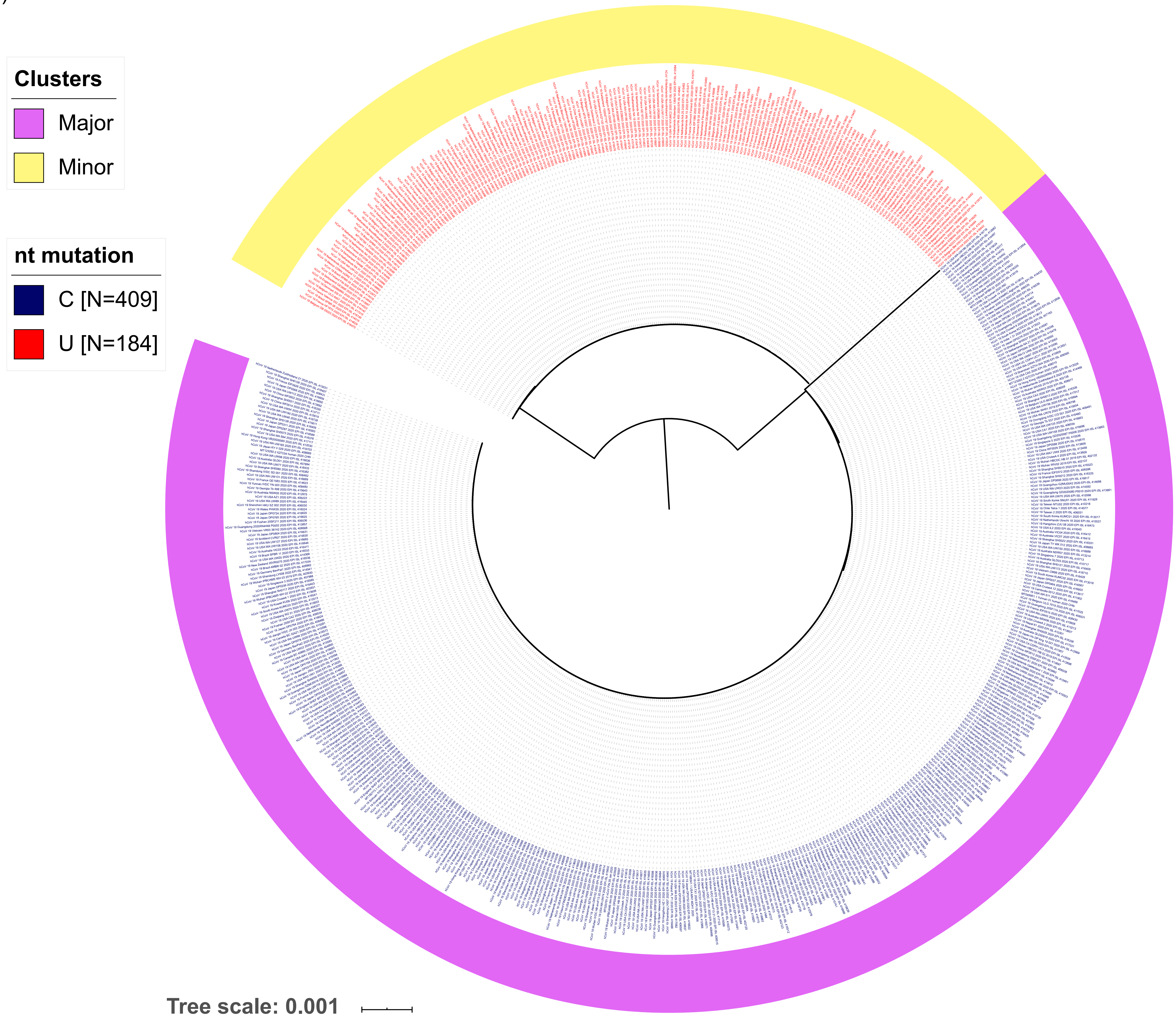
Clusters
Major
Minor
nt mutation
C [N=409]
U [N=184]
Tree scale: 0.001

e)

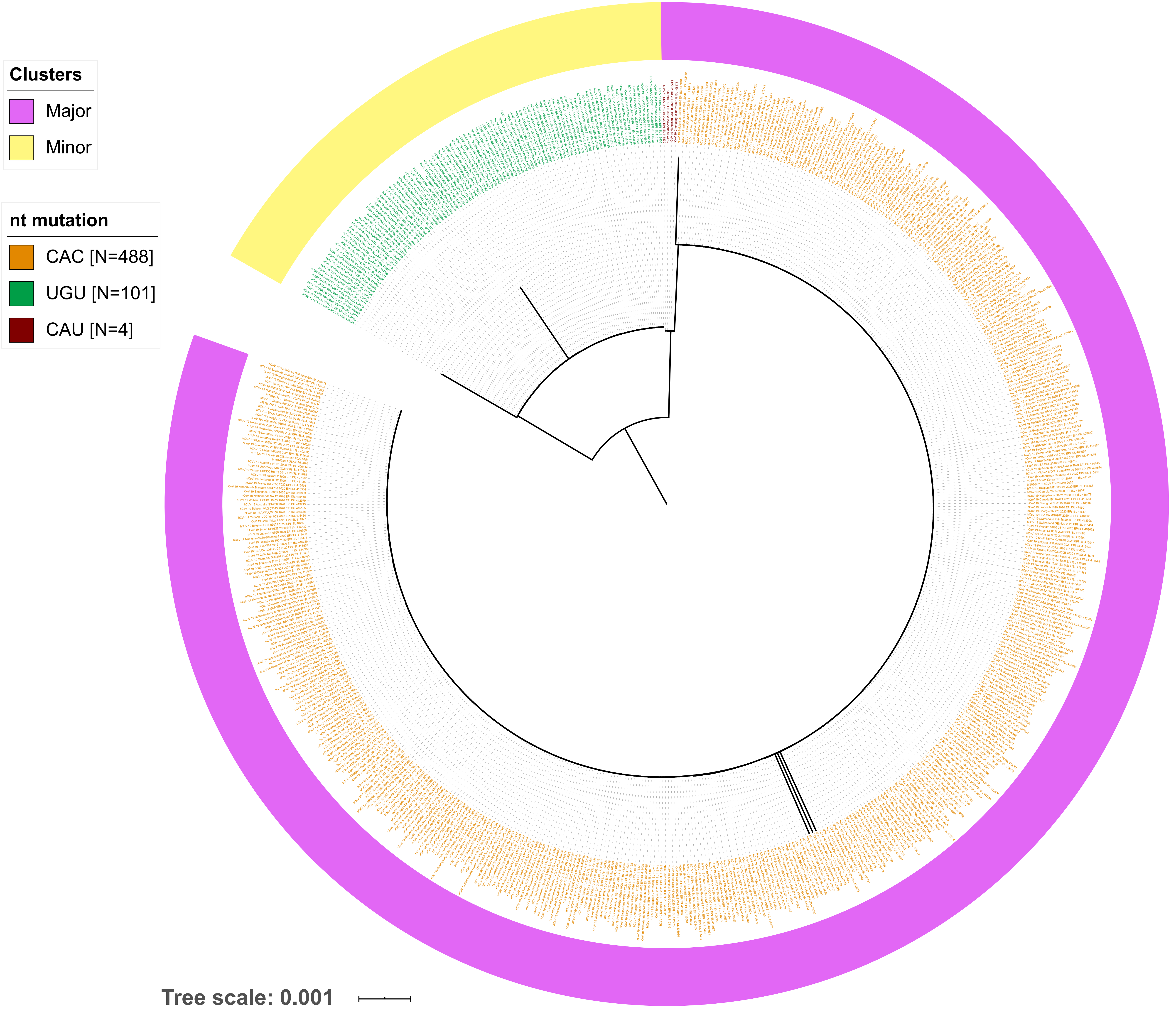
Clusters
Major
Minor
nt mutation
CAC [N=488]
UGU [N=101]
CAU [N=4]
Tree scale: 0.001

f)

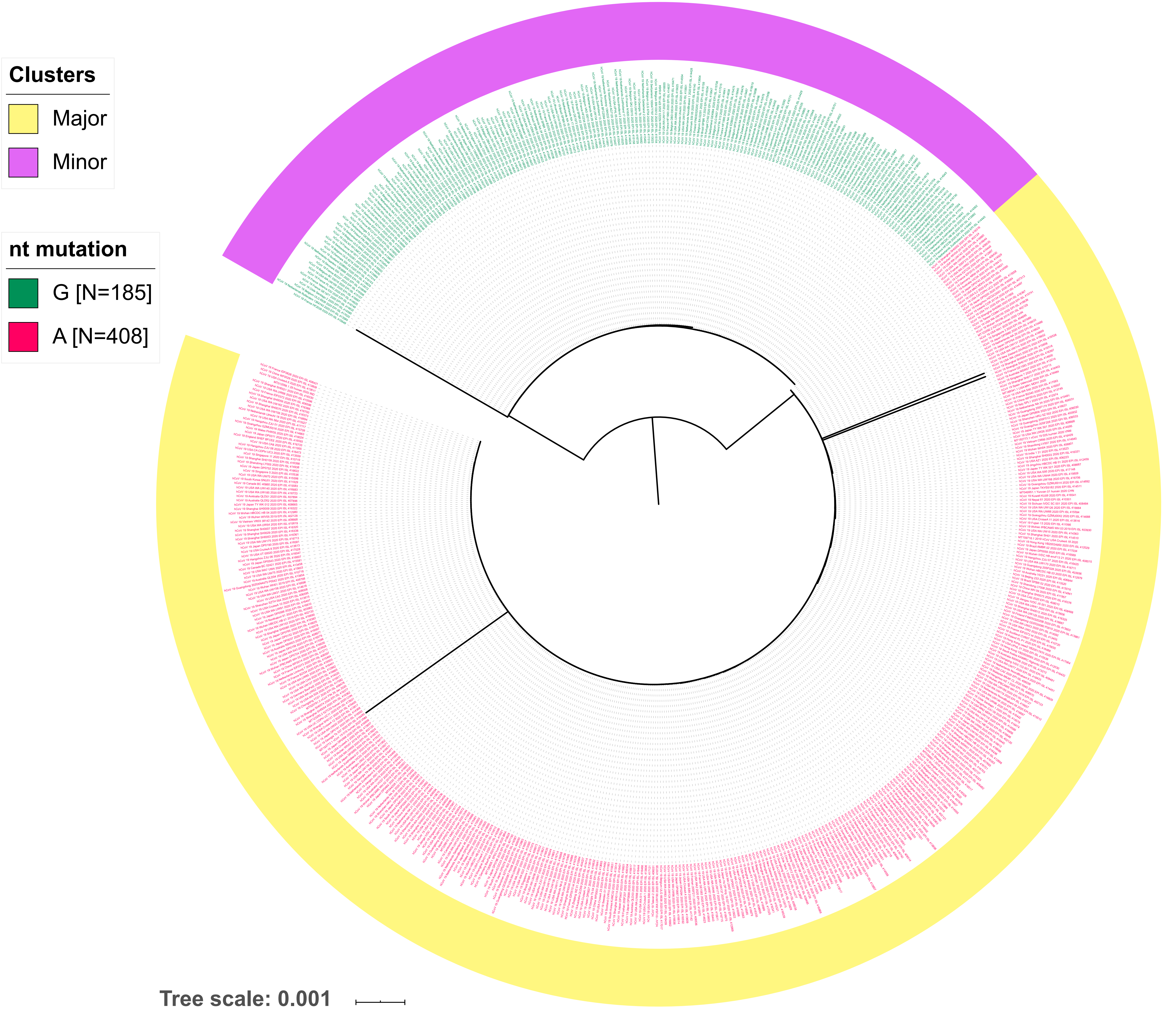

Clusters
Major
Minor
nt mutation
G [N=185]
A [N=408]
Tree scale: 0.001

g)

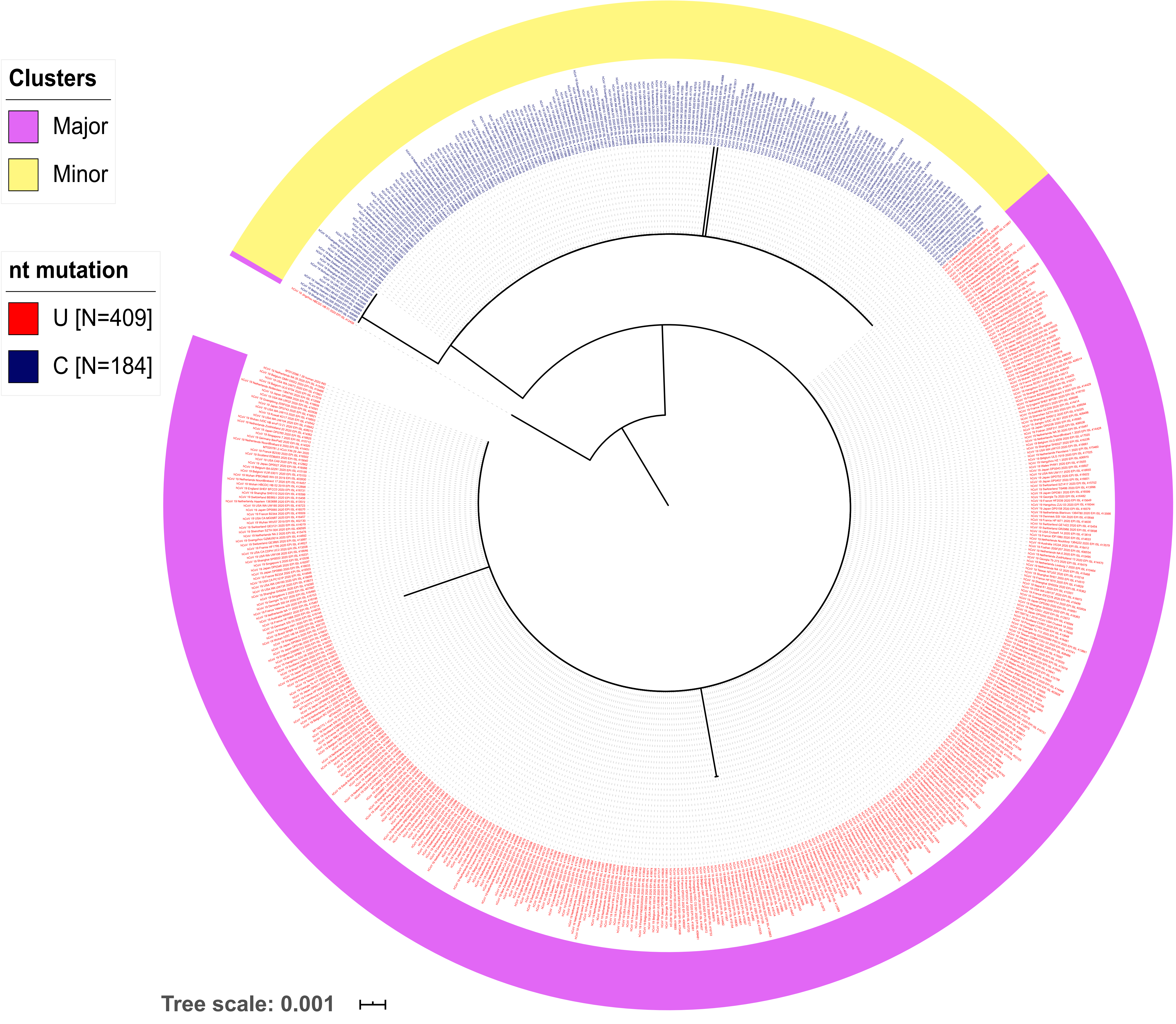
Clusters
Major
Minor
nt mutation
U [N=409]
C [N=184]
Tree scale: 0.001

h)

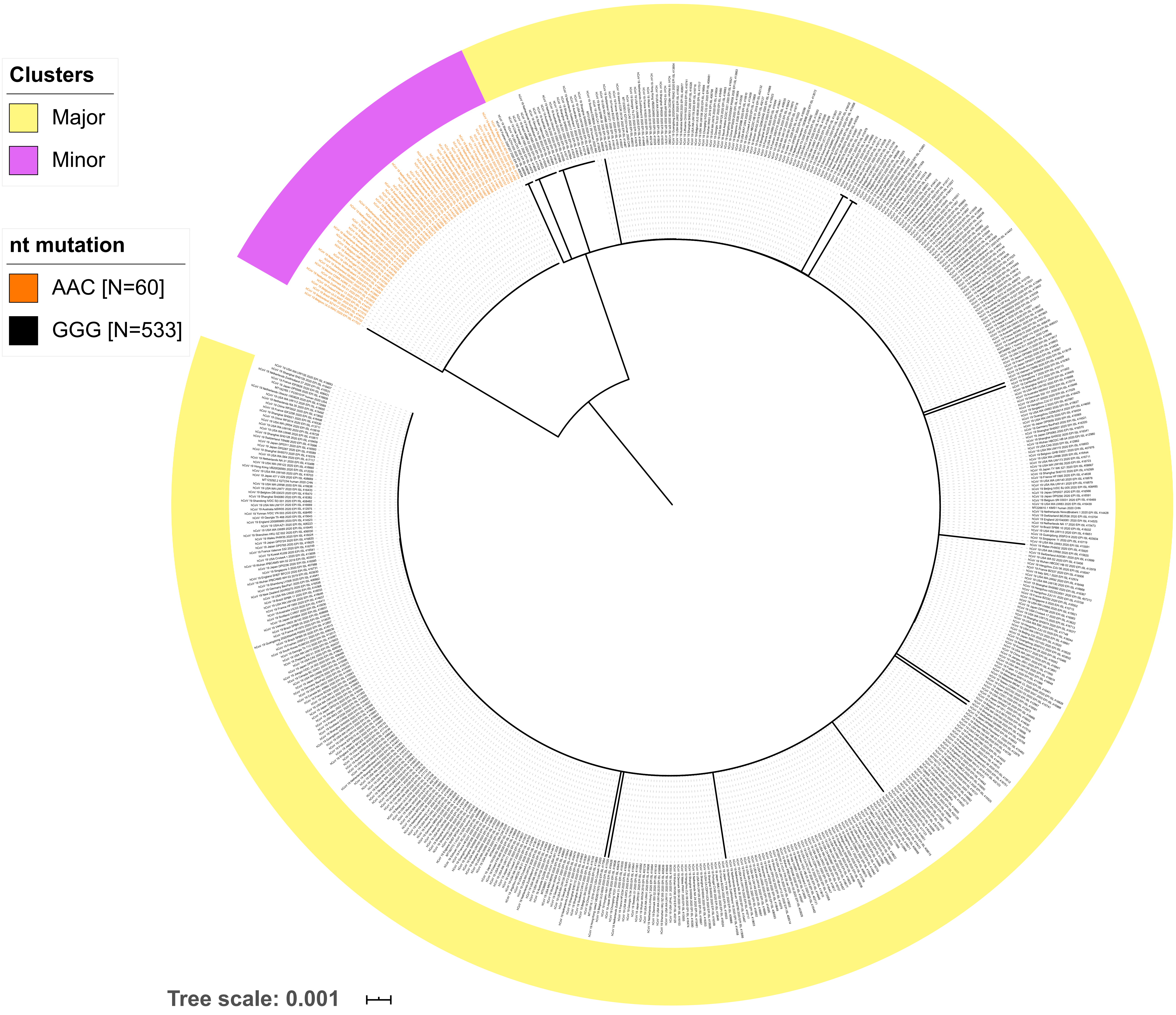
Clusters
Major
Minor
nt mutation
AAC [N=60]
GGG [N=533]
Tree scale: 0.001

**Supplementary Fig. S2 |** Maximum likelihood (ML) phylogenetic trees based on nucleotide sequences of segments with lower content of genetic variation (**a**) S1, (**b**) S3, (**c**) S5, (**d**) S7, (**e**) S9, (**f**) S11, (**g**) S13, (**h**) S15 and (**i**) S17.

a)

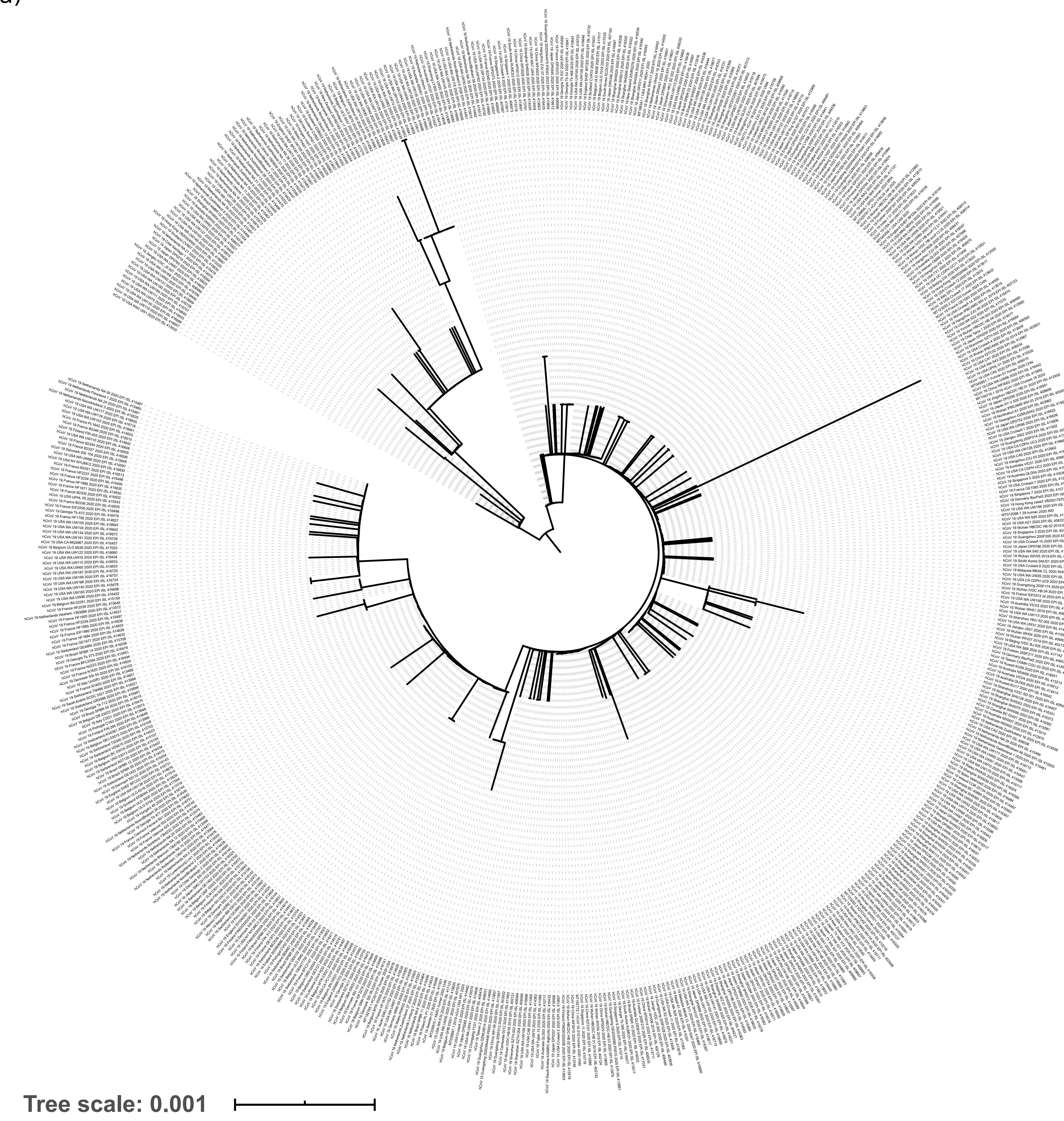

b)

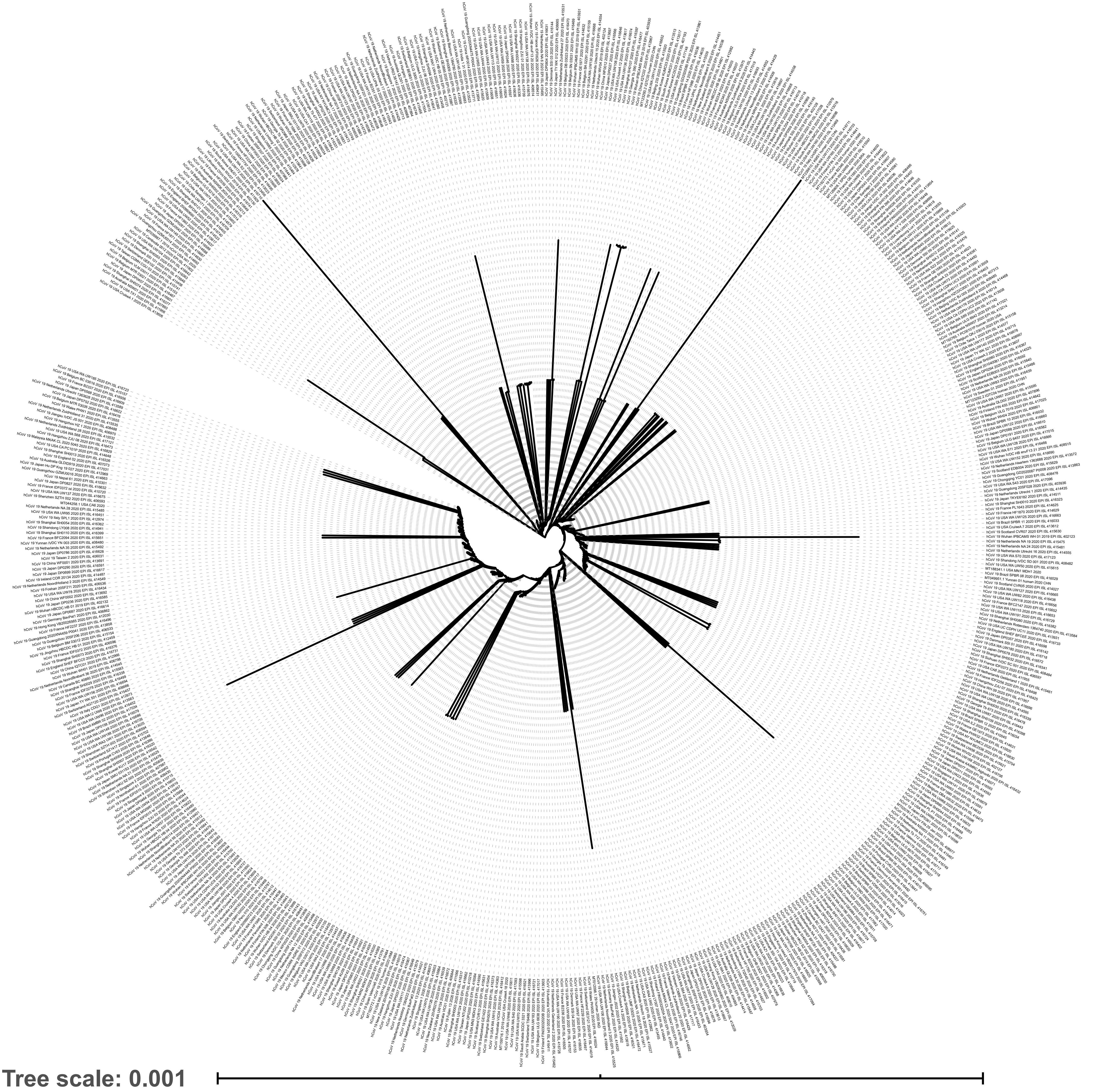

c)

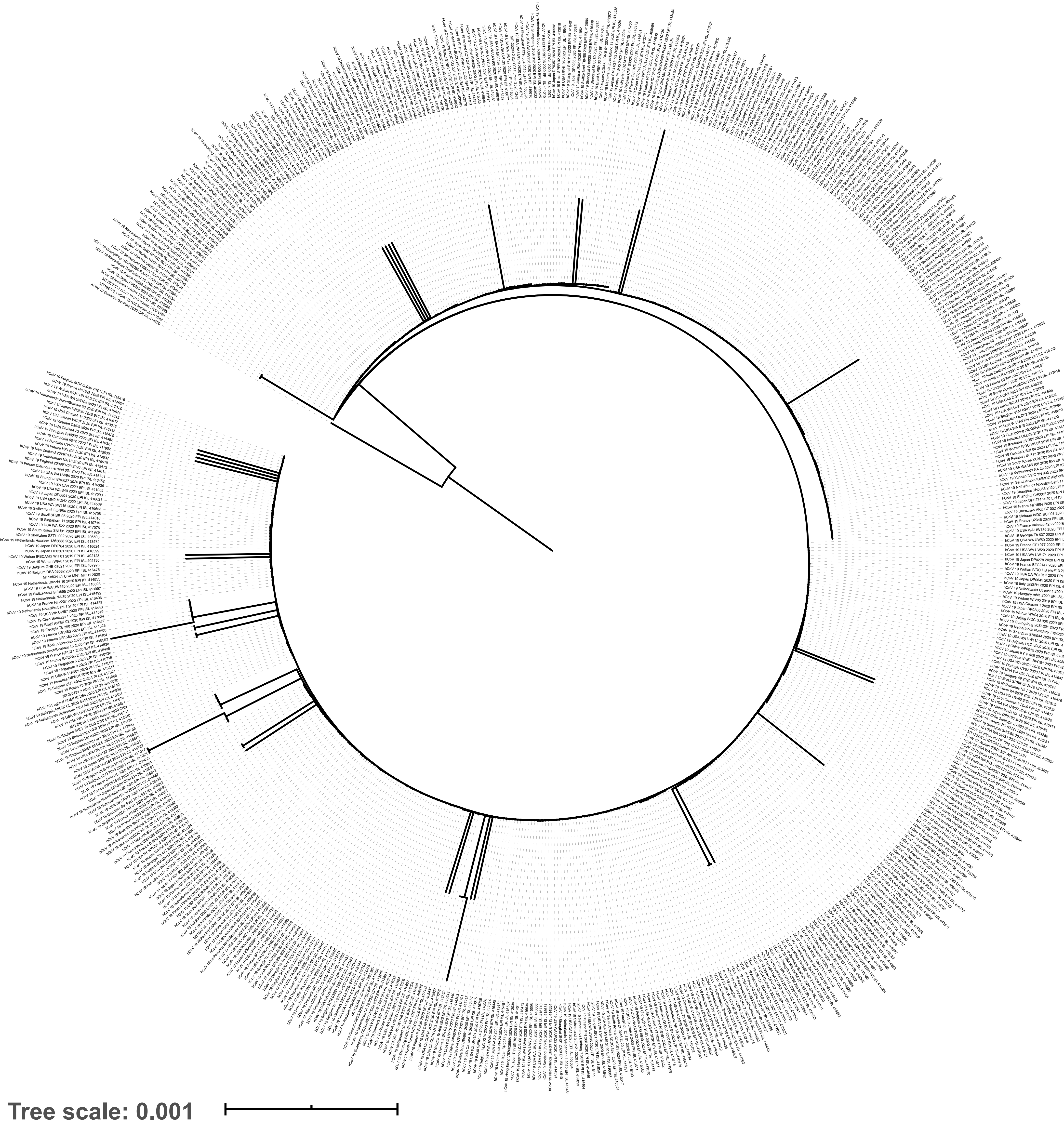

d)

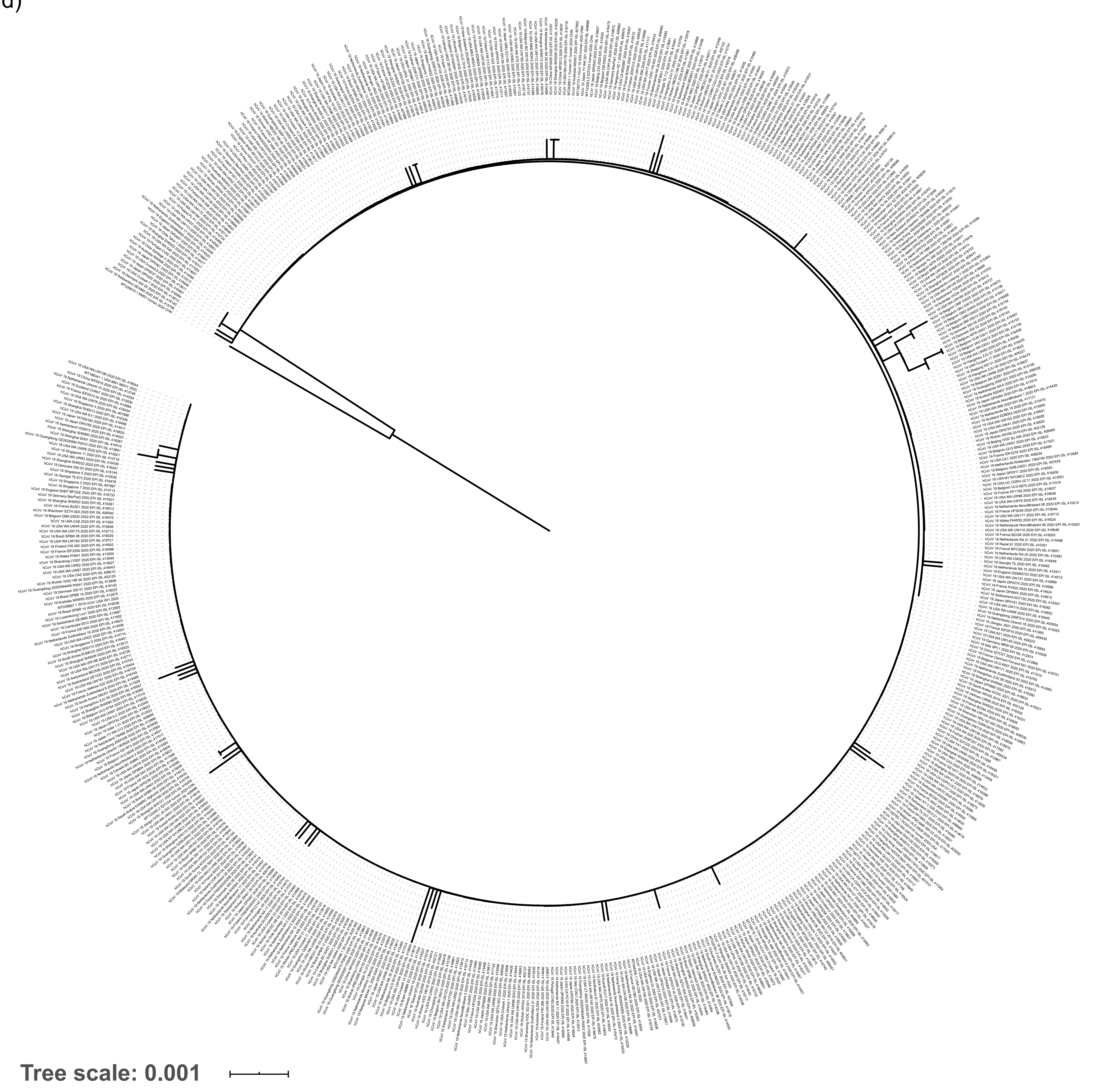

e)

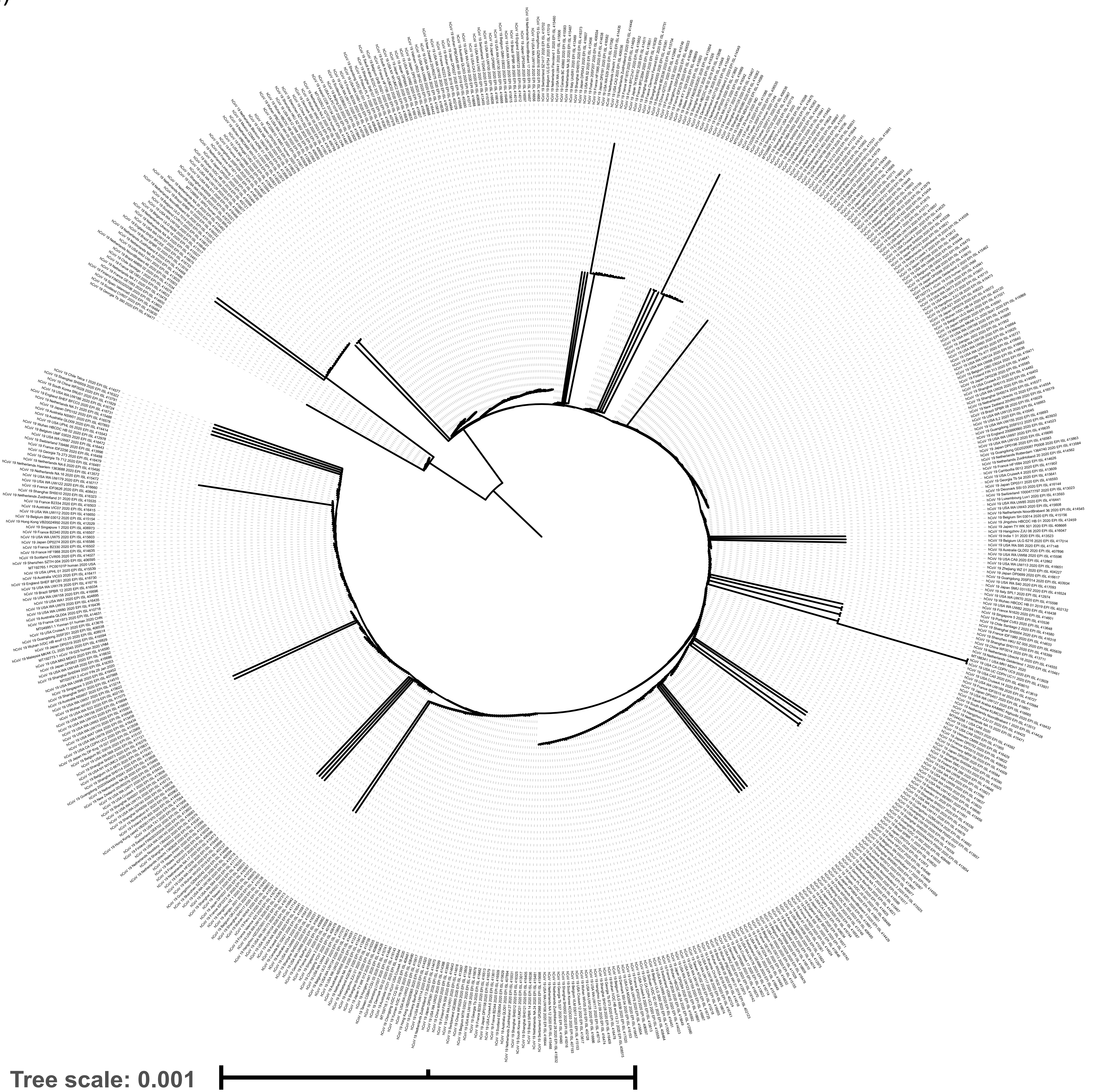

f)

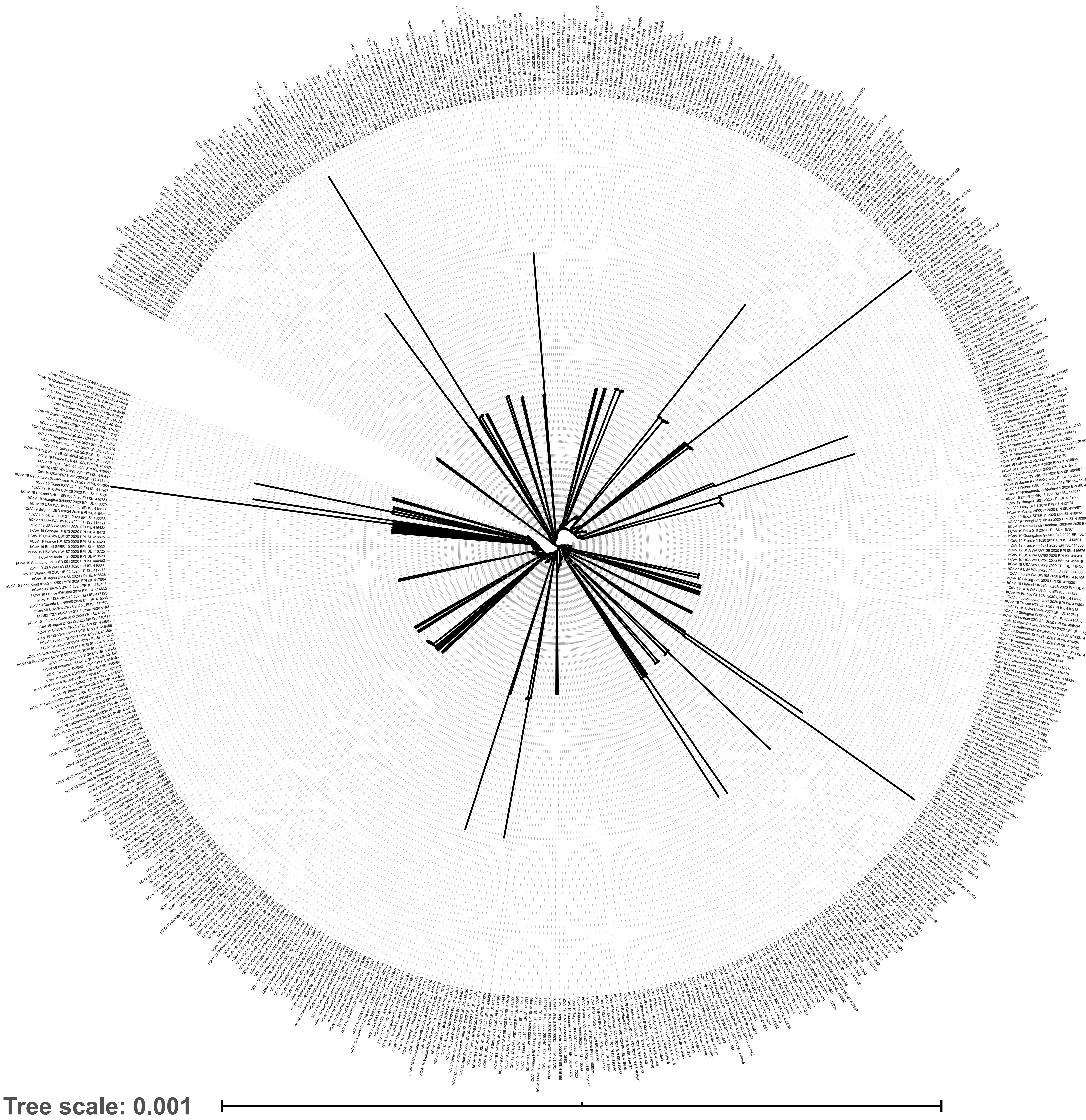

g)

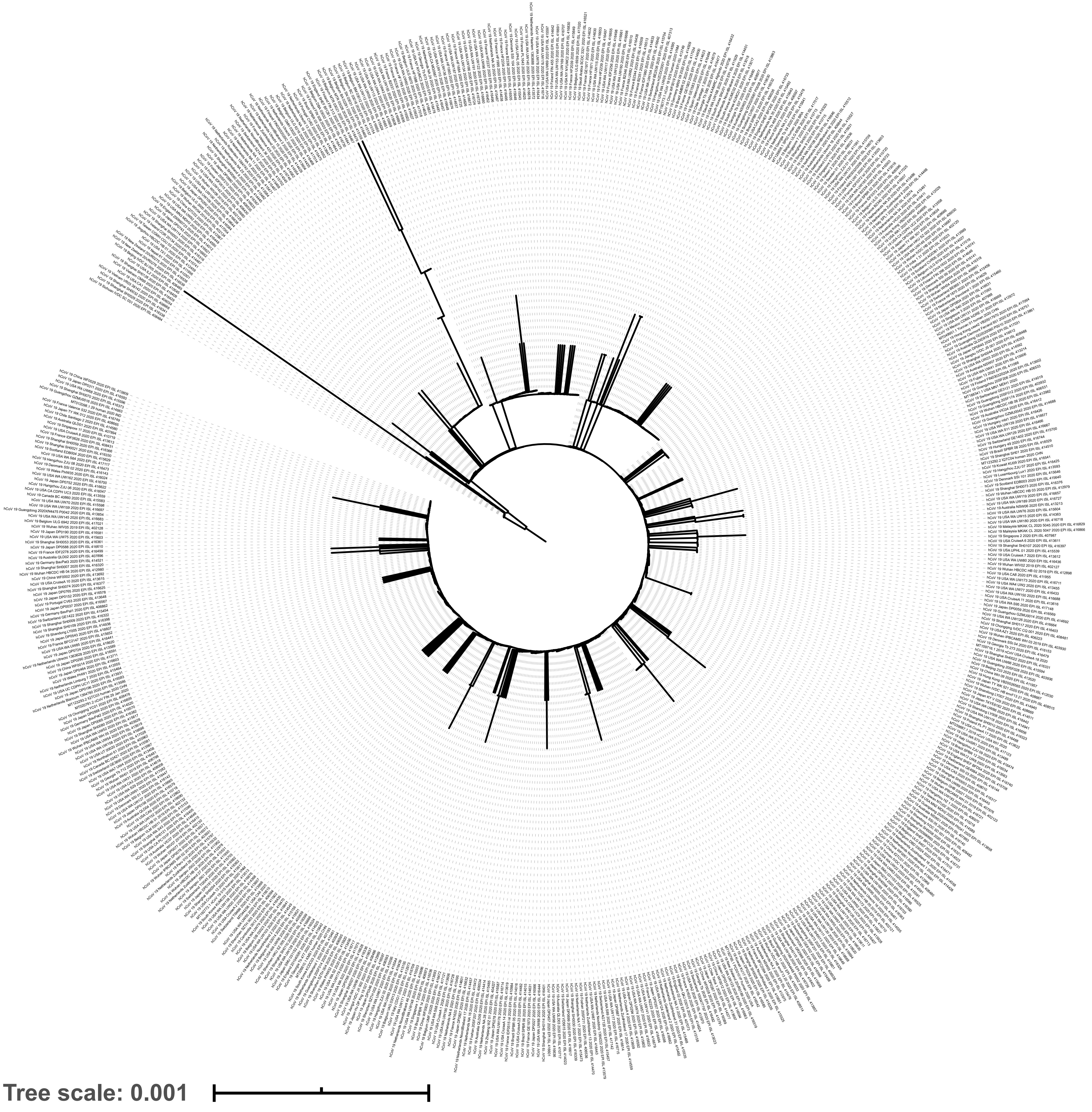

h)

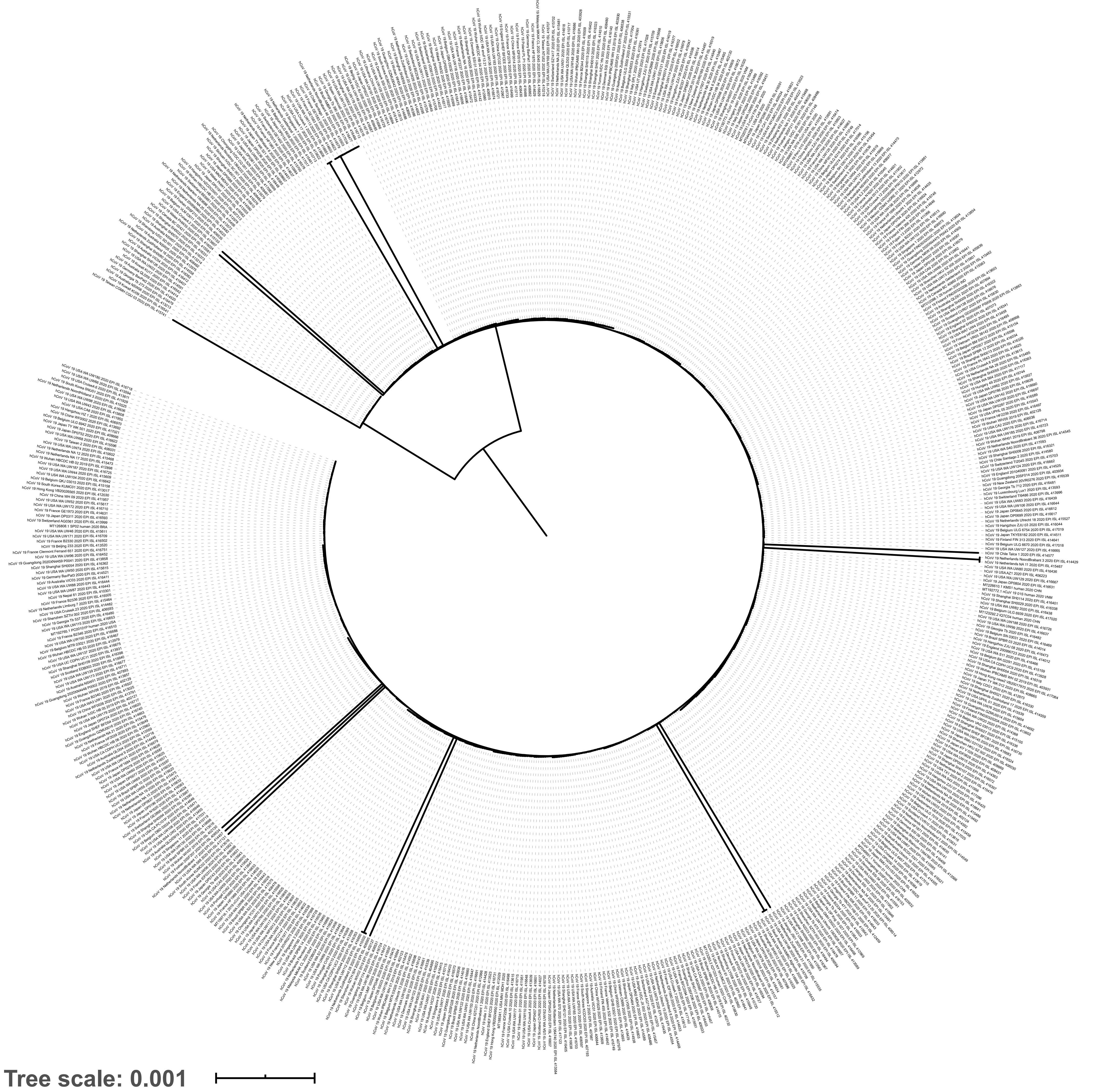

i)

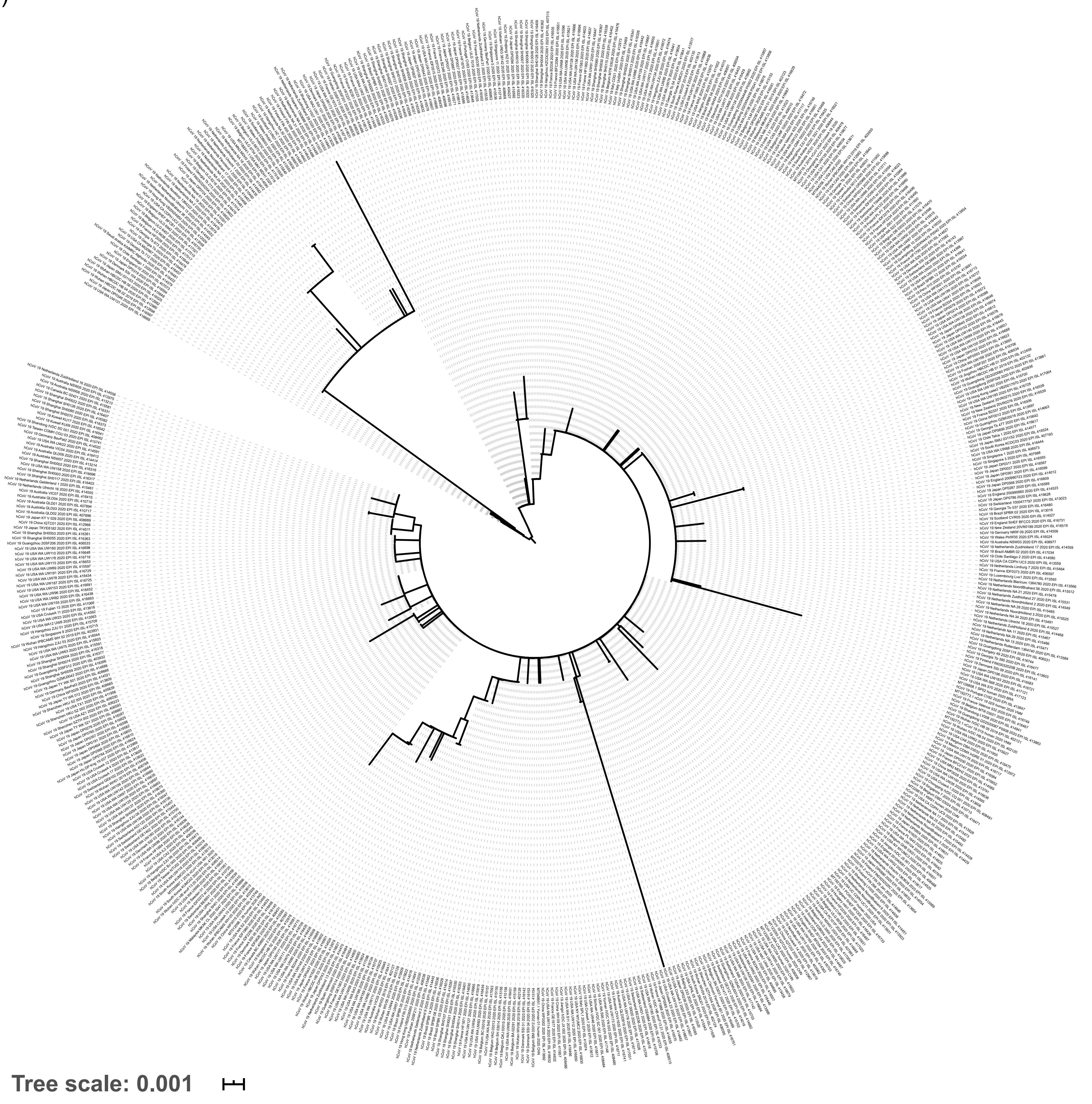

### Supplementary Information

#### The global population of SARS-CoV-2 is composed of six major subtypes

Ivair José Morais Júnior, Richard Costa Polveiro, Gabriel Medeiros Souza, Daniel Inserra Bortolin, Flávio Tetsuo Sassaki, Alison Talis Martins Lima

**Supplementary Table S1** | Genomes of SARS-CoV-2 used in this study.
Genomes of SARS-CoV-2 were retrieved from GISAID[1] and GenBank[2] on 25 March, 2020. We applied a conservative filter for high quality sequences sampled from humans by excluding those ones with indeterminate nucleotides ('N's or any other ambiguous code) and non-intact ORFs (no frameshifts, except that of *nsp*12 cistron). After filtering for high quality sequences, a total of 767 SARS-CoV-2 genomes were submitted to haplotype detection analysis in DnaSP v.6[3] and a single representative of each haplotype was used in study. Additional sequences from the same haplotype are indicated in red. The haplotypes 4, 37, 119 and 443 (31, 9, 7 and 6 genomes each, respectively) and 57, 168, 247, 318 (5 genomes, each) were the most represented in our data set. Therefore, after removing the redundant sequences, the data set totalized 593 unique SARS-CoV-2 genomes. We wish to acknowledge all researchers that deposited the SARS-CoV-2 genomes in GISAID and/or GenBank databases.

**Supplementary Table S2** | Genotypes of 593 SARS-CoV-2 genomes based on 12 widely shared polymorphisms (WSPs) detected in this study.
For each virus isolate is shown the nucleotide at the WSPs: *nsp*3-[318], *nsp*4-[228], *nsp*6-[111], *nsp*12-[967], *nsp*13-[1511], *nsp*13-[1622], *nsp*14-[21], *S*-[1,841], *ORF8*-[251], *N*-[608], *N*-[609] and *N*-[610]. Nucleotide bases A, C, G and U are indicated by colored cells in green, blue, purple and red.

| Haplotype ID | Accession ID | Virus name | Host | Location | Collection date | Originating lab | Submitting lab | Authors |
|---|---|---|---|---|---|---|---|---|
| 1 | EPI_ISL_402120 | hCoV-19/Wuhan/IVDC-HB-04/2020 | *Homo sapiens* | Asia / China / Hubei / Wuhan | 2020-01-01 | National Institute for Viral Disease Control and Prevention, China CDC | National Institute for Viral Disease Control and Prevention, China CDC | Wenjie Tan, Xiang Zhao, Wenling Wang, Xuejun Ma, Yongzhong Jiang, Roujian Lu, Ji Wang, Weimin Zhou, Peihua Niu, Peipei Liu, Faxian Zhan, Weifeng Shi, Baoying Huang, Jun Liu, Li Zhao, Yao Meng, Xiaozhou He, Fei Ye, Na Zhu, Yang Li, Jing Chen, Wenbo Xu, George F. Gao, Guizhen Wu |
| 2 | EPI_ISL_402121 | hCoV-19/Wuhan/IVDC-HB-05/2019 | *Homo sapiens* | Asia / China / Hubei / Wuhan | 2019-12-30 | National Institute for Viral Disease Control and Prevention, China CDC | National Institute for Viral Disease Control and Prevention, China CDC | Wenjie Tan, Xuejun Ma, Xiang Zhao, Wenling Wang, Yongzhong Jiang, Roujian Lu, Ji Wang, Peihua Niu, Weimin Zhou, Faxian Zhan, Weifeng Shi, Baoying Huang, Jun Liu, Li Zhao, Yao Meng, Fei Ye, Na Zhu, Xiaozhou He, Peipei Liu, Yang Li, Jing Chen, Wenbo Xu, George F. Gao, Guizhen Wu |
| 3 | EPI_ISL_402123 | hCoV-19/Wuhan/IPBCAMS-WH-01/2019 | *Homo sapiens* | Asia / China / Hubei / Wuhan | 2019-12-24 | Institute of Pathogen Biology, Chinese Academy of Medical Sciences & Peking Union Medical College | Institute of Pathogen Biology, Chinese Academy of Medical Sciences & Peking Union Medical College | Lili Ren, Jianwei Wang, Qi Jin, Zichun Xiang, Zhiqiang Wu, Chao Wu, Yiwei Liu |
| 4 | EPI_ISL_402124 | hCoV-19/Wuhan/WIV04/2019 | *Homo sapiens* | Asia / China / Hubei / Wuhan | 2019-12-30 | Wuhan Jinyintan Hospital | Wuhan Institute of Virology, Chinese Academy of Sciences | Peng Zhou, Xing-Lou Yang, Ding-Yu Zhang, Lei Zhang, Yan Zhu, Hao-Rui Si, Zhengli Shi |
| 4 | EPI_ISL_402125 | hCoV-19/Wuhan-Hu-1/2019 | *Homo sapiens* | Asia / China | 2019-12-31 | unknown | National Institute for Communicable Disease Control and Prevention (ICDC) Chinese Center for Disease Control and Prevention (China CDC) | Zhang,Y.-Z., Wu,F., Chen,Y.-M., Pei,Y.-Y., Xu,L., Wang,W., Zhao,S., Yu,B., Hu,Y., Tao,Z.-W., Song,Z.-G., Tian,J.-H., Zhang,Y.-L., Liu,Y., Zheng,J.-J., Dai,F.-H., Wang,Q.-M., She,J.-L. and Zhu,T.-Y. |
| 4 | EPI_ISL_403929 | hCoV-19/Wuhan/IPBCAMS-WH-04/2019 | *Homo sapiens* | Asia / China / Hubei / Wuhan | 2019-12-30 | Institute of Pathogen Biology, Chinese Academy of Medical Sciences & Peking Union Medical College | Institute of Pathogen Biology, Chinese Academy of Medical Sciences & Peking Union Medical College | Lili Ren, Jianwei Wang, Qi Jin, Zichun Xiang, Zhiqiang Wu, Chao Wu, Yiwei Liu |
| 4 | EPI_ISL_403963 | hCoV-19/Nonthaburi/74/2020 | *Homo sapiens* | Asia / Thailand / Nonthaburi | 2020-01-13 | Bamrasnaradura Hospital | 1. Department of Medical Sciences, Ministry of Public Health, Thailand 2. Thai Red Cross Emerging Infectious Diseases - Health Science Centre 3. Department of Disease Control, Ministry of Public Health, Thailand | Pilailuk,Okada; Siripaporn,Phuygun; Thanutsapa,Thanadachakul; Supaporn,Wacharapluesadee; Sittiporn,Parnmen; Warawan,Wongboot; Sunthareeya,Waicharoen; Rome,Buathong; Malinee,Chittaganpitch; Nanthawan,Mekha |
| 4 | EPI_ISL_404228 | hCoV-19/Zhejiang/WZ-02/2020 | *Homo sapiens* | Asia / China / Zhejiang | 2020-01-17 | Zhejiang Provincial Center for Disease Control and Prevention | Department of Microbiology, Zhejiang Provincial Center for Disease Control and Prevention | Yanjun Zhang, Yin Chen, Haiyan Mao, Junhang Pan, Xiuyu Lou, Yiyu Lu, Juying Yan, Hanping Zhu, Jian Gao, Yan Feng, Yi Sun, Hao Yan, Zhen Li, Yisheng Sun, Liming Gong, Qiong Ge, Wen Shi, Xinying Wang, Wenwu Yao, Zhangnv Yang, Fang Xu, Chen Chen, Enfu Chen, Zhen Wang, Zhiping Chen, Jianmin Jiang, Chonggao Hu |
| 4 | EPI_ISL_406716 | hCoV-19/China/WHU01/2020 | *Homo sapiens* | Asia / China / Hubei / Wuhan | 2020-01-02 | unknown | State Key Laboratory of Virology, Wuhan University | Chen,L., Liu,W., Zhang,Q., Xu,K., Ye,G., Wu,W., Sun,Z., Liu,F., Wu,K., Mei,Y., Zhang,W., Chen,Y., Li,Y., Shi,M., Lan,K. and Liu,Y. |
| 4 | EPI_ISL_406717 | hCoV-19/China/WHU02/2020 | *Homo sapiens* | Asia / China / Hubei / Wuhan | 2020-01-02 | unknown | State Key Laboratory of Virology, Wuhan University | Chen,L., Liu,W., Zhang,Q., Xu,K., Ye,G., Wu,W., Sun,Z., Liu,F., Wu,K., Mei,Y., Zhang,W., Chen,Y., Li,Y., Shi,M., Lan,K. and Liu,Y. |
| 4 | EPI_ISL_406800 | hCoV-19/Wuhan/WH03/2020 | *Homo sapiens* | Asia / China / Hubei / Wuhan | 2020-01-01 | General Hospital of Central Theater Command of People's Liberation Army of China | BGI & Institute of Microbiology, Chinese Academy of Sciences & Shandong First Medical University & Shandong Academy of Medical Sciences & General Hospital of Central Theater Command of People's Liberation Army of China | Weijun Chen, Yuhai Bi, Weifeng Shi and Zhenhong Hu |
| 4 | EPI_ISL_408479 | hCoV-19/Chongqing/ZX01/2020 | *Homo sapiens* | Asia / China / Chongqing / Zhongxian | 2020-01-23 | Zhongxian Center for Disease Control and Prevention | Chongqing Municipal Center for Disease Control and Prevention | Ye Sheng, Tang Yun, Ling Hua, Zhang Hong, Yu zhen,Chen Shuang,Tan ZhangPing, Su Kun, Li Qin, Tang Wenge, Rong Rong |
| 4 | EPI_ISL_410531 | hCoV-19/Japan/NA-20-05-1/2020 | *Homo sapiens* | Asia / Japan / Nara | 2020-01-25 | Dept. of Pathology, National Institute of Infectious Diseases | Pathogen Genomics Center, National Institute of Infectious Diseases | Tsuyoshi Sekizuka, Harutaka Katano, Shutoku Matsuyama, Naganori Nao, Kazuya Shirato, Motoi Suzuki, Hideki Hasegawa, Takaji Wakita, Makoto Takeda, Tadaki Suzuki, Makoto Kuroda |

| | | | | | | | | |
|---|---|---|---|---|---|---|---|---|
| 4 | EPI_ISL_410532 | hCoV-19/Japan/OS-20-07-1/2020 | *Homo sapiens* | Asia / Japan / Osaka | 2020-01-23 | Dept. of Pathology, National Institute of Infectious Diseases | Pathogen Genomics Center, National Institute of Infectious Diseases | Tsuyoshi Sekizuka, Harutaka Katano, Shutoku Matsuyama, Naganori Nao, Kazuya Shirato, Motoi Suzuki, Hideki Hasegawa, Takaji Wakita, Makoto Takeda, Tadaki Suzuki, Makoto Kuroda |
| 4 | EPI_ISL_411915 | hCoV-19/Taiwan/CGMH-CGU-01/2020 | *Homo sapiens* | Asia / Taiwan / Taoyuan | 2020-01-25 | Laboratory Medicine | Department of Laboratory Medicine, Lin-Kou Chang Gung Memorial Hospital, Taoyuan, Taiwan. | Kuo-Chien Tsao, Yu-Nong Gong, Shu-Li Yang, Yi-Chun Li, Chung-Guei Huang, Yhu-Chering Huang, Shin-Ru Shih |
| 4 | EPI_ISL_411927 | hCoV-19/Taiwan/4/2020 | *Homo sapiens* | Asia / Taiwan / Taipei | 2020-01-28 | Taiwan Centers for Disease Control | Taiwan Centers for Disease Control | Ji-Rong Yang, Yu-Chi-Lin, Jung-Jung Mu, Ming-Tsan-Liu |
| 4 | EPI_ISL_411953 | hCoV-19/Jiangsu/JS03/2020 | *Homo sapiens* | Asia / China / Jiangsu | 2020-01-24 | NHC Key laboratory of Enteric Pathogenic Microbiology, Institute of Pathogenic Microbiology | Jiangsu Provincial Center for Disease Control & Prevention | Kangchen Zhao, Xiaojuan Zhu, Lunbiao Cui, Tao Wu, Yiyue Ge, Bin Wu, Yin Chen, Fengcai Zhu, Baoli Zhu, Ming Wu |
| 4 | EPI_ISL_412026 | hCoV-19/Hefei/2/2020 | *Homo sapiens* | Asia / China / Anhui / Hefei | 2020-02-23 | Second Hospital of Anhui Medical University | Second Hospital of Anhui Medical University | Changtai Wang, Zhongping Liua, Zixiang Chen, Xin Huang, Mengyuan Xua, Tengfei He, Mengji Lu, Zhenhua Zhang |
| 4 | EPI_ISL_413608 | hCoV-19/USA/CruiseA-3/2020 | *Homo sapiens* | North America / USA | 2020-02-18 | unknown | Pathogen Discovery, Respiratory Viruses Branch, Division of Viral Diseases, Centers for Dieases Control and Prevention | Anna Uehara, Ying Tao, Clinton R. Paden, Krista Queen, Jing Zhang, Yan Li, Mary S. Keckler, Alison S Laufer Halpin, Haibin Wang, Jasmine Padilla, Justin Lee, Christopher A. Elkins, Susan I. Gerber, Suxiang Tong |
| 4 | EPI_ISL_413610 | hCoV-19/USA/CruiseA-5/2020 | *Homo sapiens* | North America / USA | 2020-02-21 | unknown | Pathogen Discovery, Respiratory Viruses Branch, Division of Viral Diseases, Centers for Dieases Control and Prevention | Anna Uehara, Ying Tao, Clinton R. Paden, Krista Queen, Jing Zhang, Yan Li, Mary S. Keckler, Alison S Laufer Halpin, Haibin Wang, Jasmine Padilla, Justin Lee, Christopher A. Elkins, Susan I. Gerber, Suxiang Tong |
| 4 | EPI_ISL_413614 | hCoV-19/USA/CruiseA-9/2020 | *Homo sapiens* | North America / USA | 2020-02-17 | unknown | Pathogen Discovery, Respiratory Viruses Branch, Division of Viral Diseases, Centers for Dieases Control and Prevention | Ying Tao, Clinton R. Paden, Krista Queen, Anna Uehara, Jing Zhang, Yan Li, Haibin Wang, Shifaq Kamili, Xiaoyan Lu, Brian Lynch, Senthil Kumar K. Sakthivel, Brett L. Whitaker, Lijuan Wang, Janna' R. Murray, Jasmine Padilla, Justin Lee, Susan I. Gerber, Stephen Lindstrom, Suxiang Tong |
| 4 | EPI_ISL_413618 | hCoV-19/USA/CruiseA-13/2020 | *Homo sapiens* | North America / USA | 2020-02-20 | unknown | Pathogen Discovery, Respiratory Viruses Branch, Division of Viral Diseases, Centers for Dieases Control and Prevention | Clinton R. Paden, Ying Tao, Krista Queen, Anna Uehara, Jing Zhang, Yan Li, Haibin Wang, Shifaq Kamili, Xiaoyan Lu, Brian Lynch, Senthil Kumar K. Sakthivel, Brett L. Whitaker, Lijuan Wang, Janna' R. Murray, Jasmine Padilla, Justin Lee, Susan I. Gerber, Stephen Lindstrom, Suxiang Tong |
| 4 | EPI_ISL_413620 | hCoV-19/USA/CruiseA-15/2020 | *Homo sapiens* | North America / USA | 2020-02-18 | unknown | Pathogen Discovery, Respiratory Viruses Branch, Division of Viral Diseases, Centers for Dieases Control and Prevention | Clinton R. Paden, Ying Tao, Krista Queen, Anna Uehara, Jing Zhang, Yan Li, Haibin Wang, Shifaq Kamili, Xiaoyan Lu, Brian Lynch, Senthil Kumar K. Sakthivel, Brett L. Whitaker, Lijuan Wang, Janna' R. Murray, Jasmine Padilla, Justin Lee, Susan I. Gerber, Stephen Lindstrom, Suxiang Tong |
| 4 | EPI_ISL_413621 | hCoV-19/USA/CruiseA-16/2020 | *Homo sapiens* | North America / USA | 2020-02-18 | unknown | Pathogen Discovery, Respiratory Viruses Branch, Division of Viral Diseases, Centers for Dieases Control and Prevention | Clinton R. Paden, Ying Tao, Krista Queen, Anna Uehara, Jing Zhang, Yan Li, Haibin Wang, Shifaq Kamili, Xiaoyan Lu, Brian Lynch, Senthil Kumar K. Sakthivel, Brett L. Whitaker, Lijuan Wang, Janna' R. Murray, Jasmine Padilla, Justin Lee, Susan I. Gerber, Stephen Lindstrom, Suxiang Tong |
| 4 | EPI_ISL_414479 | hCoV-19/USA/CruiseA-19/2020 | *Homo sapiens* | North America / USA | 2020-02-18 | unknown | Pathogen Discovery, Respiratory Viruses Branch, Division of Viral Diseases, Centers for Disease Control and Prevention | Ying Tao, Krista Queen, Clinton R. Paden, Anna Uehara, Jing Zhang, Yan Li, Mary S. Keckler, Alison S. Laufer Halpin, Haibin Wang, Jasmine Padilla, Justin Lee, Christopher A. Elkins, Susan I. Gerber, Suxiang Tong |
| 4 | EPI_ISL_414481 | hCoV-19/USA/CruiseA-22/2020 | *Homo sapiens* | North America / USA | 2020-02-21 | unknown | Pathogen Discovery, Respiratory Viruses Branch, Division of Viral Diseases, Centers for Disease Control and Prevention | Ying Tao, Krista Queen, Clinton R. Paden, Anna Uehara, Jing Zhang, Yan Li, Mary S. Keckler, Alison S. Laufer Halpin, Haibin Wang, Jasmine Padilla, Justin Lee, Christopher A. Elkins, Susan I. Gerber, Suxiang Tong |

| | | | | | | | | |
|---|---|---|---|---|---|---|---|---|
| 4 | EPI_ISL_415711 | hCoV-19/Hangzhou/ZJU-05/2020 | *Homo sapiens* | Asia / China / Hangzhou | 2020-01-22 | State Key Laboratory for Diagnosis and Treatment of Infectious Diseases, National Clinical Research Center for Infectious Diseases, First Affiliated Hospital, Zhejiang University School of Medicine, Hangzhou, China. 310003 | State Key Laboratory for Diagnosis and Treatment of Infectious Diseases, National Clinical Research Center for Infectious Diseases, First Affiliated Hospital, Zhejiang University School of Medicine, Hangzhou, China. 310003 | Hangping Yao, Nanping Wu, Chao Jiang, Xiangyun Lu, Linfang Cheng, Fumin Liu, Zhigang Wu, Haibo Wu, Changzhong Jin, Min Zheng, Lanjuan Li |
| 4 | EPI_ISL_416046 | hCoV-19/Hangzhou/ZJU-04/2020 | *Homo sapiens* | Asia / China / Hangzhou | 2020-01-24 | State Key Laboratory for Diagnosis and Treatment of Infectious Diseases, National Clinical Research Center for Infectious Diseases, First Affiliated Hospital, Zhejiang University School of Medicine, Hangzhou, China 310003 | State Key Laboratory for Diagnosis and Treatment of Infectious Diseases, National Clinical Research Center for Infectious Diseases, First Affiliated Hospital, Zhejiang University School of Medicine, Hangzhou, China 310003 | Hangping Yao, Nanping Wu, Chao Jiang, Xiangyun Lu, Linfang Cheng, Fumin Liu, Zhigang Wu, Haibo Wu, Changzhong Jin, Min Zheng, Lanjuan Li |
| 4 | MT192759 | SARS-CoV-2/CGMH-CGU-01/Homo sapiens2020/TWN | *Homo sapiens* | Taiwan | 2020-01-25 | not specified | not specified | Tsao,K.-C., Gong,Y.-N., Yang,S.-L., Liu,Y.-C., Huang,C.-G., Huang,Y.-C., Chen,G.-W. and Shih,S.-R. |
| 4 | NC_045512 | Wuhan-Hu-1 | *Homo sapiens* | China | 2019-12 | not specified | not specified | Wu,F., Zhao,S., Yu,B., Chen,Y.-M., Wang,W., Hu,Y., Song,Z.-G., Tao,Z.-W., Tian,J.-H., Pei,Y.-Y., Yuan,M.L., Zhang,Y.-L., Dai,F.-H., Liu,Y., Wang,Q.-M., Zheng,J.-J., Xu,L., Holmes,E.C. and Zhang,Y.-Z. |
| 4 | MT184909 | 2019-nCoV/USA-CruiseA-22/2020 | *Homo sapiens* | USA | 2020-02-21 | not specified | not specified | Tao,Y., Queen,K., Paden,C.R., Uehara,A., Zhang,J., Li,Y., Keckler,M.S., Laufer Halpin,A.S., Wang,H., Padilla,J., Lee,J., Elkins,C.A., Gerber,S.I. and Tong,S. |
| 4 | MT184907 | 2019-nCoV/USA-CruiseA-19/2020 | *Homo sapiens* | USA | 2020-02-18 | not specified | not specified | Tao,Y., Queen,K., Paden,C.R., Uehara,A., Zhang,J., Li,Y., Keckler,M.S., Laufer Halpin,A.S., Wang,H., Padilla,J., Lee,J., Elkins,C.A., Gerber,S.I. and Tong,S. |
| 4 | LR757996 | SARS-CoV-2 | *Homo sapiens* | China:Wuhan | 2020-01-01 | not specified | not specified | Hunter,C. and Wei,X. |
| 4 | MN988669 | 2019-nCoV WHU02 | *Homo sapiens* | China | 2020-01-02 | not specified | not specified | Chen,L., Liu,W., Zhang,Q., Xu,K., Ye,G., Wu,W., Sun,Z., Liu,F., Wu,K., Mei,Y., Zhang,W., Chen,Y., Li,Y., Shi,M., Lan,K. and Liu,Y. |
| 5 | EPI_ISL_402127 | hCoV-19/Wuhan/WIV02/2019 | *Homo sapiens* | Asia / China / Hubei / Wuhan | 2019-12-30 | Wuhan Jinyintan Hospital | Wuhan Institute of Virology, Chinese Academy of Sciences | Peng Zhou, Xing-Lou Yang, Ding-Yu Zhang, Lei Zhang, Yan Zhu, Hao-Rui Si, Zhengli Shi |
| 6 | EPI_ISL_402128 | hCoV-19/Wuhan/WIV05/2019 | *Homo sapiens* | Asia / China / Hubei / Wuhan | 2019-12-30 | Wuhan Jinyintan Hospital | Wuhan Institute of Virology, Chinese Academy of Sciences | Peng Zhou, Xing-Lou Yang, Ding-Yu Zhang, Lei Zhang, Yan Zhu, Hao-Rui Si, Zhengli Shi |
| 7 | EPI_ISL_402129 | hCoV-19/Wuhan/WIV06/2019 | *Homo sapiens* | Asia / China / Hubei / Wuhan | 2019-12-30 | Wuhan Jinyintan Hospital | Wuhan Institute of Virology, Chinese Academy of Sciences | Peng Zhou, Xing-Lou Yang, Ding-Yu Zhang, Lei Zhang, Yan Zhu, Hao-Rui Si, Zhengli Shi |
| 8 | EPI_ISL_402130 | hCoV-19/Wuhan/WIV07/2019 | *Homo sapiens* | Asia / China / Hubei / Wuhan | 2019-12-30 | Wuhan Jinyintan Hospital | Wuhan Institute of Virology, Chinese Academy of Sciences | Peng Zhou, Xing-Lou Yang, Ding-Yu Zhang, Lei Zhang, Yan Zhu, Hao-Rui Si, Zhengli Shi |
| 9 | EPI_ISL_402132 | hCoV-19/Wuhan/HBCDC-HB-01/2019 | *Homo sapiens* | Asia / China / Hubei / Wuhan | 2019-12-30 | Wuhan Jinyintan Hospital | Hubei Provincial Center for Disease Control and Prevention | Bin Fang, Xiang Li, Xiao Yu, Linlin Liu, Bo Yang, Faxian Zhan, Guojun Ye, Xixiang Huo, Junqiang Xu, Bo Yu, Kun Cai, Jing Li, Yongzhong Jiang. |
| 10 | EPI_ISL_403928 | hCoV-19/Wuhan/IPBCAMS-WH-05/2020 | *Homo sapiens* | Asia / China / Hubei / Wuhan | 2020-01-01 | Institute of Pathogen Biology, Chinese Academy of Medical Sciences & Peking Union Medical College | Institute of Pathogen Biology, Chinese Academy of Medical Sciences & Peking Union Medical College | Lili Ren, Jianwei Wang, Qi Jin, Zichun Xiang, Zhiqiang Wu, Chao Wu, Yiwei Liu |
| 11 | EPI_ISL_403930 | hCoV-19/Wuhan/IPBCAMS-WH-03/2019 | *Homo sapiens* | Asia / China / Hubei / Wuhan | 2019-12-30 | Institute of Pathogen Biology, Chinese Academy of Medical Sciences & Peking Union Medical College | Institute of Pathogen Biology, Chinese Academy of Medical Sciences & Peking Union Medical College | Lili Ren, Jianwei Wang, Qi Jin, Zichun Xiang, Zhiqiang Wu, Chao Wu, Yiwei Liu |

| | | | | | | | | |
|---|---|---|---|---|---|---|---|---|
| 12 | EPI_ISL_403931 | hCoV-19/Wuhan/IPBCAMS-WH-02/2019 | *Homo sapiens* | Asia / China / Hubei / Wuhan | 2019-12-30 | Institute of Pathogen Biology, Chinese Academy of Medical Sciences & Peking Union Medical College | Institute of Pathogen Biology, Chinese Academy of Medical Sciences & Peking Union Medical College | Lili Ren, Jianwei Wang, Qi Jin, Zichun Xiang, Zhiqiang Wu, Chao Wu, Yiwei Liu |
| 13 | EPI_ISL_403932 | hCoV-19/Guangdong/20SF012/2020 | *Homo sapiens* | Asia / China / Guandong / Shenzhen | 2020-01-14 | Guangdong Provincial Center for Diseases Control and Prevention; Guangdong Provincial Public Health | Department of Microbiology, Guangdong Provincial Center for Diseases Control and Prevention | Min Kang, Jie Wu, Jing Lu, Tao Liu, Baisheng Li, Shujiang Mei, Feng Ruan, Lifeng Lin, Changwen Ke, Haojie Zhong, Yingtao Zhang, Lirong Zou, Xuguang Chen, Qi Zhu, Jianpeng Xiao, Jianxiang Geng, Zhe Liu, Jianxiong Hu, Weilin Zeng, Xing Li, Yuhuang Liao, Xiujuan Tang, Songjian Xiao, Ying Wang, Yingchao Song, Xue Zhuang, Lijun Liang, Guanhao He, Huihong Deng, Tie Song, Jianfeng He, Wenjun Ma |
| 13 | EPI_ISL_403933 | hCoV-19/Guangdong/20SF013/2020 | *Homo sapiens* | Asia / China / Guandong / Shenzhen | 2020-01-15 | Guangdong Provincial Center for Diseases Control and Prevention; Guangdong Provincial Public Health | Department of Microbiology, Guangdong Provincial Center for Diseases Control and Prevention | Min Kang, Jie Wu, Jing Lu, Tao Liu, Baisheng Li, Shujiang Mei, Feng Ruan, Lifeng Lin, Changwen Ke, Haojie Zhong, Yingtao Zhang, Lirong Zou, Xuguang Chen, Qi Zhu, Jianpeng Xiao, Jianxiang Geng, Zhe Liu, Jianxiong Hu, Weilin Zeng, Xing Li, Yuhuang Liao, Xiujuan Tang, Songjian Xiao, Ying Wang, Yingchao Song, Xue Zhuang, Lijun Liang, Guanhao He, Huihong Deng, Tie Song, Jianfeng He, Wenjun Ma |
| 13 | EPI_ISL_403935 | hCoV-19/Guangdong/20SF025/2020 | *Homo sapiens* | Asia / China / Guangdong / Shenzhen | 2020-01-15 | Guangdong Provincial Center for Diseases Control and Prevention; Guangdong Provincial Public Health | Department of Microbiology, Guangdong Provincial Center for Diseases Control and Prevention | Min Kang, Jie Wu, Jing Lu, Tao Liu, Baisheng Li, Shujiang Mei, Feng Ruan, Lifeng Lin, Changwen Ke, Haojie Zhong, Yingtao Zhang, Lirong Zou, Xuguang Chen, Qi Zhu, Jianpeng Xiao, Jianxiang Geng, Zhe Liu, Jianxiong Hu, Weilin Zeng, Xing Li, Yuhuang Liao, Xiujuan Tang, Songjian Xiao, Ying Wang, Yingchao Song, Xue Zhuang, Lijun Liang, Guanhao He, Huihong Deng, Tie Song, Jianfeng He, Wenjun Ma |
| 14 | EPI_ISL_403934 | hCoV-19/Guangdong/20SF014/2020 | *Homo sapiens* | Asia / China / Guandong / Shenzhen | 2020-01-15 | Guangdong Provincial Center for Diseases Control and Prevention; Guangdong Provincial Public Health | Department of Microbiology, Guangdong Provincial Center for Diseases Control and Prevention | Min Kang, Jie Wu, Jing Lu, Tao Liu, Baisheng Li, Shujiang Mei, Feng Ruan, Lifeng Lin, Changwen Ke, Haojie Zhong, Yingtao Zhang, Lirong Zou, Xuguang Chen, Qi Zhu, Jianpeng Xiao, Jianxiang Geng, Zhe Liu, Jianxiong Hu, Weilin Zeng, Xing Li, Yuhuang Liao, Xiujuan Tang, Songjian Xiao, Ying Wang, Yingchao Song, Xue Zhuang, Lijun Liang, Guanhao He, Huihong Deng, Tie Song, Jianfeng He, Wenjun Ma |
| 15 | EPI_ISL_403936 | hCoV-19/Guangdong/20SF028/2020 | *Homo sapiens* | Asia / China / Guangdong / Zhuhai | 2020-01-17 | Guangdong Provincial Center for Diseases Control and Prevention; Guangdong Provincial Public Health | Department of Microbiology, Guangdong Provincial Center for Diseases Control and Prevention | Min Kang, Jie Wu, Jing Lu, Tao Liu, Baisheng Li, Shujiang Mei, Feng Ruan, Lifeng Lin, Changwen Ke, Haojie Zhong, Yingtao Zhang, Lirong Zou, Xuguang Chen, Qi Zhu, Jianpeng Xiao, Jianxiang Geng, Zhe Liu, Jianxiong Hu, Weilin Zeng, Xing Li, Yuhuang Liao, Xiujuan Tang, Songjian Xiao, Ying Wang, Yingchao Song, Xue Zhuang, Lijun Liang, Guanhao He, Huihong Deng, Tie Song, Jianfeng He, Wenjun Ma |
| 15 | EPI_ISL_403937 | hCoV-19/Guangdong/20SF040/2020 | *Homo sapiens* | Asia / China / Guangdong / Zhuhai | 2020-01-18 | Guangdong Provincial Center for Diseases Control and Prevention; Guangdong Provincial Public Health | Department of Microbiology, Guangdong Provincial Center for Diseases Control and Prevention | Min Kang, Jie Wu, Jing Lu, Tao Liu, Baisheng Li, Shujiang Mei, Feng Ruan, Lifeng Lin, Changwen Ke, Haojie Zhong, Yingtao Zhang, Lirong Zou, Xuguang Chen, Qi Zhu, Jianpeng Xiao, Jianxiang Geng, Zhe Liu, Jianxiong Hu, Weilin Zeng, Xing Li, Yuhuang Liao, Xiujuan Tang, Songjian Xiao, Ying Wang, Yingchao Song, Xue Zhuang, Lijun Liang, Guanhao He, Huihong Deng, Tie Song, Jianfeng He, Wenjun Ma |
| 16 | EPI_ISL_403962 | hCoV-19/Nonthaburi/61/2020 | *Homo sapiens* | Asia / Thailand / Nonthaburi | 2020-01-08 | Bamrasnaradura Hospital | 1. Department of Medical Sciences, Ministry of Public Health, Thailand 2. Thai Red Cross Emerging Infectious Diseases - Health Science Centre 3. Department of Disease Control, Ministry of Public Health, Thailand | Pilailuk,Okada; Siripaporn,Phuygun; Thanutsapa,Thanadachakul; Supaporn,Wacharapluesadee; Sittiporn,Parnmen; Warawan,Wongboot; Sunthareeya,Waicharoen; Rome,Buathong; Malinee,Chittaganpitch; Nanthawan,Mekha |
| 16 | EPI_ISL_412899 | hCoV-19/Wuhan/HBCDC-HB-03/2019 | *Homo sapiens* | Asia / China / Hubei / Wuhan | 2019-12-30 | Wuhan Jinyintan Hospital | Hubei Provincial Center for Disease Control and Prevention | Bin Fang, Xiang Li, Xiao Yu, Linlin Liu, Bo Yang, Faxian Zhan, Guojun Ye, Xixiang Huo, Junqiang Xu, Bo Yu, Kun Cai, Jing Li, Yongzhong Jiang. |

| | | | | | | | | |
|---|---|---|---|---|---|---|---|---|
| 17 | EPI_ISL_404227 | hCoV-19/Zhejiang/WZ-01/2020 | *Homo sapiens* | Asia / China / Zhejiang | 2020-01-16 | Zhejiang Provincial Center for Disease Control and Prevention | Department of Microbiology, Zhejiang Provincial Center for Disease Control and Prevention | Yin Chen, Yanjun Zhang, Haiyan Mao, Junhang Pan, Xiuyu Lou, Yiyu Lu, Juying Yan, Hanping Zhu, Jian Gao, Yan Feng, Yi Sun, Hao Yan, Zhen Li, Yisheng Sun, Liming Gong, Qiong Ge, Wen Shi, Xinying Wang, Wenwu Yao, Zhangnv Yang, Fang Xu, Chen Chen, Enfu Chen, Zhen Wang, Zhiping Chen, Jianmin Jiang, Chonggao Hu |
| 18 | EPI_ISL_404895 | hCoV-19/USA/WA1/2020 | *Homo sapiens* | North America / USA / Washington / Snohomish County | 2020-01-19 | Providence Regional Medical Center | Division of Viral Diseases, Centers for Disease Control and Prevention | Queen,K., Tao,Y., Li,Y., Paden,C.R., Lu,X., Zhang,J., Gerber,S.I., Lindstrom,S., Tong,S. |
| 18 | EPI_ISL_407214 | hCoV-19/USA/WA1-A12/2020 | *Homo sapiens* | North America / USA / Washington | 2020-01-25 | WA State Department of Health | Pathogen Discovery, Respiratory Viruses Branch, Division of Viral Diseases, Centers for Dieases Control and Prevention | Krista Queen, Azaibi Tamin, Jennifer Harcourt, Ying Tao, Clinton R. Paden, Jing Zhang, Yan Li, Anna Uehara, Xiaoyan Lu, Shifaq Kamili, Rashi Gautam, Haibin Wang, Janna' R. Murray, Susan I. Gerber, Stephen Lindstrom, Natalie Thornburg, Suxiang Tong |
| 18 | EPI_ISL_407215 | hCoV-19/USA/WA1-F6/2020 | *Homo sapiens* | North America / USA / Washington | 2020-01-25 | Washington State Department of Health | Pathogen Discovery, Respiratory Viruses Branch, Division of Viral Diseases, Centers for Dieases Control and Prevention | Krista Queen, Azaibi Tamin, Jennifer Harcourt, Ying Tao, Clinton R. Paden, Jing Zhang, Yan Li, Anna Uehara, Xiaoyan Lu, Shifaq Kamili, Rashi Gautam, Haibin Wang, Janna' R. Murray, Susan I. Gerber, Stephen Lindstrom, Natalie Thornburg, Suxiang Tong |
| 18 | EPI_ISL_411060 | hCoV-19/Fujian/8/2020 | *Homo sapiens* | Asia / China / Fujian | 2020-01-21 | Fujian Center for Disease Control and Prevention | Fujian Center for Disease Control and Prevention | Chen Wei, Zhang Yanhua, He Wenxiang, Weng Yuwei |
| 18 | MT020881 | 2019-nCoV/USA-WA1-F6/2020 | *Homo sapiens* | USA: WA | 2020-01-25 | not specified | not specified | Queen,K., Tamin,A., Harcourt,J., Tao,Y., Paden,C.R., Zhang,J., Li,Y., Uehara,A., Lu,X., Kamili,S., Gautam,R., Wang,H., Murray,J.R., Gerber,S.I., Lindstrom,S., Thornburg,N. and Tong,S. |
| 18 | MT020880 | 2019-nCoV/USA-WA1-A12/2020 | *Homo sapiens* | USA: WA | 2020-01-25 | not specified | not specified | Queen,K., Tamin,A., Harcourt,J., Tao,Y., Paden,C.R., Zhang,J., Li,Y., Uehara,A., Lu,X., Kamili,S., Gautam,R., Wang,H., Murray,J.R., Gerber,S.I., Lindstrom,S., Thornburg,N. and Tong,S. |
| 18 | MN985325 | 2019-nCoV/USA-WA1/2020 | *Homo sapiens* | USA | 2020-01-19 | not specified | not specified | Harcourt,J., Tamin,A., Lu,X., Kamili,S., Sakthivel,S.K., Murray,J., Queen,K., Tao,Y., Paden,C.R., Zhang,J., Li,Y., Uehara,A., Wang,H., Goldsmith,C., Bullock,H.A., Wang,L., Whitaker,B., Lynch,B., Gautam,R., Schindewolf,C., Lokugamage,K.G., Scharton,D., Plante,J.A., Mirchandani,D., Widen,S.G., Narayanan,K., Makino,S., Ksiazek,T.G., Plante,K.S., Weaver,S.C., Lindstrom,S., Tong,S., Menachery,V.D. and Thornburg,N.J. |
| 19 | EPI_ISL_405839 | hCoV-19/Shenzhen/HKU-SZ-005/2020 | *Homo sapiens* | Asia / China / Guangdong / Shenzhen | 2020-01-11 | The University of Hong Kong - Shenzhen Hospital | Li Ka Shing Faculty of Medicine, The University of Hong Kong | Chan,J.F.-W., Yuan,S., Kok,K.H., To,K.K.-W., Chu,H., Yang,J., Xing,F., Liu,J., Yip,C.C.-Y., Poon,R.W.-S., Tsai,H.W., Lo,S.K.-F., Chan,K.H., Poon,V.K.-M., Chan,W.M., Ip,J.D., Cai,J.P., Cheng,V.C.-C., Chen,H., Hui,C.K.-M. and Yuen,K.Y. |
| 20 | EPI_ISL_406030 | hCoV-19/Shenzhen/HKU-SZ-002/2020 | *Homo sapiens* | Asia / China / Guangdong / Shenzhen | 2020-01-10 | The University of Hong Kong - Shenzhen Hospital | Li Ka Shing Faculty of Medicine, The University of Hong Kong | Chan,J.F.-W., Yuan,S., Kok,K.H., To,K.K.-W., Chu,H., Yang,J., Xing,F., Liu,J., Yip,C.C.-Y., Poon,R.W.-S., Tsai,H.W., Lo,S.K.-F., Chan,K.H., Poon,V.K.-M., Chan,W.M., Ip,J.D., Cai,J.P., Cheng,V.C.-C., Chen,H., Hui,C.K.-M. and Yuen,K.Y. |
| 21 | EPI_ISL_406031 | hCoV-19/Taiwan/2/2020 | *Homo sapiens* | Asia / Taiwan / Kaohsiung | 2020-01-23 | Centers for Disease Control, R.O.C. (Taiwan) | Centers for Disease Control, R.O.C. (Taiwan) | Ji-Rong Yang, Yu-Chi Lin, Jung-Jung Mu, Ming-Tsan Liu, Shu-Ying Li |
| 22 | EPI_ISL_406034 | hCoV-19/USA/CA1/2020 | *Homo sapiens* | North America / USA / California / Los Angeles | 2020-01-23 | California Department of Public Health | Pathogen Discovery, Respiratory Viruses Branch, Division of Viral Diseases, Centers for Dieases Control and Prevention | Anna Uehara, Krista Queen, Ying Tao, Yan Li, Clinton R. Paden, Jing Zhang, Xiaoyan Lu, Brian Lynch, Senthil Kumar K. Sakthivel, Brett L. Whitaker, Shifaq Kamili, Lijuan Wang, Janna' R. Murray, Susan I. Gerber, Stephen Lindstrom, Suxiang Tong |
| 23 | EPI_ISL_406036 | hCoV-19/USA/CA2/2020 | *Homo sapiens* | North America / USA / California / Orange County | 2020-01-22 | California Department of Public Health | Pathogen Discovery, Respiratory Viruses Branch, Division of Viral Diseases, Centers for Dieases Control and Prevention | Anna Uehara, Krista Queen, Ying Tao, Yan Li, Clinton R. Paden, Jing Zhang, Xiaoyan Lu, Brian Lynch, Senthil Kumar K. Sakthivel, Brett L. Whitaker, Shifaq Kamili, Lijuan Wang, Janna' R. Murray, Susan I. Gerber, Stephen Lindstrom, Suxiang Tong |
| 24 | EPI_ISL_406223 | hCoV-19/USA/AZ1/2020 | *Homo sapiens* | North America / USA / Arizona / Phoenix | 2020-01-22 | Arizona Department of Health Services | Pathogen Discovery, Respiratory Viruses Branch, Division of Viral Diseases, Centers for Disease Control and Prevention | Ying Tao, Clinton R. Paden, Krista Queen, Anna Uehara, Yan Li, Jing Zhang, Xiaoyan Lu, Brian Lynch, Senthil Kumar K. Sakthivel, Brett L. Whitaker, Shifaq Kamili, Lijuan Wang, Janna' R. Murray, Susan I. Gerber, Stephen Lindstrom, Suxiang Tong |

| | | | | | | | | |
|---|---|---|---|---|---|---|---|---|
| 25 | EPI_ISL_406531 | hCoV-19/Guangdong/20SF174/2020 | *Homo sapiens* | Asia / China / Guangdong / Zhuhai | 2020-01-22 | Guangdong Provincial Center for Diseases Contorl and Prevention; Guangdong Provinical Public Health | Guangdong Provincial Center for Disease Control and Prevention | Min Kang, Jie Wu, Jing Lu, Tao Liu, Baisheng Li, Shujiang Mei, Feng Ruan, Lifeng Lin, Changwen Ke, Haojie Zhong, Yingtao Zhang, Lirong Zou, Xuguang Chen, Qi Zhu, Jianpeng Xiao, Jianxiang Geng, Zhe Liu, Jianxiong Hu, Weilin Zeng, Xing Li, Yuhuang Liao, Xiujuan Tang, Songjian Xiao, Ying Wang, Yingchao Song, Xue Zhuang, Lijun Liang, Guanhao He, Huihong Deng, Tie Song, Jianfeng He, Wenjun Ma |
| 26 | EPI_ISL_406533 | hCoV-19/Guangzhou/20SF206/2020 | *Homo sapiens* | Asia / China / Guangdong / Guangzhou | 2020-01-22 | Guangdong Provincial Center for Diseases Contorl and Prevention; Guangdong Provinical Public Health | Guangdong Provincial Center for Diseases Control and Prevention | Min Kang, Jie Wu, Jing Lu, Tao Liu, Baisheng Li, Shujiang Mei, Feng Ruan, Lifeng Lin, Changwen Ke, Haojie Zhong, Yingtao Zhang, Lirong Zou, Xuguang Chen, Qi Zhu, Jianpeng Xiao, Jianxiang Geng, Zhe Liu, Jianxiong Hu, Weilin Zeng, Xing Li, Yuhuang Liao, Xiujuan Tang, Songjian Xiao, Ying Wang, Yingchao Song, Xue Zhuang, Lijun Liang, Guanhao He, Huihong Deng, Tie Song, Jianfeng He, Wenjun Ma |
| 27 | EPI_ISL_406534 | hCoV-19/Foshan/20SF207/2020 | *Homo sapiens* | Asia / China / Guangdong / Foshan | 2020-01-22 | Guangdong Provincial Center for Diseases Control and Prevention; Guangdong Provincial Public Health | Guangdong Provincial Center for Diseases Control and Prevention | Min Kang, Jie Wu, Jing Lu, Tao Liu, Baisheng Li, Shujiang Mei, Feng Ruan, Lifeng Lin, Changwen Ke, Haojie Zhong, Yingtao Zhang, Lirong Zou, Xuguang Chen, Qi Zhu, Jianpeng Xiao, Jianxiang Geng, Zhe Liu, Jianxiong Hu, Weilin Zeng, Xing Li, Yuhuang Liao, Xiujuan Tang, Songjian Xiao, Ying Wang, Yingchao Song, Xue Zhuang, Lijun Liang, Guanhao He, Huihong Deng, Tie Song, Jianfeng He, Wenjun Ma |
| 28 | EPI_ISL_406535 | hCoV-19/Foshan/20SF210/2020 | *Homo sapiens* | Asia / China / Guangdong / Foshan | 2020-01-22 | Guangdong Provincial Center for Diseases Control and Prevention; Guangdong Provincial Public Health | Guangdong Provincial Center for Diseases Control and Prevention | Min Kang, Jie Wu, Jing Lu, Tao Liu, Baisheng Li, Shujiang Mei, Feng Ruan, Lifeng Lin, Changwen Ke, Haojie Zhong, Yingtao Zhang, Lirong Zou, Xuguang Chen, Qi Zhu, Jianpeng Xiao, Jianxiang Geng, Zhe Liu, Jianxiong Hu, Weilin Zeng, Xing Li, Yuhuang Liao, Xiujuan Tang, Songjian Xiao, Ying Wang, Yingchao Song, Xue Zhuang, Lijun Liang, Guanhao He, Huihong Deng, Tie Song, Jianfeng He, Wenjun Ma |
| 29 | EPI_ISL_406536 | hCoV-19/Foshan/20SF211/2020 | *Homo sapiens* | Asia / China / Guangdong / Foshan | 2020-01-22 | Guangdong Provincial Center for Diseases Control and Prevention; Guangdong Provincial Public Health | Guangdong Provincial Center for Diseases Control and Prevention | Min Kang, Jie Wu, Jing Lu, Tao Liu, Baisheng Li, Shujiang Mei, Feng Ruan, Lifeng Lin, Changwen Ke, Haojie Zhong, Yingtao Zhang, Lirong Zou, Xuguang Chen, Qi Zhu, Jianpeng Xiao, Jianxiang Geng, Zhe Liu, Jianxiong Hu, Weilin Zeng, Xing Li, Yuhuang Liao, Xiujuan Tang, Songjian Xiao, Ying Wang, Yingchao Song, Xue Zhuang, Lijun Liang, Guanhao He, Huihong Deng, Tie Song, Jianfeng He, Wenjun Ma |
| 30 | EPI_ISL_406538 | hCoV-19/Guangdong/20SF201/2020 | *Homo sapiens* | Asia / China / Guangdong | 2020-01-23 | Guangdong Provincial Center for Diseases Control and Prevention;Guangdong Provincial Institute of Public Health | Guangdong Provincial Center for Diseases Control and Prevention | Min Kang, Jie Wu, Jing Lu, Tao Liu, Baisheng Li, Shujiang Mei, Feng Ruan, Lifeng Lin, Changwen Ke, Haojie Zhong, Yingtao Zhang, Lirong Zou, Xuguang Chen, Qi Zhu, Jianpeng Xiao, Jianxiang Geng, Zhe Liu, Jianxiong Hu, Weilin Zeng, Xing Li, Yuhuang Liao, Xiujuan Tang, Songjian Xiao, Ying Wang, Yingchao Song, Xue Zhuang, Lijun Liang, Guanhao He, Huihong Deng, Tie Song, Jianfeng He, Wenjun Ma |
| 31 | EPI_ISL_406593 | hCoV-19/Shenzhen/SZTH-002/2020 | *Homo sapiens* | Asia / China / Guandong / Shenzhen | 2020-01-13 | Shenzhen Key Laboratory of Pathogen and Immunity, National Clinical Research Center for Infectious Disease, Shenzhen Third People's Hospital | Shenzhen Key Laboratory of Pathogen and Immunity, National Clinical Research Center for Infectious Disease, Shenzhen Third People's Hospital | Yang Yang, Chenguang Shen, Li Xing, Zhixiang Xu, Haixia Zheng, Yingxia Liu |
| 32 | EPI_ISL_406594 | hCoV-19/Shenzhen/SZTH-003/2020 | *Homo sapiens* | Asia / China / Guandong / Shenzhen | 2020-01-16 | Shenzhen Key Laboratory of Pathogen and Immunity, National Clinical Research Center for Infectious Disease, Shenzhen Third People's Hospital | Shenzhen Key Laboratory of Pathogen and Immunity, National Clinical Research Center for Infectious Disease, Shenzhen Third People's Hospital | Yang Yang, Chenguang Shen, Li Xing, Zhixiang Xu, Haixia Zheng, Yingxia Liu |

| | | | | | | | | |
|---|---|---|---|---|---|---|---|---|
| 33 | EPI_ISL_406595 | hCoV-19/Shenzhen/SZTH-004/2020 | *Homo sapiens* | Asia / China / Guandong / Shenzhen | 2020-01-16 | Shenzhen Key Laboratory of Pathogen and Immunity, National Clinical Research Center for Infectious Disease, Shenzhen Third People's Hospital | Shenzhen Key Laboratory of Pathogen and Immunity, National Clinical Research Center for Infectious Disease, Shenzhen Third People's Hospital | Yang Yang, Chenguang Shen, Li Xing, Zhixiang Xu, Haixia Zheng, Yingxia Liu |
| 34 | EPI_ISL_406596 | hCoV-19/France/IDF0372/2020 | *Homo sapiens* | Europe / France / Ile-de-France / Paris | 2020-01-23 | Department of Infectious and Tropical Diseases, Bichat Claude Bernard Hospital, Paris | National Reference Center for Viruses of Respiratory Infections, Institut Pasteur, Paris | Mélanie Albert, Marion Barbet, Sylvie Behillil, Méline Bizard, Angela Brisebarre, Flora Donati, Vincent Enouf, Maud Vanpeene, Sylvie van der Werf, Yazdan Yazdanpanah, Xavier Lescure. |
| 35 | EPI_ISL_406597 | hCoV-19/France/IDF0373/2020 | *Homo sapiens* | Europe / France / Ile-de-France / Paris | 2020-01-23 | Department of Infectious and Tropical Diseases, Bichat Claude Bernard Hospital, Paris | National Reference Center for Viruses of Respiratory Infections, Institut Pasteur, Paris | Mélanie Albert, Marion Barbet, Sylvie Behillil, Méline Bizard, Angela Brisebarre, Flora Donati, Vincent Enouf, Maud Vanpeene, Sylvie van der Werf, Yazdan Yazdanpanah, Xavier Lescure. |
| 36 | EPI_ISL_406798 | hCoV-19/Wuhan/WH01/2019 | *Homo sapiens* | Asia / China / Hubei / Wuhan | 2019-12-26 | General Hospital of Central Theater Command of People's Liberation Army of China | BGI & Institute of Microbiology, Chinese Academy of Sciences & Shandong First Medical University & Shandong Academy of Medical Sciences & General Hospital of Central Theater Command of People's Liberation Army of China | Weijun Chen, Yuhai Bi, Weifeng Shi and Zhenhong Hu |
| 37 | EPI_ISL_406801 | hCoV-19/Wuhan/WH04/2020 | *Homo sapiens* | Asia / China / Hubei / Wuhan | 2020-01-05 | General Hospital of Central Theater Command of People's Liberation Army of China | BGI & Institute of Microbiology, Chinese Academy of Sciences & Shandong First Medical University & Shandong Academy of Medical Sciences & General Hospital of Central Theater Command of People's Liberation Army of China | Weijun Chen, Yuhai Bi, Weifeng Shi and Zhenhong Hu |
| 37 | EPI_ISL_411926 | hCoV-19/Taiwan/3/2020 | *Homo sapiens* | Asia / Taiwan / Taipei | 2020-01-24 | Taiwan Centers for Disease Control | Taiwan Centers for Disease Control | Ji-Rong Yang, Yu-Chi-Lin, Jung-Jung Mu, Ming-Tsan-Liu |
| 37 | EPI_ISL_413729 | hCoV-19/China/WF0015/2020 | *Homo sapiens* | Asia / China | 2020-02 | Weifang Center for Disease Control and Prevention | Weifang Center for Disease Control and Prevention & BGI-Shenzhen | Qing Nie, Xingguang Li, Erik M Volz, Han Fu, Haowei Wang, Xiaoyue Xi, Wei Chen, Dehui Liu, Yingying Chen, Mengmeng Tian, Wei Tan, Junjie Zai, Wanying Sun, Jiandong Li, Junhua Li |
| 37 | EPI_ISL_413750 | hCoV-19/China/WF0020/2020 | *Homo sapiens* | Asia / China | 2020-02 | Weifang Center for Disease Control and Prevention | Weifang Center for Disease Control and Prevention & BGI-Shenzhen | Qing Nie, Xingguang Li, Erik M Volz, Han Fu, Haowei Wang, Xiaoyue Xi, Wei Chen, Dehui Liu, Yingying Chen, Mengmeng Tian, Wei Tan, Junjie Zai, Wanying Sun, Jiandong Li, Junhua Li |
| 37 | EPI_ISL_414689 | hCoV-19/Guangzhou/GZMU0044/2020 | *Homo sapiens* | Asia / China / Guangdong / Guangzhou | 2020-02-25 | State Key Laboratory of Respiratory Disease, National Clinical Research Center for Respiratory Disease, Guangzhou Institute of Respiratory Health, the First Affiliated Hospital of Guangzhou Medical University | The First Affiliated Hospital of Guangzhou Medical University & BGI-Shenzhen | Zhao et al |
| 37 | EPI_ISL_414690 | hCoV-19/Guangzhou/GZMU0047/2020 | *Homo sapiens* | Asia / China / Guangdong / Guangzhou | 2020-02-25 | State Key Laboratory of Respiratory Disease, National Clinical Research Center for Respiratory Disease, Guangzhou Institute of Respiratory Health, the First Affiliated Hospital of Guangzhou Medical University | The First Affiliated Hospital of Guangzhou Medical University & BGI-Shenzhen | Zhao et al |

| | | | | | | | | |
|---|---|---|---|---|---|---|---|---|
| 37 | EPI_ISL_414691 | hCoV-19/Guangzhou/GZMU0048/2020 | *Homo sapiens* | Asia / China / Guangdong / Guangzhou | 2020-02-25 | State Key Laboratory of Respiratory Disease, National Clinical Research Center for Respiratory Disease, Guangzhou Institute of Respiratory Health, the First Affiliated Hospital of Guangzhou Medical University | The First Affiliated Hospital of Guangzhou Medical University & BGI-Shenzhen | Zhao et al |
| 37 | EPI_ISL_416042 | hCoV-19/Hangzhou/ZJU-02/2020 | *Homo sapiens* | Asia / China / Hangzhou | 2020-01-26 | State Key Laboratory for Diagnosis and Treatment of Infectious Diseases, National Clinical Research Center for Infectious Diseases, First Affiliated Hospital, Zhejiang University School of Medicine, Hangzhou, China. 310003 | State Key Laboratory for Diagnosis and Treatment of Infectious Diseases, National Clinical Research Center for Infectious Diseases, First Affiliated Hospital, Zhejiang University School of Medicine, Hangzhou, China. 310003 | Hangping Yao, Nanping Wu, Chao Jiang, Xiangyun Lu, Linfang Cheng, Fumin Liu, Zhigang Wu, Haibo Wu, Changzhong Jin, Min Zheng, Lanjuan Li |
| 37 | MT066175 | SARS-CoV-2/NTU01/2020/TWN | *Homo sapiens* | Taiwan | 2020-01-31 | not specified | not specified | Yeh,S.-H., Lin,Y.-Y., Lai,Y.-Y., Li,C.-L., Chang,S.-C., Chen,P.-J. and Chang,S.-Y. |
| 38 | EPI_ISL_406844 | hCoV-19/Australia/VIC01/2020 | *Homo sapiens* | Oceania / Australia / Victoria / Clayton | 2020-01-25 | Monash Medical Centre | Collaboration between the University of Melbourne at The Peter Doherty Institute for Infection and Immunity, and the Victorian Infectious Disease Reference Laboratory | Caly,L., Seemann,T., Schultz,M., Druce,J. and Taiaroa,G |
| 39 | EPI_ISL_406862 | hCoV-19/Germany/BavPat1/2020 | *Homo sapiens* | Europe / Germany / Bavaria / Munich | 2020-01-28 | Charité Universitätsmedizin Berlin, Institute of Virology; Institut für Mikrobiologie der Bundeswehr, Munich | Charité Universitätsmedizin Berlin, Institute of Virology | Victor M Corman, Julia Schneider, Talitha Veith, Barbara Mühlemann, Markus Antwerpen, Christian Drosten, Roman Wölfel |
| 40 | EPI_ISL_406970 | hCoV-19/Hangzhou/HZ-1/2020 | *Homo sapiens* | Asia / China / Zhejiang / Hangzhou | 2020-01-20 | Hangzhou Center for Disease and Control Microbiology Lab | Hangzhou Center for Disease and Control Microbiology Lab | Yu Hua, Wang Haoqiu, Li Jun, Yu Xinfeng |
| 41 | EPI_ISL_406973 | hCoV-19/Singapore/1/2020 | *Homo sapiens* | Asia / Singapore | 2020-01-23 | Singapore General Hospital | National Public Health Laboratory | Mak, TM; Octavia S; Chavatte JM; Zhou, ZY; Cui, L; Lin, RTP |
| 42 | EPI_ISL_407073 | hCoV-19/England/02/2020 | *Homo sapiens* | Europe / United Kingdom / England | 2020-01-29 | Respiratory Virus Unit, Microbiology Services Colindale, Public Health England | Respiratory Virus Unit, Microbiology Services Colindale, Public Health England | Monica Galiano, Shahjahan Miah, Richard Myers, Angie Lackenby, Omolola Akinbami, Tiina Talts, Leena Bhaw, Kirstin Edwards, Jonathan Hubb, Joanna Ellis, Maria Zambon. |
| 43 | EPI_ISL_407193 | hCoV-19/South Korea/KCDC03/2020 | *Homo sapiens* | Asia / South Korea / Gyeonggi-do | 2020-01-25 | Korea Centers for Disease Control & Prevention (KCDC) Center for Laboratory Control of Infectious Diseases Division of Viral Diseases | Korea Centers for Disease Control & Prevention (KCDC) Center for Laboratory Control of Infectious Diseases Division of Viral Diseases | Jeong-Min Kim, Yoon-Seok Chung, Namjoo Lee, Mi-Seon Kim, SangHee Woo, Hye-Joon Jo, Sehee Park, Heui Man Kim, Myung Guk Han |
| 44 | EPI_ISL_407313 | hCoV-19/Hangzhou/HZCDC0001/2020 | *Homo sapiens* | Asia / China / Zhejiang / Hangzhou | 2020-01-19 | Hangzhou Center for Disease Control and Prevention | Hangzhou Center for Disease Control and Prevention | Jun Li, Haoqiu Wang, Hua Yu, Lingfeng Mao, Xinfen Yu, Zhou Sun, Qingxin Kong, Xin Qian, Shuchang Chen, Xuchu Wang |
| 44 | EPI_ISL_416404 | hCoV-19/Shanghai/SH0119/2020 | *Homo sapiens* | Asia / China / Shanghai | 2020-02-09 | Shanghai Public Health Clinical Center, Shanghai Medical College, Fudan University | National Research Center for Translational Medicine (Shanghai), Ruijin Hospital affiliated to Shanghai Jiao Tong University School of Medicine & Shanghai Public Health Clinical Center | Shengyue Wang, Xiaonan Zhang, Gang Lu, Yun Tan, Yun Ling, Hongzhou Lu, Saijuan Chen |
| 45 | EPI_ISL_407893 | hCoV-19/Australia/NSW01/2020 | *Homo sapiens* | Oceania / Australia / New South Wales / Sydney | 2020-01-24 | Centre for Infectious Diseases and Microbiology Laboratory Services | NSW Health Pathology - Institute of Clinical Pathology and Medical Research; Westmead Hospital; University of Sydney | Eden J-S, Carter I, Rahman H, Holmes EC, Rockett R, O’Sullivan MV, Sintchenko V, Chen SC, Maddocks S, Kok J and Dwyer DE for the 2019-nCoV Study Group |

| | | | | | | | | |
|---|---|---|---|---|---|---|---|---|
| 46 | EPI_ISL_407894 | hCoV-19/Australia/QLD01/2020 | *Homo sapiens* | Oceania / Australia / Queensland / Gold Coast | 2020-01-28 | Pathology Queensland | Public Health Virology Laboratory | Ben Huang, Alyssa Pyke, Amanda De Jong, Andrew Van Den Hurk, Carmel Taylor, David Warrilow, Doris Genge, Elisabeth Gamez, Glen Hewitson, Ian Maxwell Mackay, Inga Sultana, Jamie McMahon, Jean Barcelon, Judy Northill, Mitchell Finger, Natalie Simpson, Neelima Nair, Peter Burtonclay, Peter Moore, Sarah Wheatley, Sean Moody, Sonja Hall-Mendelin, Timothy Gardam, and Frederick Moore. |
| 47 | EPI_ISL_407896 | hCoV-19/Australia/QLD02/2020 | *Homo sapiens* | Oceania / Australia / Queensland / Gold Coast | 2020-01-30 | Pathology Queensland | Public Health Virology Laboratory | Ben Huang, Alyssa Pyke, Amanda De Jong, Andrew Van Den Hurk, Carmel Taylor, David Warrilow, Doris Genge, Elisabeth Gamez, Glen Hewitson, Ian Maxwell Mackay, Inga Sultana, Jamie McMahon, Jean Barcelon, Judy Northill, Mitchell Finger, Natalie Simpson, Neelima Nair, Peter Burtonclay, Peter Moore, Sarah Wheatley, Sean Moody, Sonja Hall-Mendelin, Timothy Gardam, and Frederick Moore. |
| 48 | EPI_ISL_407976 | hCoV-19/Belgium/GHB-03021/2020 | *Homo sapiens* | Europe / Belgium / Leuven | 2020-02-03 | KU Leuven, Clinical and Epidemiological Virology | KU Leuven, Clinical and Epidemiological Virology | Bert Vanmechelen, Elke Wollants, Annabel Rector, Els Keyaerts, Lies Laenen, Marc Van Ranst, and Piet Maes |
| 49 | EPI_ISL_407987 | hCoV-19/Singapore/2/2020 | *Homo sapiens* | Asia / Singapore | 2020-01-25 | Singapore General Hospital | Programme in Emerging Infectious Diseases, Duke-NUS Medical School | Danielle E Anderson, Martin Linster, Yan Zhuang, Jayanthi Jayakumar, Kian Sing Chan, Lynette LE Oon, Jenny GH Low, Yvonne CF Su, Linfa Wang, Gavin JD Smith |
| 49 | EPI_ISL_410537 | hCoV-19/Singapore/6/2020 | *Homo sapiens* | Asia / Singapore | 2020-02-09 | Singapore General Hospital, Molecular Laboratory, Division of Pathology | Programme in Emerging Infectious Diseases, Duke-NUS Medical School | Danielle E Anderson, Martin Linster, Yan Zhuang, Jayanthi Jayakumar, Kian Sing Chan, Lynette LE Oon, Shirin Kalimuddin, Jenny GH Low, Yvonne CF Su, Gavin JD Smith |
| 50 | EPI_ISL_407988 | hCoV-19/Singapore/3/2020 | *Homo sapiens* | Asia / Singapore | 2020-02-01 | National Centre for Infectious Diseases | Programme in Emerging Infectious Diseases, Duke-NUS Medical School | Danielle E Anderson, Martin Linster, Yan Zhuang, Jayanthi Jayakumar, David CB Lye, Yee Sin Leo, Barnaby E Young, Yvonne CF Su, Linfa Wang, Gavin JD Smith |
| 51 | EPI_ISL_408008 | hCoV-19/USA/CA3/2020 | *Homo sapiens* | North America / USA / California | 2020-01-29 | California Department of Health | Pathogen Discovery, Respiratory Viruses Branch, Division of Viral Diseases, Centers for Disease Control and Prevention | Krista Queen, Jing Zhang, Yan Li, Ying Tao, Anna Uehara, Clinton Paden, Xiaoyan Lu, Brian Lynch, Senthil Kumar K. Sakthivel, Brett L. Whitaker, Shifaq Kamili, Lijuan Wang, Janna' R. Murray, Susan I. Gerber, Stephen Lindstrom, Suxiang Tong |
| 51 | EPI_ISL_408009 | hCoV-19/USA/CA4/2020 | *Homo sapiens* | North America / USA / California | 2020-01-29 | California Department of Health | Pathogen Discovery, Respiratory Viruses Branch, Division of Viral Diseases, Centers for Dieases Control and Prevention | Krista Queen, Jing Zhang, Yan Li, Ying Tao, Anna Uehara, Clinton Paden, Xiaoyan Lu, Brian Lynch, Senthil Kumar K. Sakthivel, Brett L. Whitaker, Shifaq Kamili, Lijuan Wang, Janna' R. Murray, Susan I. Gerber, Stephen Lindstrom, Suxiang Tong |
| 51 | MT027063 | 2019-nCoV/USA-CA4/2020 | *Homo sapiens* | USA: CA | 2020-01-29 | not specified | not specified | Queen,K., Zhang,J., Li,Y., Tao,Y., Uehara,A., Paden,C.R., Lu,X., Lynch,B., Sakthivel,S.K.K., Whitaker,B.L., Kamili,S., Wang,L., Murray,J.R., Gerber,S.I., Lindstrom,S. and Tong,S. |
| 52 | EPI_ISL_408010 | hCoV-19/USA/CA5/2020 | *Homo sapiens* | North America / USA / California | 2020-01-29 | California Department of Health | Pathogen Discovery, Respiratory Viruses Branch, Division of Viral Diseases, Centers for Dieases Control and Prevention | Ying Tao, Krista Queen, Jing Zhang, Yan Li, Anna Uehara, Clinton Paden, Xiaoyan Lu, Brian Lynch, Senthil Kumar K. Sakthivel, Brett L. Whitaker, Shifaq Kamili, Lijuan Wang, Janna' R. Murray, Susan I. Gerber, Stephen Lindstrom, Suxiang Tong |
| 53 | EPI_ISL_408430 | hCoV-19/France/IDF0515/2020 | *Homo sapiens* | Europe / France / Ile-de-France / Paris | 2020-01-29 | Department of Infectious and Tropical Diseases, Bichat Claude Bernard Hospital, Paris | National Reference Center for Viruses of Respiratory Infections, Institut Pasteur, Paris | Mélanie Albert, Marion Barbet, Sylvie Behillil, Méline Bizard, Angela Brisebarre, Flora Donati, Vincent Enouf, Maud Vanpeene, Sylvie van der Werf, Yazdan Yazdanpanah, Xavier Lescure |
| 54 | EPI_ISL_408431 | hCoV-19/France/IDF0626/2020 | *Homo sapiens* | Europe / France / Ile-de-France / Paris | 2020-01-29 | Sorbonne Université, Inserm et Assistance Publique-Hôpitaux de Paris (Pitié Salpétrière) | National Reference Center for Viruses of Respiratory Infections, Institut Pasteur, Paris | Mélanie Albert, Marion Barbet, Sylvie Behillil, Méline Bizard, Angela Brisebarre, Flora Donati, Vincent Enouf, Maud Vanpeene, Sylvie van der Werf, Sonia Burrel, Anne-Geneviève Marcelin, Vincent Calvez, David Boutolleau, Elise Klément, Valérie Pourcher, Eric Caumes. |
| 55 | EPI_ISL_408478 | hCoV-19/Chongqing/YC01/2020 | *Homo sapiens* | Asia / China / Chongqinq / Yongchuan | 2020-01-21 | Yongchuan District Center for Disease Control and Prevention | Chongqing Municipal Center for Disease Control and Prevention | Ye Sheng, Tang Yun, Ling Hua,Yu zhen,Chen Shuang,Tan ZhangPing, Su Kun, Li Qing, Tang Wenge, Rong Rong |
| 56 | EPI_ISL_408480 | hCoV-19/Yunnan/IVDC-YN-003/2020 | *Homo sapiens* | Asia / China / Yunnan / Kunming | 2020-01-17 | National Institute for Viral Disease Control and Prevention, China CDC | National Institute for Viral Disease Control & Prevention, CCDC | Wenjie Tan, Xiaoqing Fu, Xiang Zhao, Wenling Wang, Peihua Niu, Roujian Lu,Yanhong Sun, Baoying Huang, Li Zhao, Fei Ye, Wenbo Xu, George F. Gao, Guizhen Wu |
| 57 | EPI_ISL_408481 | hCoV-19/Chongqing/IVDC-CQ-001/2020 | *Homo sapiens* | Asia / China / Chongqing | 2020-01-18 | National Institute for Viral Disease Control and Prevention, China CDC | National Institute for Viral Disease Control & Prevention, CCDC | Wenjie Tan, Hengqin Wang, Xiang Zhao, Wenling Wang, Peihua Niu, Roujian Lu, Sheng Ye, Baoying Huang, Li Zhao, Fei Ye, Wenbo Xu, George F. Gao, Guizhen Wu |

| | | | | | | | | |
|---|---|---|---|---|---|---|---|---|
| 57 | EPI_ISL_412968 | hCoV-19/Japan/Hu_DP_Kng_19-020/2020 | *Homo sapiens* | Asia / Japan | 2020-02-10 | unknown | Takayuki Hishiki Kanagawa Prefectural Institute of Public Health, Department of Microbiology | Hishiki,T., Suzuki,R., Sakuragi,J., Usui,K., Tanaka,Y., Kawai,J., Kogo,Y., Matsuki,Y., An,T., Hayashizaki,Y. and Takasaki,T. |
| 57 | EPI_ISL_414484 | hCoV-19/USA/CruiseA-25/2020 | *Homo sapiens* | North America / USA | 2020-02-17 | unknown | Pathogen Discovery, Respiratory Viruses Branch, Division of Viral Diseases, Centers for Disease Control and Prevention | Krista Queen, Anna Uehara, Ying Tao, Clinton R. Paden, Jing Zhang, Yan Li, Haibin Wang, Shifaq Kamili, Xiaoyan Lu, Brian Lynch, Senthil Kumar K. Sakthivel, Brett L. Whitaker, Lijuan Wang, Janna' R. Murray, Jasmine Padilla, Justin Lee, Susan I. Gerber, Stephen Lindstrom, Suxiang Tong |
| 57 | EPI_ISL_416574 | hCoV-19/Japan/DP0107/2020 | *Homo sapiens* | Asia / Japan / unknown | 2020-02-15 | Japanese Quarantine Stations | Pathogen Genomics Center, National Institute of Infectious Diseases | Tsuyoshi Sekizuka, Kentaro Itokawa, Rina Tanaka, Masanori Hashino, Tsutomu Kageyama, Shinji Saito, Ikuyo Takayama, Hideki Hasegawa, Takuri Takahashi, Hajime Kamiya, Takuya Yamagishi, Motoi Suzuki, Takaji Wakita, Makoto Kuroda |
| 57 | EPI_ISL_416577 | hCoV-19/Japan/DP0134/2020 | *Homo sapiens* | Asia / Japan / unknown | 2020-02-15 | Japanese Quarantine Stations | Pathogen Genomics Center, National Institute of Infectious Diseases | Tsuyoshi Sekizuka, Kentaro Itokawa, Rina Tanaka, Masanori Hashino, Tsutomu Kageyama, Shinji Saito, Ikuyo Takayama, Hideki Hasegawa, Takuri Takahashi, Hajime Kamiya, Takuya Yamagishi, Motoi Suzuki, Takaji Wakita, Makoto Kuroda |
| 58 | EPI_ISL_408482 | hCoV-19/Shandong/IVDC-SD-001/2020 | *Homo sapiens* | Asia / China / Shandong / Qingdao | 2020-01-19 | National Institute for Viral Disease Control and Prevention, China CDC | National Institute for Viral Disease Control & Prevention, CCDC | Wenjie Tan, Zhaoguo Wang, Xiang Zhao, Wenling Wang, Peihua Niu, Roujian Lu, Ti Liu, Baoying Huang, Li Zhao, Fei Ye, Wenbo Xu, George F. Gao, Guizhen Wu |
| 59 | EPI_ISL_408484 | hCoV-19/Sichuan/IVDC-SC-001/2020 | *Homo sapiens* | Asia / China / Sichuan / Chengdu | 2020-01-15 | National Institute for Viral Disease Control and Prevention, China CDC | National Institute for Viral Disease Control & Prevention, CCDC | Wenjie Tan, Jianan Xu, Wenling Wang, Peihua Niu, Roujian Lu, Huiping Yang, Xiang Zhao, Baoying Huang, Li Zhao, Fei Ye, Wenbo Xu, George F. Gao, Guizhen Wu |
| 60 | EPI_ISL_408485 | hCoV-19/Beijing/IVDC-BJ-005/2020 | *Homo sapiens* | Asia / China / Beijing | 2020-01-18 | National Institute for Viral Disease Control and Prevention, China CDC | National Institute for Viral Disease Control & Prevention, CCDC | Wenjie Tan,Quanyi Wang,Wenling Wang, Peihua Niu,Roujian Lu,Yang Pan,Xiang Zhao,Baoying Huang,Li Zhao,Fei Ye,Wenbo Xu,George F. Gao,Guizhen Wu |
| 61 | EPI_ISL_408486 | hCoV-19/Jiangxi/IVDC-JX-002/2020 | *Homo sapiens* | Asia / China / Jiangxi / Pingxiang | 2020-01-11 | National Institute for Viral Disease Control and Prevention, China CDC | National Institute for Viral Disease Control & Prevention, CCDC | Wenjie Tan, Yong Shi, Wenling Wang, Peihua Niu, Roujian Lu, Jianxiong Li, Xiang Zhao, Baoying Huang, Li Zhao, Fei Ye, Wenbo Xu, George F. Gao, Guizhen Wu |
| 62 | EPI_ISL_408488 | hCoV-19/Jiangsu/IVDC-JS-001/2020 | *Homo sapiens* | Asia / China / Jiangsu / Huaian | 2020-01-19 | National Institute for Viral Disease Control and Prevention, China CDC | National Institute for Viral Disease Control & Prevention, CCDC | Wenjie Tan, Shenjiao Wang, Wenling Wang, Peihua Niu, Roujian Lu, Kangchen Zhao, Xiang Zhao, Baoying Huang, Li Zhao, Fei Ye, Wenbo Xu, George F. Gao, Guizhen Wu |
| 63 | EPI_ISL_408514 | hCoV-19/Wuhan/IVDC-HB-envF13-20/2020 | Environment | Asia / China / Hubei / Wuhan | 2020-01-01 | Institute of Viral Disease Control and Prevention, China CDC | Institute of Viral Disease Control and Prevention, China CDC | William J. Liu, Peipei Liu, Xiang Zhao, Peihua Niu, Yingze Zhao, Wenwen Lei, Ziqian Xu, Shumei Zou, Wei Zhen, Beiwei Ye, Mengjie Yang, Weifeng Shi, Roujian Lu, Wenjie Tan, Zhixiao Chen, Yuchao Wu, Juan Song, Weimin Zhou, Dayan Wang, Jun Han, Wenbo Xu, George F. Gao, Guizhen Wu |
| 64 | EPI_ISL_408515 | hCoV-19/Wuhan/IVDC-HB-envF13-21/2020 | Environment | Asia / China / Hubei / Wuhan | 2020-01-01 | Institute of Viral Disease Control and Prevention, China CDC | Institute of Viral Disease Control and Prevention, China CDC | William J. Liu, Peipei Liu, Xiang Zhao, Peihua Niu, Yingze Zhao, Wenwen Lei, Ziqian Xu, Shumei Zou, Wei Zhen, Beiwei Ye, Mengjie Yang, Weifeng Shi, Roujian Lu, Wenjie Tan, Zhixiao Chen, Yuchao Wu, Juan Song, Weimin Zhou, Dayan Wang, Jun Han, Wenbo Xu, George F. Gao, Guizhen Wu |
| 65 | EPI_ISL_408665 | hCoV-19/Japan/TY-WK-012/2020 | *Homo sapiens* | Asia / Japan / Tokyo | 2020-01-29 | Dept. of Virology III, National Institute of Infectious Diseases | Pathogen Genomics Center, National Institute of Infectious Diseases | Tsuyoshi Sekizuka, Shutoku Matsuyama, Naganori Nao, Kazuya Shirato, Makoto Takeda, Makoto Kuroda |
| 66 | EPI_ISL_408666 | hCoV-19/Japan/TY-WK-501/2020 | *Homo sapiens* | Asia / Japan / Tokyo | 2020-01-31 | Dept. of Virology III, National Institute of Infectious Diseases | Pathogen Genomics Center, National Institute of Infectious Diseases | Tsuyoshi Sekizuka, Shutoku Matsuyama, Naganori Nao, Kazuya Shirato, Makoto Takeda, Makoto Kuroda |
| 67 | EPI_ISL_408667 | hCoV-19/Japan/TY-WK-521/2020 | *Homo sapiens* | Asia / Japan / Tokyo | 2020-01-31 | Dept. of Virology III, National Institute of Infectious Diseases | Pathogen Genomics Center, National Institute of Infectious Diseases | Tsuyoshi Sekizuka, Shutoku Matsuyama, Naganori Nao, Kazuya Shirato, Makoto Takeda, Makoto Kuroda |
| 68 | EPI_ISL_408668 | hCoV-19/Vietnam/VR03-38142/2020 | *Homo sapiens* | Asia / Vietnam / Thanh Hoa | 2020-01-24 | National Influenza Center - National Institute of Hygiene and Epidemiology (NIHE) | National Influenza Center - National Institute of Hygiene and Epidemiology (NIHE) | Ung Thi Hong Trang, Hoang Vu Mai Phuong, Nguyen Le Khanh Hang, Nguyen Vu Son, Le Thi Thanh, Vuong Duc Cuong, Nguyen Phuong Anh, Pham Thi Hien, Tran Thu Huong, Le Thi Quynh Mai, |
| 69 | EPI_ISL_408669 | hCoV-19/Japan/KY-V-029/2020 | *Homo sapiens* | Asia / Japan / Kyoto | 2020-01-29 | Dept. of Virology III, National Institute of Infectious Diseases | Pathogen Genomics Center, National Institute of Infectious Diseases | Tsuyoshi Sekizuka, Shutoku Matsuyama, Naganori Nao, Kazuya Shirato, Makoto Takeda, Makoto Kuroda |

| | | | | | | | | |
|---|---|---|---|---|---|---|---|---|
| 70 | EPI_ISL_408977 | hCoV-19/Australia/NSW03/2020 | *Homo sapiens* | Oceania / Australia / New South Wales / Sydney | 2020-01-25 | Serology, Virology and OTDS Laboratories (SAViD), NSW Health Pathology Randwick | NSW Health Pathology - Institute of Clinical Pathology and Medical Research; Centre for Infectious Diseases and Microbiology Laboratory Services; Westmead Hospital; University of Sydney | Eden J-S, Carter I, Rahman H, Rawlinson W, Holmes EC, Rockett R, O'Sullivan MV, Sintchenko V, Chen SC, Maddocks S, Kok J and Dwyer DE for the 2019-nCoV Study Group* |
| 71 | EPI_ISL_410045 | hCoV-19/USA/IL2/2020 | *Homo sapiens* | North America / USA / Illinois | 2020-01-28 | IL Department of Public Health Chicago Laboratory | Pathogen Discovery, Respiratory Viruses Branch, Division of Viral Diseases, Centers for Dieases Control and Prevention | Yan Li, Jing Zhang, Krista Queen, Ying Tao, Anna Uehara, Clinton R. Paden, Xiaoyan Lu, Brian Lynch, Senthil Kumar K. Sakthivel, Brett L. Whitaker, Shifaq Kamili, Lijuan Wang, Janna' R. Murray, Susan I. Gerber, Stephen Lindstrom, Suxiang Tong |
| 72 | EPI_ISL_410218 | hCoV-19/Taiwan/NTU02/2020 | *Homo sapiens* | Asia / Taiwan / Taipei | 2020-02-05 | Department of Laboratory Medicine, National Taiwan University Hospital | Microbial Genomics Core Lab, National Taiwan University Centers of Genomic and Precision Medicine | Shiou-Hwei Yeh, You-Yu Lin, Ya-Yun Lai, Chiao-Ling Li, Shan-Chwen Chang, Pei-Jer Chen, Sui-Yuan Chang |
| 73 | EPI_ISL_410301 | hCoV-19/Nepal/61/2020 | *Homo sapiens* | Asia / Nepal / Kathmandu | 2020-01-13 | National Influenza Centre, National Public Health Laboratory, Kathmandu, Nepal | The University of Hong Kong | Ranjit Sah , Runa Jha, Daniel Chu, Haogao Gu, Malik Peiris, Anup Bastola, Alfonso J. Rodriguez-Morales, Bibek Kumar Lal, Basu Dev Pandey, Leo Poon |
| 74 | EPI_ISL_410536 | hCoV-19/Singapore/5/2020 | *Homo sapiens* | Asia / Singapore | 2020-02-06 | Singapore General Hospital, Molecular Laboratory, Division of Pathology | Programme in Emerging Infectious Diseases, Duke-NUS Medical School | Danielle E Anderson, Martin Linster, Yan Zhuang, Jayanthi Jayakumar, Kian Sing Chan, Lynette LE Oon, Shirin Kalimuddin, Jenny GH Low, Yvonne CF Su, Gavin JD Smith |
| 75 | EPI_ISL_410713 | hCoV-19/Singapore/7/2020 | *Homo sapiens* | Asia / Singapore | 2020-01-27 | National Public Health Laboratory, National Centre for Infectious Diseases | National Public Health Laboratory, National Centre for Infectious Diseases | Octavia S, Mak TM, Cui L, Lin RTP |
| 76 | EPI_ISL_410715 | hCoV-19/Singapore/9/2020 | *Homo sapiens* | Asia / Singapore | 2020-02-04 | National Public Health Laboratory, National Centre for Infectious Diseases | National Public Health Laboratory, National Centre for Infectious Diseases | Octavia S, Mak TM, Cui L, Lin RTP |
| 76 | EPI_ISL_410716 | hCoV-19/Singapore/10/2020 | *Homo sapiens* | Asia / Singapore | 2020-02-04 | National Public Health Laboratory, National Centre for Infectious Diseases | National Centre for Infectious Diseases, National Centre for Infectious Diseases | Octavia S, Mak TM, Cui L, Lin RTP |
| 77 | EPI_ISL_410717 | hCoV-19/Australia/QLD03/2020 | *Homo sapiens* | Oceania / Australia / Queensland / Gold Coast | 2020-02-05 | Pathology Queensland | Public Health Virology Laboratory | Ben Huang, Alyssa Pyke, Amanda De Jong, Andrew Van Den Hurk, Carmel Taylor, David Warrilow, Doris Genge, Elisabeth Gamez, Glen Hewitson, Ian Maxwell Mackay, Inga Sultana, Jamie McMahon, Jean Barcelon, Judy Northill, Mitchell Finger, Natalie Simpson, Neelima Nair, Peter Burtonclay, Peter Moore, Sarah Wheatley, Sean Moody, Sonja Hall-Mendelin, Timothy Gardam, and Frederick Moore. |
| 78 | EPI_ISL_410718 | hCoV-19/Australia/QLD04/2020 | *Homo sapiens* | Oceania / Australia / Queensland / Gold Coast | 2020-02-05 | Pathology Queensland | Public Health Virology Laboratory | Ben Huang, Alyssa Pyke, Amanda De Jong, Andrew Van Den Hurk, Carmel Taylor, David Warrilow, Doris Genge, Elisabeth Gamez, Glen Hewitson, Ian Maxwell Mackay, Inga Sultana, Jamie McMahon, Jean Barcelon, Judy Northill, Mitchell Finger, Natalie Simpson, Neelima Nair, Peter Burtonclay, Peter Moore, Sarah Wheatley, Sean Moody, Sonja Hall-Mendelin, Timothy Gardam, and Frederick Moore. |
| 78 | EPI_ISL_411954 | hCoV-19/USA/CA7/2020 | *Homo sapiens* | North America / USA / California | 2020-02-06 | California Department of Public Health | Pathogen Discovery, Respiratory Viruses Branch, Division of Viral Diseases, Centers for Dieases Control and Prevention | Krista Queen, Anna Uehara, Jing Zhang, Yan Li, Ying Tao, Clinton R. Paden, Haibin Wang, Shifaq Kamili, Xiaoyan Lu, Brian Lynch, Senthil Kumar K. Sakthivel, Brett L. Whitaker, Lijuan Wang, Janna' R. Murray, Susan I. Gerber, Stephen Lindstrom, Suxiang Tong |
| 78 | EPI_ISL_413853 | hCoV-19/Guangdong/2020XN4243-P0035/2020 | *Homo sapiens* | Asia / China / Guangdong | 2020-01-30 | Guangdong Provincial Institution of Public Health, Guangdong Provincial Center for Disease Control and Prevention | Guangdong Provincial Institution of Public Health | Jing Lu, Louis du Plessis, Liu Zhe, Jiufeng Sun, Sarah François, Huifang Lin, Moritz Kraemer, Jingju Peng, Qianlin Xiong, Runyu Yuan, Lilian Zeng, Pingping Zhou, Chuming Liang, Tao Liu, Wei Li, Juan Su, Huanying Zheng, Kang Min, Song Tie, Bo Peng, Shisong Fang, Wenzhe Su, Kuibiao Li, Ruilin Sun, Ru bai, Xi Tang, Minfeng Liang, Nuno Faria, Josh Quick, Andrew Rambaut, Verity Hill, Wenjun Ma, Nick Loman, Oliver Pybus, Changwen Ke |
| 79 | EPI_ISL_410719 | hCoV-19/Singapore/11/2020 | *Homo sapiens* | Asia / Singapore | 2020-02-02 | National Public Health Laboratory | National Public Health Laboratory | Octavia S, Mak TM, Cui L, Lin RTP |

| | | | | | | | | |
|---|---|---|---|---|---|---|---|---|
| 80 | EPI_ISL_410720 | hCoV-19/France/IDF0372-isl/2020 | *Homo sapiens* | Europe / France / Ile-de-France / Paris | 2020-01-23 | Department of Infectious and Tropical Diseases, Bichat Claude Bernard Hospital, Paris | National Reference Center for Viruses of Respiratory Infections, Institut Pasteur, Paris | Mélanie Albert, Marion Barbet, Sylvie Behillil, Méline Bizard, Angela Brisebarre, Flora Donati, Vincent Enouf, Maud Vanpeene, Sylvie van der Werf, Yazdan Yazdanpanah, Xavier Lescure. |
| 80 | EPI_ISL_411219 | hCoV-19/France/IDF0386-islP1/2020 | *Homo sapiens* | Europe / France / Ile-de-France / Paris | 2020-01-28 | Department of Infectious and Tropical Diseases, Bichat Claude Bernard Hospital, Paris | Laboratoire Virpath, CIRI U111, UCBL1, INSERM, CNRS, ENS Lyon | Olivier Terrier, Aurélien Traversier, Julien Fouret, Yazdan Yazdanpanah, Xavier Lescure, Alexandre Gaymard, Bruno Lina, Manuel Rosa-Calatrava |
| 80 | EPI_ISL_411220 | hCoV-19/France/IDF0386-islP3/2020 | *Homo sapiens* | Europe / France / Ile-de-France / Paris | 2020-01-28 | Department of Infectious and Tropical Diseases, Bichat Claude Bernard Hospital, Paris | Laboratoire Virpath, CIRI U111, UCBL1, INSERM, CNRS, ENS Lyon | Olivier Terrier, Aurélien Traversier, Julien Fouret, Yazdan Yazdanpanah, Xavier Lescure, Alexandre Gaymard, Bruno Lina, Manuel Rosa-Calatrava |
| 81 | EPI_ISL_410984 | hCoV-19/France/IDF0515-isl/2020 | *Homo sapiens* | Europe / France / Ile-de-France / Paris | 2020-01-29 | Department of Infectious and Tropical Diseases, Bichat Claude Bernard Hospital, Paris | National Reference Center for Viruses of Respiratory Infections, Institut Pasteur, Paris | Mélanie Albert, Marion Barbet, Sylvie Behillil, Méline Bizard, Angela Brisebarre, Flora Donati, Vincent Enouf, Maud Vanpeene, Sylvie van der Werf, Yazdan Yazdanpanah, Xavier Lescure |
| 81 | EPI_ISL_411218 | hCoV-19/France/IDF0571/2020 | *Homo sapiens* | Europe / France / Ile-de-France / Paris | 2020-02-02 | Department of Infectious and Tropical Diseases, Bichat Claude Bernard Hospital, Paris | Laboratoire Virpath, CIRI U111, UCBL1, INSERM, CNRS, ENS Lyon | Olivier Terrier, Aurélien Traversier, Julien Fouret, Yazdan Yazdanpanah, Xavier Lescure, Catherine Legras-Lachuer, Alexandre Gaymard, Bruno Lina, Manuel Rosa-Calatrava |
| 82 | EPI_ISL_411066 | hCoV-19/Fujian/13/2020 | *Homo sapiens* | Asia / China / Fujian | 2020-01-22 | Fujian Center for Disease Control and Prevention | Fujian Center for Disease Control and Prevention | Chen Wei, Zhang Yanhua, He Wenxiang, Weng Yuwei |
| 83 | EPI_ISL_411902 | hCoV-19/Cambodia/0012/2020 | *Homo sapiens* | Asia / Cambodia / Sihanoukville | 2020-01-27 | Virology Unit, Institut Pasteur du Cambodge. | Virology Unit, Institut Pasteur du Cambodge (Sequencing done by: Jessica E Manning/Jennifer A Bohl at Malaria and Vector Research Research Laboratory, National Institute of Allergy and Infectious Diseases and Vida Ahyong from Chan-Zuckerberg Biohub) | Erik A Karlsson, Jennifer A Bohl, Vida Ahyong, Veasna Duong, Philippe Dussart, Jessica E Manning. |
| 84 | EPI_ISL_411929 | hCoV-19/South Korea/SNU01/2020 | *Homo sapiens* | Asia / South Korea | 2020-01 | unknown | Department of Clinical Diagnostics | Park,W.B., Kwon,N.-J., Choi,S.-J., Kang,C.K., Choe,P.G., Kim,J.Y., Yun,J., Lee,G.-W., Seong,M.-W., Kim,N., Seo,J.-S. and Oh,M.-D. |
| 85 | EPI_ISL_411950 | hCoV-19/Jiangsu/JS01/2020 | *Homo sapiens* | Asia / China / Jiangsu | 2020-01-23 | NHC Key laboratory of Enteric Pathogenic Microbiology, Institute of Pathogenic Microbiology | Jiangsu Provincial Center for Disease Control & Prevention | Lunbiao Cui,Kangchen Zhao,Xiaojuan Zhu,Yiyue Ge,Tao Wu,Bin Wu,Yin Chen,Fengcai Zhu,Baoli Zhu,Ming Wu |
| 86 | EPI_ISL_411951 | hCoV-19/Sweden/01/2020 | *Homo sapiens* | Europe / Sweden | 2020-02-07 | unknown | Unit for Laboratory Development and Technology Transfer, Public Health Agency of Sweden | Bengner,M., Palmerus,M., Lindsjo,O., Lind Karlberg,M., Monteil,V., Appelberg,S., Brave,A., Muradrasoli,S. and Tegmark-Wisell,K. |
| 87 | EPI_ISL_411952 | hCoV-19/Jiangsu/JS02/2020 | *Homo sapiens* | Asia / China / Jiangsu | 2020-01-24 | NHC Key laboratory of Enteric Pathogenic Microbiology, Institute of Pathogenic Microbiology | Jiangsu Provincial Center for Disease Control & Prevention | Kangchen Zhao, Xiaojuan Zhu, Lunbiao Cui, Tao Wu, Yiyue Ge, Bin Wu, Yin Chen, Fengcai Zhu, Baoli Zhu, Ming Wu |
| 88 | EPI_ISL_411955 | hCoV-19/USA/CA8/2020 | *Homo sapiens* | North America / USA / California | 2020-02-10 | California Department of Public Health | Pathogen Discovery, Respiratory Viruses Branch, Division of Viral Diseases, Centers for Dieases Control and Prevention | Krista Queen, Anna Uehara, Jing Zhang, Yan Li, Ying Tao, Clinton R. Paden, Haibin Wang, Shifaq Kamili, Xiaoyan Lu, Brian Lynch, Senthil Kumar K. Sakthivel, Brett L. Whitaker, Lijuan Wang, Janna' R. Murray, Susan I. Gerber, Stephen Lindstrom, Suxiang Tong |
| 89 | EPI_ISL_411956 | hCoV-19/USA/TX1/2020 | *Homo sapiens* | North America / USA / Texas | 2020-02-11 | Texas Department of State Health Services | Pathogen Discovery, Respiratory Viruses Branch, Division of Viral Diseases, Centers for Dieases Control and Prevention | Krista Queen, Anna Uehara, Jing Zhang, Yan Li, Ying Tao, Clinton R. Paden, Haibin Wang, Shifaq Kamili, Xiaoyan Lu, Brian Lynch, Senthil Kumar K. Sakthivel, Brett L. Whitaker, Lijuan Wang, Janna' R. Murray, Susan I. Gerber, Stephen Lindstrom, Suxiang Tong |
| 90 | EPI_ISL_411957 | hCoV-19/China/WH-09/2020 | *Homo sapiens* | Asia / China | 2020-01-08 | unknown | Key Laboratory of Homo sapiens Diseases, Comparative Medicine, Institute of Laboratory Animal Science | Linlin,B., Lili,R., Shuran,G., Jiangning,L., Feifei,Q., Qi,L., Fengdi,L., Jing,X., Wei,D., Pin,Y., Yanfeng,X., Yajin,Q., Hong,G., Qiang,W., Mingya,L., Guanpeng,W., Shunyi,W., Zhiqi,S., Li,G., Lan,C., Conghui,W., Ying,W., Xinming,W., Yan,X., Qi,J. and Chuan,Q. |
| 91 | EPI_ISL_412029 | hCoV-19/Hong Kong/VB20024950/2020 | *Homo sapiens* | Asia / Hong Kong | 2020-01-30 | Hong Kong Department of Health | The University of Hong Kong | Dominic N.C. Tsang, Daniel K.W. Chu, Leo L.M. Poon, Malik Peiris |
| 92 | EPI_ISL_412030 | hCoV-19/Hong Kong/VB20026565/2020 | *Homo sapiens* | Asia / Hong Kong | 2020-02-01 | Hong Kong Department of Health | School of Public Health, The University of Hon g Kong | Dominic N.C. Tsang, Daniel K.W. Chu, Leo L.M. Poon, Malik Peiris |

| | | | | | | | | |
|---|---|---|---|---|---|---|---|---|
| 93 | EPI_ISL_412459 | hCoV-19/Jingzhou/HBCDC-HB-01/2020 | *Homo sapiens* | Asia / China / Hubei / Jingzhou | 2020-01-08 | Jingzhou Center for Disease Control and Prevention | Hubei Provincial Center for Disease Control and Prevention | Bin Fang, Xiang Li, Xiao Yu, Linlin Liu, Bo Yang, Faxian Zhan, Guojun Ye, Xixiang Huo, Junqiang Xu, Bo Yu, Kun Cai, Jing Li, Maoyi Chen,Jie Hu, Chunlin Mao, Yongzhong Jiang. |
| 94 | EPI_ISL_412862 | hCoV-19/USA/CA9/2020 | *Homo sapiens* | North America / USA / California / Solano | 2020-02-23 | California Department of Public Health | Pathogen Discovery, Respiratory Viruses Branch, Division of Viral Diseases, Centers for Disease Control and Prevention | Krista Queen, Anna Uehara, Jing Zhang, Yan Li, Ying Tao, Clinton R. Paden, Haibin Wang, Shifaq Kamili, Xiaoyan Lu, Brian Lynch, Senthil Kumar K. Sakthivel, Brett L. Whitaker, Lijuan Wang, Janna' R. Murray, Jasmine Padilla, Justin Lee, Susan I. Gerber, Stephen Lindstrom, Suxiang Tong |
| 95 | EPI_ISL_412898 | hCoV-19/Wuhan/HBCDC-HB-02/2019 | *Homo sapiens* | Asia / China / Hubei / Wuhan | 2019-12-30 | Wuhan Jinyintan Hospital | Hubei Provincial Center for Disease Control and Prevention | Bin Fang, Xiang Li, Xiao Yu, Linlin Liu, Bo Yang, Faxian Zhan, Guojun Ye, Xixiang Huo, Junqiang Xu, Bo Yu, Kun Cai, Jing Li, Yongzhong Jiang. |
| 96 | EPI_ISL_412966 | hCoV-19/China/IQTC01/2020 | *Homo sapiens* | Asia / China / Guangdong / Guangzhou | 2020-02-05 | unknown | Technology Centre, Guangzhou Customs | Shi,Y., Sun,J., Zheng,K., Huang,J. and Zhao,J. |
| 97 | EPI_ISL_412967 | hCoV-19/China/IQTC02/2020 | *Homo sapiens* | Asia / China / Guangzhou | 2020-01-29 | unknown | Technology Centre, Guangzhou Customs | Shi,Y., Zheng,K., Sun,J., Huang,J., Zhu,A., Zhuang,Z., Dai,J., Chen,Z., Sun,F., Zhang,Z., Li,X. and Wang,Y. |
| 98 | EPI_ISL_412969 | hCoV-19/Japan/Hu_DP_Kng_19-027/2020 | *Homo sapiens* | Asia / Japan | 2020-02-10 | unknown | Takayuki Hishiki Kanagawa Prefectural Institute of Public Health, Department of Microbiology | Hishiki,T., Suzuki,R., Sakuragi,J., Usui,K., Tanaka,Y., Kawai,J., Kogo,Y., Matsuki,Y., An,T., Hayashizaki,Y. and Takasaki,T. |
| 99 | EPI_ISL_412970 | hCoV-19/USA/WA2/2020 | *Homo sapiens* | North America / USA / Washington / Snohomish County | 2020-02-24 | Washington State Department of Health | Seattle Flu Study | Helen Chu, Michael Boeckh, Janet Englund, Michael Famulare, Barry Lutz, Deborah Nickerson, Mark Rieder, Lea Starita, Matthew Thompson, Jay Shendure, and Trevor Bedford |
| 99 | EPI_ISL_416462 | hCoV-19/USA/WA-S7/2020 | *Homo sapiens* | North America / USA / Washington | 2020-02-24 | Seattle Flu Study | Seattle Flu Study | Chu et al |
| 99 | EPI_ISL_417095 | hCoV-19/USA/WA-S42/2020 | *Homo sapiens* | North America / USA / Washington / King County | 2020-02-28 | Washington State Department of Health | Seattle Flu Study | Chu etl al |
| 99 | EPI_ISL_417100 | hCoV-19/USA/WA-S47/2020 | *Homo sapiens* | North America / USA / Washington / Snohomish County | 2020-02-29 | Washington State Department of Health | Seattle Flu Study | Chu etl al |
| 100 | EPI_ISL_412972 | hCoV-19/Mexico/CDMX-InDRE_01/2020 | *Homo sapiens* | North America / Mexico / Mexico City | 2020-02-27 | Instituto Nacional de Enfermedades Respiratorias | Instituto de Diagnostico y Referencia Epidemiologicos (INDRE) | Ramirez-Gonzalez Ernesto, Garces-Ayala Fabiola, Araiza-Rodriguez Adnan, Mendieta-Condado Edgar, Rodriguez-Maldonado Abril, Wong-Arambula Claudia, Vazquez-Perez Joel, Martinez Arturo, Boukadida Celia, Munoz-Medina Esteban, Sanchez Alejandro, Isa Pavel, Taboada Blanca, Lopez Susana, Arias Carlos, Barrera-Badillo Gisela, Hernandez-Rivas Lucia, Lopez-Martinez Irma |
| 100 | EPI_ISL_416428 | hCoV-19/Vietnam/39607/2020 | *Homo sapiens* | Asia / Vietnam / Quangning | 2020-03-07 | National Influenza Center, National Institute of Hygiene and Epidemiology (NIHE) | National Influenza Center, National Institute of Hygiene and Epidemiology (NIHE) | Le Quynh Mai, Taichiro Takemura, Meng Ling Moi, Takeshi Nabeshima, Nguyen Le Khanh Hang, Hoang Vu Mai Phuong, Ung Thi Hong Trang, Le Thi Thanh, Nguyen Vu Son, Vuong Duc Cuong, Pham Thi Hien, Tran Thu Huong, Nguyen Phuong Anh, Pham Hong Quynh Anh, Kouichi Morita, Futoshi Hasebe, Dang Duc Anh |
| 100 | EPI_ISL_416430 | hCoV-19/Vietnam/CM295/2020 | *Homo sapiens* | Asia / Vietnam / Hanoi | 2020-03-06 | National Influenza Center, National Institute of Hygiene and Epidemiology (NIHE) | National Influenza Center, National Institute of Hygiene and Epidemiology (NIHE) | Le Quynh Mai, Taichiro Takemura, Meng Ling Moi, Takeshi Nabeshima, Nguyen Le Khanh Hang, Hoang Vu Mai Phuong, Ung Thi Hong Trang, Le Thi Thanh, Nguyen Vu Son, Vuong Duc Cuong, Pham Thi Hien, Tran Thu Huong, Nguyen Phuong Anh, Pham Hong Quynh Anh, Kouichi Morita, Futoshi Hasebe, Dang Duc Anh |

| | | | | | | | | |
|---|---|---|---|---|---|---|---|---|
| 100 | EPI_ISL_416431 | hCoV-19/Vietnam/CM296/2020 | *Homo sapiens* | Asia / Vietnam / Hanoi | 2020-03-06 | National Influenza Center, National Institute of Hygiene and Epidemiology (NIHE) | National Influenza Center, National Institute of Hygiene and Epidemiology (NIHE) | Le Quynh Mai, Taichiro Takemura, Meng Ling Moi, Takeshi Nabeshima, Nguyen Le Khanh Hang, Hoang Vu Mai Phuong, Ung Thi Hong Trang, Le Thi Thanh, Nguyen Vu Son, Vuong Duc Cuong, Pham Thi Hien, Tran Thu Huong, Nguyen Phuong Anh, Pham Hong Quynh Anh, Kouichi Morita, Futoshi Hasebe, Dang Duc Anh |
| 101 | EPI_ISL_412973 | hCoV-19/Italy/CDG1/2020 | *Homo sapiens* | Europe / Italy / Lombardy | 2020-02-20 | Department of Infectious Diseases, Istituto Superiore di Sanità, Roma , Italy | Virology Laboratory, Scientific Department, Army Medical Center | Paola Stefanelli, Stefano Fiore, Antonella Marchi, Eleonora Benedetti, Concetta Fabiani, Giovanni Faggioni, Antonella Fortunato, Riccardo De Santis, Silvia Fillo, Anna Anselmo, Andrea Ciammaruconi, Stefano Palomba, Florigio Lista |
| 102 | EPI_ISL_412974 | hCoV-19/Italy/SPL1/2020 | *Homo sapiens* | Europe / Italy / Rome | 2020-01-29 | Department of Infectious Diseases, Istituto Superiore di Sanità, Rome, Italy | Virology Laboratory, Scientific Department, Army Medical Center | Paola Stefanelli, Stefano Fiore, Antonella Marchi, Eleonora Benedetti, Concetta Fabiani, Giovanni Faggioni, Antonella Fortunato, Silvia Fillo, Riccardo De Santis, Andrea Ciammaruconi, Giancarlo Petralito, Filippo Molinari, Florigio Lista |
| 103 | EPI_ISL_412975 | hCoV-19/Australia/NSW05/2020 | *Homo sapiens* | Oceania / Australia / New South Wales / Sydney | 2020-02-28 | Centre for Infectious Diseases and Microbiology Laboratory Services | NSW Health Pathology - Institute of Clinical Pathology and Medical Research; Westmead Hospital; University of Sydney | Eden J-S, Carter I, Rahman H, Holmes EC, Rockett R, O'Sullivan MV, Sintchenko V, Chen SC, Maddocks S, Kok J and Dwyer DE for the 2019-nCoV Study Group |
| 104 | EPI_ISL_412978 | hCoV-19/Wuhan/HBCDC-HB-02/2020 | *Homo sapiens* | Asia / China / Hubei / Wuhan | 2020-01-17 | The Central Hospital Of Wuhan | Hubei Provincial Center for Disease Control and Prevention | Bin Fang, Xiang Li, Xiao Yu, Linlin Liu, Bo Yang, Faxian Zhan, Guojun Ye, Xixiang Huo, Junqiang Xu, Bo Yu, Kun Cai, Jing Li, Yongzhong Jiang. |
| 105 | EPI_ISL_412979 | hCoV-19/Wuhan/HBCDC-HB-03/2020 | *Homo sapiens* | Asia / China / Hubei / Wuhan | 2020-01-18 | Union Hospital of Tongji Medical College, Huazhong University of Science and Technology | Hubei Provincial Center for Disease Control and Prevention | Bin Fang, Xiang Li, Xiao Yu, Linlin Liu, Bo Yang, Faxian Zhan, Guojun Ye, Xixiang Huo, Junqiang Xu, Bo Yu, Kun Cai, Jing Li, Yongzhong Jiang. |
| 106 | EPI_ISL_412980 | hCoV-19/Wuhan/HBCDC-HB-04/2020 | *Homo sapiens* | Asia / China / Hubei / Wuhan | 2020-01-18 | Union Hospital of Tongji Medical College, Huazhong University of Science and Technology | Hubei Provincial Center for Disease Control and Prevention | Bin Fang, Xiang Li, Xiao Yu, Linlin Liu, Bo Yang, Faxian Zhan, Guojun Ye, Xixiang Huo, Junqiang Xu, Bo Yu, Kun Cai, Jing Li, Yongzhong Jiang. |
| 107 | EPI_ISL_412982 | hCoV-19/Wuhan/HBCDC-HB-06/2020 | *Homo sapiens* | Asia / China / Hubei / Wuhan | 2020-02-07 | Wuhan Lung Hospital | Hubei Provincial Center for Disease Control and Prevention | Bin Fang, Xiang Li, Xiao Yu, Linlin Liu, Bo Yang, Faxian Zhan, Guojun Ye, Xixiang Huo, Junqiang Xu, Bo Yu, Kun Cai, Jing Li, Yongzhong Jiang. |
| 108 | EPI_ISL_413016 | hCoV-19/Brazil/SPBR-02/2020 | *Homo sapiens* | South America / Brazil / Sao Paulo / Sao Paulo | 2020-02-28 | Hospital Israelita Albert Einstein | Instituto Adolfo Lutz, Interdisciplinary Procedures Center, Strategic Laboratory | Jaqueline Goes de Jesus, Claudio Tavares Sacchi, Fabiana Cristina Pereira dos Santos, Ingra Morales Claro, Flávia Cristina da Silva Sales, Claudia Regina Gonçalves, Joshua Quick, Maria do Carmo Sampaio Tavares Timenetsky, Nicholas James Loman, Andrew Rambaut, Ester Cerdeira Sabino, Nuno Rodrigues Faria |
| 109 | EPI_ISL_413017 | hCoV-19/South Korea/KUMC01/2020 | *Homo sapiens* | Asia / South Korea | 2020-02-06 | Department of Microbiology, Institute for Viral Diseases, College of Medicine, Korea University | Department of Microbiology, Institute for Viral Diseases, College of Medicine, Korea University | Changmin Kang, Joon-Yong Bae, Jungmin Lee, Heedo Park, Juyoung Cho, Jeonghun Kim, Gee eun Lee, Cui Chunguang, Kyeong-ryeol Shin, Dong Min Kim, Jin Il Kim, Man-Seong Park |
| 110 | EPI_ISL_413018 | hCoV-19/South Korea/KUMC02/2020 | *Homo sapiens* | Asia / South Korea | 2020-02-06 | Department of Microbiology, Institute for Viral Diseases, College of Medicine, Korea University | Department of Microbiology, Institute for Viral Diseases, College of Medicine, Korea University | Changmin Kang, Joon-Yong Bae, Jungmin Lee, Heedo Park, Juyoung Cho, Jeonghun Kim, Gee eun Lee, Cui Chunguang, Kyeong-ryeol Shin, Dong Min Kim, Jin Il Kim, Man-Seong Park |
| 111 | EPI_ISL_413023 | hCoV-19/Switzerland/1000477797/2020 | *Homo sapiens* | Europe / Switzerland / Zurich | 2020-02-29 | Division of Infectious Diseases, University Hospital Zurich | Institute of Medical Virology, University of Zurich | Stefan Schmutz, Maryam Zaheri, Verena Kufner, Gabriela Ziltener, Patrick Redli, Fiona Steiner, Jon Huder, Riccarda Capaul, Andrea Zbinden, Jürg Böni, Michael Huber, Roberto Speck, Alexandra Trkola |
| 112 | EPI_ISL_413025 | hCoV-19/USA/WA3-UW1/2020 | *Homo sapiens* | North America / USA / Washington | 2020-02-27 | Harborview Medical Center | UW Virology Lab | Pavitra Roychoudhury, Arun Nalla, Hong Xie, Keith Jerome, Alexander Greninger |
| 113 | EPI_ISL_413213 | hCoV-19/Australia/NSW06/2020 | *Homo sapiens* | Oceania / Australia / New South Wales / Sydney | 2020-02-29 | Centre for Infectious Diseases and Microbiology Laboratory Services | NSW Health Pathology - Institute of Clinical Pathology and Medical Research; Westmead Hospital; University of Sydney | Eden J-S, Carter I, Rahman H, Holmes EC, Rockett R, O'Sullivan MV, Sintchenko V, Chen SC, Maddocks S, Kok J and Dwyer DE for the 2019-nCoV Study Group* |

| | | | | | | | | |
|---|---|---|---|---|---|---|---|---|
| 114 | EPI_ISL_413214 | hCoV-19/Australia/NSW07/2020 | *Homo sapiens* | Oceania / Australia / New South Wales / Sydney | 2020-02-29 | Centre for Infectious Diseases and Microbiology Laboratory Services | NSW Health Pathology - Institute of Clinical Pathology and Medical Research; Westmead Hospital; University of Sydney | Eden J-S, Carter I, Rahman H, Holmes EC, Rockett R, O'Sullivan MV, Sintchenko V, Chen SC, Maddocks S, Kok J and Dwyer DE for the 2019-nCoV Study Group* |
| 115 | EPI_ISL_413455 | hCoV-19/USA/WA4-UW2/2020 | *Homo sapiens* | North America / USA / Washington | 2020-02-28 | Washington State Public Health Lab | University of Washington Virology Lab | Pavitra Roychoudhury, Arun Nalla, Hong Xie, Keith Jerome, Alexander Greninger |
| 116 | EPI_ISL_413456 | hCoV-19/USA/WA-S2/2020 | *Homo sapiens* | North America / USA / Washington / King County | 2020-02-20 | Seattle Flu Study | Seattle Flu Study | Chu et al |
| 116 | EPI_ISL_413457 | hCoV-19/USA/WA6-UW3/2020 | *Homo sapiens* | North America / USA / Washington | 2020-02-29 | Washington State Public Health Lab | UW Virology Lab | Pavitra Roychoudhury, Arun Nalla, Hong Xie, Keith Jerome, Alexander Greninger |
| 116 | EPI_ISL_413560 | hCoV-19/USA/WA-S3/2020 | *Homo sapiens* | North America / USA / Washington | 2020-02-28 | Seattle Flu Study | Seattle Flu Study | Chu et al |
| 116 | EPI_ISL_414366 | hCoV-19/USA/WA-UW18/2020 | *Homo sapiens* | North America / USA / Washington | 2020-03-05 | UW Virology Lab | UW Virology Lab | Pavitra Roychoudhury, Hong Xie, Keith Jerome, Alexander Greninger |
| 116 | EPI_ISL_415541 | hCoV-19/USA/UPHL-03/2020 | *Homo sapiens* | North America / USA / Utah | 2020-03-13 | Utah Public Health Laboratory | Utah Public Health Laboratory | Erin Young, Kelly Oakeson |
| 116 | EPI_ISL_415542 | hCoV-19/USA/UPHL-04/2020 | *Homo sapiens* | North America / USA / Utah | 2020-03-13 | Utah Public Health Laboratory | Utah Public Health Laboratory | Erin Young, Kelly Oakeson |
| 116 | EPI_ISL_416460 | hCoV-19/USA/WA-S5/2020 | *Homo sapiens* | North America / USA / Washington / King County | 2020-02-29 | Seattle Flu Study | Seattle Flu Study | Chu et al |
| 116 | EPI_ISL_416461 | hCoV-19/USA/WA-S6/2020 | *Homo sapiens* | North America / USA / Washington / King County | 2020-02-29 | Seattle Flu Study | Seattle Flu Study | Chu et al |
| 116 | EPI_ISL_416465 | hCoV-19/USA/WA-S10/2020 | *Homo sapiens* | North America / USA / Washington / King County | 2020-02-29 | Seattle Flu Study | Seattle Flu Study | Chu et al |
| 116 | EPI_ISL_416643 | hCoV-19/USA/WA-UW105/2020 | *Homo sapiens* | North America / USA / Washington | 2020-03-11 | UW Virology Lab | UW Virology Lab | Pavitra Roychoudhury, Hong Xie, Keith Jerome, Alexander Greninger |
| 116 | EPI_ISL_417077 | hCoV-19/USA/WA-S24/2020 | *Homo sapiens* | North America / USA / Washington / Snohomish County | 2020-03-02 | Washington State Department of Health | Seattle Flu Study | Chu etl al |
| 116 | EPI_ISL_417091 | hCoV-19/USA/WA-S38/2020 | *Homo sapiens* | North America / USA / Washington / Snohomish County | 2020-03-04 | Washington State Department of Health | Seattle Flu Study | Chu etl al |
| 116 | EPI_ISL_417105 | hCoV-19/USA/WA-S52/2020 | *Homo sapiens* | North America / USA / Washington / Snohomish County | 2020-03-03 | Washington State Department of Health | Seattle Flu Study | Chu etl al |
| 116 | EPI_ISL_417108 | hCoV-19/USA/WA-S55/2020 | *Homo sapiens* | North America / USA / Washington / King County | 2020-02-29 | Washington State Department of Health | Seattle Flu Study | Chu etl al |

| | | | | | | | | |
|---|---|---|---|---|---|---|---|---|
| 116 | EPI_ISL_417124 | hCoV-19/USA/WA-S71/2020 | *Homo sapiens* | North America / USA / Washington / King County | 2020-03-05 | Washington State Department of Health | Seattle Flu Study | Chu etl al |
| 116 | EPI_ISL_417128 | hCoV-19/USA/WA-S75/2020 | *Homo sapiens* | North America / USA / Washington / King County | 2020-03-05 | Washington State Department of Health | Seattle Flu Study | Chu etl al |
| 116 | EPI_ISL_417135 | hCoV-19/USA/WA-S82/2020 | *Homo sapiens* | North America / USA / Washington / King County | 2020-02-22 | Washington State Department of Health | Seattle Flu Study | Chu etl al |
| 116 | EPI_ISL_417141 | hCoV-19/USA/WA-S88/2020 | *Homo sapiens* | North America / USA / Washington / King County | 2020-03-01 | Washington State Department of Health | Seattle Flu Study | Chu etl al |
| 116 | EPI_ISL_417144 | hCoV-19/USA/WA-S91/2020 | *Homo sapiens* | North America / USA / Washington / Snohomish County | 2020-03-02 | Washington State Department of Health | Seattle Flu Study | Chu etl al |
| 116 | EPI_ISL_417154 | hCoV-19/USA/WA-S101/2020 | *Homo sapiens* | North America / USA / Washington / King County | 2020-02-28 | Washington State Department of Health | Seattle Flu Study | Chu etl al |
| 116 | EPI_ISL_417156 | hCoV-19/USA/WA-S103/2020 | *Homo sapiens* | North America / USA / Washington / King County | 2020-02-28 | Washington State Department of Health | Seattle Flu Study | Chu etl al |
| 116 | EPI_ISL_417169 | hCoV-19/USA/WA-S116/2020 | *Homo sapiens* | North America / USA / Washington | 2020-03-02 | Washington State Department of Health | Seattle Flu Study | Chu etl al |
| 117 | EPI_ISL_413458 | hCoV-19/USA/WA7-UW4/2020 | *Homo sapiens* | North America / USA / Washington | 2020-03-01 | Washington State Public Health Lab | UW Virology Lab | Pavitra Roychoudhury, Arun Nalla, Hong Xie, Keith Jerome, Alexander Greninger |
| 118 | EPI_ISL_413489 | hCoV-19/Italy/UniSR1/2020 | *Homo sapiens* | Europe / Italy / Lombardy / Milan | 2020-03-03 | Laboratorio di Microbiologia e Virologia, Università Vita-Salute San Raffaele, Milano | Laboratorio di Microbiologia e Virologia, Università Vita-Salute San Raffaele, Milano | R.A Diotti, E. Criscuolo, M. Castelli, V. Caputo, R. Ferrarese, M. Sampaolo, E. Boeri, I. Negri, V. Amato, G. Lo Raso, C. Di Resta, R. Burioni, M. Clementi, N. Mancini & N. Clementi |
| 119 | EPI_ISL_413513 | hCoV-19/South Korea/KUMC03/2020 | *Homo sapiens* | Asia / South Korea | 2020-02-27 | Division of Infectious Diseases, Department of Internal Medicine, Korea University College of Medicine | Department of Microbiology, Institute for Viral Diseases, College of Medicine, Korea University | Changmin Kang, Joon-Yong Bae, Jungmin Lee, Jin Gu Yoon, Heedo Park, Juyoung Cho, Jeonghun Kim, Gee Eun Lee, Cui Chunguang, Kyeong-ryeol Shin, Ji Yun Noh, Joon Young Song, Hee Jin Cheong, Woo Joo Kim, Jin Il Kim, Man-Seong Park |
| 119 | EPI_ISL_413514 | hCoV-19/South Korea/KUMC04/2020 | *Homo sapiens* | Asia / South Korea | 2020-02-27 | Department of Microbiology, Institute for Viral Diseases, College of Medicine, Korea University | Department of Microbiology, Institute for Viral Diseases, College of Medicine, Korea University | Changmin Kang, Joon-Yong Bae, Jungmin Lee, Jin Gu Yoon, Heedo Park, Juyoung Cho, Jeonghun Kim, Gee Eun Lee, Cui Chunguang, Kyeong-ryeol Shin, Ji Yun Noh, Joon Young Song, Hee Jin Cheong, Woo Joo Kim, Jin Il Kim, Man-Seong Park |
| 119 | EPI_ISL_413515 | hCoV-19/South Korea/KUMC05/2020 | *Homo sapiens* | Asia / South Korea | 2020-02-27 | Division of Infectious Diseases, Department of Internal Medicine, Korea University College of Medicine | Department of Microbiology, Institute for Viral Diseases, College of Medicine, Korea University | Changmin Kang, Joon-Yong Bae, Jungmin Lee, Jin Gu Yoon, Heedo Park, Juyoung Cho, Jeonghun Kim, Gee Eun Lee, Cui Chunguang, Kyeong-ryeol Shin, Ji Yun Noh, Joon Young Song, Hee Jin Cheong, Woo Joo Kim, Jin Il Kim, Man-Seong Park |
| 119 | EPI_ISL_413516 | hCoV-19/South Korea/KUMC06/2020 | *Homo sapiens* | Asia / South Korea | 2020-02-27 | Department of Microbiology, Institute for Viral Diseases, College of Medicine, Korea University | Department of Microbiology, Institute for Viral Diseases, College of Medicine, Korea University | Changmin Kang, Joon-Yong Bae, Jungmin Lee, Jin Gu Yoon, Heedo Park, Juyoung Cho, Jeonghun Kim, Gee Eun Lee, Cui Chunguang, Kyeong-ryeol Shin, Ji Yun Noh, Joon Young Song, Hee Jin Cheong, Woo Joo Kim, Jin Il Kim, Man-Seong Park |
| 119 | EPI_ISL_413518 | hCoV-19/Beijing/105/2020 | *Homo sapiens* | Asia / China / Beijing | 2020-01-26 | unknown | Infectious Disease Control Center | Li,J., Li,L., Li,Z., Qiu,S., Song,H., Li,P. and Li,P. |

| | | | | | | | | |
|---|---|---|---|---|---|---|---|---|
| 119 | EPI_ISL_413519 | hCoV-19/Beijing/231/2020 | *Homo sapiens* | Asia / China / Beijing | 2020-01-28 | unknown | Infectious Disease Control Center | Li,J., Li,L., Li,Z., Qiu,S., Song,H., Li,P. and Li,P. |
| 119 | EPI_ISL_413521 | hCoV-19/Beijing/235/2020 | *Homo sapiens* | Asia / China / Beijing | 2020-01-28 | unknown | Infectious Disease Control Center | Li,J., Li,L., Li,Z., Qiu,S., Song,H., Li,P. and Li,P. |
| 120 | EPI_ISL_413520 | hCoV-19/Beijing/233/2020 | *Homo sapiens* | Asia / China / Beijing | 2020-01-28 | unknown | Infectious Disease Control Center | Li,J., Li,L., Li,Z., Qiu,S., Song,H., Li,P. and Li,P. |
| 121 | EPI_ISL_413523 | hCoV-19/India/1-31/2020 | *Homo sapiens* | Asia / India / Kerala | 2020-01-31 | Indian Council of Medical Research-National Institute of Virology | National Influenza Center, Indian Council of Medical Research-National Institute of Virology | Potdar V, Yadav PD, Choudhary ML, Shete-Aich A |
| 122 | EPI_ISL_413555 | hCoV-19/Wales/PHW1/2020 | *Homo sapiens* | Europe / United Kingdom / Wales | 2020-02-27 | Wales Specialist Virology Centre | Public Health Wales Microbiology Cardiff | Catherine Moore, Cen Sabu, Joanne Watkins, Sally Corden, Tom Connor |
| 122 | EPI_ISL_414587 | hCoV-19/Ireland/Limerick-19935/2020 | *Homo sapiens* | Europe / Ireland / Limerick | 2020-03-03 | UCD National Virus Reference Laboratory | UCD National Virus Reference Laboratory | Michael Carr, Gabriel Gonzalez, Jonathan Dean, Suzie Coughlan, Alison Murphy, Kevin Byrne, Ken Wolfe, Jeff Connell, Brendan Loftus, Cillian F De Gascun |
| 123 | EPI_ISL_413558 | hCoV-19/USA/CA-CDPH-UC2/2020 | *Homo sapiens* | North America / USA / California / Solano County | 2020-02-27 | California Department of Public Health | Chiu Laboratory, UCSF-Abbott Viral Diagnostics and Discovery Center, University of California, San Francisco | Xianding Deng, Scot Federman, Guixia Yu, Chao-Yang Pan, Hugo Guevara, Alicia Sotomayor-Gonzalez, Allan Gopez, Wei Gu, Steve Miller, Debra A. Wadford, and Charles Y. Chiu |
| 124 | EPI_ISL_413559 | hCoV-19/USA/CA-CDPH-UC3/2020 | *Homo sapiens* | North America / USA / California / Solano County | 2020-02-27 | California Department of Public Health | Chiu Laboratory, UCSF-Abbott Viral Diagnostics and Discovery Center, University of California, San Francisco | Xianding Deng, Scot Federman, Guixia Yu, Chao-Yang Pan, Hugo Guevara, Alicia Sotomayor-Gonzalez, Allan Gopez, Wei Gu, Steve Miller, Debra A. Wadford, and Charles Y. Chiu |
| 125 | EPI_ISL_413563 | hCoV-19/USA/WA12-UW8/2020 | *Homo sapiens* | North America / USA / Washington | 2020-03-03 | UW Virology Lab | UW Virology Lab | Pavitra Roychoudhury, Hong Xie, Keith Jerome, Alexander Greninger |
| 126 | EPI_ISL_413566 | hCoV-19/Netherlands/Blaricum_1364780/2020 | *Homo sapiens* | Europe / Netherlands / Blaricum | 2020-03-02 | MHC Gooi & Vechtstreek | Erasmus Medical Center | David Nieuwenhuijse, Bas Oude Munnink, Reina Sikkema, Claudia Schapendonk, Irina Chestakova, Anne van der Linden, Mark Pronk, Pascal Lexmond, Corien Swaan, Manon Haverkate, Madelief Mollers, Mart Stein, Sandra Kengne Kamga Mobou, Jeroen van Kampen, Jolanda Voermans, Aura Timen, Corine GeurtsvanKessel, Annemiek van der Eijk, Richard Molenkamp, Marion Koopmans, on behalf of the Dutch national COVID-19 response team. |
| 126 | EPI_ISL_413591 | hCoV-19/Netherlands/Zeewolde_1365080/2020 | *Homo sapiens* | Europe / Netherlands / Zeewolde | 2020-03-02 | MHC Flevoland | Erasmus Medical Center | David Nieuwenhuijse, Bas Oude Munnink, Reina Sikkema, Claudia Schapendonk, Irina Chestakova, Anne van der Linden, Mark Pronk, Pascal Lexmond, Corien Swaan, Manon Haverkate, Madelief Mollers, Mart Stein, Sandra Kengne Kamga Mobou, Jeroen van Kampen, Jolanda Voermans, Aura Timen, Corine GeurtsvanKessel, Annemiek van der Eijk, Richard Molenkamp, Marion Koopmans, on behalf of the Dutch national COVID-19 response team. |
| 126 | EPI_ISL_414443 | hCoV-19/Netherlands/Utrecht_11/2020 | *Homo sapiens* | Europe / Netherlands / Utrecht | 2020-03-03 | Dutch COVID-19 response team | Erasmus Medical Center | David Nieuwenhuijse, Bas Oude Munnink, Reina Sikkema, Claudia Schapendonk, Irina Chestakova, Anne van der Linden, Mark Pronk, Pascal Lexmond, Corien Swaan, Manon Haverkate, Madelief Mollers, Mart Stein, Sandra Kengne Kamga Mobou, Jeroen van Kampen, Jolanda Voermans, Aura Timen, Corine GeurtsvanKessel, Annemiek van der Eijk, Richard Molenkamp, Marion Koopmans, on behalf of the Dutch national COVID-19 response team. |
| 127 | EPI_ISL_413572 | hCoV-19/Netherlands/Haarlem_1363688/2020 | *Homo sapiens* | Europe / Netherlands / Haarlem | 2020-03-01 | MHC Kennemerland | Erasmus Medical Center | David Nieuwenhuijse, Bas Oude Munnink, Reina Sikkema, Claudia Schapendonk, Irina Chestakova, Anne van der Linden, Mark Pronk, Pascal Lexmond, Corien Swaan, Manon Haverkate, Madelief Mollers, Mart Stein, Sandra Kengne Kamga Mobou, Jeroen van Kampen, Jolanda Voermans, Aura Timen, Corine GeurtsvanKessel, Annemiek van der Eijk, Richard Molenkamp, Marion Koopmans, on behalf of the Dutch national COVID-19 response team. |

| | | | | | | | | |
|---|---|---|---|---|---|---|---|---|
| 128 | EPI_ISL_413579 | hCoV-19/Netherlands/Nootdorp_1364222/2020 | *Homo sapiens* | Europe / Netherlands / Nootdorp | 2020-03-03 | MHC Haaglanden | Erasmus Medical Center | David Nieuwenhuijse, Bas Oude Munnink, Reina Sikkema, Claudia Schapendonk, Irina Chestakova, Anne van der Linden, Mark Pronk, Pascal Lexmond, Corien Swaan, Manon Haverkate, Madelief Mollers, Mart Stein, Sandra Kengne Kamga Mobou, Jeroen van Kampen, Jolanda Voermans, Aura Timen, Corine GeurtsvanKessel, Annemiek van der Eijk, Richard Molenkamp, Marion Koopmans, on behalf of the Dutch national COVID-19 response team. |
| 128 | EPI_ISL_413587 | hCoV-19/Netherlands/Tilburg_1364286/2020 | *Homo sapiens* | Europe / Netherlands / Tilburg | 2020-03-03 | Foundation Elisabeth-Tweesteden Ziekenhuis | Erasmus Medical Center | David Nieuwenhuijse, Bas Oude Munnink, Reina Sikkema, Claudia Schapendonk, Irina Chestakova, Anne van der Linden, Mark Pronk, Pascal Lexmond, Corien Swaan, Manon Haverkate, Madelief Mollers, Mart Stein, Sandra Kengne Kamga Mobou, Jeroen van Kampen, Jolanda Voermans, Aura Timen, Corine GeurtsvanKessel, Annemiek van der Eijk, Richard Molenkamp, Marion Koopmans, on behalf of the Dutch national COVID-19 response team. |
| 129 | EPI_ISL_413584 | hCoV-19/Netherlands/Rotterdam_1364740/2020 | *Homo sapiens* | Europe / Netherlands / Rotterdam | 2020-03-03 | unknown | Erasmus Medical Center | David Nieuwenhuijse, Bas Oude Munnink, Reina Sikkema, Claudia Schapendonk, Irina Chestakova, Anne van der Linden, Mark Pronk, Pascal Lexmond, Corien Swaan, Manon Haverkate, Madelief Mollers, Mart Stein, Sandra Kengne Kamga Mobou, Jeroen van Kampen, Jolanda Voermans, Aura Timen, Corine GeurtsvanKessel, Annemiek van der Eijk, Richard Molenkamp, Marion Koopmans, on behalf of the Dutch national COVID-19 response team. |
| 129 | EPI_ISL_414423 | hCoV-19/Netherlands/Gelderland_1/2020 | *Homo sapiens* | Europe / Netherlands / Gelderland | 2020-03-02 | Dutch COVID-19 response team | Erasmus Medical Center | David Nieuwenhuijse, Bas Oude Munnink, Reina Sikkema, Claudia Schapendonk, Irina Chestakova, Anne van der Linden, Mark Pronk, Pascal Lexmond, Corien Swaan, Manon Haverkate, Madelief Mollers, Mart Stein, Sandra Kengne Kamga Mobou, Jeroen van Kampen, Jolanda Voermans, Aura Timen, Corine GeurtsvanKessel, Annemiek van der Eijk, Richard Molenkamp, Marion Koopmans, on behalf of the Dutch national COVID-19 response team. |
| 129 | EPI_ISL_414433 | hCoV-19/Netherlands/NoordHolland_1/2020 | *Homo sapiens* | Europe / Netherlands / Noord Holland | 2020-03-03 | Dutch COVID-19 response team | Erasmus Medical Center | David Nieuwenhuijse, Bas Oude Munnink, Reina Sikkema, Claudia Schapendonk, Irina Chestakova, Anne van der Linden, Mark Pronk, Pascal Lexmond, Corien Swaan, Manon Haverkate, Madelief Mollers, Mart Stein, Sandra Kengne Kamga Mobou, Jeroen van Kampen, Jolanda Voermans, Aura Timen, Corine GeurtsvanKessel, Annemiek van der Eijk, Richard Molenkamp, Marion Koopmans, on behalf of the Dutch national COVID-19 response team. |
| 129 | EPI_ISL_414439 | hCoV-19/Netherlands/Utrecht_5/2020 | *Homo sapiens* | Europe / Netherlands / Utrecht | 2020-03-02 | Dutch COVID-19 response team | Erasmus Medical Center | David Nieuwenhuijse, Bas Oude Munnink, Reina Sikkema, Claudia Schapendonk, Irina Chestakova, Anne van der Linden, Mark Pronk, Pascal Lexmond, Corien Swaan, Manon Haverkate, Madelief Mollers, Mart Stein, Sandra Kengne Kamga Mobou, Jeroen van Kampen, Jolanda Voermans, Aura Timen, Corine GeurtsvanKessel, Annemiek van der Eijk, Richard Molenkamp, Marion Koopmans, on behalf of the Dutch national COVID-19 response team. |
| 129 | EPI_ISL_414446 | hCoV-19/Netherlands/ZuidHolland_10/2020 | *Homo sapiens* | Europe / Netherlands / Zuid Holland | 2020-03-03 | Dutch COVID-19 response team | Erasmus Medical Center | David Nieuwenhuijse, Bas Oude Munnink, Reina Sikkema, Claudia Schapendonk, Irina Chestakova, Anne van der Linden, Mark Pronk, Pascal Lexmond, Corien Swaan, Manon Haverkate, Madelief Mollers, Mart Stein, Sandra Kengne Kamga Mobou, Jeroen van Kampen, Jolanda Voermans, Aura Timen, Corine GeurtsvanKessel, Annemiek van der Eijk, Richard Molenkamp, Marion Koopmans, on behalf of the Dutch national COVID-19 response team. |

| | | | | | | | | |
|---|---|---|---|---|---|---|---|---|
| 129 | EPI_ISL_415463 | hCoV-19/Netherlands/Gelderland_3/2020 | *Homo sapiens* | Europe / Netherlands / Gelderland | 2020-03-09 | Dutch COVID-19 response team | Erasmus Medical Center | David Nieuwenhuijse, Bas Oude Munnink, Reina Sikkema, Claudia Schapendonk, Irina Chestakova, Anne van der Linden, Mark Pronk, Pascal Lexmond, Corien Swaan, Manon Haverkate, Madelief Mollers, Mart Stein, Sandra Kengne Kamga Mobou, Jeroen van Kampen, Jolanda Voermans, Aura Timen, Corine GeurtsvanKessel, Annemiek van der Eijk, Richard Molenkamp, Marion Koopmans, on behalf of the Dutch national COVID-19 response team. |
| 129 | EPI_ISL_415496 | hCoV-19/Netherlands/NA_7/2020 | *Homo sapiens* | Europe / Netherlands | 2020-03-09 | Dutch COVID-19 response team | Erasmus Medical Center | David Nieuwenhuijse, Bas Oude Munnink, Reina Sikkema, Claudia Schapendonk, Irina Chestakova, Anne van der Linden, Mark Pronk, Pascal Lexmond, Corien Swaan, Manon Haverkate, Madelief Mollers, Mart Stein, Sandra Kengne Kamga Mobou, Jeroen van Kampen, Jolanda Voermans, Aura Timen, Corine GeurtsvanKessel, Annemiek van der Eijk, Richard Molenkamp, Marion Koopmans, on behalf of the Dutch national COVID-19 response team. |
| 129 | EPI_ISL_415502 | hCoV-19/Netherlands/NoordBrabant_45/2020 | *Homo sapiens* | Europe / Netherlands / Noord Brabant | 2020-03-09 | Dutch COVID-19 response team | Erasmus Medical Center | David Nieuwenhuijse, Bas Oude Munnink, Reina Sikkema, Claudia Schapendonk, Irina Chestakova, Anne van der Linden, Mark Pronk, Pascal Lexmond, Corien Swaan, Manon Haverkate, Madelief Mollers, Mart Stein, Sandra Kengne Kamga Mobou, Jeroen van Kampen, Jolanda Voermans, Aura Timen, Corine GeurtsvanKessel, Annemiek van der Eijk, Richard Molenkamp, Marion Koopmans, on behalf of the Dutch national COVID-19 response team. |
| 129 | EPI_ISL_415514 | hCoV-19/Netherlands/NoordBrabant_58/2020 | *Homo sapiens* | Europe / Netherlands / Noord Brabant | 2020-03-11 | Dutch COVID-19 response team | Erasmus Medical Center | David Nieuwenhuijse, Bas Oude Munnink, Reina Sikkema, Claudia Schapendonk, Irina Chestakova, Anne van der Linden, Mark Pronk, Pascal Lexmond, Corien Swaan, Manon Haverkate, Madelief Mollers, Mart Stein, Sandra Kengne Kamga Mobou, Jeroen van Kampen, Jolanda Voermans, Aura Timen, Corine GeurtsvanKessel, Annemiek van der Eijk, Richard Molenkamp, Marion Koopmans, on behalf of the Dutch national COVID-19 response team. |
| 129 | EPI_ISL_415523 | hCoV-19/Netherlands/NoordBrabant_67/2020 | *Homo sapiens* | Europe / Netherlands / Noord Brabant | 2020 | Dutch COVID-19 response team | Erasmus Medical Center | David Nieuwenhuijse, Bas Oude Munnink, Reina Sikkema, Claudia Schapendonk, Irina Chestakova, Anne van der Linden, Mark Pronk, Pascal Lexmond, Corien Swaan, Manon Haverkate, Madelief Mollers, Mart Stein, Sandra Kengne Kamga Mobou, Jeroen van Kampen, Jolanda Voermans, Aura Timen, Corine GeurtsvanKessel, Annemiek van der Eijk, Richard Molenkamp, Marion Koopmans, on behalf of the Dutch national COVID-19 response team. |
| 129 | EPI_ISL_415529 | hCoV-19/Netherlands/ZuidHolland_25/2020 | *Homo sapiens* | Europe / Netherlands / Zuid Holland | 2020-03-09 | Dutch COVID-19 response team | Erasmus Medical Center | David Nieuwenhuijse, Bas Oude Munnink, Reina Sikkema, Claudia Schapendonk, Irina Chestakova, Anne van der Linden, Mark Pronk, Pascal Lexmond, Corien Swaan, Manon Haverkate, Madelief Mollers, Mart Stein, Sandra Kengne Kamga Mobou, Jeroen van Kampen, Jolanda Voermans, Aura Timen, Corine GeurtsvanKessel, Annemiek van der Eijk, Richard Molenkamp, Marion Koopmans, on behalf of the Dutch national COVID-19 response team. |
| 129 | EPI_ISL_415533 | hCoV-19/Netherlands/ZuidHolland_29/2020 | *Homo sapiens* | Europe / Netherlands / Zuid Holland | 2020-03-09 | Dutch COVID-19 response team | Erasmus Medical Center | David Nieuwenhuijse, Bas Oude Munnink, Reina Sikkema, Claudia Schapendonk, Irina Chestakova, Anne van der Linden, Mark Pronk, Pascal Lexmond, Corien Swaan, Manon Haverkate, Madelief Mollers, Mart Stein, Sandra Kengne Kamga Mobou, Jeroen van Kampen, Jolanda Voermans, Aura Timen, Corine GeurtsvanKessel, Annemiek van der Eijk, Richard Molenkamp, Marion Koopmans, on behalf of the Dutch national COVID-19 response team. |

| | | | | | | | | |
|---|---|---|---|---|---|---|---|---|
| 130 | EPI_ISL_413589 | hCoV-19/Netherlands/Utrecht_1363628/2020 | *Homo sapiens* | Europe / Netherlands / Utrecht | 2020-03-01 | MHC Utrecht | Erasmus Medical Center | David Nieuwenhuijse, Bas Oude Munnink, Reina Sikkema, Claudia Schapendonk, Irina Chestakova, Anne van der Linden, Mark Pronk, Pascal Lexmond, Corien Swaan, Manon Haverkate, Madelief Mollers, Mart Stein, Sandra Kengne Kamga Mobou, Jeroen van Kampen, Jolanda Voermans, Aura Timen, Corine GeurtsvanKessel, Annemiek van der Eijk, Richard Molenkamp, Marion Koopmans, on behalf of the Dutch national COVID-19 response team. |
| 131 | EPI_ISL_413593 | hCoV-19/Luxembourg/Lux1/2020 | *Homo sapiens* | Europe / Luxembourg | 2020-02-29 | Laboratoire National de Santé | Erasmus Medical Center | David Nieuwenhuijse, Bas Oude Munnink, Reina Sikkema, Claudia Schapendonk, Irina Chestakova, Anne van der Linden, Mark Pronk, Pascal Lexmond, T. Abdelrahman, G. Fournier, J. Mossong, T. Nguyen, Jeroen van Kampen, Jolanda Voermans, Corine GeurtsvanKessel, Annemiek van der Eijk, Richard Molenkamp, Marion Koopmans, on behalf of the Dutch national COVID-19 response team. |
| 132 | EPI_ISL_413602 | hCoV-19/Finland/FIN03032020A/2020 | *Homo sapiens* | Europe / Finland / Helsinki | 2020-03-03 | Department of Virology and Immunology, University of Helsinki and Helsinki University Hospital, Huslab Finland | Department of Virology, Faculty of Medicine, University of Helsinki, Helsinki, Finland | Teemu Smura, Hannimari Kallio-Kokko, Olli Vapalahti |
| 133 | EPI_ISL_413603 | hCoV-19/Finland/FIN03032020B/2020 | *Homo sapiens* | Europe / Finland / Helsinki | 2020-03-03 | Department of Virology and Immunology, University of Helsinki and Helsinki University Hospital, Huslab Finland | Department of Virology, Faculty of Medicine, University of Helsinki, Helsinki, Finland | Teemu Smura, Hannimari Kallio-Kokko, Olli Vapalahti |
| 134 | EPI_ISL_413604 | hCoV-19/Finland/FIN03032020C/2020 | *Homo sapiens* | Europe / Finland / Helsinki | 2020-03-03 | Department of Virology and Immunology, University of Helsinki and Helsinki University Hospital, Huslab Finland | Department of Virology, Faculty of Medicine, University of Helsinki, Helsinki, Finland | Teemu Smura, Hannimari Kallio-Kokko, Olli Vapalahti |
| 135 | EPI_ISL_413606 | hCoV-19/USA/CruiseA-1/2020 | *Homo sapiens* | North America / USA | 2020-02-17 | unknown | Pathogen Discovery, Respiratory Viruses Branch, Division of Viral Diseases, Centers for Dieases Control and Prevention | Anna Uehara, Ying Tao, Clinton R. Paden, Krista Queen, Jing Zhang, Yan Li, Mary S. Keckler, Alison S Laufer Halpin, Haibin Wang, Jasmine Padilla, Justin Lee, Christopher A. Elkins, Susan I. Gerber, Suxiang Tong |
| 136 | EPI_ISL_413607 | hCoV-19/USA/CruiseA-2/2020 | *Homo sapiens* | North America / USA | 2020-02-18 | unknown | Pathogen Discovery, Respiratory Viruses Branch, Division of Viral Diseases, Centers for Dieases Control and Prevention | Anna Uehara, Ying Tao, Clinton R. Paden, Krista Queen, Jing Zhang, Yan Li, Mary S. Keckler, Alison S Laufer Halpin, Haibin Wang, Jasmine Padilla, Justin Lee, Christopher A. Elkins, Susan I. Gerber, Suxiang Tong |
| 137 | EPI_ISL_413609 | hCoV-19/USA/CruiseA-4/2020 | *Homo sapiens* | North America / USA | 2020-02-21 | unknown | Pathogen Discovery, Respiratory Viruses Branch, Division of Viral Diseases, Centers for Dieases Control and Prevention | Anna Uehara, Ying Tao, Clinton R. Paden, Krista Queen, Jing Zhang, Yan Li, Mary S. Keckler, Alison S Laufer Halpin, Haibin Wang, Jasmine Padilla, Justin Lee, Christopher A. Elkins, Susan I. Gerber, Suxiang Tong |
| 138 | EPI_ISL_413611 | hCoV-19/USA/CruiseA-6/2020 | *Homo sapiens* | North America / USA | 2020-02-21 | unknown | Pathogen Discovery, Respiratory Viruses Branch, Division of Viral Diseases, Centers for Dieases Control and Prevention | Anna Uehara, Ying Tao, Clinton R. Paden, Krista Queen, Jing Zhang, Yan Li, Mary S. Keckler, Alison S Laufer Halpin, Haibin Wang, Jasmine Padilla, Justin Lee, Christopher A. Elkins, Susan I. Gerber, Suxiang Tong |
| 139 | EPI_ISL_413612 | hCoV-19/USA/CruiseA-7/2020 | *Homo sapiens* | North America / USA | 2020-02-17 | unknown | Pathogen Discovery, Respiratory Viruses Branch, Division of Viral Diseases, Centers for Dieases Control and Prevention | Ying Tao, Clinton R. Paden, Krista Queen, Anna Uehara, Jing Zhang, Yan Li, Haibin Wang, Shifaq Kamili, Xiaoyan Lu, Brian Lynch, Senthil Kumar K. Sakthivel, Brett L. Whitaker, Lijuan Wang, Janna' R. Murray, Jasmine Padilla, Justin Lee, Susan I. Gerber, Stephen Lindstrom, Suxiang Tong |
| 140 | EPI_ISL_413613 | hCoV-19/USA/CruiseA-8/2020 | *Homo sapiens* | North America / USA | 2020-02-17 | unknown | Pathogen Discovery, Respiratory Viruses Branch, Division of Viral Diseases, Centers for Dieases Control and Prevention | Ying Tao, Clinton R. Paden, Krista Queen, Anna Uehara, Jing Zhang, Yan Li, Haibin Wang, Shifaq Kamili, Xiaoyan Lu, Brian Lynch, Senthil Kumar K. Sakthivel, Brett L. Whitaker, Lijuan Wang, Janna' R. Murray, Jasmine Padilla, Justin Lee, Susan I. Gerber, Stephen Lindstrom, Suxiang Tong |

| | | | | | | | | |
|---|---|---|---|---|---|---|---|---|
| 141 | EPI_ISL_413615 | hCoV-19/USA/CruiseA-10/2020 | *Homo sapiens* | North America / USA | 2020-02-17 | unknown | Pathogen Discovery, Respiratory Viruses Branch, Division of Viral Diseases, Centers for Dieases Control and Prevention | Ying Tao, Clinton R. Paden, Krista Queen, Anna Uehara, Jing Zhang, Yan Li, Haibin Wang, Shifaq Kamili, Xiaoyan Lu, Brian Lynch, Senthil Kumar K. Sakthivel, Brett L. Whitaker, Lijuan Wang, Janna' R. Murray, Jasmine Padilla, Justin Lee, Susan I. Gerber, Stephen Lindstrom, Suxiang Tong |
| 142 | EPI_ISL_413616 | hCoV-19/USA/CruiseA-11/2020 | *Homo sapiens* | North America / USA | 2020-02-17 | unknown | Pathogen Discovery, Respiratory Viruses Branch, Division of Viral Diseases, Centers for Dieases Control and Prevention | Ying Tao, Clinton R. Paden, Krista Queen, Anna Uehara, Jing Zhang, Yan Li, Haibin Wang, Shifaq Kamili, Xiaoyan Lu, Brian Lynch, Senthil Kumar K. Sakthivel, Brett L. Whitaker, Lijuan Wang, Janna' R. Murray, Jasmine Padilla, Justin Lee, Susan I. Gerber, Stephen Lindstrom, Suxiang Tong |
| 143 | EPI_ISL_413617 | hCoV-19/USA/CruiseA-12/2020 | *Homo sapiens* | North America / USA | 2020-02-20 | unknown | Pathogen Discovery, Respiratory Viruses Branch, Division of Viral Diseases, Centers for Dieases Control and Prevention | Ying Tao, Clinton R. Paden, Krista Queen, Anna Uehara, Jing Zhang, Yan Li, Haibin Wang, Shifaq Kamili, Xiaoyan Lu, Brian Lynch, Senthil Kumar K. Sakthivel, Brett L. Whitaker, Lijuan Wang, Janna' R. Murray, Jasmine Padilla, Justin Lee, Susan I. Gerber, Stephen Lindstrom, Suxiang Tong |
| 144 | EPI_ISL_413619 | hCoV-19/USA/CruiseA-14/2020 | *Homo sapiens* | North America / USA | 2020-02-25 | unknown | Pathogen Discovery, Respiratory Viruses Branch, Division of Viral Diseases, Centers for Dieases Control and Prevention | Clinton R. Paden, Ying Tao, Krista Queen, Anna Uehara, Jing Zhang, Yan Li, Haibin Wang, Shifaq Kamili, Xiaoyan Lu, Brian Lynch, Senthil Kumar K. Sakthivel, Brett L. Whitaker, Lijuan Wang, Janna' R. Murray, Jasmine Padilla, Justin Lee, Susan I. Gerber, Stephen Lindstrom, Suxiang Tong |
| 145 | EPI_ISL_413622 | hCoV-19/USA/CruiseA-17/2020 | *Homo sapiens* | North America / USA | 2020-02-24 | unknown | Pathogen Discovery, Respiratory Viruses Branch, Division of Viral Diseases, Centers for Dieases Control and Prevention | Clinton R. Paden, Ying Tao, Krista Queen, Anna Uehara, Jing Zhang, Yan Li, Haibin Wang, Shifaq Kamili, Xiaoyan Lu, Brian Lynch, Senthil Kumar K. Sakthivel, Brett L. Whitaker, Lijuan Wang, Janna' R. Murray, Jasmine Padilla, Justin Lee, Susan I. Gerber, Stephen Lindstrom, Suxiang Tong |
| 146 | EPI_ISL_413647 | hCoV-19/Portugal/CV62/2020 | *Homo sapiens* | Europe / Portugal | 2020-03-01 | Centro Hospital do Porto, E.P.E. - H. Geral de Santo Antonio | Instituto Nacional de Saude (INSA) | Raquel Guiomar, Inês Costa, Pedro Pechirra, Joana Mendonça, Luís Vieira, Helena Ramos, Joana Isidro, Vítor Borges, João Paulo Gomes |
| 147 | EPI_ISL_413648 | hCoV-19/Portugal/CV63/2020 | *Homo sapiens* | Europe / Portugal | 2020-03-01 | Centro Hospitalar e Universitário de Sao Joao, Porto | Instituto Nacional de Saude (INSA) | Raquel Guiomar, Inês Costa, Pedro Pechirra, Joana Mendonça, Luís Vieira, João Tiago Guimarães, Joana Isidro, Vítor Borges, João Paulo Gomes |
| 148 | EPI_ISL_413691 | hCoV-19/China/WF0001/2020 | *Homo sapiens* | Asia / China | 2020-01 | Weifang Center for Disease Control and Prevention | Weifang Center for Disease Control and Prevention & BGI-Shenzhen | Qing Nie, Xingguang Li, Erik M Volz, Han Fu, Haowei Wang, Xiaoyue Xi, Wei Chen, Dehui Liu, Yingying Chen, Mengmeng Tian, Wei Tan, Junjie Zai, Wanying Sun, Jiandong Li, Junhua Li |
| 148 | EPI_ISL_413746 | hCoV-19/China/WF0016/2020 | *Homo sapiens* | Asia / China | 2020-02 | Weifang Center for Disease Control and Prevention | Weifang Center for Disease Control and Prevention & BGI-Shenzhen | Qing Nie, Xingguang Li, Erik M Volz, Han Fu, Haowei Wang, Xiaoyue Xi, Wei Chen, Dehui Liu, Yingying Chen, Mengmeng Tian, Wei Tan, Junjie Zai, Wanying Sun, Jiandong Li, Junhua Li |
| 148 | EPI_ISL_413748 | hCoV-19/China/WF0018/2020 | *Homo sapiens* | Asia / China | 2020-02 | Weifang Center for Disease Control and Prevention | Weifang Center for Disease Control and Prevention & BGI-Shenzhen | Qing Nie, Xingguang Li, Erik M Volz, Han Fu, Haowei Wang, Xiaoyue Xi, Wei Chen, Dehui Liu, Yingying Chen, Mengmeng Tian, Wei Tan, Junjie Zai, Wanying Sun, Jiandong Li, Junhua Li |
| 149 | EPI_ISL_413692 | hCoV-19/China/WF0002/2020 | *Homo sapiens* | Asia / China | 2020-01 | Weifang Center for Disease Control and Prevention | Weifang Center for Disease Control and Prevention & BGI-Shenzhen | Qing Nie, Xingguang Li, Erik M Volz, Han Fu, Haowei Wang, Xiaoyue Xi, Wei Chen, Dehui Liu, Yingying Chen, Mengmeng Tian, Wei Tan, Junjie Zai, Wanying Sun, Jiandong Li, Junhua Li |
| 149 | EPI_ISL_413694 | hCoV-19/China/WF0004/2020 | *Homo sapiens* | Asia / China | 2020-01 | Weifang Center for Disease Control and Prevention | Weifang Center for Disease Control and Prevention & BGI-Shenzhen | Qing Nie, Xingguang Li, Erik M Volz, Han Fu, Haowei Wang, Xiaoyue Xi, Wei Chen, Dehui Liu, Yingying Chen, Mengmeng Tian, Wei Tan, Junjie Zai, Wanying Sun, Jiandong Li, Junhua Li |
| 150 | EPI_ISL_413693 | hCoV-19/China/WF0003/2020 | *Homo sapiens* | Asia / China | 2020-01 | Weifang Center for Disease Control and Prevention | Weifang Center for Disease Control and Prevention & BGI-Shenzhen | Qing Nie, Xingguang Li, Erik M Volz, Han Fu, Haowei Wang, Xiaoyue Xi, Wei Chen, Dehui Liu, Yingying Chen, Mengmeng Tian, Wei Tan, Junjie Zai, Wanying Sun, Jiandong Li, Junhua Li |
| 151 | EPI_ISL_413697 | hCoV-19/China/WF0012/2020 | *Homo sapiens* | Asia / China | 2020-02 | Weifang Center for Disease Control and Prevention | Weifang Center for Disease Control and Prevention & BGI-Shenzhen | Qing Nie, Xingguang Li, Erik M Volz, Han Fu, Haowei Wang, Xiaoyue Xi, Wei Chen, Dehui Liu, Yingying Chen, Mengmeng Tian, Wei Tan, Junjie Zai, Wanying Sun, Jiandong Li, Junhua Li |
| 151 | EPI_ISL_413761 | hCoV-19/China/WF0026/2020 | *Homo sapiens* | Asia / China | 2020-02 | Weifang Center for Disease Control and Prevention | Weifang Center for Disease Control and Prevention & BGI-Shenzhen | Qing Nie, Xingguang Li, Erik M Volz, Han Fu, Haowei Wang, Xiaoyue Xi, Wei Chen, Dehui Liu, Yingying Chen, Mengmeng Tian, Wei Tan, Junjie Zai, Wanying Sun, Jiandong Li, Junhua Li |
| 152 | EPI_ISL_413711 | hCoV-19/China/WF0014/2020 | *Homo sapiens* | Asia / China | 2020-02 | Weifang Center for Disease Control and Prevention | Weifang Center for Disease Control and Prevention & BGI-Shenzhen | Qing Nie, Xingguang Li, Erik M Volz, Han Fu, Haowei Wang, Xiaoyue Xi, Wei Chen, Dehui Liu, Yingying Chen, Mengmeng Tian, Wei Tan, Junjie Zai, Wanying Sun, Jiandong Li, Junhua Li |

| | | | | | | | | |
|---|---|---|---|---|---|---|---|---|
| 153 | EPI_ISL_413749 | hCoV-19/China/WF0019/2020 | *Homo sapiens* | Asia / China | 2020-02 | Weifang Center for Disease Control and Prevention | Weifang Center for Disease Control and Prevention & BGI-Shenzhen | Qing Nie, Xingguang Li, Erik M Volz, Han Fu, Haowei Wang, Xiaoyue Xi, Wei Chen, Dehui Liu, Yingying Chen, Mengmeng Tian, Wei Tan, Junjie Zai, Wanying Sun, Jiandong Li, Junhua Li |
| 154 | EPI_ISL_413791 | hCoV-19/China/WF0028/2020 | *Homo sapiens* | Asia / China | 2020-02 | Weifang Center for Disease Control and Prevention | Weifang Center for Disease Control and Prevention & BGI-Shenzhen | Qing Nie, Xingguang Li, Erik M Volz, Han Fu, Haowei Wang, Xiaoyue Xi, Wei Chen, Dehui Liu, Yingying Chen, Mengmeng Tian, Wei Tan, Junjie Zai, Wanying Sun, Jiandong Li, Junhua Li |
| 155 | EPI_ISL_413809 | hCoV-19/China/WF0029/2020 | *Homo sapiens* | Asia / China | 2020-02 | Weifang Center for Disease Control and Prevention | Weifang Center for Disease Control and Prevention & BGI-Shenzhen | Qing Nie, Xingguang Li, Erik M Volz, Han Fu, Haowei Wang, Xiaoyue Xi, Wei Chen, Dehui Liu, Yingying Chen, Mengmeng Tian, Wei Tan, Junjie Zai, Wanying Sun, Jiandong Li, Junhua Li |
| 156 | EPI_ISL_413854 | hCoV-19/Guangdong/2020XN4475-P0042/2020 | *Homo sapiens* | Asia / China / Guangdong | 2020-01-30 | Guangdong Provincial Institution of Public Health, Guangdong Provinical Center for Disease Control and Prevention | Guangdong Provincial Institution of Public Health | Jing Lu, Louis du Plessis, Liu Zhe, Jiufeng Sun, Sarah François, Huifang Lin, Moritz Kraemer, Jingju Peng, Qianlin Xiong, Runyu Yuan, Lilian Zeng, Pingping Zhou, Chuming Liang, Tao Liu, Wei Li, Juan Su, Huanying Zheng, Kang Min, Song Tie, Bo Peng, Shisong Fang, Wenzhe Su, Kuibiao Li, Ruilin Sun, Ru bai, Xi Tang, Minfeng Liang, Nuno Faria, Josh Quick, Andrew Rambaut, Verity Hill, Wenjun Ma, Nick Loman, Oliver Pybus, Changwen Ke |
| 157 | EPI_ISL_413857 | hCoV-19/Guangdong/2020XN4448-P0002/2020 | *Homo sapiens* | Asia / China / Guangdong | 2020-01-31 | Guangdong Provincial Institution of Public Health, Guangdong Provinical Center for Disease Control and Prevention | Guangdong Provincial Institution of Public Health | Jing Lu, Louis du Plessis, Liu Zhe, Jiufeng Sun, Sarah François, Huifang Lin, Moritz Kraemer, Jingju Peng, Qianlin Xiong, Runyu Yuan, Lilian Zeng, Pingping Zhou, Chuming Liang, Tao Liu, Wei Li, Juan Su, Huanying Zheng, Kang Min, Song Tie, Bo Peng, Shisong Fang, Wenzhe Su, Kuibiao Li, Ruilin Sun, Ru bai, Xi Tang, Minfeng Liang, Nuno Faria, Josh Quick, Andrew Rambaut, Verity Hill, Wenjun Ma, Nick Loman, Oliver Pybus, Changwen Ke |
| 158 | EPI_ISL_413858 | hCoV-19/Guangdong/2020XN4459-P0041/2020 | *Homo sapiens* | Asia / China / Guangdong | 2020-01-30 | Guangdong Provincial Institution of Public Health, Guangdong Provinical Center for Disease Control and Prevention | Guangdong Provincial Institution of Public Health | Jing Lu, Louis du Plessis, Liu Zhe, Jiufeng Sun, Sarah François, Huifang Lin, Moritz Kraemer, Jingju Peng, Qianlin Xiong, Runyu Yuan, Lilian Zeng, Pingping Zhou, Chuming Liang, Tao Liu, Wei Li, Juan Su, Huanying Zheng, Kang Min, Song Tie, Bo Peng, Shisong Fang, Wenzhe Su, Kuibiao Li, Ruilin Sun, Ru bai, Xi Tang, Minfeng Liang, Nuno Faria, Josh Quick, Andrew Rambaut, Verity Hill, Wenjun Ma, Nick Loman, Oliver Pybus, Changwen Ke |
| 159 | EPI_ISL_413861 | hCoV-19/Guangdong/GD2020080-P0010/2020 | *Homo sapiens* | Asia / China / Guangdong | 2020-02-01 | Guangdong Provincial Institution of Public Health, Guangdong Provinical Center for Disease Control and Prevention | Guangdong Provincial Institution of Public Health | Jing Lu, Louis du Plessis, Liu Zhe, Jiufeng Sun, Sarah François, Huifang Lin, Moritz Kraemer, Jingju Peng, Qianlin Xiong, Runyu Yuan, Lilian Zeng, Pingping Zhou, Chuming Liang, Tao Liu, Wei Li, Juan Su, Huanying Zheng, Kang Min, Song Tie, Bo Peng, Shisong Fang, Wenzhe Su, Kuibiao Li, Ruilin Sun, Ru bai, Xi Tang, Minfeng Liang, Nuno Faria, Josh Quick, Andrew Rambaut, Verity Hill, Wenjun Ma, Nick Loman, Oliver Pybus, Changwen Ke |
| 160 | EPI_ISL_413863 | hCoV-19/Guangdong/GD2020087-P0008/2020 | *Homo sapiens* | Asia / China / Guangdong | 2020-02-01 | Guangdong Provincial Institution of Public Health, Guangdong Provinical Center for Disease Control and Prevention | Guangdong Provincial Institution of Public Health | Jing Lu, Louis du Plessis, Liu Zhe, Jiufeng Sun, Sarah François, Huifang Lin, Moritz Kraemer, Jingju Peng, Qianlin Xiong, Runyu Yuan, Lilian Zeng, Pingping Zhou, Chuming Liang, Tao Liu, Wei Li, Juan Su, Huanying Zheng, Kang Min, Song Tie, Bo Peng, Shisong Fang, Wenzhe Su, Kuibiao Li, Ruilin Sun, Ru bai, Xi Tang, Minfeng Liang, Nuno Faria, Josh Quick, Andrew Rambaut, Verity Hill, Wenjun Ma, Nick Loman, Oliver Pybus, Changwen Ke |
| 161 | EPI_ISL_413928 | hCoV-19/USA/CA-CDPH-UC9/2020 | *Homo sapiens* | North America / USA / California / San Francisco | 2020-03-05 | California Department of Public Health | Chiu Laboratory UCSF-Abbott Viral Diagnostics and Discovery Center | Xianding Deng, Scot Federman, Guixia Yu, Chao-Yang Pan, Hugo Guevara, Alicia Sotomayor-Gonzalez, Allan Gopez, Wei Gu, Steve Miller, Debra A. Wadford, and Charles Y. Chiu |
| 162 | EPI_ISL_413931 | hCoV-19/USA/UC-CDPH-UC11/2020 | *Homo sapiens* | North America / USA / California / San Francisco | 2020-03-05 | California Department of Public Health | Chiu Laboratory UCSF-Abbott Viral Diagnostics and Discovery Center University of California, San Francisco | Xianding Deng, Scot Federman, Guixia Yu, Chao-Yang Pan, Hugo Guevara, Alicia Sotomayor-Gonzalez, Allan Gopez, Wei Gu, Steve Miller, Debra A. Wadford, and Charles Y. Chiu |
| 163 | EPI_ISL_413996 | hCoV-19/Switzerland/TI9486/2020 | *Homo sapiens* | Europe / Switzerland / Tessin | 2020-02-24 | Laboratoire de Virologie, HUG | Swiss National Reference Centre for Influenza | LAUBSCHER Florian et al. |
| 164 | EPI_ISL_413997 | hCoV-19/Switzerland/GE3895/2020 | *Homo sapiens* | Europe / Switzerland / Geneva | 2020-02-26 | Laboratoire de Virologie, HUG | Swiss National Reference Centre for Influenza | LAUBSCHER Florian et al. |

| | | | | | | | | |
|---|---|---|---|---|---|---|---|---|
| 165 | EPI_ISL_413999 | hCoV-19/Switzerland/AG0361/2020 | *Homo sapiens* | Europe / Switzerland / Argovie | 2020-02-27 | Laboratoire de Virologie, HUG | Swiss National Reference Centre for Influenza | LAUBSCHER Florian et al. |
| 166 | EPI_ISL_414012 | hCoV-19/England/200990723/2020 | *Homo sapiens* | Europe / United Kingdom / England | 2020-02-27 | Respiratory Virus Unit, Microbiology Services Colindale, Public Health England | Respiratory Virus Unit, Microbiology Services Colindale, Public Health England | Monica Galiano, Shahjahan Miah, Angie Lackenby, Omolola Akinbami, Tiina Talts, Leena Bhaw, Richard Myers, Steven Platt, Kirstin Edwards, Jonathan Hubb, Joanna Ellis, Maria Zambon |
| 167 | EPI_ISL_414014 | hCoV-19/Brazil/SPBR-03/2020 | *Homo sapiens* | South America / Brazil / Sao Paulo | 2020-03-02 | Hospital Israelita Albert Einstein | Instituto Adolfo Lutz, Interdiciplinary Procedures Center, Strategic Laboratory | Claudio Tavares Sacchi, Claudia Regina Gonçalves, Katia Correia dos Santos, Carlos Henrique Camargo, Maria do Carmo Sampaito Tavares Timenetsky, Terezinha Maria de Paiva, Ester Cerdeira Sabino |
| 167 | EPI_ISL_414017 | hCoV-19/Brazil/SPBR-04/2020 | *Homo sapiens* | South America / Brazil / Sao Paulo | 2020-03-04 | Hospital São Joaquim Beneficencia Portuguesa | Instituto Adolfo Lutz, Interdiciplinary Procedures Center, Strategic Laboratory | Claudio Tavares Sacchi, Claudia Regina Gonçalves, Fabiana Cristina Pereira dos Santos, Carlos Henrique Camargo, Maria do Carmo Sampaito Tavares Timenetsky, Daniela Bernardes Borges da Silva, Terezinha Maria de Paiva, Ester Cerdeira Sabino |
| 167 | EPI_ISL_416028 | hCoV-19/Brazil/SPBR-07/2020 | *Homo sapiens* | South America / Brazil / Sao Paulo / Sao Paulo | 2020-03-03 | National Influenza Center - Instituto Adolfo Lutz | Instituto Adolfo Lutz, Interdiciplinary Procedures Center, Strategic Laboratory | Claudio Tavares Sacchi, Claudia Regina Gonçalves, Carlos Henrique Camargo, Fabiana Cristina Pereira dos Santos, Daniela Bernardes Borges da Silva, Simone Guadagnucci Morillo, Adriano Abbud, Adriana Bugno, Maria do Carmo Sampaio Tavares Timenetsky, Terezinha Maria de Paiva |
| 168 | EPI_ISL_414015 | hCoV-19/Brazil/SPBR-06/2020 | *Homo sapiens* | South America / Brazil / Sao Paulo / Sao Paulo | 2020-02-29 | Hospital São Joaquim Beneficencia Portuguesa | Instituto Adolfo Lutz, Interdiciplinary Procedures Center, Strategic Laboratory | Claudio Tavares Sacchi, Claudia Regina Gonçalves, SimoneGuadagnucci Morillo, Carlos Henrique Camargo, Maria do Carmo Sampaito Tavares Timenetsky, Fabiana Cristina Pereira dos Santos Terezinha Maria de Paiva, Ester Cerdeira Sabino |
| 168 | EPI_ISL_416658 | hCoV-19/USA/WA-UW120/2020 | *Homo sapiens* | North America / USA / Washington | 2020-03-11 | UW Virology Lab | UW Virology Lab | Pavitra Roychoudhury, Hong Xie, Keith Jerome, Alexander Greninger |
| 168 | EPI_ISL_416742 | hCoV-19/Czech Republic/ChVir1630/2020 | *Homo sapiens* | Europe / Czech Republic / Prague | 2020-02 | NRL for Influenza, Centrum Epidemiology and Microbiology of National Institute of Public Health, Czech Republic | Charite Universitaetsmedizin Berlin, Institute of Virology | Victor M Corman, Julia Schneider, Jörn Beheim-Schwarzbach, Talitha Veith, Barbara Muehlemann, Terry Jones, Akexander Nagy, Jaromira Vecerova, Dusan Trnka, Ludmila Novakova, Helena Jirincova, Christian Drosten |
| 168 | EPI_ISL_416832 | hCoV-19/USA/NY-NYUMC4/2020 | *Homo sapiens* | North America / USA / New York City | 2020-03-16 | NYU Langone Health | Department of Pathology and Medicine, New York University School of Medicine | John Chen, Dacia Dimartino, Xiaojun Feng, Adriana Heguy, Megan Hogan, Emily Huang, George Jour, Christian Marier, Matt Maurano, Mark Mulligan, Peter Meyn, Marie Samanovic-Golden, Amy Rapkiewicz, Guomiao Shen, Matija Snuderl, Gael Westby, Paul Zappile |
| 168 | EPI_ISL_417032 | hCoV-19/Australia/QLDID920/2020 | *Homo sapiens* | Oceania / Australia / Queensland / Rockhampton | 2020-03-11 | Rockhampton Base Hospital | Public Health Virology Laboratory | Bixing Huang, Alyssa Pyke, Amanda De Jong, Andrew Van Den Hurk, Carmel Taylor, David Warrilow, Doris Genge, Elisabeth Gamez, Glen Hewitson, Ian Maxwell Mackay, Inga Sultana, Jamie McMahon, Jean Barcelon, Judy Northill, Mitchell Finger, Natalie Simpson, Neelima Nair, Peter Burtonclay, Peter Moore, Sarah Wheatley, Sean Moody, Sonja Hall-Mendelin, Timothy Gardam, and Frederick Moore |
| 169 | EPI_ISL_414016 | hCoV-19/Brazil/SPBR-05/2020 | *Homo sapiens* | South America / Brazil / Sao Paulo / Sao Paulo | 2020-02-29 | Hospital São Joaquim Beneficencia Portuguesa | Instituto Adolfo Lutz, Interdiciplinary Procedures Center, Strategic Laboratory | Claudio Tavares Sacchi, Claudia Regina Gonçalves, Audrey Cilli, Carlos Henrique Camargo, Maria do Carmo Sampaito Tavares Timenetsky, Daniela Bernardes Borges da Silva, Terezinha Maria de Paiva, Ester Cerdeira Sabino |
| 169 | EPI_ISL_416031 | hCoV-19/Brazil/SPBR-09/2020 | *Homo sapiens* | South America / Brazil / Sao Paulo / Sao Paulo | 2020-03-04 | National Influenza Center - Instituto Adolfo Lutz | Instituto Adolfo Lutz, Interdiciplinary Procedures Center, Strategic Laboratory | Claudio Tavares Sacchi, Claudia Regina Gonçalves, Carlos Henrique Camargo, Fabiana Cristina Pereira dos Santos, Daniela Bernardes Borges da Silva, Simone Guadagnucci Morillo, Adriano Abbud, Adriana Bugno, Maria do Carmo Sampaio Tavares Timenetsky, Terezinha Maria de Paiva |
| 170 | EPI_ISL_414019 | hCoV-19/Switzerland/GE3121/2020 | *Homo sapiens* | Europe / Switzerland / Geneva | 2020-02-27 | Laboratoire de Virologie, HUG | Swiss National Reference Centre for Influenza | LAUBSCHER Florian et al. |

| | | | | | | | | |
|---|---|---|---|---|---|---|---|---|
| 170 | EPI_ISL_414020 | hCoV-19/Switzerland/GE5373/2020 | *Homo sapiens* | Europe / Switzerland / Geneva | 2020-02-27 | Laboratoire de Virologie, HUG | Swiss National Reference Centre for Influenza | LAUBSCHER Florian et al. |
| 171 | EPI_ISL_414023 | hCoV-19/Switzerland/VD5615/2020 | *Homo sapiens* | Europe / Switzerland / Vaud | 2020-03-01 | Laboratoire de Virologie, HUG | Swiss National Reference Centre for Influenza | LAUBSCHER Florian et al. |
| 171 | EPI_ISL_415459 | hCoV-19/Switzerland/VD0503/2020 | *Homo sapiens* | Europe / Switzerland / Genève | 2020-02-29 | Hôpitaux universitaires de Genève Laboratoire de Virologie | Hôpitaux universitaires de Genève Laboratoire de Virologie | Laubscher F. |
| 172 | EPI_ISL_414027 | hCoV-19/Scotland/CVR05/2020 | *Homo sapiens* | Europe / United Kingdom / Scotland | 2020-03-04 | West of Scotland Specialist Virology Centre, NHSGGC | MRC-University of Glasgow Centre for Virus Research | Emma Thomson, Antonia Ho; Kathy Smollett, Daniel Mair, Stephen Carmichael, Ana da Silva Filipe; Richard Orton, David L Robertson; Alasdair MacLean, Rory Gunson. |
| 173 | EPI_ISL_414363 | hCoV-19/USA/WA-UW15/2020 | *Homo sapiens* | North America / USA / Washington | 2020-03-04 | UW Virology Lab | UW Virology Lab | Pavitra Roychoudhury, Hong Xie, Keith Jerome, Alexander Greninger |
| 174 | EPI_ISL_414368 | hCoV-19/USA/WA-UW20/2020 | *Homo sapiens* | North America / USA / Washington | 2020-03-05 | UW Virology Lab | UW Virology Lab | Pavitra Roychoudhury, Hong Xie, Keith Jerome, Alexander Greninger |
| 175 | EPI_ISL_414414 | hCoV-19/Australia/QLD09/2020 | *Homo sapiens* | Oceania / Australia / Queensland / Gold Coast | 2020-02-29 | Pathology Queensland | Public Health Virology Laboratory | Bixing Huang, Alyssa Pyke, Amanda De Jong, Andrew Van Den Hurk, Carmel Taylor, David Warrilow, Doris Genge, Elisabeth Gamez, Glen Hewitson, Ian Maxwell Mackay, Inga Sultana, Jamie McMahon, Jean Barcelon, Judy Northill, Mitchell Finger, Natalie Simpson, Neelima Nair, Peter Burtonclay, Peter Moore, Sarah Wheatley, Sean Moody, Sonja Hall-Mendelin, Timothy Gardam, and Frederick Moore |
| 176 | EPI_ISL_414428 | hCoV-19/Netherlands/NoordBrabant_1/2020 | *Homo sapiens* | Europe / Netherlands / Noord Brabant | 2020-03-02 | Dutch COVID-19 response team | Erasmus Medical Center | David Nieuwenhuijse, Bas Oude Munnink, Reina Sikkema, Claudia Schapendonk, Irina Chestakova, Anne van der Linden, Mark Pronk, Pascal Lexmond, Corien Swaan, Manon Haverkate, Madelief Mollers, Mart Stein, Sandra Kengne Kamga Mobou, Jeroen van Kampen, Jolanda Voermans, Aura Timen, Corine GeurtsvanKessel, Annemiek van der Eijk, Richard Molenkamp, Marion Koopmans, on behalf of the Dutch national COVID-19 response team. |
| 177 | EPI_ISL_414429 | hCoV-19/Netherlands/NoordBrabant_3/2020 | *Homo sapiens* | Europe / Netherlands / Noord Brabant | 2020-03-02 | Dutch COVID-19 response team | Erasmus Medical Center | David Nieuwenhuijse, Bas Oude Munnink, Reina Sikkema, Claudia Schapendonk, Irina Chestakova, Anne van der Linden, Mark Pronk, Pascal Lexmond, Corien Swaan, Manon Haverkate, Madelief Mollers, Mart Stein, Sandra Kengne Kamga Mobou, Jeroen van Kampen, Jolanda Voermans, Aura Timen, Corine GeurtsvanKessel, Annemiek van der Eijk, Richard Molenkamp, Marion Koopmans, on behalf of the Dutch national COVID-19 response team. |
| 177 | EPI_ISL_415498 | hCoV-19/Netherlands/NA_9/2020 | *Homo sapiens* | Europe / Netherlands | 2020-03-09 | Dutch COVID-19 response team | Erasmus Medical Center | David Nieuwenhuijse, Bas Oude Munnink, Reina Sikkema, Claudia Schapendonk, Irina Chestakova, Anne van der Linden, Mark Pronk, Pascal Lexmond, Corien Swaan, Manon Haverkate, Madelief Mollers, Mart Stein, Sandra Kengne Kamga Mobou, Jeroen van Kampen, Jolanda Voermans, Aura Timen, Corine GeurtsvanKessel, Annemiek van der Eijk, Richard Molenkamp, Marion Koopmans, on behalf of the Dutch national COVID-19 response team. |
| 178 | EPI_ISL_414435 | hCoV-19/Netherlands/Utrecht_1/2020 | *Homo sapiens* | Europe / Netherlands / Utrecht | 2020-03-03 | Dutch COVID-19 response team | Erasmus Medical Center | David Nieuwenhuijse, Bas Oude Munnink, Reina Sikkema, Claudia Schapendonk, Irina Chestakova, Anne van der Linden, Mark Pronk, Pascal Lexmond, Corien Swaan, Manon Haverkate, Madelief Mollers, Mart Stein, Sandra Kengne Kamga Mobou, Jeroen van Kampen, Jolanda Voermans, Aura Timen, Corine GeurtsvanKessel, Annemiek van der Eijk, Richard Molenkamp, Marion Koopmans, on behalf of the Dutch national COVID-19 response team. |

| | | | | | | | | |
|---|---|---|---|---|---|---|---|---|
| 179 | EPI_ISL_414445 | hCoV-19/Netherlands/ZuidHolland_9/2020 | *Homo sapiens* | Europe / Netherlands / Zuid Holland | 2020-03-03 | Dutch COVID-19 response team | Erasmus Medical Center | David Nieuwenhuijse, Bas Oude Munnink, Reina Sikkema, Claudia Schapendonk, Irina Chestakova, Anne van der Linden, Mark Pronk, Pascal Lexmond, Corien Swaan, Manon Haverkate, Madelief Mollers, Mart Stein, Sandra Kengne Kamga Mobou, Jeroen van Kampen, Jolanda Voermans, Aura Timen, Corine GeurtsvanKessel, Annemiek van der Eijk, Richard Molenkamp, Marion Koopmans, on behalf of the Dutch national COVID-19 response team. |
| 179 | EPI_ISL_415480 | hCoV-19/Netherlands/NA_23/2020 | *Homo sapiens* | Europe / Netherlands | 2020-03-09 | Dutch COVID-19 response team | Erasmus Medical Center | David Nieuwenhuijse, Bas Oude Munnink, Reina Sikkema, Claudia Schapendonk, Irina Chestakova, Anne van der Linden, Mark Pronk, Pascal Lexmond, Corien Swaan, Manon Haverkate, Madelief Mollers, Mart Stein, Sandra Kengne Kamga Mobou, Jeroen van Kampen, Jolanda Voermans, Aura Timen, Corine GeurtsvanKessel, Annemiek van der Eijk, Richard Molenkamp, Marion Koopmans, on behalf of the Dutch national COVID-19 response team. |
| 180 | EPI_ISL_414451 | hCoV-19/Netherlands/NoordBrabant_6/2020 | *Homo sapiens* | Europe / Netherlands / Noord Brabant | 2020-03-06 | Dutch COVID-19 response team | Erasmus Medical Center | David Nieuwenhuijse, Bas Oude Munnink, Reina Sikkema, Claudia Schapendonk, Irina Chestakova, Anne van der Linden, Mark Pronk, Pascal Lexmond, Corien Swaan, Manon Haverkate, Madelief Mollers, Mart Stein, Sandra Kengne Kamga Mobou, Jeroen van Kampen, Jolanda Voermans, Aura Timen, Corine GeurtsvanKessel, Annemiek van der Eijk, Richard Molenkamp, Marion Koopmans, on behalf of the Dutch national COVID-19 response team. |
| 181 | EPI_ISL_414457 | hCoV-19/Netherlands/NoordBrabant_17/2020 | *Homo sapiens* | Europe / Netherlands / Noord Brabant | 2020-03-06 | Dutch COVID-19 response team | Erasmus Medical Center | David Nieuwenhuijse, Bas Oude Munnink, Reina Sikkema, Claudia Schapendonk, Irina Chestakova, Anne van der Linden, Mark Pronk, Pascal Lexmond, Corien Swaan, Manon Haverkate, Madelief Mollers, Mart Stein, Sandra Kengne Kamga Mobou, Jeroen van Kampen, Jolanda Voermans, Aura Timen, Corine GeurtsvanKessel, Annemiek van der Eijk, Richard Molenkamp, Marion Koopmans, on behalf of the Dutch national COVID-19 response team. |
| 182 | EPI_ISL_414468 | hCoV-19/Netherlands/ZuidHolland_8/2020 | *Homo sapiens* | Europe / Netherlands / Zuid Holland | 2020-03-06 | Dutch COVID-19 response team | Erasmus Medical Center | David Nieuwenhuijse, Bas Oude Munnink, Reina Sikkema, Claudia Schapendonk, Irina Chestakova, Anne van der Linden, Mark Pronk, Pascal Lexmond, Corien Swaan, Manon Haverkate, Madelief Mollers, Mart Stein, Sandra Kengne Kamga Mobou, Jeroen van Kampen, Jolanda Voermans, Aura Timen, Corine GeurtsvanKessel, Annemiek van der Eijk, Richard Molenkamp, Marion Koopmans, on behalf of the Dutch national COVID-19 response team. |
| 183 | EPI_ISL_414470 | hCoV-19/Netherlands/ZuidHolland_13/2020 | *Homo sapiens* | Europe / Netherlands / Zuid Holland | 2020-03-06 | Dutch COVID-19 response team | Erasmus Medical Center | David Nieuwenhuijse, Bas Oude Munnink, Reina Sikkema, Claudia Schapendonk, Irina Chestakova, Anne van der Linden, Mark Pronk, Pascal Lexmond, Corien Swaan, Manon Haverkate, Madelief Mollers, Mart Stein, Sandra Kengne Kamga Mobou, Jeroen van Kampen, Jolanda Voermans, Aura Timen, Corine GeurtsvanKessel, Annemiek van der Eijk, Richard Molenkamp, Marion Koopmans, on behalf of the Dutch national COVID-19 response team. |
| 184 | EPI_ISL_414482 | hCoV-19/USA/CruiseA-23/2020 | *Homo sapiens* | North America / USA | 2020-02-18 | unknown | Pathogen Discovery, Respiratory Viruses Branch, Division of Viral Diseases, Centers for Disease Control and Prevention | Krista Queen, Anna Uehara, Ying Tao, Clinton R. Paden, Jing Zhang, Yan Li, Haibin Wang, Shifaq Kamili, Xiaoyan Lu, Brian Lynch, Senthil Kumar K. Sakthivel, Brett L. Whitaker, Lijuan Wang, Janna' R. Murray, Jasmine Padilla, Justin Lee, Susan I. Gerber, Stephen Lindstrom, Suxiang Tong |
| 185 | EPI_ISL_414487 | hCoV-19/Ireland/COR-20134/2020 | *Homo sapiens* | Europe / Ireland / Cork | 2020-03-04 | UCD National Virus Reference Laboratory | UCD National Virus Reference Laboratory | Michael Carr, Gabriel Gonzalez, Jonathan Dean, Suzie Coughlan, Alison Murphy, Kevin Byrne, Ken Wolfe, Jeff Connell, Brendan Loftus, Cillian F De Gascun |

| | | | | | | | | |
|---|---|---|---|---|---|---|---|---|
| 186 | EPI_ISL_414509 | hCoV-19/Germany/NRW-09/2020 | *Homo sapiens* | Europe / Germany / North Rhine Westphalia / Heinsberg District | 2020-02-28 | Center of Medical Microbiology, Virology, and Hospital Hygiene, University of Duesseldorf | Center of Medical Microbiology, Virology, and Hospital Hygiene, University of Duesseldorf | Ortwin Adams, Marcel Andree, Alexander Dilthey, Torsten Feldt, Sandra Hauka, Torsten Houwaart, Björn-Erik Jensen, Detlef Kindgen-Milles, Malte Kohns Vasconcelos, Klaus Pfeffer, Tina Senff, Daniel Strelow, Jörg Timm, Andreas Walker, Tobias Wienemann |
| 187 | EPI_ISL_414510 | hCoV-19/Shanghai/SH01/2020 | *Homo sapiens* | Asia / China / Shanghai | 2020-02-02 | unknown | Key Laboratory of Medical Molecular Virology (MOE/NHC/CAMS) | Zhang,R., Yi,Z., Wang,Y., Teng,Z., Xu,W., Song,W., Cai,X., Sun,Z., Gu,C., Zhou,Y., Chen,H., Ye,R., Han,W., Zhu,Y., Feng,F., Fang,F., Li,C., Zhang,X., Qu,D., Fu,C., Xie,Y. and Yuan,Z. |
| 188 | EPI_ISL_414511 | hCoV-19/Japan/TKYE6182/2020 | *Homo sapiens* | Asia / Japan | 2020-01 | unknown | Ryota Kumagai Tokyo Metropolitan Institute of Public Health | Kumagai,R., Yoshida,I., Nagashima,M., Chiba,T. and Sadamasu,K. |
| 189 | EPI_ISL_414520 | hCoV-19/Germany/BavPat2/2020 | *Homo sapiens* | Europe / Germany / Munich | 2020-03-02 | Bundeswehr Institute of Microbiology | Bundeswehr Institute of Microbiology | Mathias C Walter, Markus H Antwerpen and Roman Wölfel |
| 190 | EPI_ISL_414521 | hCoV-19/Germany/BavPat3/2020 | *Homo sapiens* | Europe / Germany / Munich | 2020-03-02 | Bundeswehr Institute of Microbiology | Bundeswehr Institute of Microbiology | Mathias C Walter, Markus H Antwerpen and Roman Wölfel |
| 191 | EPI_ISL_414523 | hCoV-19/England/200990660/2020 | *Homo sapiens* | Europe / United Kingdom / England | 2020-02-27 | Respiratory Virus Unit, Microbiology Services Colindale, Public Health England | Respiratory Virus Unit, Microbiology Services Colindale, Public Health England | Monica Galiano, Shahjahan Miah, Angie Lackenby, Omolola Akinbami, Tiina Talts, Leena Bhaw, Richard Myers, Steven Platt, Kirstin Edwards, Jonathan Hubb, Joanna Ellis, Maria Zambon |
| 191 | EPI_ISL_414526 | hCoV-19/England/201040141/2020 | *Homo sapiens* | Europe / United Kingdom / England | 2020-03-03 | Respiratory Virus Unit, Microbiology Services Colindale, Public Health England | Respiratory Virus Unit, Microbiology Services Colindale, Public Health England | Monica Galiano, Shahjahan Miah, Angie Lackenby, Omolola Akinbami, Tiina Talts, Leena Bhaw, Richard Myers, Steven Platt, Kirstin Edwards, Jonathan Hubb, Joanna Ellis, Maria Zambon |
| 192 | EPI_ISL_414525 | hCoV-19/England/201040081/2020 | *Homo sapiens* | Europe / United Kingdom / England | 2020-03-02 | Respiratory Virus Unit, Microbiology Services Colindale, Public Health England | Respiratory Virus Unit, Microbiology Services Colindale, Public Health England | Monica Galiano, Shahjahan Miah, Angie Lackenby, Omolola Akinbami, Tiina Talts, Leena Bhaw, Richard Myers, Steven Platt, Kirstin Edwards, Jonathan Hubb, Joanna Ellis, Maria Zambon |
| 193 | EPI_ISL_414545 | hCoV-19/Netherlands/NoordBrabant_36/2020 | *Homo sapiens* | Europe / Netherlands / Noord Brabant | 2020-03-03 | Dutch COVID-19 response team | Erasmus Medical Center | David Nieuwenhuijse, Bas Oude Munnink, Reina Sikkema, Claudia Schapendonk, Irina Chestakova, Anne van der Linden, Mark Pronk, Pascal Lexmond, Corien Swaan, Manon Haverkate, Madelief Mollers, Mart Stein, Sandra Kengne Kamga Mobou, Jeroen van Kampen, Jolanda Voermans, Aura Timen, Corine GeurtsvanKessel, Annemiek van der Eijk, Richard Molenkamp, Marion Koopmans, on behalf of the Dutch national COVID-19 response team. |
| 193 | EPI_ISL_414557 | hCoV-19/Netherlands/ZuidHolland_15/2020 | *Homo sapiens* | Europe / Netherlands / Zuid Holland | 2020-03-08 | Dutch COVID-19 response team | Erasmus Medical Center | David Nieuwenhuijse, Bas Oude Munnink, Reina Sikkema, Claudia Schapendonk, Irina Chestakova, Anne van der Linden, Mark Pronk, Pascal Lexmond, Corien Swaan, Manon Haverkate, Madelief Mollers, Mart Stein, Sandra Kengne Kamga Mobou, Jeroen van Kampen, Jolanda Voermans, Aura Timen, Corine GeurtsvanKessel, Annemiek van der Eijk, Richard Molenkamp, Marion Koopmans, on behalf of the Dutch national COVID-19 response team. |
| 193 | EPI_ISL_414561 | hCoV-19/Netherlands/ZuidHolland_19/2020 | *Homo sapiens* | Europe / Netherlands / Zuid Holland | 2020-03-05 | Dutch COVID-19 response team | Erasmus Medical Center | David Nieuwenhuijse, Bas Oude Munnink, Reina Sikkema, Claudia Schapendonk, Irina Chestakova, Anne van der Linden, Mark Pronk, Pascal Lexmond, Corien Swaan, Manon Haverkate, Madelief Mollers, Mart Stein, Sandra Kengne Kamga Mobou, Jeroen van Kampen, Jolanda Voermans, Aura Timen, Corine GeurtsvanKessel, Annemiek van der Eijk, Richard Molenkamp, Marion Koopmans, on behalf of the Dutch national COVID-19 response team. |

| | | | | | | | | |
|---|---|---|---|---|---|---|---|---|
| 194 | EPI_ISL_414549 | hCoV-19/Netherlands/NoordHolland_2/2020 | *Homo sapiens* | Europe / Netherlands / Noord Holland | 2020-03-03 | Dutch COVID-19 response team | Erasmus Medical Center | David Nieuwenhuijse, Bas Oude Munnink, Reina Sikkema, Claudia Schapendonk, Irina Chestakova, Anne van der Linden, Mark Pronk, Pascal Lexmond, Corien Swaan, Manon Haverkate, Madelief Mollers, Mart Stein, Sandra Kengne Kamga Mobou, Jeroen van Kampen, Jolanda Voermans, Aura Timen, Corine GeurtsvanKessel, Annemiek van der Eijk, Richard Molenkamp, Marion Koopmans, on behalf of the Dutch national COVID-19 response team. |
| 195 | EPI_ISL_414552 | hCoV-19/Netherlands/Utrecht_13/2020 | *Homo sapiens* | Europe / Netherlands / Utrecht | 2020-03-07 | Dutch COVID-19 response team | Erasmus Medical Center | David Nieuwenhuijse, Bas Oude Munnink, Reina Sikkema, Claudia Schapendonk, Irina Chestakova, Anne van der Linden, Mark Pronk, Pascal Lexmond, Corien Swaan, Manon Haverkate, Madelief Mollers, Mart Stein, Sandra Kengne Kamga Mobou, Jeroen van Kampen, Jolanda Voermans, Aura Timen, Corine GeurtsvanKessel, Annemiek van der Eijk, Richard Molenkamp, Marion Koopmans, on behalf of the Dutch national COVID-19 response team. |
| 195 | EPI_ISL_414564 | hCoV-19/Netherlands/ZuidHolland_22/2020 | *Homo sapiens* | Europe / Netherlands / Zuid Holland | 2020-03-08 | Dutch COVID-19 response team | Erasmus Medical Center | David Nieuwenhuijse, Bas Oude Munnink, Reina Sikkema, Claudia Schapendonk, Irina Chestakova, Anne van der Linden, Mark Pronk, Pascal Lexmond, Corien Swaan, Manon Haverkate, Madelief Mollers, Mart Stein, Sandra Kengne Kamga Mobou, Jeroen van Kampen, Jolanda Voermans, Aura Timen, Corine GeurtsvanKessel, Annemiek van der Eijk, Richard Molenkamp, Marion Koopmans, on behalf of the Dutch national COVID-19 response team. |
| 196 | EPI_ISL_414554 | hCoV-19/Netherlands/Utrecht_15/2020 | *Homo sapiens* | Europe / Netherlands / Utrecht | 2020-03-08 | Dutch COVID-19 response team | Erasmus Medical Center | David Nieuwenhuijse, Bas Oude Munnink, Reina Sikkema, Claudia Schapendonk, Irina Chestakova, Anne van der Linden, Mark Pronk, Pascal Lexmond, Corien Swaan, Manon Haverkate, Madelief Mollers, Mart Stein, Sandra Kengne Kamga Mobou, Jeroen van Kampen, Jolanda Voermans, Aura Timen, Corine GeurtsvanKessel, Annemiek van der Eijk, Richard Molenkamp, Marion Koopmans, on behalf of the Dutch national COVID-19 response team. |
| 197 | EPI_ISL_414555 | hCoV-19/Netherlands/Utrecht_16/2020 | *Homo sapiens* | Europe / Netherlands / Utrecht | 2020-03-08 | Dutch COVID-19 response team | Erasmus Medical Center | David Nieuwenhuijse, Bas Oude Munnink, Reina Sikkema, Claudia Schapendonk, Irina Chestakova, Anne van der Linden, Mark Pronk, Pascal Lexmond, Corien Swaan, Manon Haverkate, Madelief Mollers, Mart Stein, Sandra Kengne Kamga Mobou, Jeroen van Kampen, Jolanda Voermans, Aura Timen, Corine GeurtsvanKessel, Annemiek van der Eijk, Richard Molenkamp, Marion Koopmans, on behalf of the Dutch national COVID-19 response team. |
| 198 | EPI_ISL_414558 | hCoV-19/Netherlands/ZuidHolland_16/2020 | *Homo sapiens* | Europe / Netherlands / Zuid Holland | 2020-03-06 | Dutch COVID-19 response team | Erasmus Medical Center | David Nieuwenhuijse, Bas Oude Munnink, Reina Sikkema, Claudia Schapendonk, Irina Chestakova, Anne van der Linden, Mark Pronk, Pascal Lexmond, Corien Swaan, Manon Haverkate, Madelief Mollers, Mart Stein, Sandra Kengne Kamga Mobou, Jeroen van Kampen, Jolanda Voermans, Aura Timen, Corine GeurtsvanKessel, Annemiek van der Eijk, Richard Molenkamp, Marion Koopmans, on behalf of the Dutch national COVID-19 response team. |
| 199 | EPI_ISL_414559 | hCoV-19/Netherlands/ZuidHolland_17/2020 | *Homo sapiens* | Europe / Netherlands / Zuid Holland | 2020-03-07 | Dutch COVID-19 response team | Erasmus Medical Center | David Nieuwenhuijse, Bas Oude Munnink, Reina Sikkema, Claudia Schapendonk, Irina Chestakova, Anne van der Linden, Mark Pronk, Pascal Lexmond, Corien Swaan, Manon Haverkate, Madelief Mollers, Mart Stein, Sandra Kengne Kamga Mobou, Jeroen van Kampen, Jolanda Voermans, Aura Timen, Corine GeurtsvanKessel, Annemiek van der Eijk, Richard Molenkamp, Marion Koopmans, on behalf of the Dutch national COVID-19 response team. |

| | | | | | | | | |
|---|---|---|---|---|---|---|---|---|
| 200 | EPI_ISL_414562 | hCoV-19/Netherlands/ZuidHolland_20/2020 | *Homo sapiens* | Europe / Netherlands / Zuid Holland | 2020-03-03 | Dutch COVID-19 response team | Erasmus Medical Center | David Nieuwenhuijse, Bas Oude Munnink, Reina Sikkema, Claudia Schapendonk, Irina Chestakova, Anne van der Linden, Mark Pronk, Pascal Lexmond, Corien Swaan, Manon Haverkate, Madelief Mollers, Mart Stein, Sandra Kengne Kamga Mobou, Jeroen van Kampen, Jolanda Voermans, Aura Timen, Corine GeurtsvanKessel, Annemiek van der Eijk, Richard Molenkamp, Marion Koopmans, on behalf of the Dutch national COVID-19 response team. |
| 201 | EPI_ISL_414577 | hCoV-19/Chile/Talca-1/2020 | *Homo sapiens* | South America / Chile / Talca | 2020-03-02 | Hospital de Talca, Chile | Instituto de Salud Publica de Chile | Andrés E. Castillo, Bárbara Parra, Paz Tapia, Alejandra Acevedo, Jaime Lagos, Winston Andrade, Loredana Arata, Gabriel Leal, Gisselle Barra, Carolina Tambley, Javier Tognarelli, Patricia Bustos, Soledad Ulloa, Rodrigo Fasce, Jorge Fernández. |
| 202 | EPI_ISL_414579 | hCoV-19/Chile/Santiago-1/2020 | *Homo sapiens* | South America / Chile / Santiago | 2020-03-03 | Clinica Alemana de Santiago, Chile | Instituto de Salud Publica de Chile | Andrés E. Castillo, Bárbara Parra, Paz Tapia, Alejandra Acevedo, Jaime Lagos, Winston Andrade, Loredana Arata, Gabriel Leal, Gisselle Barra, Carolina Tambley, Javier Tognarelli, Patricia Bustos, Soledad Ulloa, Rodrigo Fasce, Jorge Fernández. |
| 203 | EPI_ISL_414580 | hCoV-19/Chile/Santiago-2/2020 | *Homo sapiens* | South America / Chile / Santiago | 2020-03-05 | Clinica Santa Maria, Santiago, Chile | Instituto de Salud Publica de Chile | Andrés E. Castillo, Bárbara Parra, Paz Tapia, Alejandra Acevedo, Jaime Lagos, Winston Andrade, Loredana Arata, Gabriel Leal, Gisselle Barra, Carolina Tambley, Javier Tognarelli, Patricia Bustos, Soledad Ulloa, Rodrigo Fasce, Jorge Fernández. |
| 204 | EPI_ISL_414589 | hCoV-19/USA/MN2-MDH2/2020 | *Homo sapiens* | North America / USA / Minnesota | 2020-03-07 | Minnesota Department of Health, Public Health Laboratory | Minnesota Department of Health, Public Health Laboratory | Matt Plumb, Jake Garfin and Xiong Wang |
| 205 | EPI_ISL_414590 | hCoV-19/USA/MN3-MDH3/2020 | *Homo sapiens* | North America / USA / Minnesota | 2020-03-09 | Minnesota Department of Health, Public Health Laboratory | Minnesota Department of Health, Public Health Laboratory | Matt Plumb, Jake Garfin and Xiong Wang |
| 206 | EPI_ISL_414591 | hCoV-19/USA/WA-UW22/2020 | *Homo sapiens* | North America / USA / Washington / Kirkland | 2020-03-06 | UW Virology Lab | UW Virology Lab | Pavitra Roychoudhury, Hong Xie, Keith Jerome, Alexander Greninger |
| 207 | EPI_ISL_414592 | hCoV-19/USA/WA-UW23/2020 | *Homo sapiens* | North America / USA / Washington / Tacoma | 2020-03-06 | UW Virology Lab | UW Virology Lab | Pavitra Roychoudhury, Hong Xie, Keith Jerome, Alexander Greninger |
| 208 | EPI_ISL_414595 | hCoV-19/USA/WA-UW26/2020 | *Homo sapiens* | North America / USA / Washington / Kirkland | 2020-03-05 | UW Virology Lab | UW Virology Lab | Pavitra Roychoudhury, Hong Xie, Keith Jerome, Alexander Greninger |
| 209 | EPI_ISL_414600 | hCoV-19/France/GE1583/2020 | *Homo sapiens* | Europe / France / Grand-Est / Strasbourg | 2020-02-26 | Laboratoire de Virologie Institut de Virologie - INSERM U 1109 Hôpitaux Universitaires de Strasbourg | National Reference Center for Viruses of Respiratory Infections, Institut Pasteur, Paris | Mélnie Albert, Marion Barbet, Sylvie Behillil, Méline Bizard, Angela Brisebarre, Flora Donati Vincent Enouf, Maud Vanpeene, Sylvie van der Werf, Samira Fafi-Kremer |
| 210 | EPI_ISL_414601 | hCoV-19/France/N1620/2020 | *Homo sapiens* | Europe / France / Normandie / Rouen | 2020-02-27 | "Centre Hositalier Universitaire de Rouen Laboratoire de Virologie" | National Reference Center for Viruses of Respiratory Infections, Institut Pasteur, Paris | Mélnie Albert, Marion Barbet, Sylvie Behillil, Méline Bizard, Angela Brisebarre, Flora Donati Vincent Enouf, Maud Vanpeene, Sylvie van der Werf, Jean-Christophe Plantier |
| 211 | EPI_ISL_414618 | hCoV-19/USA/WA-UW31/2020 | *Homo sapiens* | North America / USA | 2020-03-08 | UW Virology Lab | UW Virology Lab | Pavitra Roychoudhury, Hong Xie, Keith Jerome, Alexander Greninger |
| 212 | EPI_ISL_414623 | hCoV-19/France/GE1583/2020 | *Homo sapiens* | Europe / France / Grand-Est / Strasbourg | 2020-02-25 | Laboratoire de Virologie Institut de Virologie - INSERM U 1109 Hôpitaux Universitaires de Strasbourg | National Reference Center for Viruses of Respiratory Infections, Institut Pasteur, Paris | Mélnie Albert, Marion Barbet, Sylvie Behillil, Méline Bizard, Angela Brisebarre, Flora Donati Vincent Enouf, Maud Vanpeene, Sylvie van der Werf, Samira Fafi-Kremer |
| 213 | EPI_ISL_414624 | hCoV-19/France/N1620/2020 | *Homo sapiens* | Europe / France / Normandie / Rouen | 2020-02-26 | Centre Hositalier Universitaire de Rouen Laboratoire de Virologie | National Reference Center for Viruses of Respiratory Infections, Institut Pasteur, Paris | Mélnie Albert, Marion Barbet, Sylvie Behillil, Méline Bizard, Angela Brisebarre, Flora Donati Vincent Enouf, Maud Vanpeene, Sylvie van der Werf, Jean-Christophe Plantier |
| 214 | EPI_ISL_414625 | hCoV-19/France/PL1643/2020 | *Homo sapiens* | Europe / France / Pays de la Loire / Nantes | 2020-02-26 | Centre Hospitalier Régional Universitaire de Nantes Laboratoire de Virologie | National Reference Center for Viruses of Respiratory Infections, Institut Pasteur, Paris | Mélnie Albert, Marion Barbet, Sylvie Behillil, Méline Bizard, Angela Brisebarre, Flora Donati Vincent Enouf, Maud Vanpeene, Sylvie van der Werf, Marianne Coste-Burel |

| | | | | | | | | |
|---|---|---|---|---|---|---|---|---|
| 215 | EPI_ISL_414626 | hCoV-19/France/HF1684/2020 | *Homo sapiens* | Europe / France / Hauts de France / Crépy en Valois | 2020-02-29 | unknown | National Reference Center for Viruses of Respiratory Infections, Institut Pasteur, Paris | Mélnie Albert, Marion Barbet, Sylvie Behillil, Méline Bizard, Angela Brisebarre, Flora Donati Vincent Enouf, Maud Vanpeene, Sylvie van der Werf |
| 216 | EPI_ISL_414627 | hCoV-19/France/HF1795/2020 | *Homo sapiens* | Europe / France / Hauts de France / Compiègne | 2020-03-02 | Centre Hospitalier Compiègne Laboratoire de Biologie | National Reference Center for Viruses of Respiratory Infections, Institut Pasteur, Paris | Mélnie Albert, Marion Barbet, Sylvie Behillil, Méline Bizard, Angela Brisebarre, Flora Donati Vincent Enouf, Maud Vanpeene, Sylvie van der Werf, Raulin Olivia |
| 217 | EPI_ISL_414629 | hCoV-19/France/HF1870/2020 | *Homo sapiens* | Europe / France / Hauts de France / Compiègne | 2020-03-03 | Centre Hospitalier Compiègne Laboratoire de Biologie | National Reference Center for Viruses of Respiratory Infections, Institut Pasteur, Paris | Mélnie Albert, Marion Barbet, Sylvie Behillil, Méline Bizard, Angela Brisebarre, Flora Donati Vincent Enouf, Maud Vanpeene, Sylvie van der Werf, Raulin Olivia |
| 218 | EPI_ISL_414630 | hCoV-19/France/HF1871/2020 | *Homo sapiens* | Europe / France / Hauts de France / Compiègne | 2020-03-03 | Centre Hospitalier Compiègne Laboratoire de Biologie | National Reference Center for Viruses of Respiratory Infections, Institut Pasteur, Paris | Mélnie Albert, Marion Barbet, Sylvie Behillil, Méline Bizard, Angela Brisebarre, Flora Donati Vincent Enouf, Maud Vanpeene, Sylvie van der Werf, Raulin Olivia |
| 219 | EPI_ISL_414631 | hCoV-19/France/GE1973/2020 | *Homo sapiens* | Europe / France / Grand-Est / Reims | 2020-03-04 | Hôpital Robert Debré Laboratoire de Virologie | National Reference Center for Viruses of Respiratory Infections, Institut Pasteur, Paris | Mélnie Albert, Marion Barbet, Sylvie Behillil, Méline Bizard, Angela Brisebarre, Flora Donati Vincent Enouf, Maud Vanpeene, Sylvie van der Werf, Laurent Andreoletti |
| 220 | EPI_ISL_414632 | hCoV-19/France/GE1977/2020 | *Homo sapiens* | Europe / France / Grand-Est / Reims | 2020-03-04 | Hôpital Robert Debré Laboratoire de Virologie | National Reference Center for Viruses of Respiratory Infections, Institut Pasteur, Paris | Mélnie Albert, Marion Barbet, Sylvie Behillil, Méline Bizard, Angela Brisebarre, Flora Donati Vincent Enouf, Maud Vanpeene, Sylvie van der Werf, Laurent Andreoletti |
| 221 | EPI_ISL_414633 | hCoV-19/France/IDF1980/2020 | *Homo sapiens* | Europe / France / Ile-de-France / Pontoise | 2020-03-04 | Centre Hospitalier René Dubois Laboratoire de Microbiologie - Bât A | National Reference Center for Viruses of Respiratory Infections, Institut Pasteur, Paris | Mélnie Albert, Marion Barbet, Sylvie Behillil, Méline Bizard, Angela Brisebarre, Flora Donati Vincent Enouf, Maud Vanpeene, Sylvie van der Werf, Pascale Martres |
| 222 | EPI_ISL_414635 | hCoV-19/France/HF1988/2020 | *Homo sapiens* | Europe / France / Hauts de France / Compiègne | 2020-03-04 | Centre Hospitalier Compiègne Laboratoire de Biologie | National Reference Center for Viruses of Respiratory Infections, Institut Pasteur, Paris | Mélnie Albert, Marion Barbet, Sylvie Behillil, Méline Bizard, Angela Brisebarre, Flora Donati Vincent Enouf, Maud Vanpeene, Sylvie van der Werf, Raulin Olivia |
| 223 | EPI_ISL_414637 | hCoV-19/France/HF1993/2020 | *Homo sapiens* | Europe / France / Hauts de France / Compiègne | 2020-03-04 | Centre Hospitalier Compiègne Laboratoire de Biologie | National Reference Center for Viruses of Respiratory Infections, Institut Pasteur, Paris | Mélnie Albert, Marion Barbet, Sylvie Behillil, Méline Bizard, Angela Brisebarre, Flora Donati Vincent Enouf, Maud Vanpeene, Sylvie van der Werf, Raulin Olivia |
| 224 | EPI_ISL_414638 | hCoV-19/France/HF1995/2020 | *Homo sapiens* | Europe / France / Hauts de France / Compiègne | 2020-03-04 | Centre Hospitalier Compiègne Laboratoire de Biologie | National Reference Center for Viruses of Respiratory Infections, Institut Pasteur, Paris | Mélnie Albert, Marion Barbet, Sylvie Behillil, Méline Bizard, Angela Brisebarre, Flora Donati Vincent Enouf, Maud Vanpeene, Sylvie van der Werf, Raulin Olivia |
| 225 | EPI_ISL_414641 | hCoV-19/Finland/FIN-313/2020 | *Homo sapiens* | Europe / Finland | 2020-03-05 | Department of Virology and Immunology, University of Helsinki and Helsinki University Hospital, Huslab Finland | Department of Virology, Faculty of Medicine, University of Helsinki, Helsinki, Finland | Teemu Smura, Hannimari Kallio-Kokko, Olli Vapalahti |
| 226 | EPI_ISL_414642 | hCoV-19/Finland/FIN-455/2020 | *Homo sapiens* | Europe / Finland | 2020-03-08 | Department of Virology and Immunology, University of Helsinki and Helsinki University Hospital, Huslab Finland | Department of Virology, Faculty of Medicine, University of Helsinki, Helsinki, Finland | Teemu Smura, Hannimari Kallio-Kokko, Olli Vapalahti |
| 227 | EPI_ISL_414643 | hCoV-19/Finland/FIN-508/2020 | *Homo sapiens* | Europe / Finland | 2020-03-07 | Department of Virology and Immunology, University of Helsinki and Helsinki University Hospital, Huslab Finland | Department of Virology, Faculty of Medicine, University of Helsinki, Helsinki, Finland | Teemu Smura, Hannimari Kallio-Kokko, Olli Vapalahti |
| 228 | EPI_ISL_414646 | hCoV-19/Finland/FIN-266/2020 | *Homo sapiens* | Europe / Finland | 2020-03-04 | Department of Virology and Immunology, University of Helsinki and Helsinki University Hospital, Huslab Finland | Department of Virology, Faculty of Medicine, University of Helsinki, Helsinki, Finland | Teemu Smura, Hannimari Kallio-Kokko, Olli Vapalahti |
| 229 | EPI_ISL_414648 | hCoV-19/USA/CA-PC101P/2020 | *Homo sapiens* | North America / USA / California / San Diego County | 2020-03-11 | Andersen Lab, The Scripps Research Institute | Andersen Lab, The Scripps Research Institute | Mark Zeller, Catie Anderson, Emily Spender, Sarah Topol, Raphaelle Klitting, Refugio Robles-Sikisaka, Karthik Gangavarapu, Laura Nicholson, Kristian Andersen |

| | | | | | | | | |
|---|---|---|---|---|---|---|---|---|
| 230 | EPI_ISL_414663 | hCoV-19/Guangzhou/GZMU0016/2020 | *Homo sapiens* | Asia / China / Guangdong / Guangzhou | 2020-02-25 | State Key Laboratory of Respiratory Disease, National Clinical Research Center for Respiratory Disease, Guangzhou Institute of Respiratory Health, the First Affiliated Hospital of Guangzhou Medical University | the First Affiliated Hospital of Guangzhou Medical University & BGI-Shenzhen | Zhao et al |
| 231 | EPI_ISL_414688 | hCoV-19/Guangzhou/GZMU0042/2020 | *Homo sapiens* | Asia / China / Guangdong / Guangzhou | 2020-02-25 | State Key Laboratory of Respiratory Disease, National Clinical Research Center for Respiratory Disease, Guangzhou Institute of Respiratory Health, the First Affiliated Hospital of Guangzhou Medical University | The First Affiliated Hospital of Guangzhou Medical University & BGI-Shenzhen | Zhao et al |
| 232 | EPI_ISL_414692 | hCoV-19/Guangzhou/GZMU0014/2020 | *Homo sapiens* | Asia / China / Guangdong / Guangzhou | 2020-02-25 | State Key Laboratory of Respiratory Disease, National Clinical Research Center for Respiratory Disease, Guangzhou Institute of Respiratory Health, the First Affiliated Hospital of Guangzhou Medical University | The First Affiliated Hospital of Guangzhou Medical University & BGI-Shenzhen | Zhao et al |
| 233 | EPI_ISL_414938 | hCoV-19/Shandong/LY005/2020 | *Homo sapiens* | Asia / China / Shandong | 2020-01-24 | Shandong Provincial Center for Disease Control and Prevention | Beijing Institute of Microbiology and Epidemiology | Xiao-Lin Jiang, Xiao-Li Zhang, Xiang-Na Zhao, Cun-Bao Li, Jie Lei, Zeng-Qiang Kou, Wen-Kui Sun, Yang Hang, Feng Gao, Sheng-Xiang Ji, Can-Fang Lin, Bo Pang, Ming-Xiao Yao, Guo-Lin Wang, Lin Yao, Li-Jun Duan, Xiao Wei, Dian-Ming Kang, Mai-Juan Ma |
| 234 | EPI_ISL_414940 | hCoV-19/Shandong/LY007/2020 | *Homo sapiens* | Asia / China / Shandong | 2020-01-25 | Shandong Provincial Center for Disease Control and Prevention | Beijing Institute of Microbiology and Epidemiology | Xiao-Lin Jiang, Xiao-Li Zhang, Xiang-Na Zhao, Cun-Bao Li, Jie Lei, Zeng-Qiang Kou, Wen-Kui Sun, Yang Hang, Feng Gao, Sheng-Xiang Ji, Can-Fang Lin, Bo Pang, Ming-Xiao Yao, Guo-Lin Wang, Lin Yao, Li-Jun Duan, Xiao Wei, Dian-Ming Kang, Mai-Juan Ma |
| 235 | EPI_ISL_414941 | hCoV-19/Shandong/LY008/2020 | *Homo sapiens* | Asia / China / Shandong | 2020-01-30 | Shandong Provincial Center for Disease Control and Prevention | Beijing Institute of Microbiology and Epidemiology | Xiao-Lin Jiang, Xiao-Li Zhang, Xiang-Na Zhao, Cun-Bao Li, Jie Lei, Zeng-Qiang Kou, Wen-Kui Sun, Yang Hang, Feng Gao, Sheng-Xiang Ji, Can-Fang Lin, Bo Pang, Ming-Xiao Yao, Guo-Lin Wang, Lin Yao, Li-Jun Duan, Xiao Wei, Dian-Ming Kang, Mai-Juan Ma |
| 236 | EPI_ISL_415153 | hCoV-19/Belgium/VLM-03011/2020 | *Homo sapiens* | Europe / Belgium / Huldenberg | 2020-03-03 | KU Leuven, Clinical and Epidemiological Virology | KU Leuven, Clinical and Epidemiological Virology | Bert Vanmechelen, Joan Marti-Carreras, Tony Wawina, Marc Van Ranst, Piet Maes |
| 237 | EPI_ISL_415154 | hCoV-19/Belgium/BM-03012/2020 | *Homo sapiens* | Europe / Belgium / Kraainem | 2020-03-01 | KU Leuven, Clincal and Epidemiological Virology | KU Leuven, Clincal and Epidemiological Virology | Bert Vanmechelen, Joan Marti-Careras, Tony Wawina, Marc Van Ranst, Piet Maes. |
| 238 | EPI_ISL_415155 | hCoV-19/Belgium/VAG-03013/2020 | *Homo sapiens* | Europe / Belgium / Huldenberg | 2020-03-01 | KU Leuven, Clinical and Epidemiological Virology | KU Leuven, Clinical and Epidemiological Virology | Bert Vanmechelen, Joan Marti-Carreras, Tony Wawina, Marc Van Ranst, Piet Maes |
| 239 | EPI_ISL_415156 | hCoV-19/Belgium/SH-03014/2020 | *Homo sapiens* | Europe / Belgium / Huldenberg | 2020-03-01 | KU Leuven, Clinical and Epidemiological Virology | KU Leuven, Clinical and Epidemiological Virology | Bert Vanmechelen, Joan Marti-Carreras, Tony Wawina, Piet Maes |
| 240 | EPI_ISL_415157 | hCoV-19/Belgium/BC-03016/2020 | *Homo sapiens* | Europe / Belgium / Sint-Niklaas | 2020-03-01 | KU Leuven, Clinical and Epidemiological Virology | KU Leuven, Clinical and Epidemiological Virology | Bert Vanmechelen, Joan Marti-Carreras, Tony Wawina, Piet Maes |
| 241 | EPI_ISL_415158 | hCoV-19/Belgium/QKJ-03015/2020 | *Homo sapiens* | Europe / Belgium / Brussels | 2020-03-01 | KU Leuven, Clinical and Epidemiological Virology | KU Leuven, Clinical and Epidemiological Virology | Bert Vanmechelen, Joan Marti-Carreras, Tony Wawina, Piet Maes |
| 242 | EPI_ISL_415159 | hCoV-19/Belgium/BA-02291/2020 | *Homo sapiens* | Europe / Belgium / Leuven | 2020-02-29 | KU Leuven, Clinical and Epidemiological Virology | KU Leuven, Clinical and Epidemiological Virology | Bert Vanmechelen, Joan Marti-Carreras, Tony Wawina, Piet Maes |

| | | | | | | | | |
|---|---|---|---|---|---|---|---|---|
| 243 | EPI_ISL_415454 | hCoV-19/Switzerland/GE1422/2020 | *Homo sapiens* | Europe / Switzerland | 2020-02-28 | Hôpitaux universitaires de Genève Laboratoire de Virologie | Hôpitaux universitaires de Genève Laboratoire de Virologie | Laubscher F. |
| 244 | EPI_ISL_415456 | hCoV-19/Switzerland/BE6651/2020 | *Homo sapiens* | Europe / Switzerland | 2020-02-29 | Hôpitaux universitaires de Genève Laboratoire de Virologie | Hôpitaux universitaires de Genève Laboratoire de Virologie | Laubscher F. |
| 245 | EPI_ISL_415457 | hCoV-19/Switzerland/AG7120/2020 | *Homo sapiens* | Europe / Switzerland | 2020-02-29 | Hôpitaux universitaires de Genève Laboratoire de Virologie | Hôpitaux universitaires de Genève Laboratoire de Virologie | Laubscher F. |
| 245 | EPI_ISL_416743 | hCoV-19/Czech Republic/ChVir1912/2020 | *Homo sapiens* | Europe / Czech Republic / Prague | 2020-03 | NRL for Influenza, Centrum Epidemiology and Microbiology of National Institute of Public Health, Czech Republic | Charite Universitaetsmedizin Berlin, Institute of Virology | Victor M Corman, Julia Schneider, Jörn Beheim-Schwarzbach, Talitha Veith, Barbara Muehlemann, Terry Jones, Akexander Nagy, Jaromira Vecerova, Dusan Trnka, Ludmila Novakova, Helena Jirincova, Christian Drosten |
| 246 | EPI_ISL_415458 | hCoV-19/Switzerland/GE8102/2020 | *Homo sapiens* | Europe / Switzerland | 2020-03-01 | Hôpitaux universitaires de Genève Laboratoire de Virologie | Hôpitaux universitaires de Genève Laboratoire de Virologie | Laubscher F. |
| 246 | EPI_ISL_416035 | hCoV-19/Brazil/SPBR-13/2020 | *Homo sapiens* | South America / Brazil / Sao Paulo / Sao Paulo | 2020-03-05 | National Influenza Center - Instituto Adolfo Lutz | Instituto Adolfo Lutz, Interdiciplinary Procedures Center, Strategic Laboratory | Claudio Tavares Sacchi, Claudia Regina Gonçalves, Carlos Henrique Camargo, Erica Valessa Ramos Gomes, Fabiana Cristina Pereira dos Santos, Daniela Bernardes Borges da Silva, Simone Guadagnucci Morillo, Adriano Abbud, Adriana Bugno, Maria do Carmo Sampaio Tavares Timenetsky, Terezinha Maria de Paiva |
| 247 | EPI_ISL_415460 | hCoV-19/Netherlands/Flevoland_1/2020 | *Homo sapiens* | Europe / Netherlands / Flevoland | 2020-03-09 | Dutch COVID-19 response team | Erasmus Medical Center | David Nieuwenhuijse, Bas Oude Munnink, Reina Sikkema, Claudia Schapendonk, Irina Chestakova, Anne van der Linden, Mark Pronk, Pascal Lexmond, Corien Swaan, Manon Haverkate, Madelief Mollers, Mart Stein, Sandra Kengne Kamga Mobou, Jeroen van Kampen, Jolanda Voermans, Aura Timen, Corine GeurtsvanKessel, Annemiek van der Eijk, Richard Molenkamp, Marion Koopmans, on behalf of the Dutch national COVID-19 response team. |
| 247 | EPI_ISL_415465 | hCoV-19/Netherlands/NA_1/2020 | *Homo sapiens* | Europe / Netherlands | 2020-03-10 | Dutch COVID-19 response team | Erasmus Medical Center | David Nieuwenhuijse, Bas Oude Munnink, Reina Sikkema, Claudia Schapendonk, Irina Chestakova, Anne van der Linden, Mark Pronk, Pascal Lexmond, Corien Swaan, Manon Haverkate, Madelief Mollers, Mart Stein, Sandra Kengne Kamga Mobou, Jeroen van Kampen, Jolanda Voermans, Aura Timen, Corine GeurtsvanKessel, Annemiek van der Eijk, Richard Molenkamp, Marion Koopmans, on behalf of the Dutch national COVID-19 response team. |
| 247 | EPI_ISL_415466 | hCoV-19/Netherlands/NA_10/2020 | *Homo sapiens* | Europe / Netherlands | 2020-03-09 | Dutch COVID-19 response team | Erasmus Medical Center | David Nieuwenhuijse, Bas Oude Munnink, Reina Sikkema, Claudia Schapendonk, Irina Chestakova, Anne van der Linden, Mark Pronk, Pascal Lexmond, Corien Swaan, Manon Haverkate, Madelief Mollers, Mart Stein, Sandra Kengne Kamga Mobou, Jeroen van Kampen, Jolanda Voermans, Aura Timen, Corine GeurtsvanKessel, Annemiek van der Eijk, Richard Molenkamp, Marion Koopmans, on behalf of the Dutch national COVID-19 response team. |
| 247 | EPI_ISL_415469 | hCoV-19/Netherlands/NA_13/2020 | *Homo sapiens* | Europe / Netherlands | 2020-03-10 | Dutch COVID-19 response team | Erasmus Medical Center | David Nieuwenhuijse, Bas Oude Munnink, Reina Sikkema, Claudia Schapendonk, Irina Chestakova, Anne van der Linden, Mark Pronk, Pascal Lexmond, Corien Swaan, Manon Haverkate, Madelief Mollers, Mart Stein, Sandra Kengne Kamga Mobou, Jeroen van Kampen, Jolanda Voermans, Aura Timen, Corine GeurtsvanKessel, Annemiek van der Eijk, Richard Molenkamp, Marion Koopmans, on behalf of the Dutch national COVID-19 response team. |

| | | | | | | | | |
|---|---|---|---|---|---|---|---|---|
| 247 | EPI_ISL_415515 | hCoV-19/Netherlands/NoordBrabant_59/2020 | *Homo sapiens* | Europe / Netherlands / Noord Brabant | 2020-03-11 | Dutch COVID-19 response team | Erasmus Medical Center | David Nieuwenhuijse, Bas Oude Munnink, Reina Sikkema, Claudia Schapendonk, Irina Chestakova, Anne van der Linden, Mark Pronk, Pascal Lexmond, Corien Swaan, Manon Haverkate, Madelief Mollers, Mart Stein, Sandra Kengne Kamga Mobou, Jeroen van Kampen, Jolanda Voermans, Aura Timen, Corine GeurtsvanKessel, Annemiek van der Eijk, Richard Molenkamp, Marion Koopmans, on behalf of the Dutch national COVID-19 response team. |
| 248 | EPI_ISL_415461 | hCoV-19/Netherlands/Gelderland_1/2020 | *Homo sapiens* | Europe / Netherlands / Gelderland | 2020-03-10 | Dutch COVID-19 response team | Erasmus Medical Center | David Nieuwenhuijse, Bas Oude Munnink, Reina Sikkema, Claudia Schapendonk, Irina Chestakova, Anne van der Linden, Mark Pronk, Pascal Lexmond, Corien Swaan, Manon Haverkate, Madelief Mollers, Mart Stein, Sandra Kengne Kamga Mobou, Jeroen van Kampen, Jolanda Voermans, Aura Timen, Corine GeurtsvanKessel, Annemiek van der Eijk, Richard Molenkamp, Marion Koopmans, on behalf of the Dutch national COVID-19 response team. |
| 248 | EPI_ISL_415526 | hCoV-19/Netherlands/Utrecht_17/2020 | *Homo sapiens* | Europe / Netherlands / Utrecht | 2020-03-10 | Dutch COVID-19 response team | Erasmus Medical Center | David Nieuwenhuijse, Bas Oude Munnink, Reina Sikkema, Claudia Schapendonk, Irina Chestakova, Anne van der Linden, Mark Pronk, Pascal Lexmond, Corien Swaan, Manon Haverkate, Madelief Mollers, Mart Stein, Sandra Kengne Kamga Mobou, Jeroen van Kampen, Jolanda Voermans, Aura Timen, Corine GeurtsvanKessel, Annemiek van der Eijk, Richard Molenkamp, Marion Koopmans, on behalf of the Dutch national COVID-19 response team. |
| 249 | EPI_ISL_415462 | hCoV-19/Netherlands/Gelderland_2/2020 | *Homo sapiens* | Europe / Netherlands / Gelderland | 2020-03-09 | Dutch COVID-19 response team | Erasmus Medical Center | David Nieuwenhuijse, Bas Oude Munnink, Reina Sikkema, Claudia Schapendonk, Irina Chestakova, Anne van der Linden, Mark Pronk, Pascal Lexmond, Corien Swaan, Manon Haverkate, Madelief Mollers, Mart Stein, Sandra Kengne Kamga Mobou, Jeroen van Kampen, Jolanda Voermans, Aura Timen, Corine GeurtsvanKessel, Annemiek van der Eijk, Richard Molenkamp, Marion Koopmans, on behalf of the Dutch national COVID-19 response team. |
| 249 | EPI_ISL_415518 | hCoV-19/Netherlands/NoordBrabant_62/2020 | *Homo sapiens* | Europe / Netherlands / Noord Brabant | 2020 | Dutch COVID-19 response team | Erasmus Medical Center | David Nieuwenhuijse, Bas Oude Munnink, Reina Sikkema, Claudia Schapendonk, Irina Chestakova, Anne van der Linden, Mark Pronk, Pascal Lexmond, Corien Swaan, Manon Haverkate, Madelief Mollers, Mart Stein, Sandra Kengne Kamga Mobou, Jeroen van Kampen, Jolanda Voermans, Aura Timen, Corine GeurtsvanKessel, Annemiek van der Eijk, Richard Molenkamp, Marion Koopmans, on behalf of the Dutch national COVID-19 response team. |
| 249 | EPI_ISL_415520 | hCoV-19/Netherlands/NoordBrabant_64/2020 | *Homo sapiens* | Europe / Netherlands / Noord Brabant | 2020 | Dutch COVID-19 response team | Erasmus Medical Center | David Nieuwenhuijse, Bas Oude Munnink, Reina Sikkema, Claudia Schapendonk, Irina Chestakova, Anne van der Linden, Mark Pronk, Pascal Lexmond, Corien Swaan, Manon Haverkate, Madelief Mollers, Mart Stein, Sandra Kengne Kamga Mobou, Jeroen van Kampen, Jolanda Voermans, Aura Timen, Corine GeurtsvanKessel, Annemiek van der Eijk, Richard Molenkamp, Marion Koopmans, on behalf of the Dutch national COVID-19 response team. |
| 250 | EPI_ISL_415464 | hCoV-19/Netherlands/Limburg_7/2020 | *Homo sapiens* | Europe / Netherlands / Limburg | 2020 | Dutch COVID-19 response team | Erasmus Medical Center | David Nieuwenhuijse, Bas Oude Munnink, Reina Sikkema, Claudia Schapendonk, Irina Chestakova, Anne van der Linden, Mark Pronk, Pascal Lexmond, Corien Swaan, Manon Haverkate, Madelief Mollers, Mart Stein, Sandra Kengne Kamga Mobou, Jeroen van Kampen, Jolanda Voermans, Aura Timen, Corine GeurtsvanKessel, Annemiek van der Eijk, Richard Molenkamp, Marion Koopmans, on behalf of the Dutch national COVID-19 response team. |

| | | | | | | | | |
|---|---|---|---|---|---|---|---|---|
| 251 | EPI_ISL_415467 | hCoV-19/Netherlands/NA_11/2020 | *Homo sapiens* | Europe / Netherlands | 2020-03-10 | Dutch COVID-19 response team | Erasmus Medical Center | David Nieuwenhuijse, Bas Oude Munnink, Reina Sikkema, Claudia Schapendonk, Irina Chestakova, Anne van der Linden, Mark Pronk, Pascal Lexmond, Corien Swaan, Manon Haverkate, Madelief Mollers, Mart Stein, Sandra Kengne Kamga Mobou, Jeroen van Kampen, Jolanda Voermans, Aura Timen, Corine GeurtsvanKessel, Annemiek van der Eijk, Richard Molenkamp, Marion Koopmans, on behalf of the Dutch national COVID-19 response team. |
| 251 | EPI_ISL_415483 | hCoV-19/Netherlands/NA_26/2020 | *Homo sapiens* | Europe / Netherlands | 2020-03-09 | Dutch COVID-19 response team | Erasmus Medical Center | David Nieuwenhuijse, Bas Oude Munnink, Reina Sikkema, Claudia Schapendonk, Irina Chestakova, Anne van der Linden, Mark Pronk, Pascal Lexmond, Corien Swaan, Manon Haverkate, Madelief Mollers, Mart Stein, Sandra Kengne Kamga Mobou, Jeroen van Kampen, Jolanda Voermans, Aura Timen, Corine GeurtsvanKessel, Annemiek van der Eijk, Richard Molenkamp, Marion Koopmans, on behalf of the Dutch national COVID-19 response team. |
| 252 | EPI_ISL_415468 | hCoV-19/Netherlands/NA_12/2020 | *Homo sapiens* | Europe / Netherlands | 2020-03-10 | Dutch COVID-19 response team | Erasmus Medical Center | David Nieuwenhuijse, Bas Oude Munnink, Reina Sikkema, Claudia Schapendonk, Irina Chestakova, Anne van der Linden, Mark Pronk, Pascal Lexmond, Corien Swaan, Manon Haverkate, Madelief Mollers, Mart Stein, Sandra Kengne Kamga Mobou, Jeroen van Kampen, Jolanda Voermans, Aura Timen, Corine GeurtsvanKessel, Annemiek van der Eijk, Richard Molenkamp, Marion Koopmans, on behalf of the Dutch national COVID-19 response team. |
| 252 | EPI_ISL_415511 | hCoV-19/Netherlands/NoordBrabant_55/2020 | *Homo sapiens* | Europe / Netherlands / Noord Brabant | 2020-03-09 | Dutch COVID-19 response team | Erasmus Medical Center | David Nieuwenhuijse, Bas Oude Munnink, Reina Sikkema, Claudia Schapendonk, Irina Chestakova, Anne van der Linden, Mark Pronk, Pascal Lexmond, Corien Swaan, Manon Haverkate, Madelief Mollers, Mart Stein, Sandra Kengne Kamga Mobou, Jeroen van Kampen, Jolanda Voermans, Aura Timen, Corine GeurtsvanKessel, Annemiek van der Eijk, Richard Molenkamp, Marion Koopmans, on behalf of the Dutch national COVID-19 response team. |
| 252 | EPI_ISL_415530 | hCoV-19/Netherlands/ZuidHolland_26/2020 | *Homo sapiens* | Europe / Netherlands / Zuid Holland | 2020-03-09 | Dutch COVID-19 response team | Erasmus Medical Center | David Nieuwenhuijse, Bas Oude Munnink, Reina Sikkema, Claudia Schapendonk, Irina Chestakova, Anne van der Linden, Mark Pronk, Pascal Lexmond, Corien Swaan, Manon Haverkate, Madelief Mollers, Mart Stein, Sandra Kengne Kamga Mobou, Jeroen van Kampen, Jolanda Voermans, Aura Timen, Corine GeurtsvanKessel, Annemiek van der Eijk, Richard Molenkamp, Marion Koopmans, on behalf of the Dutch national COVID-19 response team. |
| 253 | EPI_ISL_415471 | hCoV-19/Netherlands/NA_15/2020 | *Homo sapiens* | Europe / Netherlands | 2020-03-11 | Dutch COVID-19 response team | Erasmus Medical Center | David Nieuwenhuijse, Bas Oude Munnink, Reina Sikkema, Claudia Schapendonk, Irina Chestakova, Anne van der Linden, Mark Pronk, Pascal Lexmond, Corien Swaan, Manon Haverkate, Madelief Mollers, Mart Stein, Sandra Kengne Kamga Mobou, Jeroen van Kampen, Jolanda Voermans, Aura Timen, Corine GeurtsvanKessel, Annemiek van der Eijk, Richard Molenkamp, Marion Koopmans, on behalf of the Dutch national COVID-19 response team. |
| 254 | EPI_ISL_415472 | hCoV-19/Netherlands/NA_16/2020 | *Homo sapiens* | Europe / Netherlands | 2020-03-11 | Dutch COVID-19 response team | Erasmus Medical Center | David Nieuwenhuijse, Bas Oude Munnink, Reina Sikkema, Claudia Schapendonk, Irina Chestakova, Anne van der Linden, Mark Pronk, Pascal Lexmond, Corien Swaan, Manon Haverkate, Madelief Mollers, Mart Stein, Sandra Kengne Kamga Mobou, Jeroen van Kampen, Jolanda Voermans, Aura Timen, Corine GeurtsvanKessel, Annemiek van der Eijk, Richard Molenkamp, Marion Koopmans, on behalf of the Dutch national COVID-19 response team. |

| | | | | | | | | |
|---|---|---|---|---|---|---|---|---|
| 255 | EPI_ISL_415473 | hCoV-19/Netherlands/NA_17/2020 | *Homo sapiens* | Europe / Netherlands | 2020-03-09 | Dutch COVID-19 response team | Erasmus Medical Center | David Nieuwenhuijse, Bas Oude Munnink, Reina Sikkema, Claudia Schapendonk, Irina Chestakova, Anne van der Linden, Mark Pronk, Pascal Lexmond, Corien Swaan, Manon Haverkate, Madelief Mollers, Mart Stein, Sandra Kengne Kamga Mobou, Jeroen van Kampen, Jolanda Voermans, Aura Timen, Corine GeurtsvanKessel, Annemiek van der Eijk, Richard Molenkamp, Marion Koopmans, on behalf of the Dutch national COVID-19 response team. |
| 255 | EPI_ISL_415474 | hCoV-19/Netherlands/NA_18/2020 | *Homo sapiens* | Europe / Netherlands | 2020-03-09 | Dutch COVID-19 response team | Erasmus Medical Center | David Nieuwenhuijse, Bas Oude Munnink, Reina Sikkema, Claudia Schapendonk, Irina Chestakova, Anne van der Linden, Mark Pronk, Pascal Lexmond, Corien Swaan, Manon Haverkate, Madelief Mollers, Mart Stein, Sandra Kengne Kamga Mobou, Jeroen van Kampen, Jolanda Voermans, Aura Timen, Corine GeurtsvanKessel, Annemiek van der Eijk, Richard Molenkamp, Marion Koopmans, on behalf of the Dutch national COVID-19 response team. |
| 255 | EPI_ISL_415499 | hCoV-19/Netherlands/NoordBrabant_41/2020 | *Homo sapiens* | Europe / Netherlands / Noord Brabant | 2020 | Dutch COVID-19 response team | Erasmus Medical Center | David Nieuwenhuijse, Bas Oude Munnink, Reina Sikkema, Claudia Schapendonk, Irina Chestakova, Anne van der Linden, Mark Pronk, Pascal Lexmond, Corien Swaan, Manon Haverkate, Madelief Mollers, Mart Stein, Sandra Kengne Kamga Mobou, Jeroen van Kampen, Jolanda Voermans, Aura Timen, Corine GeurtsvanKessel, Annemiek van der Eijk, Richard Molenkamp, Marion Koopmans, on behalf of the Dutch national COVID-19 response team. |
| 256 | EPI_ISL_415475 | hCoV-19/Netherlands/NA_19/2020 | *Homo sapiens* | Europe / Netherlands | 2020-03-12 | Dutch COVID-19 response team | Erasmus Medical Center | David Nieuwenhuijse, Bas Oude Munnink, Reina Sikkema, Claudia Schapendonk, Irina Chestakova, Anne van der Linden, Mark Pronk, Pascal Lexmond, Corien Swaan, Manon Haverkate, Madelief Mollers, Mart Stein, Sandra Kengne Kamga Mobou, Jeroen van Kampen, Jolanda Voermans, Aura Timen, Corine GeurtsvanKessel, Annemiek van der Eijk, Richard Molenkamp, Marion Koopmans, on behalf of the Dutch national COVID-19 response team. |
| 257 | EPI_ISL_415476 | hCoV-19/Netherlands/NA_2/2020 | *Homo sapiens* | Europe / Netherlands | 2020-03-10 | Dutch COVID-19 response team | Erasmus Medical Center | David Nieuwenhuijse, Bas Oude Munnink, Reina Sikkema, Claudia Schapendonk, Irina Chestakova, Anne van der Linden, Mark Pronk, Pascal Lexmond, Corien Swaan, Manon Haverkate, Madelief Mollers, Mart Stein, Sandra Kengne Kamga Mobou, Jeroen van Kampen, Jolanda Voermans, Aura Timen, Corine GeurtsvanKessel, Annemiek van der Eijk, Richard Molenkamp, Marion Koopmans, on behalf of the Dutch national COVID-19 response team. |
| 258 | EPI_ISL_415478 | hCoV-19/Netherlands/NA_21/2020 | *Homo sapiens* | Europe / Netherlands | 2020-03-08 | Dutch COVID-19 response team | Erasmus Medical Center | David Nieuwenhuijse, Bas Oude Munnink, Reina Sikkema, Claudia Schapendonk, Irina Chestakova, Anne van der Linden, Mark Pronk, Pascal Lexmond, Corien Swaan, Manon Haverkate, Madelief Mollers, Mart Stein, Sandra Kengne Kamga Mobou, Jeroen van Kampen, Jolanda Voermans, Aura Timen, Corine GeurtsvanKessel, Annemiek van der Eijk, Richard Molenkamp, Marion Koopmans, on behalf of the Dutch national COVID-19 response team. |
| 259 | EPI_ISL_415481 | hCoV-19/Netherlands/NA_24/2020 | *Homo sapiens* | Europe / Netherlands | 2020-03-08 | Dutch COVID-19 response team | Erasmus Medical Center | David Nieuwenhuijse, Bas Oude Munnink, Reina Sikkema, Claudia Schapendonk, Irina Chestakova, Anne van der Linden, Mark Pronk, Pascal Lexmond, Corien Swaan, Manon Haverkate, Madelief Mollers, Mart Stein, Sandra Kengne Kamga Mobou, Jeroen van Kampen, Jolanda Voermans, Aura Timen, Corine GeurtsvanKessel, Annemiek van der Eijk, Richard Molenkamp, Marion Koopmans, on behalf of the Dutch national COVID-19 response team. |

| | | | | | | | | |
|---|---|---|---|---|---|---|---|---|
| 260 | EPI_ISL_415482 | hCoV-19/Netherlands/NA_25/2020 | *Homo sapiens* | Europe / Netherlands | 2020-03-09 | Dutch COVID-19 response team | Erasmus Medical Center | David Nieuwenhuijse, Bas Oude Munnink, Reina Sikkema, Claudia Schapendonk, Irina Chestakova, Anne van der Linden, Mark Pronk, Pascal Lexmond, Corien Swaan, Manon Haverkate, Madelief Mollers, Mart Stein, Sandra Kengne Kamga Mobou, Jeroen van Kampen, Jolanda Voermans, Aura Timen, Corine GeurtsvanKessel, Annemiek van der Eijk, Richard Molenkamp, Marion Koopmans, on behalf of the Dutch national COVID-19 response team. |
| 261 | EPI_ISL_415485 | hCoV-19/Netherlands/NA_28/2020 | *Homo sapiens* | Europe / Netherlands | 2020-03-12 | Dutch COVID-19 response team | Erasmus Medical Center | David Nieuwenhuijse, Bas Oude Munnink, Reina Sikkema, Claudia Schapendonk, Irina Chestakova, Anne van der Linden, Mark Pronk, Pascal Lexmond, Corien Swaan, Manon Haverkate, Madelief Mollers, Mart Stein, Sandra Kengne Kamga Mobou, Jeroen van Kampen, Jolanda Voermans, Aura Timen, Corine GeurtsvanKessel, Annemiek van der Eijk, Richard Molenkamp, Marion Koopmans, on behalf of the Dutch national COVID-19 response team. |
| 262 | EPI_ISL_415486 | hCoV-19/Netherlands/NA_29/2020 | *Homo sapiens* | Europe / Netherlands | 2020-03-13 | Dutch COVID-19 response team | Erasmus Medical Center | David Nieuwenhuijse, Bas Oude Munnink, Reina Sikkema, Claudia Schapendonk, Irina Chestakova, Anne van der Linden, Mark Pronk, Pascal Lexmond, Corien Swaan, Manon Haverkate, Madelief Mollers, Mart Stein, Sandra Kengne Kamga Mobou, Jeroen van Kampen, Jolanda Voermans, Aura Timen, Corine GeurtsvanKessel, Annemiek van der Eijk, Richard Molenkamp, Marion Koopmans, on behalf of the Dutch national COVID-19 response team. |
| 263 | EPI_ISL_415487 | hCoV-19/Netherlands/NA_30/2020 | *Homo sapiens* | Europe / Netherlands | 2020-03-13 | Dutch COVID-19 response team | Erasmus Medical Center | David Nieuwenhuijse, Bas Oude Munnink, Reina Sikkema, Claudia Schapendonk, Irina Chestakova, Anne van der Linden, Mark Pronk, Pascal Lexmond, Corien Swaan, Manon Haverkate, Madelief Mollers, Mart Stein, Sandra Kengne Kamga Mobou, Jeroen van Kampen, Jolanda Voermans, Aura Timen, Corine GeurtsvanKessel, Annemiek van der Eijk, Richard Molenkamp, Marion Koopmans, on behalf of the Dutch national COVID-19 response team. |
| 264 | EPI_ISL_415488 | hCoV-19/Netherlands/NA_31/2020 | *Homo sapiens* | Europe / Netherlands | 2020-03-13 | Dutch COVID-19 response team | Erasmus Medical Center | David Nieuwenhuijse, Bas Oude Munnink, Reina Sikkema, Claudia Schapendonk, Irina Chestakova, Anne van der Linden, Mark Pronk, Pascal Lexmond, Corien Swaan, Manon Haverkate, Madelief Mollers, Mart Stein, Sandra Kengne Kamga Mobou, Jeroen van Kampen, Jolanda Voermans, Aura Timen, Corine GeurtsvanKessel, Annemiek van der Eijk, Richard Molenkamp, Marion Koopmans, on behalf of the Dutch national COVID-19 response team. |
| 265 | EPI_ISL_415491 | hCoV-19/Netherlands/NA_34/2020 | *Homo sapiens* | Europe / Netherlands | 2020-03-07 | Dutch COVID-19 response team | Erasmus Medical Center | David Nieuwenhuijse, Bas Oude Munnink, Reina Sikkema, Claudia Schapendonk, Irina Chestakova, Anne van der Linden, Mark Pronk, Pascal Lexmond, Corien Swaan, Manon Haverkate, Madelief Mollers, Mart Stein, Sandra Kengne Kamga Mobou, Jeroen van Kampen, Jolanda Voermans, Aura Timen, Corine GeurtsvanKessel, Annemiek van der Eijk, Richard Molenkamp, Marion Koopmans, on behalf of the Dutch national COVID-19 response team. |
| 266 | EPI_ISL_415492 | hCoV-19/Netherlands/NA_35/2020 | *Homo sapiens* | Europe / Netherlands | 2020-03-10 | Dutch COVID-19 response team | Erasmus Medical Center | David Nieuwenhuijse, Bas Oude Munnink, Reina Sikkema, Claudia Schapendonk, Irina Chestakova, Anne van der Linden, Mark Pronk, Pascal Lexmond, Corien Swaan, Manon Haverkate, Madelief Mollers, Mart Stein, Sandra Kengne Kamga Mobou, Jeroen van Kampen, Jolanda Voermans, Aura Timen, Corine GeurtsvanKessel, Annemiek van der Eijk, Richard Molenkamp, Marion Koopmans, on behalf of the Dutch national COVID-19 response team. |

| | | | | | | | | |
|---|---|---|---|---|---|---|---|---|
| 267 | EPI_ISL_415495 | hCoV-19/Netherlands/NA_6/2020 | *Homo sapiens* | Europe / Netherlands | 2020-03-10 | Dutch COVID-19 response team | Erasmus Medical Center | David Nieuwenhuijse, Bas Oude Munnink, Reina Sikkema, Claudia Schapendonk, Irina Chestakova, Anne van der Linden, Mark Pronk, Pascal Lexmond, Corien Swaan, Manon Haverkate, Madelief Mollers, Mart Stein, Sandra Kengne Kamga Mobou, Jeroen van Kampen, Jolanda Voermans, Aura Timen, Corine GeurtsvanKessel, Annemiek van der Eijk, Richard Molenkamp, Marion Koopmans, on behalf of the Dutch national COVID-19 response team. |
| 268 | EPI_ISL_415503 | hCoV-19/Netherlands/NoordBrabant_46/2020 | *Homo sapiens* | Europe / Netherlands / Noord Brabant | 2020-03-11 | Dutch COVID-19 response team | Erasmus Medical Center | David Nieuwenhuijse, Bas Oude Munnink, Reina Sikkema, Claudia Schapendonk, Irina Chestakova, Anne van der Linden, Mark Pronk, Pascal Lexmond, Corien Swaan, Manon Haverkate, Madelief Mollers, Mart Stein, Sandra Kengne Kamga Mobou, Jeroen van Kampen, Jolanda Voermans, Aura Timen, Corine GeurtsvanKessel, Annemiek van der Eijk, Richard Molenkamp, Marion Koopmans, on behalf of the Dutch national COVID-19 response team. |
| 269 | EPI_ISL_415512 | hCoV-19/Netherlands/NoordBrabant_56/2020 | *Homo sapiens* | Europe / Netherlands / Noord Brabant | 2020-03-09 | Dutch COVID-19 response team | Erasmus Medical Center | David Nieuwenhuijse, Bas Oude Munnink, Reina Sikkema, Claudia Schapendonk, Irina Chestakova, Anne van der Linden, Mark Pronk, Pascal Lexmond, Corien Swaan, Manon Haverkate, Madelief Mollers, Mart Stein, Sandra Kengne Kamga Mobou, Jeroen van Kampen, Jolanda Voermans, Aura Timen, Corine GeurtsvanKessel, Annemiek van der Eijk, Richard Molenkamp, Marion Koopmans, on behalf of the Dutch national COVID-19 response team. |
| 270 | EPI_ISL_415525 | hCoV-19/Netherlands/NoordHolland_3/2020 | *Homo sapiens* | Europe / Netherlands / Noord Holland | 2020-03-12 | Dutch COVID-19 response team | Erasmus Medical Center | David Nieuwenhuijse, Bas Oude Munnink, Reina Sikkema, Claudia Schapendonk, Irina Chestakova, Anne van der Linden, Mark Pronk, Pascal Lexmond, Corien Swaan, Manon Haverkate, Madelief Mollers, Mart Stein, Sandra Kengne Kamga Mobou, Jeroen van Kampen, Jolanda Voermans, Aura Timen, Corine GeurtsvanKessel, Annemiek van der Eijk, Richard Molenkamp, Marion Koopmans, on behalf of the Dutch national COVID-19 response team. |
| 271 | EPI_ISL_415527 | hCoV-19/Netherlands/Utrecht_18/2020 | *Homo sapiens* | Europe / Netherlands / Utrecht | 2020-03-12 | Dutch COVID-19 response team | Erasmus Medical Center | David Nieuwenhuijse, Bas Oude Munnink, Reina Sikkema, Claudia Schapendonk, Irina Chestakova, Anne van der Linden, Mark Pronk, Pascal Lexmond, Corien Swaan, Manon Haverkate, Madelief Mollers, Mart Stein, Sandra Kengne Kamga Mobou, Jeroen van Kampen, Jolanda Voermans, Aura Timen, Corine GeurtsvanKessel, Annemiek van der Eijk, Richard Molenkamp, Marion Koopmans, on behalf of the Dutch national COVID-19 response team. |
| 272 | EPI_ISL_415531 | hCoV-19/Netherlands/ZuidHolland_27/2020 | *Homo sapiens* | Europe / Netherlands / Zuid Holland | 2020-03-09 | Dutch COVID-19 response team | Erasmus Medical Center | David Nieuwenhuijse, Bas Oude Munnink, Reina Sikkema, Claudia Schapendonk, Irina Chestakova, Anne van der Linden, Mark Pronk, Pascal Lexmond, Corien Swaan, Manon Haverkate, Madelief Mollers, Mart Stein, Sandra Kengne Kamga Mobou, Jeroen van Kampen, Jolanda Voermans, Aura Timen, Corine GeurtsvanKessel, Annemiek van der Eijk, Richard Molenkamp, Marion Koopmans, on behalf of the Dutch national COVID-19 response team. |
| 273 | EPI_ISL_415532 | hCoV-19/Netherlands/ZuidHolland_28/2020 | *Homo sapiens* | Europe / Netherlands / Zuid Holland | 2020-03-09 | Dutch COVID-19 response team | Erasmus Medical Center | David Nieuwenhuijse, Bas Oude Munnink, Reina Sikkema, Claudia Schapendonk, Irina Chestakova, Anne van der Linden, Mark Pronk, Pascal Lexmond, Corien Swaan, Manon Haverkate, Madelief Mollers, Mart Stein, Sandra Kengne Kamga Mobou, Jeroen van Kampen, Jolanda Voermans, Aura Timen, Corine GeurtsvanKessel, Annemiek van der Eijk, Richard Molenkamp, Marion Koopmans, on behalf of the Dutch national COVID-19 response team. |

| | | | | | | | | |
|---|---|---|---|---|---|---|---|---|
| 274 | EPI_ISL_415535 | hCoV-19/Netherlands/ZuidHolland_31/2020 | *Homo sapiens* | Europe / Netherlands / Zuid Holland | 2020-03-12 | Dutch COVID-19 response team | Erasmus Medical Center | David Nieuwenhuijse, Bas Oude Munnink, Reina Sikkema, Claudia Schapendonk, Irina Chestakova, Anne van der Linden, Mark Pronk, Pascal Lexmond, Corien Swaan, Manon Haverkate, Madelief Mollers, Mart Stein, Sandra Kengne Kamga Mobou, Jeroen van Kampen, Jolanda Voermans, Aura Timen, Corine GeurtsvanKessel, Annemiek van der Eijk, Richard Molenkamp, Marion Koopmans, on behalf of the Dutch national COVID-19 response team. |
| 275 | EPI_ISL_415539 | hCoV-19/USA/UPHL-01/2020 | *Homo sapiens* | North America / USA / Utah | 2020-03-13 | Utah Public Health Laboratory | Utah Public Health Laboratory | Erin Young, Kelly Oakeson |
| 276 | EPI_ISL_415543 | hCoV-19/USA/UPHL-05/2020 | *Homo sapiens* | North America / USA / Utah | 2020-03-13 | Utah Public Health Laboratory | Utah Public Health Laboratory | Erin Young, Kelly Oakeson |
| 276 | EPI_ISL_415544 | hCoV-19/USA/UPHL-06/2020 | *Homo sapiens* | North America / USA / Utah | 2020-03-13 | Utah Public Health Laboratory | Utah Public Health Laboratory | Erin Young, Kelly Oakeson |
| 277 | EPI_ISL_415581 | hCoV-19/Canada/BC_02421/2020 | *Homo sapiens* | North America / Canada / British Columbia | 2020-03-01 | BCCDC Public Health Laboratory | BCCDC Public Health Laboratory | Harrigan, Prystajecky, Krajden, Lee, Kamelian, Lapointe, Choi, Hoang, Sekirov, Levett, Tyson, Snutch, Loman, Quick, Li, Gilmour |
| 278 | EPI_ISL_415583 | hCoV-19/Canada/BC_40860/2020 | *Homo sapiens* | North America / Canada / British Columbia | 2020-03-03 | BCCDC Public Health Laboratory | BCCDC Public Health Laboratory | Harrigan, Prystajecky, Krajden, Lee, Kamelian, Lapointe, Choi, Hoang, Sekirov, Levett, Tyson, Snutch, Loman, Quick, Li, Gilmour |
| 279 | EPI_ISL_415591 | hCoV-19/USA/WA-UW63/2020 | *Homo sapiens* | North America / USA / Washington | 2020-03-10 | UW Virology Lab | UW Virology Lab | Pavitra Roychoudhury, Hong Xie, Keith Jerome, Alexander Greninger |
| 280 | EPI_ISL_415594 | hCoV-19/USA/WA-UW66/2020 | *Homo sapiens* | North America / USA / Washington | 2020-03-10 | UW Virology Lab | UW Virology Lab | Pavitra Roychoudhury, Hong Xie, Keith Jerome, Alexander Greninger |
| 281 | EPI_ISL_415595 | hCoV-19/USA/WA-UW67/2020 | *Homo sapiens* | North America / USA / Washington | 2020-03-09 | UW Virology Lab | UW Virology Lab | Pavitra Roychoudhury, Hong Xie, Keith Jerome, Alexander Greninger |
| 282 | EPI_ISL_415596 | hCoV-19/USA/WA-UW68/2020 | *Homo sapiens* | North America / USA / Washington | 2020-03-09 | UW Virology Lab | UW Virology Lab | Pavitra Roychoudhury, Hong Xie, Keith Jerome, Alexander Greninger |
| 283 | EPI_ISL_415597 | hCoV-19/USA/WA-UW69/2020 | *Homo sapiens* | North America / USA / Washington / Seattle | 2020-03-10 | UW Virology Lab | UW Virology Lab | Pavitra Roychoudhury, Hong Xie, Keith Jerome, Alexander Greninger |
| 284 | EPI_ISL_415598 | hCoV-19/USA/WA-UW70/2020 | *Homo sapiens* | North America / USA / Washington | 2020-03-10 | UW Virology Lab | UW Virology Lab | Pavitra Roychoudhury, Hong Xie, Keith Jerome, Alexander Greninger |
| 285 | EPI_ISL_415601 | hCoV-19/USA/WA-UW73/2020 | *Homo sapiens* | North America / USA / Washington | 2020-03-10 | UW Virology Lab | UW Virology Lab | Pavitra Roychoudhury, Hong Xie, Keith Jerome, Alexander Greninger |
| 286 | EPI_ISL_415602 | hCoV-19/USA/WA-UW74/2020 | *Homo sapiens* | North America / USA / Washington / Seattle | 2020-03-10 | UW Virology Lab | UW Virology Lab | Pavitra Roychoudhury, Hong Xie, Keith Jerome, Alexander Greninger |
| 287 | EPI_ISL_415603 | hCoV-19/USA/WA-UW75/2020 | *Homo sapiens* | North America / USA / Washington / Seattle | 2020-03-10 | UW Virology Lab | UW Virology Lab | Pavitra Roychoudhury, Hong Xie, Keith Jerome, Alexander Greninger |
| 288 | EPI_ISL_415604 | hCoV-19/USA/WA-UW76/2020 | *Homo sapiens* | North America / USA / Washington | 2020-03-10 | UW Virology Lab | UW Virology Lab | Pavitra Roychoudhury, Hong Xie, Keith Jerome, Alexander Greninger |
| 289 | EPI_ISL_415606 | hCoV-19/USA/WA-UW41/2020 | *Homo sapiens* | North America / USA / Washington | 2020-03-08 | UW Virology Lab | UW Virology Lab | Pavitra Roychoudhury, Hong Xie, Keith Jerome, Alexander Greninger |

| | | | | | | | | |
|---|---|---|---|---|---|---|---|---|
| 290 | EPI_ISL_415608 | hCoV-19/USA/WA-UW43/2020 | *Homo sapiens* | North America / USA / Washington | 2020-03-08 | UW Virology Lab | UW Virology Lab | Pavitra Roychoudhury, Hong Xie, Keith Jerome, Alexander Greninger |
| 291 | EPI_ISL_415609 | hCoV-19/USA/WA-UW44/2020 | *Homo sapiens* | North America / USA / Washington | 2020-03-08 | UW Virology Lab | UW Virology Lab | Pavitra Roychoudhury, Hong Xie, Keith Jerome, Alexander Greninger |
| 292 | EPI_ISL_415611 | hCoV-19/USA/WA-UW46/2020 | *Homo sapiens* | North America / USA / Washington | 2020-03-09 | UW Virology Lab | UW Virology Lab | Pavitra Roychoudhury, Hong Xie, Keith Jerome, Alexander Greninger |
| 293 | EPI_ISL_415615 | hCoV-19/USA/WA-UW50/2020 | *Homo sapiens* | North America / USA / Washington | 2020-03-08 | UW Virology Lab | UW Virology Lab | Pavitra Roychoudhury, Hong Xie, Keith Jerome, Alexander Greninger |
| 294 | EPI_ISL_415617 | hCoV-19/USA/WA-UW52/2020 | *Homo sapiens* | North America / USA / Washington | 2020-03-09 | UW Virology Lab | UW Virology Lab | Pavitra Roychoudhury, Hong Xie, Keith Jerome, Alexander Greninger |
| 295 | EPI_ISL_415619 | hCoV-19/USA/WA-UW54/2020 | *Homo sapiens* | North America / USA / Washington | 2020-03-09 | UW Virology Lab | UW Virology Lab | Pavitra Roychoudhury, Hong Xie, Keith Jerome, Alexander Greninger |
| 296 | EPI_ISL_415620 | hCoV-19/USA/WA-UW55/2020 | *Homo sapiens* | North America / USA / Washington | 2020-03-09 | UW Virology Lab | UW Virology Lab | Pavitra Roychoudhury, Hong Xie, Keith Jerome, Alexander Greninger |
| 297 | EPI_ISL_415621 | hCoV-19/USA/WA-UW56/2020 | *Homo sapiens* | North America / USA / Washington / Seattle | 2020-03-09 | UW Virology Lab | UW Virology Lab | Pavitra Roychoudhury, Hong Xie, Keith Jerome, Alexander Greninger |
| 298 | EPI_ISL_415622 | hCoV-19/USA/WA-UW57/2020 | *Homo sapiens* | North America / USA / Washington | 2020-03-09 | UW Virology Lab | UW Virology Lab | Pavitra Roychoudhury, Hong Xie, Keith Jerome, Alexander Greninger |
| 299 | EPI_ISL_415624 | hCoV-19/USA/WA-UW59/2020 | *Homo sapiens* | North America / USA / Washington | 2020-03-09 | UW Virology Lab | UW Virology Lab | Pavitra Roychoudhury, Hong Xie, Keith Jerome, Alexander Greninger |
| 300 | EPI_ISL_415625 | hCoV-19/USA/WA-UW60/2020 | *Homo sapiens* | North America / USA / Washington | 2020-03-09 | UW Virology Lab | UW Virology Lab | Pavitra Roychoudhury, Hong Xie, Keith Jerome, Alexander Greninger |
| 301 | EPI_ISL_415627 | hCoV-19/USA/WA-UW62/2020 | *Homo sapiens* | North America / USA / Washington | 2020-03-09 | UW Virology Lab | UW Virology Lab | Pavitra Roychoudhury, Hong Xie, Keith Jerome, Alexander Greninger |
| 302 | EPI_ISL_415629 | hCoV-19/Scotland/EDB004/2020 | *Homo sapiens* | Europe / United Kingdom / Scotland | 2020-03-04 | Virology Department, Royal Infirmary of Edinburgh, NHS Lothian | Virology Department, Royal Infirmary of Edinburgh, NHS Lothian | McHugh M, Dewar R, O'Toole Á, Rambaut A, Williams TC, Templeton K |
| 303 | EPI_ISL_415630 | hCoV-19/Scotland/CVR07/2020 | *Homo sapiens* | Europe / United Kingdom / Scotland | 2020-03-09 | West of Scotland Specialist Virology Centre, NHSGGC | MRC-University of Glasgow Centre for Virus Research | Kathy Smollett, Daniel Mair, Stephen Carmichael, Ana da Silva Filipe; Richard Orton, David L Robertson; Alasdair MacLean, Rory Gunson; Natasha Jesudason, Kathy Li, Antonia Ho; Emma Thomson. |
| 304 | EPI_ISL_415631 | hCoV-19/Scotland/CVR10/2020 | *Homo sapiens* | Europe / United Kingdom / Scotland | 2020-03-10 | West of Scotland Specialist Virology Centre, NHSGGC | MRC-University of Glasgow Centre for Virus Research | Kathy Smollett, Daniel Mair, Stephen Carmichael, Ana da Silva Filipe; Richard Orton, David L Robertson; Alasdair MacLean, Rory Gunson; Natasha Jesudason, Kathy Li, Antonia Ho; Emma Thomson. |
| 305 | EPI_ISL_415640 | hCoV-19/Scotland/EDB003/2020 | *Homo sapiens* | Europe / United Kingdom / Scotland | 2020-03-08 | Virology Department, Royal Infirmary of Edinburgh, NHS Lothian | Virology Department, Royal Infirmary of Edinburgh, NHS Lothian | McHugh M, Dewar R, O'Toole Á, Rambaut A, Williams TC, Templeton K |

| | | | | | | | | |
|---|---|---|---|---|---|---|---|---|
| 306 | EPI_ISL_415641 | hCoV-19/Georgia/Tb-54/2020 | *Homo sapiens* | Asia / Georgia / Tbilisi | 2020-02-27 | R. G. Lugar Center for Public Health Research, National Center for Disease Control and Public Health (NCDC) of Georgia. | R. G. Lugar Center for Public Health Research, National Center for Disease Control and Public Health (NCDC) of Georgia. | Nato Kotaria, Marine Murtskhvaladze, Ann Machablishvili, Lela Sabadze, Mari Gavashelidze, Ana Papkiauri, Meri Pantsulaia, Gvantsa Brachveli, Tata Imnadze, Tamar Jashiashvili, Tea Tevdoradze, Ketevan Sidamonidze, Ekaterine Khmaladze, Ekaterine Zhghenti, Roena Sukhiashvili, Mariam Zakalashvili, Lela Urushadze, Magda Dgebuadze, Giorgi Tomashvili, Davit Tsaguria, Ekaterine Zangaladze, Nino Berishvili, Gvantsa Chanturia, Adam Kotorashvili, Maia Alkhazashvili, Irma Burjanadze, Anna Kasradze, Khatuna Zakhashvili, Paata Imnadze, Amiran Gamkrelidze. |
| 307 | EPI_ISL_415642 | hCoV-19/Georgia/Tb-477/2020 | *Homo sapiens* | Asia / Georgia / Tbilisi | 2020-03-10 | R. G. Lugar Center for Public Health Research, National Center for Disease Control and Public Health (NCDC) of Georgia. | R. G. Lugar Center for Public Health Research, National Center for Disease Control and Public Health (NCDC) of Georgia. | Nato Kotaria, Marine Murtskhvaladze, Ann Machablishvili, Lela Sabadze, Mari Gavashelidze, Ana Papkiauri, Meri Pantsulaia, Gvantsa Brachveli, Tata Imnadze, Tamar Jashiashvili, Tea Tevdoradze, Ketevan Sidamonidze, Ekaterine Khmaladze, Ekaterine Zhghenti, Roena Sukhiashvili, Mariam Zakalashvili, Lela Urushadze, Magda Dgebuadze, Giorgi Tomashvili, Davit Tsaguria, Ekaterine Zangaladze, Nino Berishvili, Gvantsa Chanturia, Adam Kotorashvili, Maia Alkhazashvili, Irma Burjanadze, Anna Kasradze, Khatuna Zakhashvili, Paata Imnadze, Amiran Gamkrelidze. |
| 308 | EPI_ISL_415643 | hCoV-19/Georgia/Tb-468/2020 | *Homo sapiens* | Asia / Georgia / Tbilisi | 2020-03-10 | R. G. Lugar Center for Public Health Research, National Center for Disease Control and Public Health (NCDC) of Georgia. | R. G. Lugar Center for Public Health Research, National Center for Disease Control and Public Health (NCDC) of Georgia. | Nato Kotaria, Marine Murtskhvaladze, Ann Machablishvili, Lela Sabadze, Mari Gavashelidze, Ana Papkiauri, Meri Pantsulaia, Gvantsa Brachveli, Tata Imnadze, Tamar Jashiashvili, Tea Tevdoradze, Ketevan Sidamonidze, Ekaterine Khmaladze, Ekaterine Zhghenti, Roena Sukhiashvili, Mariam Zakalashvili, Lela Urushadze, Magda Dgebuadze, Giorgi Tomashvili, Davit Tsaguria, Ekaterine Zangaladze, Nino Berishvili, Gvantsa Chanturia, Adam Kotorashvili, Maia Alkhazashvili, Irma Burjanadze, Anna Kasradze, Khatuna Zakhashvili, Paata Imnadze, Amiran Gamkrelidze. |
| 309 | EPI_ISL_415646 | hCoV-19/Denmark/SSI-101/2020 | *Homo sapiens* | Europe / Denmark | 2020-03-03 | Department of Virus and Microbiological Special diagnostics, Statens Serum Institut, Copenhagen, Denmark. | ViFU | Morten Rasmussen, Maiken Worsoe Rosenstierne , Anders Fomsgaard |
| 310 | EPI_ISL_415648 | hCoV-19/Denmark/SSI-104/2020 | *Homo sapiens* | Europe / Denmark | 2020-03-03 | Department of Virus and Microbiological Special diagnostics, Statens Serum Institut, Copenhagen, Denmark. | ViFU | Morten Rasmussen, Maiken Worsoe Rosenstierne , Anders Fomsgaard |
| 311 | EPI_ISL_415649 | hCoV-19/France/HF2039/2020 | *Homo sapiens* | Europe / France / Hauts-de-France / Crépy-en -Valois | 2020-03-05 | unknown | National Reference Center for Viruses of Respiratory Infections, Institut Pasteur, Paris | Mélnie Albert, Marion Barbet, Sylvie Behillil, Méline Bizard, Angela Brisebarre, Flora Donati Vincent Enouf, Maud Vanpeene, Sylvie van der Werf |
| 312 | EPI_ISL_415651 | hCoV-19/France/BFC2094/2020 | *Homo sapiens* | Europe / France / Bourgogne-France-Comté / Montreux-Chateau | 2020-03-05 | Unknown | National Reference Center for Viruses of Respiratory Infections, Institut Pasteur, Paris | Mélnie Albert, Marion Barbet, Sylvie Behillil, Méline Bizard, Angela Brisebarre, Flora Donati Vincent Enouf, Maud Vanpeene, Sylvie van der Werf |
| 313 | EPI_ISL_415652 | hCoV-19/France/BFC2147/2020 | *Homo sapiens* | Europe / France / Bourgogne-France-Comté / Thise | 2020-03-05 | unknown | National Reference Center for Viruses of Respiratory Infections, Institut Pasteur, Paris | Mélnie Albert, Marion Barbet, Sylvie Behillil, Méline Bizard, Angela Brisebarre, Flora Donati Vincent Enouf, Maud Vanpeene, Sylvie van der Werf |
| 314 | EPI_ISL_415698 | hCoV-19/Switzerland/GR2988/2020 | *Homo sapiens* | Europe / Switzerland | 2020-02-27 | Hôpitaux universitaires de Genève Laboratoire de Virologie | Hôpitaux universitaires de Genève Laboratoire de Virologie | Laubscher F. |

| | | | | | | | | |
|---|---|---|---|---|---|---|---|---|
| 314 | EPI_ISL_415699 | hCoV-19/Switzerland/GR3043/2020 | *Homo sapiens* | Europe / Switzerland | 2020-02-27 | Hôpitaux universitaires de Genève Laboratoire de Virologie | Hôpitaux universitaires de Genève Laboratoire de Virologie | Laubscher F. |
| 315 | EPI_ISL_415700 | hCoV-19/Switzerland/GE1402/2020 | *Homo sapiens* | Europe / Switzerland | 2020-02-28 | Hôpitaux universitaires de Genève Laboratoire de Virologie | Hôpitaux universitaires de Genève Laboratoire de Virologie | Laubscher F. |
| 316 | EPI_ISL_415702 | hCoV-19/Switzerland/SZ1417/2020 | *Homo sapiens* | Europe / Switzerland | 2020-03-02 | Hôpitaux universitaires de Genève Laboratoire de Virologie | Hôpitaux universitaires de Genève Laboratoire de Virologie | Laubscher F. |
| 317 | EPI_ISL_415703 | hCoV-19/Switzerland/TI2045/2020 | *Homo sapiens* | Europe / Switzerland | 2020-03-01 | Hôpitaux universitaires de Genève Laboratoire de Virologie | Hôpitaux universitaires de Genève Laboratoire de Virologie | Laubscher F. |
| 318 | EPI_ISL_415704 | hCoV-19/Switzerland/BE2536/2020 | *Homo sapiens* | Europe / Switzerland | 2020-03-04 | Hôpitaux universitaires de Genève Laboratoire de Virologie | Hôpitaux universitaires de Genève Laboratoire de Virologie | Laubscher F. |
| 318 | EPI_ISL_415705 | hCoV-19/Switzerland/GE4135/2020 | *Homo sapiens* | Europe / Switzerland | 2020-03-06 | Hôpitaux universitaires de Genève Laboratoire de Virologie | Hôpitaux universitaires de Genève Laboratoire de Virologie | Laubscher F. |
| 318 | EPI_ISL_415706 | hCoV-19/Switzerland/GE06207/2020 | *Homo sapiens* | Europe / Switzerland | 2020-03-06 | Hôpitaux universitaires de Genève Laboratoire de Virologie | Hôpitaux universitaires de Genève Laboratoire de Virologie | Laubscher F. |
| 318 | EPI_ISL_415707 | hCoV-19/Switzerland/GE6679/2020 | *Homo sapiens* | Europe / Switzerland | 2020-03-08 | Hôpitaux universitaires de Genève Laboratoire de Virologie | Hôpitaux universitaires de Genève Laboratoire de Virologie | Laubscher F. |
| 318 | EPI_ISL_416493 | hCoV-19/France/HF2196/2020 | *Homo sapiens* | Europe / France / Hauts de France / Château-Thierry | 2020-03-08 | CH Jean de Navarre Laboratoire de Biologie | National Reference Center for Viruses of Respiratory Infections, Institut Pasteur, Paris | Mélnie Albert, Marion Barbet, Sylvie Behillil, Méline Bizard, Angela Brisebarre, Flora Donati, Etienne Simon-Lorière, Vincent Enouf, Maud Vanpeene, Sylvie van der Werf |
| 319 | EPI_ISL_415708 | hCoV-19/Switzerland/GE4984/2020 | *Homo sapiens* | Europe / Switzerland | 2020-03-07 | Hôpitaux universitaires de Genève Laboratoire de Virologie | Hôpitaux universitaires de Genève Laboratoire de Virologie | Laubscher F. |
| 320 | EPI_ISL_415709 | hCoV-19/Hangzhou/ZJU-01/2020 | *Homo sapiens* | Asia / China / Hangzhou | 2020-01-25 | State Key Laboratory for Diagnosis and Treatment of Infectious Diseases, National Clinical Research Center for Infectious Diseases, First Affiliated Hospital, Zhejiang University School of Medicine, Hangzhou, China. 310003 | State Key Laboratory for Diagnosis and Treatment of Infectious Diseases, National Clinical Research Center for Infectious Diseases, First Affiliated Hospital, Zhejiang University School of Medicine, Hangzhou, China. 310003 | Hangping Yao, Nanping Wu, Chao Jiang, Xiangyun Lu, Linfang Cheng, Fumin Liu, Zhigang Wu, Haibo Wu, Changzhong Jin, Min Zheng, Lanjuan Li |
| 321 | EPI_ISL_415741 | hCoV-19/Taiwan/CGMH-CGU-03/2020 | *Homo sapiens* | Asia / Taiwan / Taoyuan | 2020-02-26 | Laboratory Medicine | Department of Laboratory Medicine, Lin-Kou Chang Gung Memorial Hospital, Taoyuan, Taiwan | Kuo-Chien Tsao, Yu-Nong Gong, Shu-Li Yang, Yi-Chun Liu, Chung-Guei Huang, Po-Wei Huang, Mei-Jen Hsiao, Cheng-Ta Yang, Cheng-Hsun Chiu, Chi-Hsien Huang, Kuang-Tso Le, Shu-Min Lin, Peng-Nien Huang, Kuo-Ming Lee, Guang-Wu Chen, Shin-Ru Shih |
| 321 | EPI_ISL_415743 | hCoV-19/Taiwan/CGMH-CGU-05/2020 | *Homo sapiens* | Asia / Taiwan / Taoyuan | 2020-02-27 | Laboratory Medicine | Department of Laboratory Medicine, Lin-Kou Chang Gung Memorial Hospital, Taoyuan, Taiwan | Kuo-Chien Tsao, Yu-Nong Gong, Shu-Li Yang, Yi-Chun Liu, Chung-Guei Huang, Po-Wei Huang, Mei-Jen Hsiao, Cheng-Ta Yang, Cheng-Hsun Chiu, Chi-Hsien Huang, Kuang-Tso Le, Shu-Min Lin, Peng-Nien Huang, Kuo-Ming Lee, Guang-Wu Chen, Shin-Ru Shih |
| 322 | EPI_ISL_415787 | hCoV-19/Peru/010/2020 | *Homo sapiens* | South America / Peru / Lima | 2020-03-10 | Laboratorio de Referencia Nacional de Virus Respiratorio. Instituto Nacional de Salud. Peru | Laboratorio de Referencia Nacional de Biotecnologia y Biologia Molecular.Instituto Nacional de Salud.Peru | Carlos Padilla Rojas, Priscila Lope Pari, Karolyn Vega Chozo, Johanna Balbuena Torres, Omar Caceres Rey, Hemri Bailon Calderon, Maribel Huaringa Nuñez, Nancy Rojas Serrano |
| 323 | EPI_ISL_415920 | hCoV-19/Wales/PHW32/2020 | *Homo sapiens* | Europe / United Kingdom / Wales | 2020-03-12 | Wales Specialist Virology Centre | Public Health Wales Microbiology Cardiff | Catherine Moore, Joanne Watkins, Sally Corden, Tom Connor |
| 324 | EPI_ISL_416024 | hCoV-19/Wales/PHW35/2020 | *Homo sapiens* | Europe / United Kingdom / Wales | 2020-03-11 | Wales Specialist Virology Centre | Public Health Wales Microbiology Cardiff | Catherine Moore, Joanne Watkins, Sally Corden, Tom Connor |

| | | | | | | | | |
|---|---|---|---|---|---|---|---|---|
| 325 | EPI_ISL_416029 | hCoV-19/Brazil/SPBR-08/2020 | *Homo sapiens* | South America / Brazil / Sao Paulo / Sao Paulo | 2020-03-04 | Laboiratório Fleury | Instituto Adolfo Lutz, Interdiciplinary Procedures Center, Strategic Laboratory | Claudio Tavares Sacchi, Claudia Regina Gonçalves, Carlos Henrique Camargo, Fabiana Cristina Pereira dos Santos, Daniela Bernardes Borges da Silva, Simone Guadagnucci Morillo, Adriano Abbud, Adriana Bugno, Maria do Carmo Sampaio Tavares Timenetsky, Terezinha Maria de Paiva |
| 326 | EPI_ISL_416032 | hCoV-19/Brazil/SPBR-10/2020 | *Homo sapiens* | South America / Brazil / Distrito Federal / Brasilia | 2020-03-04 | National Influenza Center - Instituto Adolfo Lutz | Instituto Adolfo Lutz, Interdiciplinary Procedures Center, Strategic Laboratory | Claudio Tavares Sacchi, Claudia Regina Gonçalves, Carlos Henrique Camargo, Fabiana Cristina Pereira dos Santos, Daniela Bernardes Borges da Silva, Simone Guadagnucci Morillo, Adriano Abbud, Adriana Bugno, Maria do Carmo Sampaio Tavares Timenetsky, Terezinha Maria de Paiva |
| 327 | EPI_ISL_416033 | hCoV-19/Brazil/SPBR-11/2020 | *Homo sapiens* | South America / Brazil / Sao Paulo / Sao Paulo | 2020-03-03 | Hospital Israelita Albert Einstein | Instituto Adolfo Lutz, Interdiciplinary Procedures Center, Strategic Laboratory | Claudio Tavares Sacchi, Claudia Regina Gonçalves, Carlos Henrique Camargo, Erica Valessa Ramos Gomes, Fabiana Cristina Pereira dos Santos, Daniela Bernardes Borges da Silva, Simone Guadagnucci Morillo, Adriano Abbud, Adriana Bugno, Maria do Carmo Sampaio Tavares Timenetsky, Terezinha Maria de Paiva |
| 328 | EPI_ISL_416034 | hCoV-19/Brazil/SPBR-12/2020 | *Homo sapiens* | South America / Brazil / Sao Paulo / Sao Paulo | 2020-03-04 | Hospital Israelita Albert Einstein | Instituto Adolfo Lutz, Interdiciplinary Procedures Center, Strategic Laboratory | Claudio Tavares Sacchi, Claudia Regina Gonçalves, Carlos Henrique Camargo, Erica Valessa Ramos Gomes, Fabiana Cristina Pereira dos Santos, Daniela Bernardes Borges da Silva, Simone Guadagnucci Morillo, Adriano Abbud, Adriana Bugno, Maria do Carmo Sampaio Tavares Timenetsky, Terezinha Maria de Paiva |
| 329 | EPI_ISL_416036 | hCoV-19/Brazil/SPBR-14/2020 | *Homo sapiens* | South America / Brazil / Sao Paulo / Sao Paulo | 2020-03-05 | National Influenza Center - Instituto Adolfo Lutz | Instituto Adolfo Lutz, Interdiciplinary Procedures Center, Strategic Laboratory | Claudio Tavares Sacchi, Claudia Regina Gonçalves, Carlos Henrique Camargo, Erica Valessa Ramos Gomes, Fabiana Cristina Pereira dos Santos, Daniela Bernardes Borges da Silva, Simone Guadagnucci Morillo, Adriano Abbud, Adriana Bugno, Maria do Carmo Sampaio Tavares Timenetsky, Terezinha Maria de Paiva |
| 330 | EPI_ISL_416044 | hCoV-19/Hangzhou/ZJU-03/2020 | *Homo sapiens* | Asia / China / Hangzhou | 2020-01-25 | State Key Laboratory for Diagnosis and Treatment of Infectious Diseases, National Clinical Research Center for Infectious Diseases, First Affiliated Hospital, Zhejiang University School of Medicine, Hangzhou, China 310003 | State Key Laboratory for Diagnosis and Treatment of Infectious Diseases, National Clinical Research Center for Infectious Diseases, First Affiliated Hospital, Zhejiang University School of Medicine, Hangzhou, China 310003 | Hangping Yao, Nanping Wu, Chao Jiang, Xiangyun Lu, Linfang Cheng, Fumin Liu, Zhigang Wu, Haibo Wu, Changzhong Jin, Min Zheng, Lanjuan Li |
| 331 | EPI_ISL_416047 | hCoV-19/Hangzhou/ZJU-06/2020 | *Homo sapiens* | Asia / China / Hangzhou | 2020-02-02 | State Key Laboratory for Diagnosis and Treatment of Infectious Diseases, National Clinical Research Center for Infectious Diseases, First Affiliated Hospital, Zhejiang University School of Medicine, Hangzhou, China 310003 | State Key Laboratory for Diagnosis and Treatment of Infectious Diseases, National Clinical Research Center for Infectious Diseases, First Affiliated Hospital, Zhejiang University School of Medicine, Hangzhou, China 310003 | Hangping Yao, Nanping Wu, Chao Jiang, Xiangyun Lu, Linfang Cheng, Fumin Liu, Zhigang Wu, Haibo Wu, Changzhong Jin, Min Zheng, Lanjuan Li |
| 332 | EPI_ISL_416140 | hCoV-19/Denmark/SSI-05/2020 | *Homo sapiens* | Europe / Denmark / Copenhagen | 2020-03-02 | Department of Virus and Microbiological Special diagnostics, Statens Serum Institut, Copenhagen, Denmark. | Statens Serum Institute | Morten Rasmussen, Maiken Worsoe Rosenstierne , Anders Fomsgaard |
| 333 | EPI_ISL_416141 | hCoV-19/Denmark/SSI-09/2020 | *Homo sapiens* | Europe / Denmark / Copenhagen | 2020-03-03 | Department of Virus and Microbiological Special diagnostics, Statens Serum Institut, Copenhagen, Denmark. | Statens Serum Institute | Morten Rasmussen, Maiken Worsoe Rosenstierne , Anders Fomsgaard |

| | | | | | | | | |
|---|---|---|---|---|---|---|---|---|
| 334 | EPI_ISL_416142 | hCoV-19/Denmark/SSI-01/2020 | *Homo sapiens* | Europe / Denmark / Copenhagen | 2020-02-26 | Department of Virus and Microbiological Special diagnostics, Statens Serum Institut, Copenhagen, Denmark. | Statens Serum Institute | Morten Rasmussen, Maiken Worsoe Rosenstierne , Anders Fomsgaard |
| 335 | EPI_ISL_416143 | hCoV-19/Denmark/SSI-02/2020 | *Homo sapiens* | Europe / Denmark / Copenhagen | 2020-02-28 | Department of Virus and Microbiological Special diagnostics, Statens Serum Institut, Copenhagen, Denmark. | ViFU | Morten Rasmussen, Maiken Worsoe Rosenstierne , Anders Fomsgaard |
| 336 | EPI_ISL_416144 | hCoV-19/Denmark/SSI-03/2020 | *Homo sapiens* | Europe / Denmark / Copenhagen | 2020-03-01 | Department of Virus and Microbiological Special diagnostics, Statens Serum Institut, Copenhagen, Denmark. | ViFU | Morten Rasmussen, Maiken Worsoe Rosenstierne , Anders Fomsgaard |
| 337 | EPI_ISL_416153 | hCoV-19/Denmark/SSI-04/2020 | *Homo sapiens* | Europe / Denmark / Copenhagen | 2020-03-02 | Department of Virus and Microbiological Special diagnostics, Statens Serum Institut, Copenhagen, Denmark. | ViFU | Morten Rasmussen, Maiken Worsoe Rosenstierne , Anders Fomsgaard |
| 338 | EPI_ISL_416316 | hCoV-19/Shanghai/SH0002/2020 | *Homo sapiens* | Asia / China / Shanghai | 2020-01-25 | Shanghai Public Health Clinical Center, Shanghai Medical College, Fudan University | National Research Center for Translational Medicine (Shanghai), Ruijin Hospital affiliated to Shanghai Jiao Tong University School of Medicine & Shanghai Public Health Clinical Center | Shengyue Wang, Xiaonan Zhang, Gang Lu, Yun Tan, Yun Ling, Hongzhou Lu, Saijuan Chen |
| 339 | EPI_ISL_416317 | hCoV-19/Shanghai/SH0003/2020 | *Homo sapiens* | Asia / China / Shanghai | 2020-01-25 | Shanghai Public Health Clinical Center, Shanghai Medical College, Fudan University | National Research Center for Translational Medicine (Shanghai), Ruijin Hospital affiliated to Shanghai Jiao Tong University School of Medicine & Shanghai Public Health Clinical Center | Shengyue Wang, Xiaonan Zhang, Gang Lu, Yun Tan, Yun Ling, Hongzhou Lu, Saijuan Chen |
| 340 | EPI_ISL_416318 | hCoV-19/Shanghai/SH0004/2020 | *Homo sapiens* | Asia / China / Shanghai | 2020-01-28 | Shanghai Public Health Clinical Center, Shanghai Medical College, Fudan University | National Research Center for Translational Medicine (Shanghai), Ruijin Hospital affiliated to Shanghai Jiao Tong University School of Medicine & Shanghai Public Health Clinical Center | Shengyue Wang, Xiaonan Zhang, Gang Lu, Yun Tan, Yun Ling, Hongzhou Lu, Saijuan Chen |
| 341 | EPI_ISL_416320 | hCoV-19/Shanghai/SH0007/2020 | *Homo sapiens* | Asia / China / Shanghai | 2020-01-28 | Shanghai Public Health Clinical Center, Shanghai Medical College, Fudan University | National Research Center for Translational Medicine (Shanghai), Ruijin Hospital affiliated to Shanghai Jiao Tong University School of Medicine & Shanghai Public Health Clinical Center | Shengyue Wang, Xiaonan Zhang, Gang Lu, Yun Tan, Yun Ling, Hongzhou Lu, Saijuan Chen |
| 342 | EPI_ISL_416321 | hCoV-19/Shanghai/SH0008/2020 | *Homo sapiens* | Asia / China / Shanghai | 2020-01-28 | Shanghai Public Health Clinical Center, Shanghai Medical College, Fudan University | National Research Center for Translational Medicine (Shanghai), Ruijin Hospital affiliated to Shanghai Jiao Tong University School of Medicine & Shanghai Public Health Clinical Center | Shengyue Wang, Xiaonan Zhang, Gang Lu, Yun Tan, Yun Ling, Hongzhou Lu, Saijuan Chen |
| 343 | EPI_ISL_416322 | hCoV-19/Shanghai/SH0009/2020 | *Homo sapiens* | Asia / China / Shanghai | 2020-01-29 | Shanghai Public Health Clinical Center, Shanghai Medical College, Fudan University | National Research Center for Translational Medicine (Shanghai), Ruijin Hospital affiliated to Shanghai Jiao Tong University School of Medicine & Shanghai Public Health Clinical Center | Shengyue Wang, Xiaonan Zhang, Gang Lu, Yun Tan, Yun Ling, Hongzhou Lu, Saijuan Chen |
| 344 | EPI_ISL_416323 | hCoV-19/Shanghai/SH0010/2020 | *Homo sapiens* | Asia / China / Shanghai | 2020-01-29 | Shanghai Public Health Clinical Center, Shanghai Medical College, Fudan University | National Research Center for Translational Medicine (Shanghai), Ruijin Hospital affiliated to Shanghai Jiao Tong University School of Medicine & Shanghai Public Health Clinical Center | Shengyue Wang, Xiaonan Zhang, Gang Lu, Yun Tan, Yun Ling, Hongzhou Lu, Saijuan Chen |

| | | | | | | | | |
|---|---|---|---|---|---|---|---|---|
| 345 | EPI_ISL_416325 | hCoV-19/Shanghai/SH0012/2020 | *Homo sapiens* | Asia / China / Shanghai | 2020-02-02 | Shanghai Public Health Clinical Center, Shanghai Medical College, Fudan University | National Research Center for Translational Medicine (Shanghai), Ruijin Hospital affiliated to Shanghai Jiao Tong University School of Medicine & Shanghai Public Health Clinical Center | Shengyue Wang, Xiaonan Zhang, Gang Lu, Yun Tan, Yun Ling, Hongzhou Lu, Saijuan Chen |
| 346 | EPI_ISL_416326 | hCoV-19/Shanghai/SH0013/2020 | *Homo sapiens* | Asia / China / Shanghai | 2020-01-30 | Shanghai Public Health Clinical Center, Shanghai Medical College, Fudan University | National Research Center for Translational Medicine (Shanghai), Ruijin Hospital affiliated to Shanghai Jiao Tong University School of Medicine & Shanghai Public Health Clinical Center | Shengyue Wang, Xiaonan Zhang, Gang Lu, Yun Tan, Yun Ling, Hongzhou Lu, Saijuan Chen |
| 347 | EPI_ISL_416330 | hCoV-19/Shanghai/SH0021/2020 | *Homo sapiens* | Asia / China / Shanghai | 2020-02-02 | Shanghai Public Health Clinical Center, Shanghai Medical College, Fudan University | National Research Center for Translational Medicine (Shanghai), Ruijin Hospital affiliated to Shanghai Jiao Tong University School of Medicine & Shanghai Public Health Clinical Center | Shengyue Wang, Xiaonan Zhang, Gang Lu, Yun Tan, Yun Ling, Hongzhou Lu, Saijuan Chen |
| 348 | EPI_ISL_416331 | hCoV-19/Shanghai/SH0022/2020 | *Homo sapiens* | Asia / China / Shanghai | 2020-01-30 | Shanghai Public Health Clinical Center, Shanghai Medical College, Fudan University | National Research Center for Translational Medicine (Shanghai), Ruijin Hospital affiliated to Shanghai Jiao Tong University School of Medicine & Shanghai Public Health Clinical Center | Shengyue Wang, Xiaonan Zhang, Gang Lu, Yun Tan, Yun Ling, Hongzhou Lu, Saijuan Chen |
| 348 | EPI_ISL_416332 | hCoV-19/Shanghai/SH0023/2020 | *Homo sapiens* | Asia / China / Shanghai | 2020-01-30 | Shanghai Public Health Clinical Center, Shanghai Medical College, Fudan University | National Research Center for Translational Medicine (Shanghai), Ruijin Hospital affiliated to Shanghai Jiao Tong University School of Medicine & Shanghai Public Health Clinical Center | Shengyue Wang, Xiaonan Zhang, Gang Lu, Yun Tan, Yun Ling, Hongzhou Lu, Saijuan Chen |
| 349 | EPI_ISL_416335 | hCoV-19/Shanghai/SH0026/2020 | *Homo sapiens* | Asia / China / Shanghai | 2020-02-02 | Shanghai Public Health Clinical Center, Shanghai Medical College, Fudan University | National Research Center for Translational Medicine (Shanghai), Ruijin Hospital affiliated to Shanghai Jiao Tong University School of Medicine & Shanghai Public Health Clinical Center | Shengyue Wang, Xiaonan Zhang, Gang Lu, Yun Tan, Yun Ling, Hongzhou Lu, Saijuan Chen |
| 350 | EPI_ISL_416336 | hCoV-19/Shanghai/SH0027/2020 | *Homo sapiens* | Asia / China / Shanghai | 2020-02-01 | Shanghai Public Health Clinical Center, Shanghai Medical College, Fudan University | National Research Center for Translational Medicine (Shanghai), Ruijin Hospital affiliated to Shanghai Jiao Tong University School of Medicine & Shanghai Public Health Clinical Center | Shengyue Wang, Xiaonan Zhang, Gang Lu, Yun Tan, Yun Ling, Hongzhou Lu, Saijuan Chen |
| 351 | EPI_ISL_416338 | hCoV-19/Shanghai/SH0029/2020 | *Homo sapiens* | Asia / China / Shanghai | 2020-02-01 | Shanghai Public Health Clinical Center, Shanghai Medical College, Fudan University | National Research Center for Translational Medicine (Shanghai), Ruijin Hospital affiliated to Shanghai Jiao Tong University School of Medicine & Shanghai Public Health Clinical Center | Shengyue Wang, Xiaonan Zhang, Gang Lu, Yun Tan, Yun Ling, Hongzhou Lu, Saijuan Chen |
| 351 | EPI_ISL_416381 | hCoV-19/Shanghai/SH0079/2020 | *Homo sapiens* | Asia / China / Shanghai | 2020-02-01 | Shanghai Public Health Clinical Center, Shanghai Medical College, Fudan University | National Research Center for Translational Medicine (Shanghai), Ruijin Hospital affiliated to Shanghai Jiao Tong University School of Medicine & Shanghai Public Health Clinical Center | Shengyue Wang, Xiaonan Zhang, Gang Lu, Yun Tan, Yun Ling, Hongzhou Lu, Saijuan Chen |
| 352 | EPI_ISL_416339 | hCoV-19/Shanghai/SH0030/2020 | *Homo sapiens* | Asia / China / Shanghai | 2020-02-03 | Shanghai Public Health Clinical Center, Shanghai Medical College, Fudan University | National Research Center for Translational Medicine (Shanghai), Ruijin Hospital affiliated to Shanghai Jiao Tong University School of Medicine & Shanghai Public Health Clinical Center | Shengyue Wang, Xiaonan Zhang, Gang Lu, Yun Tan, Yun Ling, Hongzhou Lu, Saijuan Chen |
| 353 | EPI_ISL_416341 | hCoV-19/Shanghai/SH0032/2020 | *Homo sapiens* | Asia / China / Shanghai | 2020-02-01 | Shanghai Public Health Clinical Center, Shanghai Medical College, Fudan University | National Research Center for Translational Medicine (Shanghai), Ruijin Hospital affiliated to Shanghai Jiao Tong University School of Medicine & Shanghai Public Health Clinical Center | Shengyue Wang, Xiaonan Zhang, Gang Lu, Yun Tan, Yun Ling, Hongzhou Lu, Saijuan Chen |

| | | | | | | | | |
|---|---|---|---|---|---|---|---|---|
| 354 | EPI_ISL_416353 | hCoV-19/Shanghai/SH0044/2020 | *Homo sapiens* | Asia / China / Shanghai | 2020-02-04 | Shanghai Public Health Clinical Center, Shanghai Medical College, Fudan University | National Research Center for Translational Medicine (Shanghai), Ruijin Hospital affiliated to Shanghai Jiao Tong University School of Medicine & Shanghai Public Health Clinical Center | Shengyue Wang, Xiaonan Zhang, Gang Lu, Yun Tan, Yun Ling, Hongzhou Lu, Saijuan Chen |
| 354 | EPI_ISL_416380 | hCoV-19/Shanghai/SH0077/2020 | *Homo sapiens* | Asia / China / Shanghai | 2020-02-09 | Shanghai Public Health Clinical Center, Shanghai Medical College, Fudan University | National Research Center for Translational Medicine (Shanghai), Ruijin Hospital affiliated to Shanghai Jiao Tong University School of Medicine & Shanghai Public Health Clinical Center | Shengyue Wang, Xiaonan Zhang, Gang Lu, Yun Tan, Yun Ling, Hongzhou Lu, Saijuan Chen |
| 354 | EPI_ISL_416393 | hCoV-19/Shanghai/SH0101/2020 | *Homo sapiens* | Asia / China / Shanghai | 2020-02-09 | Shanghai Public Health Clinical Center, Shanghai Medical College, Fudan University | National Research Center for Translational Medicine (Shanghai), Ruijin Hospital affiliated to Shanghai Jiao Tong University School of Medicine & Shanghai Public Health Clinical Center | Shengyue Wang, Xiaonan Zhang, Gang Lu, Yun Tan, Yun Ling, Hongzhou Lu, Saijuan Chen |
| 354 | EPI_ISL_416394 | hCoV-19/Shanghai/SH0102/2020 | *Homo sapiens* | Asia / China / Shanghai | 2020-02-14 | Shanghai Public Health Clinical Center, Shanghai Medical College, Fudan University | National Research Center for Translational Medicine (Shanghai), Ruijin Hospital affiliated to Shanghai Jiao Tong University School of Medicine & Shanghai Public Health Clinical Center | Shengyue Wang, Xiaonan Zhang, Gang Lu, Yun Tan, Yun Ling, Hongzhou Lu, Saijuan Chen |
| 355 | EPI_ISL_416361 | hCoV-19/Shanghai/SH0053/2020 | *Homo sapiens* | Asia / China / Shanghai | 2020-02-01 | Shanghai Public Health Clinical Center, Shanghai Medical College, Fudan University | National Research Center for Translational Medicine (Shanghai), Ruijin Hospital affiliated to Shanghai Jiao Tong University School of Medicine & Shanghai Public Health Clinical Center | Shengyue Wang, Xiaonan Zhang, Gang Lu, Yun Tan, Yun Ling, Hongzhou Lu, Saijuan Chen |
| 356 | EPI_ISL_416352 | hCoV-19/Shanghai/SH0043/2020 | *Homo sapiens* | Asia / China / Shanghai | 2020-01-30 | Shanghai Public Health Clinical Center, Shanghai Medical College, Fudan University | National Research Center for Translational Medicine (Shanghai), Ruijin Hospital affiliated to Shanghai Jiao Tong University School of Medicine & Shanghai Public Health Clinical Center | Shengyue Wang, Xiaonan Zhang, Gang Lu, Yun Tan, Yun Ling, Hongzhou Lu, Saijuan Chen |
| 356 | EPI_ISL_416362 | hCoV-19/Shanghai/SH0054/2020 | *Homo sapiens* | Asia / China / Shanghai | 2020-02-02 | Shanghai Public Health Clinical Center, Shanghai Medical College, Fudan University | National Research Center for Translational Medicine (Shanghai), Ruijin Hospital affiliated to Shanghai Jiao Tong University School of Medicine & Shanghai Public Health Clinical Center | Shengyue Wang, Xiaonan Zhang, Gang Lu, Yun Tan, Yun Ling, Hongzhou Lu, Saijuan Chen |
| 357 | EPI_ISL_416363 | hCoV-19/Shanghai/SH0055/2020 | *Homo sapiens* | Asia / China / Shanghai | 2020-02-08 | Shanghai Public Health Clinical Center, Shanghai Medical College, Fudan University | National Research Center for Translational Medicine (Shanghai), Ruijin Hospital affiliated to Shanghai Jiao Tong University School of Medicine & Shanghai Public Health Clinical Center | Shengyue Wang, Xiaonan Zhang, Gang Lu, Yun Tan, Yun Ling, Hongzhou Lu, Saijuan Chen |
| 358 | EPI_ISL_416366 | hCoV-19/Shanghai/SH0059/2020 | *Homo sapiens* | Asia / China / Shanghai | 2020-01-30 | Shanghai Public Health Clinical Center, Shanghai Medical College, Fudan University | National Research Center for Translational Medicine (Shanghai), Ruijin Hospital affiliated to Shanghai Jiao Tong University School of Medicine & Shanghai Public Health Clinical Center | Shengyue Wang, Xiaonan Zhang, Gang Lu, Yun Tan, Yun Ling, Hongzhou Lu, Saijuan Chen |
| 359 | EPI_ISL_416367 | hCoV-19/Shanghai/SH0060/2020 | *Homo sapiens* | Asia / China / Shanghai | 2020-02-01 | Shanghai Public Health Clinical Center, Shanghai Medical College, Fudan University | National Research Center for Translational Medicine (Shanghai), Ruijin Hospital affiliated to Shanghai Jiao Tong University School of Medicine & Shanghai Public Health Clinical Center | Shengyue Wang, Xiaonan Zhang, Gang Lu, Yun Tan, Yun Ling, Hongzhou Lu, Saijuan Chen |
| 360 | EPI_ISL_416373 | hCoV-19/Shanghai/SH0070/2020 | *Homo sapiens* | Asia / China / Shanghai | 2020-02-07 | Shanghai Public Health Clinical Center, Shanghai Medical College, Fudan University | National Research Center for Translational Medicine (Shanghai), Ruijin Hospital affiliated to Shanghai Jiao Tong University School of Medicine & Shanghai Public Health Clinical Center | Shengyue Wang, Xiaonan Zhang, Gang Lu, Yun Tan, Yun Ling, Hongzhou Lu, Saijuan Chen |

| | | | | | | | | |
|---|---|---|---|---|---|---|---|---|
| 361 | EPI_ISL_416376 | hCoV-19/Shanghai/SH0073/2020 | *Homo sapiens* | Asia / China / Shanghai | 2020-02-05 | Shanghai Public Health Clinical Center, Shanghai Medical College, Fudan University | National Research Center for Translational Medicine (Shanghai), Ruijin Hospital affiliated to Shanghai Jiao Tong University School of Medicine & Shanghai Public Health Clinical Center | Shengyue Wang, Xiaonan Zhang, Gang Lu, Yun Tan, Yun Ling, Hongzhou Lu, Saijuan Chen |
| 362 | EPI_ISL_416377 | hCoV-19/Shanghai/SH0074/2020 | *Homo sapiens* | Asia / China / Shanghai | 2020-02-07 | Shanghai Public Health Clinical Center, Shanghai Medical College, Fudan University | National Research Center for Translational Medicine (Shanghai), Ruijin Hospital affiliated to Shanghai Jiao Tong University School of Medicine & Shanghai Public Health Clinical Center | Shengyue Wang, Xiaonan Zhang, Gang Lu, Yun Tan, Yun Ling, Hongzhou Lu, Saijuan Chen |
| 363 | EPI_ISL_416382 | hCoV-19/Shanghai/SH0080/2020 | *Homo sapiens* | Asia / China / Shanghai | 2020-02-08 | Shanghai Public Health Clinical Center, Shanghai Medical College, Fudan University | National Research Center for Translational Medicine (Shanghai), Ruijin Hospital affiliated to Shanghai Jiao Tong University School of Medicine & Shanghai Public Health Clinical Center | Shengyue Wang, Xiaonan Zhang, Gang Lu, Yun Tan, Yun Ling, Hongzhou Lu, Saijuan Chen |
| 364 | EPI_ISL_416390 | hCoV-19/Shanghai/SH0094/2020 | *Homo sapiens* | Asia / China / Shanghai | 2020-01-31 | Shanghai Public Health Clinical Center, Shanghai Medical College, Fudan University | National Research Center for Translational Medicine (Shanghai), Ruijin Hospital affiliated to Shanghai Jiao Tong University School of Medicine & Shanghai Public Health Clinical Center | Shengyue Wang, Xiaonan Zhang, Gang Lu, Yun Tan, Yun Ling, Hongzhou Lu, Saijuan Chen |
| 365 | EPI_ISL_416397 | hCoV-19/Shanghai/SH0107/2020 | *Homo sapiens* | Asia / China / Shanghai | 2020-02-02 | Shanghai Public Health Clinical Center, Shanghai Medical College, Fudan University | National Research Center for Translational Medicine (Shanghai), Ruijin Hospital affiliated to Shanghai Jiao Tong University School of Medicine & Shanghai Public Health Clinical Center | Shengyue Wang, Xiaonan Zhang, Gang Lu, Yun Tan, Yun Ling, Hongzhou Lu, Saijuan Chen |
| 366 | EPI_ISL_416398 | hCoV-19/Shanghai/SH0109/2020 | *Homo sapiens* | Asia / China / Shanghai | 2020-02-02 | Shanghai Public Health Clinical Center, Shanghai Medical College, Fudan University | National Research Center for Translational Medicine (Shanghai), Ruijin Hospital affiliated to Shanghai Jiao Tong University School of Medicine & Shanghai Public Health Clinical Center | Shengyue Wang, Xiaonan Zhang, Gang Lu, Yun Tan, Yun Ling, Hongzhou Lu, Saijuan Chen |
| 367 | EPI_ISL_416399 | hCoV-19/Shanghai/SH0110/2020 | *Homo sapiens* | Asia / China / Shanghai | 2020-02-13 | Shanghai Public Health Clinical Center, Shanghai Medical College, Fudan University | National Research Center for Translational Medicine (Shanghai), Ruijin Hospital affiliated to Shanghai Jiao Tong University School of Medicine & Shanghai Public Health Clinical Center | Shengyue Wang, Xiaonan Zhang, Gang Lu, Yun Tan, Yun Ling, Hongzhou Lu, Saijuan Chen |
| 368 | EPI_ISL_416401 | hCoV-19/Shanghai/SH0114/2020 | *Homo sapiens* | Asia / China / Shanghai | 2020-02-02 | Shanghai Public Health Clinical Center, Shanghai Medical College, Fudan University | National Research Center for Translational Medicine (Shanghai), Ruijin Hospital affiliated to Shanghai Jiao Tong University School of Medicine & Shanghai Public Health Clinical Center | Shengyue Wang, Xiaonan Zhang, Gang Lu, Yun Tan, Yun Ling, Hongzhou Lu, Saijuan Chen |
| 369 | EPI_ISL_416402 | hCoV-19/Shanghai/SH0115/2020 | *Homo sapiens* | Asia / China / Shanghai | 2020-02-11 | Shanghai Public Health Clinical Center, Shanghai Medical College, Fudan University | National Research Center for Translational Medicine (Shanghai), Ruijin Hospital affiliated to Shanghai Jiao Tong University School of Medicine & Shanghai Public Health Clinical Center | Shengyue Wang, Xiaonan Zhang, Gang Lu, Yun Tan, Yun Ling, Hongzhou Lu, Saijuan Chen |
| 370 | EPI_ISL_416403 | hCoV-19/Shanghai/SH0117/2020 | *Homo sapiens* | Asia / China / Shanghai | 2020-02-02 | Shanghai Public Health Clinical Center, Shanghai Medical College, Fudan University | National Research Center for Translational Medicine (Shanghai), Ruijin Hospital affiliated to Shanghai Jiao Tong University School of Medicine & Shanghai Public Health Clinical Center | Shengyue Wang, Xiaonan Zhang, Gang Lu, Yun Tan, Yun Ling, Hongzhou Lu, Saijuan Chen |
| 371 | EPI_ISL_416405 | hCoV-19/Shanghai/SH0121/2020 | *Homo sapiens* | Asia / China / Shanghai | 2020-02-02 | Shanghai Public Health Clinical Center, Shanghai Medical College, Fudan University | National Research Center for Translational Medicine (Shanghai), Ruijin Hospital affiliated to Shanghai Jiao Tong University School of Medicine & Shanghai Public Health Clinical Center | Shengyue Wang, Xiaonan Zhang, Gang Lu, Yun Tan, Yun Ling, Hongzhou Lu, Saijuan Chen |

| | | | | | | | | |
|---|---|---|---|---|---|---|---|---|
| 372 | EPI_ISL_416407 | hCoV-19/Shanghai/SH0126/2020 | *Homo sapiens* | Asia / China / Shanghai | 2020-02-15 | Shanghai Public Health Clinical Center, Shanghai Medical College, Fudan University | National Research Center for Translational Medicine (Shanghai), Ruijin Hospital affiliated to Shanghai Jiao Tong University School of Medicine & Shanghai Public Health Clinical Center | Shengyue Wang, Xiaonan Zhang, Gang Lu, Yun Tan, Yun Ling, Hongzhou Lu, Saijuan Chen |
| 373 | EPI_ISL_416409 | hCoV-19/Shanghai/SH0128/2020 | *Homo sapiens* | Asia / China / Shanghai | 2020-02-02 | Shanghai Public Health Clinical Center, Shanghai Medical College, Fudan University | National Research Center for Translational Medicine (Shanghai), Ruijin Hospital affiliated to Shanghai Jiao Tong University School of Medicine & Shanghai Public Health Clinical Center | Shengyue Wang, Xiaonan Zhang, Gang Lu, Yun Tan, Yun Ling, Hongzhou Lu, Saijuan Chen |
| 374 | EPI_ISL_416411 | hCoV-19/Australia/VIC03/2020 | *Homo sapiens* | Oceania / Australia / Victoria / Melbourne | 2020-01-25 | Victorian Infectious Diseases Reference Laboratory (VIDRL) | Victorian Infectious Diseases Reference Laboratory and Microbiological Diagnostic Unit Public Health Laboratory, Doherty Institute | Caly L., Seemann T., Schultz M., Druce J., Taiaroa, G. |
| 375 | EPI_ISL_416412 | hCoV-19/Australia/VIC04/2020 | *Homo sapiens* | Oceania / Australia / Victoria / Melbourne | 2020-03-02 | Victorian Infectious Diseases Reference Laboratory (VIDRL) | Victorian Infectious Diseases Reference Laboratory and Microbiological Diagnostic Unit Public Health Laboratory, Doherty Institute | Caly L., Seemann T., Schultz M., Druce J., Taiaroa, G. |
| 376 | EPI_ISL_416415 | hCoV-19/Australia/VIC07/2020 | *Homo sapiens* | Oceania / Australia / Victoria / Melbourne | 2020-02-08 | Victorian Infectious Diseases Reference Laboratory (VIDRL) | Victorian Infectious Diseases Reference Laboratory and Microbiological Diagnostic Unit Public Health Laboratory, Doherty Institute | Caly L., Seemann T., Schultz M., Druce J., Taiaroa, G. |
| 377 | EPI_ISL_416425 | hCoV-19/Hangzhou/ZJU-07/2020 | *Homo sapiens* | Asia / China / Hangzhou | 2020-02-03 | State Key Laboratory for Diagnosis and Treatment of Infectious Diseases, National Clinical Research Center for Infectious Diseases, First Affiliated Hospital, Zhejiang University School of Medicine, Hangzhou, China 310003 | State Key Laboratory for Diagnosis and Treatment of Infectious Diseases, National Clinical Research Center for Infectious Diseases, First Affiliated Hospital, Zhejiang University School of Medicine, Hangzhou, China 310003 | Hangping Yao, Nanping Wu, Chao Jiang, Xiangyun Lu, Linfang Cheng, Fumin Liu, Zhigang Wu, Haibo Wu, Changzhong Jin, Min Zheng, Lanjuan Li |
| 378 | EPI_ISL_416426 | hCoV-19/Hungary/mbl1/2020 | *Homo sapiens* | Europe / Hungary / Budapest | 2020-03-17 | Virological Research Group, Szentágothai Research Centre, University of Pécs | Bioinformatics Research Group, Szentágothai Research Centre, University of Pécs | Péter Urbán, Endre Gábor Tóth, Gábor Kemenesi, Róbert Herczeg, Attila Gyenesei, Ferenc Jakab |
| 379 | EPI_ISL_416429 | hCoV-19/Vietnam/CM99/2020 | *Homo sapiens* | Asia / Vietnam / Vinhphuc | 2020-02-11 | National Influenza Center, National Institute of Hygiene and Epidemiology (NIHE) | National Influenza Center, National Institute of Hygiene and Epidemiology (NIHE) | Le Quynh Mai, Taichiro Takemura, Meng Ling Moi, Takeshi Nabeshima, Nguyen Le Khanh Hang, Hoang Vu Mai Phuong, Ung Thi Hong Trang, Le Thi Thanh, Nguyen Vu Son, Vuong Duc Cuong, Pham Thi Hien, Tran Thu Huong, Nguyen Phuong Anh, Pham Hong Quynh Anh, Kouichi Morita, Futoshi Hasebe, Dang Duc Anh |
| 380 | EPI_ISL_416432 | hCoV-19/Saudi Arabia/KAIMRC-Alghoribi/2020 | *Homo sapiens* | Asia / Saudi Arabia / Riyadh | 2020-03-07 | Clinical Microbiology Lab | Infectious Disease Research Department, King Abdullah International Medical Research Center (KAIMRC) | Majed Alghoribi, Sadeem Alhayli, Abdulrahman Alswaji, Liliane Okdah, Sameera Al Johani, Michel Doumith |
| 381 | EPI_ISL_416433 | hCoV-19/USA/WA-UW77/2020 | *Homo sapiens* | North America / USA / Washington | 2020-03-10 | UW Virology Lab | UW Virology Lab | Pavitra Roychoudhury, Hong Xie, Keith Jerome, Alexander Greninger |
| 382 | EPI_ISL_416434 | hCoV-19/USA/WA-UW78/2020 | *Homo sapiens* | North America / USA / Washington | 2020-03-10 | UW Virology Lab | UW Virology Lab | Pavitra Roychoudhury, Hong Xie, Keith Jerome, Alexander Greninger |
| 383 | EPI_ISL_416435 | hCoV-19/USA/WA-UW79/2020 | *Homo sapiens* | North America / USA / Washington | 2020-03-10 | UW Virology Lab | UW Virology Lab | Pavitra Roychoudhury, Hong Xie, Keith Jerome, Alexander Greninger |
| 384 | EPI_ISL_416436 | hCoV-19/USA/WA-UW80/2020 | *Homo sapiens* | North America / USA / Washington | 2020-03-10 | UW Virology Lab | UW Virology Lab | Pavitra Roychoudhury, Hong Xie, Keith Jerome, Alexander Greninger |

| | | | | | | | | |
|---|---|---|---|---|---|---|---|---|
| 385 | EPI_ISL_416438 | hCoV-19/USA/WA-UW82/2020 | *Homo sapiens* | North America / USA / Washington | 2020-03-10 | UW Virology Lab | UW Virology Lab | Pavitra Roychoudhury, Hong Xie, Keith Jerome, Alexander Greninger |
| 386 | EPI_ISL_416439 | hCoV-19/USA/WA-UW83/2020 | *Homo sapiens* | North America / USA / Washington | 2020-03-10 | UW Virology Lab | UW Virology Lab | Pavitra Roychoudhury, Hong Xie, Keith Jerome, Alexander Greninger |
| 387 | EPI_ISL_416441 | hCoV-19/USA/WA-UW85/2020 | *Homo sapiens* | North America / USA / Washington | 2020-03-10 | UW Virology Lab | UW Virology Lab | Pavitra Roychoudhury, Hong Xie, Keith Jerome, Alexander Greninger |
| 388 | EPI_ISL_416442 | hCoV-19/USA/WA-UW86/2020 | *Homo sapiens* | North America / USA / Washington | 2020-03-10 | UW Virology Lab | UW Virology Lab | Pavitra Roychoudhury, Hong Xie, Keith Jerome, Alexander Greninger |
| 389 | EPI_ISL_416443 | hCoV-19/USA/WA-UW87/2020 | *Homo sapiens* | North America / USA / Washington | 2020-03-10 | UW Virology Lab | UW Virology Lab | Pavitra Roychoudhury, Hong Xie, Keith Jerome, Alexander Greninger |
| 390 | EPI_ISL_416444 | hCoV-19/USA/WA-UW88/2020 | *Homo sapiens* | North America / USA / Washington | 2020-03-10 | UW Virology Lab | UW Virology Lab | Pavitra Roychoudhury, Hong Xie, Keith Jerome, Alexander Greninger |
| 391 | EPI_ISL_416445 | hCoV-19/USA/WA-UW89/2020 | *Homo sapiens* | North America / USA / Washington | 2020-03-10 | UW Virology Lab | UW Virology Lab | Pavitra Roychoudhury, Hong Xie, Keith Jerome, Alexander Greninger |
| 392 | EPI_ISL_416447 | hCoV-19/USA/WA-UW91/2020 | *Homo sapiens* | North America / USA / Washington | 2020-03-10 | UW Virology Lab | UW Virology Lab | Pavitra Roychoudhury, Hong Xie, Keith Jerome, Alexander Greninger |
| 393 | EPI_ISL_416448 | hCoV-19/USA/WA-UW92/2020 | *Homo sapiens* | North America / USA / Washington | 2020-03-11 | UW Virology Lab | UW Virology Lab | Pavitra Roychoudhury, Hong Xie, Keith Jerome, Alexander Greninger |
| 394 | EPI_ISL_416451 | hCoV-19/USA/WA-UW95/2020 | *Homo sapiens* | North America / USA / Washington | 2020-03-10 | UW Virology Lab | UW Virology Lab | Pavitra Roychoudhury, Hong Xie, Keith Jerome, Alexander Greninger |
| 395 | EPI_ISL_416452 | hCoV-19/USA/WA-UW96/2020 | *Homo sapiens* | North America / USA / Washington | 2020-03-10 | UW Virology Lab | UW Virology Lab | Pavitra Roychoudhury, Hong Xie, Keith Jerome, Alexander Greninger |
| 396 | EPI_ISL_416457 | hCoV-19/USA/CA-MG0987/2020 | *Homo sapiens* | North America / USA / California / San Diego County | 2020-03-18 | Andersen Lab, The Scripps Research Institute | Andersen Lab, The Scripps Research Institute | Mark Zeller, Catie Anderson, Emily Spender, Sarah Topol, Raphaelle Klitting, Refugio Robles-Sikisaka, Karthik Gangavarapu, Laura Nicholson, Kristian Andersen |
| 397 | EPI_ISL_416466 | hCoV-19/USA/WA-S11/2020 | *Homo sapiens* | North America / USA / Washington / King County | 2020-03-03 | Seattle Flu Study | Seattle Flu Study | Chu et al |
| 398 | EPI_ISL_416467 | hCoV-19/Belgium/MTR-03021/2020 | *Homo sapiens* | Europe / Belgium / Holsbeek | 2020-03-02 | KU Leuven, Clinical and Epidemiological Virology | KU Leuven, Clinical and Epidemiological Virology | Bert Vanmechelen, Tony Wawina, Joan Marti-Carreras, Piet Maes |
| 399 | EPI_ISL_416468 | hCoV-19/Belgium/GMH-03022/2020 | *Homo sapiens* | Europe / Belgium / Holsbeek | 2020-03-02 | KU Leuven, Clinical and Epidemiological Virology | KU Leuven, Clinical and Epidemiological Virology | Bert Vanmechelen, Tony Wawina, Joan Marti-Carreras, Piet Maes |
| 400 | EPI_ISL_416469 | hCoV-19//Belgium/SN-03031/2020 | *Homo sapiens* | Europe / Belgium / Kessel-Lo | 2020-03-03 | KU Leuven, Clinical and Epidemiological Virology | KU Leuven, Clinical and Epidemiological Virology | Bert Vanmechelen, Tony Wawina, Joan Marti-Carreras, Piet Maes |
| 401 | EPI_ISL_416470 | hCoV-19/Belgium/DB-03023/2020 | *Homo sapiens* | Europe / Belgium / Couthuin | 2020-03-02 | KU Leuven, Clinical and Epidemiological Virology | KU Leuven, Clinical and Epidemiological Virology | Bert Vanmechelen, Tony Wawina, Joan Marti-Carreras, Piet Maes |
| 402 | EPI_ISL_416471 | hCoV-19/Belgium/DBD-03024/2020 | *Homo sapiens* | Europe / Belgium / Kessel-Lo | 2020-03-02 | KU Leuven, Clinical and Epidemiological Virology | KU Leuven, Clinical and Epidemiological Virology | Bert Vanmechelen, Tony Wawina, Joan Marti-Carreras, Piet Maes |
| 403 | EPI_ISL_416472 | hCoV-19/Belgium/UMF-03025/2020 | *Homo sapiens* | Europe / Belgium / Kessel-Lo | 2020-03-02 | KU Leuven, Clinical and Epidemiological Virology | KU Leuven, Clinical and Epidemiological Virology | Bert Vanmechelen, Tony Wawina, Joan Marti-Carreras, Piet Maes |

| | | | | | | | | |
|---|---|---|---|---|---|---|---|---|
| 404 | EPI_ISL_416473 | hCoV-19/Hangzhou/ZJU-08/2020 | *Homo sapiens* | Asia / China / Hangzhou | 2020-01-26 | State Key Laboratory for Diagnosis and Treatment of Infectious Diseases, National Clinical Research Center for Infectious Diseases, First Affiliated Hospital, Zhejiang University School of Medicine, Hangzhou, China 310003 | State Key Laboratory for Diagnosis and Treatment of Infectious Diseases, National Clinical Research Center for Infectious Diseases, First Affiliated Hospital, Zhejiang University School of Medicine, Hangzhou, China 310003 | Hangping Yao, Nanping Wu, Chao Jiang, Xiangyun Lu, Linfang Cheng, Fumin Liu, Zhigang Wu, Haibo Wu, Changzhong Jin, Min Zheng, Lanjuan Li |
| 405 | EPI_ISL_416474 | hCoV-19/Hangzhou/ZJU-09/2020 | *Homo sapiens* | Asia / China / Hangzhou | 2020-01-28 | State Key Laboratory for Diagnosis and Treatment of Infectious Diseases, National Clinical Research Center for Infectious Diseases, First Affiliated Hospital, Zhejiang University School of Medicine, Hangzhou, China 310003 | State Key Laboratory for Diagnosis and Treatment of Infectious Diseases, National Clinical Research Center for Infectious Diseases, First Affiliated Hospital, Zhejiang University School of Medicine, Hangzhou, China 310003 | Hangping Yao, Nanping Wu, Chao Jiang, Xiangyun Lu, Linfang Cheng, Fumin Liu, Zhigang Wu, Haibo Wu, Changzhong Jin, Min Zheng, Lanjuan Li |
| 406 | EPI_ISL_416475 | hCoV-19/Belgium/DBA-03032/2020 | *Homo sapiens* | Europe / Belgium / Leuven | 2020-03-03 | KU Leuven, Clinical and Epidemiological Virology | KU Leuven, Clinical and Epidemiological Virology | Bert Vanmechelen, Tony Wawina, Joan Marti-Carreras, Piet Maes |
| 407 | EPI_ISL_416476 | hCoV-19/Belgium/MTR-03026/2020 | *Homo sapiens* | Europe / Belgium / Holsbeek | 2020-03-02 | KU Leuven, Clinical and Epidemiological Virology | KU Leuven, Clinical and Epidemiological Virology | Bert Vanmechelen, Tony Wawina, Joan Marti-Carreras, Piet Maes |
| 408 | EPI_ISL_416477 | hCoV-19/Georgia/Tb-390/2020 | *Homo sapiens* | Asia / Georgia / Tbilisi | 2020-03-08 | R. G. Lugar Center for Public Health Research, National Center for Disease Control and Public Health (NCDC) of Georgia. | R. G. Lugar Center for Public Health Research, National Center for Disease Control and Public Health (NCDC) of Georgia. | Marine Murtskhvaladze, Nato Kotaria, Ann Machablishvili, Lela Sabadze, Mari Gavashelidze, Ana Papkiauri, Meri Pantsulaia, Gvantsa Brachveli, Tata Imnadze, Tamar Jashiashvili, Tea Tevdoradze, Ketevan Sidamonidze, Ekaterine Khmaladze, Ekaterine Zhghenti, Roena Sukhiashvili, Mariam Zakalashvili, Lela Urushadze, Magda Dgebuadze, Giorgi Tomashvili, Davit Tsaguria, Ekaterine Zangaladze, Nino Berishvili, Gvantsa Chanturia, Adam Kotorashvili, Maia Alkhazashvili, Irma Burjanadze, Anna Kasradze, Khatuna Zakhashvili, Paata Imnadze, Amiran Gamkrelidze. |
| 409 | EPI_ISL_416478 | hCoV-19/Georgia/Tb-673/2020 | *Homo sapiens* | Asia / Georgia / Tbilisi | 2020-03-14 | R. G. Lugar Center for Public Health Research, National Center for Disease Control and Public Health (NCDC) of Georgia. | R. G. Lugar Center for Public Health Research, National Center for Disease Control and Public Health (NCDC) of Georgia. | Marine Murtskhvaladze, Nato Kotaria, Ann Machablishvili, Lela Sabadze, Mari Gavashelidze, Ana Papkiauri, Meri Pantsulaia, Gvantsa Brachveli, Tata Imnadze, Tamar Jashiashvili, Tea Tevdoradze, Ketevan Sidamonidze, Ekaterine Khmaladze, Ekaterine Zhghenti, Roena Sukhiashvili, Mariam Zakalashvili, Lela Urushadze, Magda Dgebuadze, Giorgi Tomashvili, Davit Tsaguria, Ekaterine Zangaladze, Nino Berishvili, Gvantsa Chanturia, Adam Kotorashvili, Maia Alkhazashvili, Irma Burjanadze, Anna Kasradze, Khatuna Zakhashvili, Paata Imnadze, Amiran Gamkrelidze. |
| 410 | EPI_ISL_416479 | hCoV-19/Georgia/Tb-273/2020 | *Homo sapiens* | Asia / Georgia / Tbilisi | 2020-03-05 | R. G. Lugar Center for Public Health Research, National Center for Disease Control and Public Health (NCDC) of Georgia. | R. G. Lugar Center for Public Health Research, National Center for Disease Control and Public Health (NCDC) of Georgia. | Marine Murtskhvaladze, Nato Kotaria, Ann Machablishvili, Lela Sabadze, Mari Gavashelidze, Ana Papkiauri, Meri Pantsulaia, Gvantsa Brachveli, Tata Imnadze, Tamar Jashiashvili, Tea Tevdoradze, Ketevan Sidamonidze, Ekaterine Khmaladze, Ekaterine Zhghenti, Roena Sukhiashvili, Mariam Zakalashvili, Lela Urushadze, Magda Dgebuadze, Giorgi Tomashvili, Davit Tsaguria, Ekaterine Zangaladze, Nino Berishvili, Gvantsa Chanturia, Adam Kotorashvili, Maia Alkhazashvili, Irma Burjanadze, Anna Kasradze, Khatuna Zakhashvili, Paata Imnadze, Amiran Gamkrelidze. |

| | | | | | | | | |
|---|---|---|---|---|---|---|---|---|
| 411 | EPI_ISL_416480 | hCoV-19/Georgia/Tb-537/2020 | *Homo sapiens* | Asia / Georgia / Tbilisi | 2020-03-11 | R. G. Lugar Center for Public Health Research, National Center for Disease Control and Public Health (NCDC) of Georgia. | R. G. Lugar Center for Public Health Research, National Center for Disease Control and Public Health (NCDC) of Georgia. | Ann Machablishvili, Nato Kotaria, Marine Murtskhvaladze, Lela Sabadze, Mari Gavashelidze, Ana Papkiauri, Meri Pantsulaia, Gvantsa Brachveli, Tata Imnadze, Tamar Jashiashvili, Tea Tevdoradze, Ketevan Sidamonidze, Ekaterine Khmaladze, Ekaterine Zhghenti, Roena Sukhiashvili, Mariam Zakalashvili, Lela Urushadze, Magda Dgebuadze, Giorgi Tomashvili, Davit Tsaguria, Ekaterine Zangaladze, Nino Berishvili, Gvantsa Chanturia, Adam Kotorashvili, Maia Alkhazashvili, Irma Burjanadze, Anna Kasradze, Khatuna Zakhashvili, Paata Imnadze, Amiran Gamkrelidze. |
| 412 | EPI_ISL_416481 | hCoV-19/Georgia/Tb-712/2020 | *Homo sapiens* | Asia / Georgia / Tbilisi | 2020-03-16 | R. G. Lugar Center for Public Health Research, National Center for Disease Control and Public Health (NCDC) of Georgia. | R. G. Lugar Center for Public Health Research, National Center for Disease Control and Public Health (NCDC) of Georgia. | Gvantsa Chanturia, Marine Murtskhvaladze, Nato Kotaria, Ann Machablishvili, Lela Sabadze, Mari Gavashelidze, Ana Papkiauri, Meri Pantsulaia, Gvantsa Brachveli, Tata Imnadze, Tamar Jashiashvili, Tea Tevdoradze, Ketevan Sidamonidze, Ekaterine Khmaladze, Ekaterine Zhghenti, Roena Sukhiashvili, Mariam Zakalashvili, Lela Urushadze, Magda Dgebuadze, Giorgi Tomashvili, Davit Tsaguria, Ekaterine Zangaladze, Nino Berishvili, Adam Kotorashvili, Maia Alkhazashvili, Irma Burjanadze, Anna Kasradze, Khatuna Zakhashvili, Paata Imnadze, Amiran Gamkrelidze. |
| 413 | EPI_ISL_416482 | hCoV-19/Georgia/Tb/2020 | *Homo sapiens* | Asia / Georgia / Tbilisi | 2020-03-13 | R. G. Lugar Center for Public Health Research, National Center for Disease Control and Public Health (NCDC) of Georgia. | R. G. Lugar Center for Public Health Research, National Center for Disease Control and Public Health (NCDC) of Georgia. | Adam Kotorashvili, Marine Murtskhvaladze, Nato Kotaria, Ann Machablishvili, Lela Sabadze, Mari Gavashelidze, Ana Papkiauri, Meri Pantsulaia, Gvantsa Brachveli, Tata Imnadze, Tamar Jashiashvili, Tea Tevdoradze, Ketevan Sidamonidze, Ekaterine Khmaladze, Ekaterine Zhghenti, Roena Sukhiashvili, Mariam Zakalashvili, Lela Urushadze, Magda Dgebuadze, Giorgi Tomashvili, Davit Tsaguria, Ekaterine Zangaladze, Nino Berishvili, Gvantsa Chanturia, Maia Alkhazashvili, Irma Burjanadze, Anna Kasradze, Khatuna Zakhashvili, Paata Imnadze, Amiran Gamkrelidze. |
| 414 | EPI_ISL_416484 | hCoV-19/Spain/Valencia5/2020 | *Homo sapiens* | Europe/Spain/Comunitat_Valenciana/Valencia | 2020-02-27 | Servicio de Microbiología. Consorcio Hospital General Universitario de Valencia | Sequencing and Bioinformatics Service and Molecular Epidemiology Research Group. FISABIO-Public Health | Maria Dolores Ocete, Concepcion Gimeno, Giuseppe D'Auria, Griselda De Marco, Neris Garcia-Gonzalez, Maria Alma Bracho, Fernando Gonzalez-Candelas |
| 414 | EPI_ISL_416487 | hCoV-19/Spain/Valencia8/2020 | *Homo sapiens* | Europe/Spain/Comunitat_Valenciana/Valencia | 2020-03-04 | Servicio de Microbiología. Consorcio Hospital General Universitario de Valencia | Sequencing and Bioinformatics Service and Molecular Epidemiology Research Group. FISABIO-Public Health | Giuseppe D'Auria, Griselda De Marco, Neris Garcia-Gonzalez, Maria Alma Bracho, Maria Dolores Ocete, Concepcion Gimeno, Fernando Gonzalez-Candelas |
| 415 | EPI_ISL_416488 | hCoV-19/Poland/PL_P1/2020 | *Homo sapiens* | Europe / Poland / Zielonogorskie | 2020-03-03 | ViroGenetics - BSL3 Laboratory of Virology; Homo sapiens Genome Variation Research Group & Genomics Centre MCB; Bioinformatics Research Group Department of Virology | ViroGenetics - BSL3 Laboratory of Virology; Homo sapiens Genome Variation Research Group & Genomics Centre MCB; Bioinformatics Research Group Department of Virology | Aleksandra Milewska, Ewelina Pośpiech, Agata Jarosz, Adrianna Klajmon, Kamila Marszałek, Katarzyna Pancer, Magdalena Rzeczkowska, Tomasz Wołkowicz, Katarzyna Zacharczuk, Agnieszka Kołakowska-Kulesza, Natalia Wolaniuk, Ewelina Hallman-Szelińska, Paweł P Łabaj, Wojciech Branicki, Krzysztof Pyrć |
| 416 | EPI_ISL_416494 | hCoV-19/France/N2223/2020 | *Homo sapiens* | Europe / France / Normandie / Rouen | 2020-03-04 | Centre Hositalier Universitaire de Rouen Laboratoire de Virologie | National Reference Center for Viruses of Respiratory Infections, Institut Pasteur, Paris | Mélnie Albert, Marion Barbet, Sylvie Behillil, Méline Bizard, Angela Brisebarre, Flora Donati, Etienne Simon-Lorière, Vincent Enouf, Maud Vanpeene, Sylvie van der Werf, Jean-Christophe Plantier |
| 417 | EPI_ISL_416495 | hCoV-19/France/HF2234/2020 | *Homo sapiens* | Europe / France / Hauts de France / Compiègne | 2020-03-10 | Centre Hospitalier Compiègne Laboratoire de Biologie | National Reference Center for Viruses of Respiratory Infections, Institut Pasteur, Paris | Mélnie Albert, Marion Barbet, Sylvie Behillil, Méline Bizard, Angela Brisebarre, Flora Donati, Etienne Simon-Lorière, Vincent Enouf, Maud Vanpeene, Sylvie van der Werf, Raulin Olivia |
| 418 | EPI_ISL_416496 | hCoV-19/France/HF2237/2020 | *Homo sapiens* | Europe / France / Hauts de France / Compiègne | 2020-03-10 | Centre Hospitalier Compiègne Laboratoire de Biologie | National Reference Center for Viruses of Respiratory Infections, Institut Pasteur, Paris | Mélnie Albert, Marion Barbet, Sylvie Behillil, Méline Bizard, Angela Brisebarre, Flora Donati, Etienne Simon-Lorière, Vincent Enouf, Maud Vanpeene, Sylvie van der Werf, Raulin Olivia |
| 419 | EPI_ISL_416497 | hCoV-19/France/HF2239/2020 | *Homo sapiens* | Europe / France / Hauts de France / Compiègne | 2020-03-10 | Centre Hospitalier Compiègne Laboratoire de Biologie | National Reference Center for Viruses of Respiratory Infections, Institut Pasteur, Paris | Mélnie Albert, Marion Barbet, Sylvie Behillil, Méline Bizard, Angela Brisebarre, Flora Donati, Etienne Simon-Lorière, Vincent Enouf, Maud Vanpeene, Sylvie van der Werf, Raulin Olivia |

| | | | | | | | | |
|---|---|---|---|---|---|---|---|---|
| 420 | EPI_ISL_416498 | hCoV-19/France/IDF2256/2020 | *Homo sapiens* | Europe / France / Ile de France / Garches | 2020-03-11 | Institut Médico légal- Hop R. Poincaré | National Reference Center for Viruses of Respiratory Infections, Institut Pasteur, Paris | Mélnie Albert, Marion Barbet, Sylvie Behillil, Méline Bizard, Angela Brisebarre, Flora Donati, Etienne Simon-Lorière, Vincent Enouf, Maud Vanpeene, Sylvie van der Werf |
| 421 | EPI_ISL_416499 | hCoV-19/France/IDF2278/2020 | *Homo sapiens* | Europe / France / Ile de France / Longjumeau | 2020-03-11 | LABM GH nord Essonne | National Reference Center for Viruses of Respiratory Infections, Institut Pasteur, Paris | Mélnie Albert, Marion Barbet, Sylvie Behillil, Méline Bizard, Angela Brisebarre, Flora Donati, Etienne Simon-Lorière, Vincent Enouf, Maud Vanpeene, Sylvie van der Werf |
| 422 | EPI_ISL_416502 | hCoV-19/France/B2330/2020 | *Homo sapiens* | Europe / France / Bretagne / Rennes | 2020-02-26 | CHRU Pontchaillou - Laboratoire de Virologie | National Reference Center for Viruses of Respiratory Infections, Institut Pasteur, Paris | Mélnie Albert, Marion Barbet, Sylvie Behillil, Méline Bizard, Angela Brisebarre, Flora Donati, Etienne Simon-Lorière, Vincent Enouf, Maud Vanpeene, Sylvie van der Werf, Gisèle Lagathu |
| 423 | EPI_ISL_416503 | hCoV-19/France/B2334/2020 | *Homo sapiens* | Europe / France / Bretagne / Rennes | 2020-03-01 | CHRU Pontchaillou - Laboratoire de Virologie | National Reference Center for Viruses of Respiratory Infections, Institut Pasteur, Paris | Mélnie Albert, Marion Barbet, Sylvie Behillil, Méline Bizard, Angela Brisebarre, Flora Donati, Etienne Simon-Lorière, Vincent Enouf, Maud Vanpeene, Sylvie van der Werf, Gisèle Lagathu |
| 424 | EPI_ISL_416505 | hCoV-19/France/B2336/2020 | *Homo sapiens* | Europe / France / Bretagne / Rennes | 2020-03-02 | CHRU Pontchaillou - Laboratoire de Virologie | National Reference Center for Viruses of Respiratory Infections, Institut Pasteur, Paris | Mélnie Albert, Marion Barbet, Sylvie Behillil, Méline Bizard, Angela Brisebarre, Flora Donati, Etienne Simon-Lorière, Vincent Enouf, Maud Vanpeene, Sylvie van der Werf, Gisèle Lagathu |
| 425 | EPI_ISL_416506 | hCoV-19/France/B2337/2020 | *Homo sapiens* | Europe / France / Bretagne / Rennes | 2020-03-03 | CHRU Pontchaillou - Laboratoire de Virologie | National Reference Center for Viruses of Respiratory Infections, Institut Pasteur, Paris | Mélnie Albert, Marion Barbet, Sylvie Behillil, Méline Bizard, Angela Brisebarre, Flora Donati, Etienne Simon-Lorière, Vincent Enouf, Maud Vanpeene, Sylvie van der Werf, Gisèle Lagathu |
| 426 | EPI_ISL_416507 | hCoV-19/France/B2340/2020 | *Homo sapiens* | Europe / France / Bretagne / Rennes | 2020-03-05 | CHRU Pontchaillou - Laboratoire de Virologie | National Reference Center for Viruses of Respiratory Infections, Institut Pasteur, Paris | Mélnie Albert, Marion Barbet, Sylvie Behillil, Méline Bizard, Angela Brisebarre, Flora Donati, Etienne Simon-Lorière, Vincent Enouf, Maud Vanpeene, Sylvie van der Werf, Gisèle Lagathu |
| 427 | EPI_ISL_416509 | hCoV-19/France/B2344/2020 | *Homo sapiens* | Europe / France / Bretagne / Rennes | 2020-03-06 | CHRU Pontchaillou - Laboratoire de Virologie | National Reference Center for Viruses of Respiratory Infections, Institut Pasteur, Paris | Mélnie Albert, Marion Barbet, Sylvie Behillil, Méline Bizard, Angela Brisebarre, Flora Donati, Etienne Simon-Lorière, Vincent Enouf, Maud Vanpeene, Sylvie van der Werf, Gisèle Lagathu |
| 428 | EPI_ISL_416510 | hCoV-19/France/B2346/2020 | *Homo sapiens* | Europe / France / Bretagne / Rennes | 2020-03-06 | CHRU Pontchaillou - Laboratoire de Virologie | National Reference Center for Viruses of Respiratory Infections, Institut Pasteur, Paris | Mélnie Albert, Marion Barbet, Sylvie Behillil, Méline Bizard, Angela Brisebarre, Flora Donati, Etienne Simon-Lorière, Vincent Enouf, Maud Vanpeene, Sylvie van der Werf, Gisèle Lagathu |
| 429 | EPI_ISL_416513 | hCoV-19/France/B2351/2020 | *Homo sapiens* | Europe / France / Bretagne / Rennes | 2020-03-07 | CHRU Pontchaillou - Laboratoire de Virologie | National Reference Center for Viruses of Respiratory Infections, Institut Pasteur, Paris | Mélnie Albert, Marion Barbet, Sylvie Behillil, Méline Bizard, Angela Brisebarre, Flora Donati, Etienne Simon-Lorière, Vincent Enouf, Maud Vanpeene, Sylvie van der Werf, Gisèle Lagathu |
| 430 | EPI_ISL_416519 | hCoV-19/New Zealand/20VR0189/2020 | *Homo sapiens* | Oceania / New Zealand / Auckland | 2020-03-02 | Auckland Hospital | Institute of Environmental Science and Research (ESR) | Matt Storey, Xiaoyun Ren, Gary McAuliffe, Sally Roberts, Matthew Blakiston, Erasmus Smit, Lauren Jelly, Joep de Ligt |
| 430 | EPI_ISL_416732 | hCoV-19/England/SHEF-BFCDF/2020 | *Homo sapiens* | Europe / England / London | 2020-03-03 | Virology Department, Sheffield Teaching Hospitals NHS Foundation Trust | Department of Infection, Immunity and Cardiovascular Disease, The Florey Institute, The Medical School, University of Sheffield | Thushan de Silva, Matthew Parker, Adri Angyal, Rebecca Brown, Matthew Wyles, Mehmet Yavuz, Mohammad Raza, Cariad Evans |
| 430 | EPI_ISL_416734 | hCoV-19/England/SHEF-BFCFD/2020 | *Homo sapiens* | Europe / England / South Yorkshire | 2020-03-09 | Virology Department, Sheffield Teaching Hospitals NHS Foundation Trust | Department of Infection, Immunity and Cardiovascular Disease, The Florey Institute, The Medical School, University of Sheffield | Thushan de Silva, Matthew Parker, Adri Angyal, Rebecca Brown, Matthew Wyles, Mehmet Yavuz, Mohammad Raza, Cariad Evans |
| 431 | EPI_ISL_416521 | hCoV-19/Saudi Arabia/SCDC-3321/2020 | *Homo sapiens* | Asia / Saudi Arabia | 2020-03-10 | Public Health Laboratory | Public Health Laboratory, Saudi CDC | Albarrag, A |
| 431 | EPI_ISL_416522 | hCoV-19/Saudi Arabia/SCDC-3324/2020 | *Homo sapiens* | Asia / Saudi Arabia | 2020-03-10 | Public Health Laboratory, Saudi CDC | Public Health Laboratory, Saudi CDC | Albarrag,A |
| 432 | EPI_ISL_416524 | hCoV-19/Japan/SMU-0311S2/2020 | *Homo sapiens* | Asia / Japan / Saitama | 2020-03-11 | Saitama Medical University Hospital | Saitama Medical University | Kazuo Imai |
| 433 | EPI_ISL_416525 | hCoV-19/Japan/SMU-0311S3/2020 | *Homo sapiens* | Asia / Japan / Saitama | 2020-03-11 | Saitama Medical University | Saitama Medical University | Kazuo Imai |
| 434 | EPI_ISL_416538 | hCoV-19/New Zealand/20VR0275/2020 | *Homo sapiens* | Oceania / New Zealand / Wellington | 2020-03-15 | Wellington Hospital | Institute of Environmental Science and Research (ESR) | Wellington SCL, Wellington Hospital, Riddiford Street, Newtown, Wellington 6021, New Zealand |
| 435 | EPI_ISL_416539 | hCoV-19/New Zealand/20VR0276/2020 | *Homo sapiens* | Oceania / New Zealand / Wellington | 2020-03-15 | Wellington Hospital | Institute of Environmental Science and Research (ESR) | Matt Storey, Xiaoyun Ren, Craig Thornley, Maxim Bloomfield, Erasmus Smit, Lauren Jelly, Joep de Ligt |
| 436 | EPI_ISL_416541 | hCoV-19/Kuwait/KU09/2020 | *Homo sapiens* | Asia / Kuwait / Dasman | 2020-03-02 | Dasman Diabetes Institute and Virology Laboratory Ministry of Health | Dasman Diabetes Institute | Fahd Al-Mulla, Sumi John, Rasheeba Iqbal, Motasem Melhem, Ebaa AlOzairi, Sara Al-Qabandi, Qais Al-Duwairi |

| | | | | | | | | |
|---|---|---|---|---|---|---|---|---|
| 437 | EPI_ISL_416542 | hCoV-19/Kuwait/KU17/2020 | *Homo sapiens* | Asia / Kuwait / Dasman | 2020-03-02 | Dasman Diabetes Institute | Dasman Diabetes Institute | Fahd Al-Mulla, Sumi John, Rasheeba Iqbal, Motasem Melhem, Ebaa AlOzairi, Sara Al-Qabandi, Qais Al-Duwairi |
| 438 | EPI_ISL_416566 | hCoV-19/Japan/DP0027/2020 | *Homo sapiens* | Asia / Japan / unknown | 2020-02-15 | Japanese Quarantine Stations | Pathogen Genomics Center, National Institute of Infectious Diseases | Tsuyoshi Sekizuka, Kentaro Itokawa, Rina Tanaka, Masanori Hashino, Tsutomu Kageyama, Shinji Saito, Ikuyo Takayama, Hideki Hasegawa, Takuri Takahashi, Hajime Kamiya, Takuya Yamagishi, Motoi Suzuki, Takaji Wakita, Makoto Kuroda |
| 439 | EPI_ISL_416567 | hCoV-19/Japan/DP0037/2020 | *Homo sapiens* | Asia / Japan / unknown | 2020-02-15 | Japanese Quarantine Stations | Pathogen Genomics Center, National Institute of Infectious Diseases | Tsuyoshi Sekizuka, Kentaro Itokawa, Rina Tanaka, Masanori Hashino, Tsutomu Kageyama, Shinji Saito, Ikuyo Takayama, Hideki Hasegawa, Takuri Takahashi, Hajime Kamiya, Takuya Yamagishi, Motoi Suzuki, Takaji Wakita, Makoto Kuroda |
| 440 | EPI_ISL_416569 | hCoV-19/Japan/DP0059/2020 | *Homo sapiens* | Asia / Japan / unknown | 2020-02-15 | Japanese Quarantine Stations | Pathogen Genomics Center, National Institute of Infectious Diseases | Tsuyoshi Sekizuka, Kentaro Itokawa, Rina Tanaka, Masanori Hashino, Tsutomu Kageyama, Shinji Saito, Ikuyo Takayama, Hideki Hasegawa, Takuri Takahashi, Hajime Kamiya, Takuya Yamagishi, Motoi Suzuki, Takaji Wakita, Makoto Kuroda |
| 441 | EPI_ISL_416570 | hCoV-19/Japan/DP0065/2020 | *Homo sapiens* | Asia / Japan / unknown | 2020-02-15 | Japanese Quarantine Stations | Pathogen Genomics Center, National Institute of Infectious Diseases | Tsuyoshi Sekizuka, Kentaro Itokawa, Rina Tanaka, Masanori Hashino, Tsutomu Kageyama, Shinji Saito, Ikuyo Takayama, Hideki Hasegawa, Takuri Takahashi, Hajime Kamiya, Takuya Yamagishi, Motoi Suzuki, Takaji Wakita, Makoto Kuroda |
| 442 | EPI_ISL_416571 | hCoV-19/Japan/DP0077/2020 | *Homo sapiens* | Asia / Japan / unknown | 2020-02-15 | Japanese Quarantine Stations | Pathogen Genomics Center, National Institute of Infectious Diseases | Tsuyoshi Sekizuka, Kentaro Itokawa, Rina Tanaka, Masanori Hashino, Tsutomu Kageyama, Shinji Saito, Ikuyo Takayama, Hideki Hasegawa, Takuri Takahashi, Hajime Kamiya, Takuya Yamagishi, Motoi Suzuki, Takaji Wakita, Makoto Kuroda |
| 443 | EPI_ISL_416572 | hCoV-19/Japan/DP0078/2020 | *Homo sapiens* | Asia / Japan / unknown | 2020-02-15 | Japanese Quarantine Stations | Pathogen Genomics Center, National Institute of Infectious Diseases | Tsuyoshi Sekizuka, Kentaro Itokawa, Rina Tanaka, Masanori Hashino, Tsutomu Kageyama, Shinji Saito, Ikuyo Takayama, Hideki Hasegawa, Takuri Takahashi, Hajime Kamiya, Takuya Yamagishi, Motoi Suzuki, Takaji Wakita, Makoto Kuroda |
| 443 | EPI_ISL_416573 | hCoV-19/Japan/DP0104/2020 | *Homo sapiens* | Asia / Japan / unknown | 2020-02-15 | Japanese Quarantine Stations | Pathogen Genomics Center, National Institute of Infectious Diseases | Tsuyoshi Sekizuka, Kentaro Itokawa, Rina Tanaka, Masanori Hashino, Tsutomu Kageyama, Shinji Saito, Ikuyo Takayama, Hideki Hasegawa, Takuri Takahashi, Hajime Kamiya, Takuya Yamagishi, Motoi Suzuki, Takaji Wakita, Makoto Kuroda |
| 443 | EPI_ISL_416575 | hCoV-19/Japan/DP0121/2020 | *Homo sapiens* | Asia / Japan / unknown | 2020-02-15 | Japanese Quarantine Stations | Pathogen Genomics Center, National Institute of Infectious Diseases | Tsuyoshi Sekizuka, Kentaro Itokawa, Rina Tanaka, Masanori Hashino, Tsutomu Kageyama, Shinji Saito, Ikuyo Takayama, Hideki Hasegawa, Takuri Takahashi, Hajime Kamiya, Takuya Yamagishi, Motoi Suzuki, Takaji Wakita, Makoto Kuroda |
| 443 | EPI_ISL_416576 | hCoV-19/Japan/DP0133/2020 | *Homo sapiens* | Asia / Japan / unknown | 2020-02-15 | Japanese Quarantine Stations | Pathogen Genomics Center, National Institute of Infectious Diseases | Tsuyoshi Sekizuka, Kentaro Itokawa, Rina Tanaka, Masanori Hashino, Tsutomu Kageyama, Shinji Saito, Ikuyo Takayama, Hideki Hasegawa, Takuri Takahashi, Hajime Kamiya, Takuya Yamagishi, Motoi Suzuki, Takaji Wakita, Makoto Kuroda |
| 443 | EPI_ISL_416580 | hCoV-19/Japan/DP0184/2020 | *Homo sapiens* | Asia / Japan / unknown | 2020-02-15 | Japanese Quarantine Stations | Pathogen Genomics Center, National Institute of Infectious Diseases | Tsuyoshi Sekizuka, Kentaro Itokawa, Rina Tanaka, Masanori Hashino, Tsutomu Kageyama, Shinji Saito, Ikuyo Takayama, Hideki Hasegawa, Takuri Takahashi, Hajime Kamiya, Takuya Yamagishi, Motoi Suzuki, Takaji Wakita, Makoto Kuroda |
| 443 | EPI_ISL_416629 | hCoV-19/Japan/DP0802/2020 | *Homo sapiens* | Asia / Japan / unknown | 2020-02-17 | Japanese Quarantine Stations | Pathogen Genomics Center, National Institute of Infectious Diseases | Tsuyoshi Sekizuka, Kentaro Itokawa, Rina Tanaka, Masanori Hashino, Tsutomu Kageyama, Shinji Saito, Ikuyo Takayama, Hideki Hasegawa, Takuri Takahashi, Hajime Kamiya, Takuya Yamagishi, Motoi Suzuki, Takaji Wakita, Makoto Kuroda |
| 444 | EPI_ISL_416578 | hCoV-19/Japan/DP0152/2020 | *Homo sapiens* | Asia / Japan / unknown | 2020-02-15 | Japanese Quarantine Stations | Pathogen Genomics Center, National Institute of Infectious Diseases | Tsuyoshi Sekizuka, Kentaro Itokawa, Rina Tanaka, Masanori Hashino, Tsutomu Kageyama, Shinji Saito, Ikuyo Takayama, Hideki Hasegawa, Takuri Takahashi, Hajime Kamiya, Takuya Yamagishi, Motoi Suzuki, Takaji Wakita, Makoto Kuroda |
| 445 | EPI_ISL_416579 | hCoV-19/Japan/DP0158/2020 | *Homo sapiens* | Asia / Japan / unknown | 2020-02-15 | Japanese Quarantine Stations | Pathogen Genomics Center, National Institute of Infectious Diseases | Tsuyoshi Sekizuka, Kentaro Itokawa, Rina Tanaka, Masanori Hashino, Tsutomu Kageyama, Shinji Saito, Ikuyo Takayama, Hideki Hasegawa, Takuri Takahashi, Hajime Kamiya, Takuya Yamagishi, Motoi Suzuki, Takaji Wakita, Makoto Kuroda |
| 446 | EPI_ISL_416581 | hCoV-19/Japan/DP0190/2020 | *Homo sapiens* | Asia / Japan / unknown | 2020-02-15 | Japanese Quarantine Stations | Pathogen Genomics Center, National Institute of Infectious Diseases | Tsuyoshi Sekizuka, Kentaro Itokawa, Rina Tanaka, Masanori Hashino, Tsutomu Kageyama, Shinji Saito, Ikuyo Takayama, Hideki Hasegawa, Takuri Takahashi, Hajime Kamiya, Takuya Yamagishi, Motoi Suzuki, Takaji Wakita, Makoto Kuroda |

| | | | | | | | | |
|---|---|---|---|---|---|---|---|---|
| 447 | EPI_ISL_416582 | hCoV-19/Japan/DP0191/2020 | *Homo sapiens* | Asia / Japan / unknown | 2020-02-15 | Japanese Quarantine Stations | Pathogen Genomics Center, National Institute of Infectious Diseases | Tsuyoshi Sekizuka, Kentaro Itokawa, Rina Tanaka, Masanori Hashino, Tsutomu Kageyama, Shinji Saito, Ikuyo Takayama, Hideki Hasegawa, Takuri Takahashi, Hajime Kamiya, Takuya Yamagishi, Motoi Suzuki, Takaji Wakita, Makoto Kuroda |
| 448 | EPI_ISL_416583 | hCoV-19/Japan/DP0196/2020 | *Homo sapiens* | Asia / Japan / unknown | 2020-02-15 | Japanese Quarantine Stations | Pathogen Genomics Center, National Institute of Infectious Diseases | Tsuyoshi Sekizuka, Kentaro Itokawa, Rina Tanaka, Masanori Hashino, Tsutomu Kageyama, Shinji Saito, Ikuyo Takayama, Hideki Hasegawa, Takuri Takahashi, Hajime Kamiya, Takuya Yamagishi, Motoi Suzuki, Takaji Wakita, Makoto Kuroda |
| 449 | EPI_ISL_416584 | hCoV-19/Japan/DP0200/2020 | *Homo sapiens* | Asia / Japan / unknown | 2020-02-15 | Japanese Quarantine Stations | Pathogen Genomics Center, National Institute of Infectious Diseases | Tsuyoshi Sekizuka, Kentaro Itokawa, Rina Tanaka, Masanori Hashino, Tsutomu Kageyama, Shinji Saito, Ikuyo Takayama, Hideki Hasegawa, Takuri Takahashi, Hajime Kamiya, Takuya Yamagishi, Motoi Suzuki, Takaji Wakita, Makoto Kuroda |
| 450 | EPI_ISL_416585 | hCoV-19/Japan/DP0236/2020 | *Homo sapiens* | Asia / Japan / unknown | 2020-02-16 | Japanese Quarantine Stations | Pathogen Genomics Center, National Institute of Infectious Diseases | Tsuyoshi Sekizuka, Kentaro Itokawa, Rina Tanaka, Masanori Hashino, Tsutomu Kageyama, Shinji Saito, Ikuyo Takayama, Hideki Hasegawa, Takuri Takahashi, Hajime Kamiya, Takuya Yamagishi, Motoi Suzuki, Takaji Wakita, Makoto Kuroda |
| 451 | EPI_ISL_416586 | hCoV-19/Japan/DP0274/2020 | *Homo sapiens* | Asia / Japan / unknown | 2020-02-16 | Japanese Quarantine Stations | Pathogen Genomics Center, National Institute of Infectious Diseases | Tsuyoshi Sekizuka, Kentaro Itokawa, Rina Tanaka, Masanori Hashino, Tsutomu Kageyama, Shinji Saito, Ikuyo Takayama, Hideki Hasegawa, Takuri Takahashi, Hajime Kamiya, Takuya Yamagishi, Motoi Suzuki, Takaji Wakita, Makoto Kuroda |
| 451 | EPI_ISL_416595 | hCoV-19/Japan/DP0328/2020 | *Homo sapiens* | Asia / Japan / unknown | 2020-02-16 | Japanese Quarantine Stations | Pathogen Genomics Center, National Institute of Infectious Diseases | Tsuyoshi Sekizuka, Kentaro Itokawa, Rina Tanaka, Masanori Hashino, Tsutomu Kageyama, Shinji Saito, Ikuyo Takayama, Hideki Hasegawa, Takuri Takahashi, Hajime Kamiya, Takuya Yamagishi, Motoi Suzuki, Takaji Wakita, Makoto Kuroda |
| 451 | EPI_ISL_416606 | hCoV-19/Japan/DP0482/2020 | *Homo sapiens* | Asia / Japan / unknown | 2020-02-16 | Japanese Quarantine Stations | Pathogen Genomics Center, National Institute of Infectious Diseases | Tsuyoshi Sekizuka, Kentaro Itokawa, Rina Tanaka, Masanori Hashino, Tsutomu Kageyama, Shinji Saito, Ikuyo Takayama, Hideki Hasegawa, Takuri Takahashi, Hajime Kamiya, Takuya Yamagishi, Motoi Suzuki, Takaji Wakita, Makoto Kuroda |
| 451 | EPI_ISL_416634 | hCoV-19/Japan/DP0890/2020 | *Homo sapiens* | Asia / Japan / unknown | 2020-02-17 | Japanese Quarantine Stations | Pathogen Genomics Center, National Institute of Infectious Diseases | Tsuyoshi Sekizuka, Kentaro Itokawa, Rina Tanaka, Masanori Hashino, Tsutomu Kageyama, Shinji Saito, Ikuyo Takayama, Hideki Hasegawa, Takuri Takahashi, Hajime Kamiya, Takuya Yamagishi, Motoi Suzuki, Takaji Wakita, Makoto Kuroda |
| 452 | EPI_ISL_416587 | hCoV-19/Japan/DP0278/2020 | *Homo sapiens* | Asia / Japan / unknown | 2020-02-16 | Japanese Quarantine Stations | Pathogen Genomics Center, National Institute of Infectious Diseases | Tsuyoshi Sekizuka, Kentaro Itokawa, Rina Tanaka, Masanori Hashino, Tsutomu Kageyama, Shinji Saito, Ikuyo Takayama, Hideki Hasegawa, Takuri Takahashi, Hajime Kamiya, Takuya Yamagishi, Motoi Suzuki, Takaji Wakita, Makoto Kuroda |
| 453 | EPI_ISL_416589 | hCoV-19/Japan/DP0287/2020 | *Homo sapiens* | Asia / Japan / unknown | 2020-02-16 | Japanese Quarantine Stations | Pathogen Genomics Center, National Institute of Infectious Diseases | Tsuyoshi Sekizuka, Kentaro Itokawa, Rina Tanaka, Masanori Hashino, Tsutomu Kageyama, Shinji Saito, Ikuyo Takayama, Hideki Hasegawa, Takuri Takahashi, Hajime Kamiya, Takuya Yamagishi, Motoi Suzuki, Takaji Wakita, Makoto Kuroda |
| 454 | EPI_ISL_416591 | hCoV-19/Japan/DP0290/2020 | *Homo sapiens* | Asia / Japan / unknown | 2020-02-16 | Japanese Quarantine Stations | Pathogen Genomics Center, National Institute of Infectious Diseases | Tsuyoshi Sekizuka, Kentaro Itokawa, Rina Tanaka, Masanori Hashino, Tsutomu Kageyama, Shinji Saito, Ikuyo Takayama, Hideki Hasegawa, Takuri Takahashi, Hajime Kamiya, Takuya Yamagishi, Motoi Suzuki, Takaji Wakita, Makoto Kuroda |
| 455 | EPI_ISL_416592 | hCoV-19/Japan/DP0294/2020 | *Homo sapiens* | Asia / Japan / unknown | 2020-02-16 | Japanese Quarantine Stations | Pathogen Genomics Center, National Institute of Infectious Diseases | Tsuyoshi Sekizuka, Kentaro Itokawa, Rina Tanaka, Masanori Hashino, Tsutomu Kageyama, Shinji Saito, Ikuyo Takayama, Hideki Hasegawa, Takuri Takahashi, Hajime Kamiya, Takuya Yamagishi, Motoi Suzuki, Takaji Wakita, Makoto Kuroda |
| 456 | EPI_ISL_416593 | hCoV-19/Japan/DP0311/2020 | *Homo sapiens* | Asia / Japan / unknown | 2020-02-16 | Japanese Quarantine Stations | Pathogen Genomics Center, National Institute of Infectious Diseases | Tsuyoshi Sekizuka, Kentaro Itokawa, Rina Tanaka, Masanori Hashino, Tsutomu Kageyama, Shinji Saito, Ikuyo Takayama, Hideki Hasegawa, Takuri Takahashi, Hajime Kamiya, Takuya Yamagishi, Motoi Suzuki, Takaji Wakita, Makoto Kuroda |
| 457 | EPI_ISL_416594 | hCoV-19/Japan/DP0319/2020 | *Homo sapiens* | Asia / Japan / unknown | 2020-02-16 | Japanese Quarantine Stations | Pathogen Genomics Center, National Institute of Infectious Diseases | Tsuyoshi Sekizuka, Kentaro Itokawa, Rina Tanaka, Masanori Hashino, Tsutomu Kageyama, Shinji Saito, Ikuyo Takayama, Hideki Hasegawa, Takuri Takahashi, Hajime Kamiya, Takuya Yamagishi, Motoi Suzuki, Takaji Wakita, Makoto Kuroda |

| | | | | | | | | |
|---|---|---|---|---|---|---|---|---|
| 457 | EPI_ISL_416596 | hCoV-19/Japan/DP0344/2020 | *Homo sapiens* | Asia / Japan / unknown | 2020-02-16 | Japanese Quarantine Stations | Pathogen Genomics Center, National Institute of Infectious Diseases | Tsuyoshi Sekizuka, Kentaro Itokawa, Rina Tanaka, Masanori Hashino, Tsutomu Kageyama, Shinji Saito, Ikuyo Takayama, Hideki Hasegawa, Takuri Takahashi, Hajime Kamiya, Takuya Yamagishi, Motoi Suzuki, Takaji Wakita, Makoto Kuroda |
| 458 | EPI_ISL_416597 | hCoV-19/Japan/DP0346/2020 | *Homo sapiens* | Asia / Japan / unknown | 2020-02-16 | Japanese Quarantine Stations | Pathogen Genomics Center, National Institute of Infectious Diseases | Tsuyoshi Sekizuka, Kentaro Itokawa, Rina Tanaka, Masanori Hashino, Tsutomu Kageyama, Shinji Saito, Ikuyo Takayama, Hideki Hasegawa, Takuri Takahashi, Hajime Kamiya, Takuya Yamagishi, Motoi Suzuki, Takaji Wakita, Makoto Kuroda |
| 459 | EPI_ISL_416598 | hCoV-19/Japan/DP0357/2020 | *Homo sapiens* | Asia / Japan / unknown | 2020-02-16 | Japanese Quarantine Stations | Pathogen Genomics Center, National Institute of Infectious Diseases | Tsuyoshi Sekizuka, Kentaro Itokawa, Rina Tanaka, Masanori Hashino, Tsutomu Kageyama, Shinji Saito, Ikuyo Takayama, Hideki Hasegawa, Takuri Takahashi, Hajime Kamiya, Takuya Yamagishi, Motoi Suzuki, Takaji Wakita, Makoto Kuroda |
| 460 | EPI_ISL_416599 | hCoV-19/Japan/DP0361/2020 | *Homo sapiens* | Asia / Japan / unknown | 2020-02-16 | Japanese Quarantine Stations | Pathogen Genomics Center, National Institute of Infectious Diseases | Tsuyoshi Sekizuka, Kentaro Itokawa, Rina Tanaka, Masanori Hashino, Tsutomu Kageyama, Shinji Saito, Ikuyo Takayama, Hideki Hasegawa, Takuri Takahashi, Hajime Kamiya, Takuya Yamagishi, Motoi Suzuki, Takaji Wakita, Makoto Kuroda |
| 461 | EPI_ISL_416601 | hCoV-19/Japan/DP0457/2020 | *Homo sapiens* | Asia / Japan / unknown | 2020-02-16 | Japanese Quarantine Stations | Pathogen Genomics Center, National Institute of Infectious Diseases | Tsuyoshi Sekizuka, Kentaro Itokawa, Rina Tanaka, Masanori Hashino, Tsutomu Kageyama, Shinji Saito, Ikuyo Takayama, Hideki Hasegawa, Takuri Takahashi, Hajime Kamiya, Takuya Yamagishi, Motoi Suzuki, Takaji Wakita, Makoto Kuroda |
| 462 | EPI_ISL_416603 | hCoV-19/Japan/DP0464/2020 | *Homo sapiens* | Asia / Japan / unknown | 2020-02-16 | Japanese Quarantine Stations | Pathogen Genomics Center, National Institute of Infectious Diseases | Tsuyoshi Sekizuka, Kentaro Itokawa, Rina Tanaka, Masanori Hashino, Tsutomu Kageyama, Shinji Saito, Ikuyo Takayama, Hideki Hasegawa, Takuri Takahashi, Hajime Kamiya, Takuya Yamagishi, Motoi Suzuki, Takaji Wakita, Makoto Kuroda |
| 463 | EPI_ISL_416607 | hCoV-19/Japan/DP0543/2020 | *Homo sapiens* | Asia / Japan / unknown | 2020-02-17 | Japanese Quarantine Stations | Pathogen Genomics Center, National Institute of Infectious Diseases | Tsuyoshi Sekizuka, Kentaro Itokawa, Rina Tanaka, Masanori Hashino, Tsutomu Kageyama, Shinji Saito, Ikuyo Takayama, Hideki Hasegawa, Takuri Takahashi, Hajime Kamiya, Takuya Yamagishi, Motoi Suzuki, Takaji Wakita, Makoto Kuroda |
| 464 | EPI_ISL_416609 | hCoV-19/Japan/DP0568/2020 | *Homo sapiens* | Asia / Japan / unknown | 2020-02-17 | Japanese Quarantine Stations | Pathogen Genomics Center, National Institute of Infectious Diseases | Tsuyoshi Sekizuka, Kentaro Itokawa, Rina Tanaka, Masanori Hashino, Tsutomu Kageyama, Shinji Saito, Ikuyo Takayama, Hideki Hasegawa, Takuri Takahashi, Hajime Kamiya, Takuya Yamagishi, Motoi Suzuki, Takaji Wakita, Makoto Kuroda |
| 465 | EPI_ISL_416610 | hCoV-19/Japan/DP0588/2020 | *Homo sapiens* | Asia / Japan / unknown | 2020-02-17 | Japanese Quarantine Stations | Pathogen Genomics Center, National Institute of Infectious Diseases | Tsuyoshi Sekizuka, Kentaro Itokawa, Rina Tanaka, Masanori Hashino, Tsutomu Kageyama, Shinji Saito, Ikuyo Takayama, Hideki Hasegawa, Takuri Takahashi, Hajime Kamiya, Takuya Yamagishi, Motoi Suzuki, Takaji Wakita, Makoto Kuroda |
| 466 | EPI_ISL_416612 | hCoV-19/Japan/DP0645/2020 | *Homo sapiens* | Asia / Japan / unknown | 2020-02-17 | Japanese Quarantine Stations | Pathogen Genomics Center, National Institute of Infectious Diseases | Tsuyoshi Sekizuka, Kentaro Itokawa, Rina Tanaka, Masanori Hashino, Tsutomu Kageyama, Shinji Saito, Ikuyo Takayama, Hideki Hasegawa, Takuri Takahashi, Hajime Kamiya, Takuya Yamagishi, Motoi Suzuki, Takaji Wakita, Makoto Kuroda |
| 467 | EPI_ISL_416614 | hCoV-19/Japan/DP0687/2020 | *Homo sapiens* | Asia / Japan / unknown | 2020-02-17 | Japanese Quarantine Stations | Pathogen Genomics Center, National Institute of Infectious Diseases | Tsuyoshi Sekizuka, Kentaro Itokawa, Rina Tanaka, Masanori Hashino, Tsutomu Kageyama, Shinji Saito, Ikuyo Takayama, Hideki Hasegawa, Takuri Takahashi, Hajime Kamiya, Takuya Yamagishi, Motoi Suzuki, Takaji Wakita, Makoto Kuroda |
| 468 | EPI_ISL_416617 | hCoV-19/Japan/DP0699/2020 | *Homo sapiens* | Asia / Japan / unknown | 2020-02-17 | Japanese Quarantine Stations | Pathogen Genomics Center, National Institute of Infectious Diseases | Tsuyoshi Sekizuka, Kentaro Itokawa, Rina Tanaka, Masanori Hashino, Tsutomu Kageyama, Shinji Saito, Ikuyo Takayama, Hideki Hasegawa, Takuri Takahashi, Hajime Kamiya, Takuya Yamagishi, Motoi Suzuki, Takaji Wakita, Makoto Kuroda |
| 469 | EPI_ISL_416620 | hCoV-19/Japan/DP0724/2020 | *Homo sapiens* | Asia / Japan / unknown | 2020-02-17 | Japanese Quarantine Stations | Pathogen Genomics Center, National Institute of Infectious Diseases | Tsuyoshi Sekizuka, Kentaro Itokawa, Rina Tanaka, Masanori Hashino, Tsutomu Kageyama, Shinji Saito, Ikuyo Takayama, Hideki Hasegawa, Takuri Takahashi, Hajime Kamiya, Takuya Yamagishi, Motoi Suzuki, Takaji Wakita, Makoto Kuroda |
| 470 | EPI_ISL_416621 | hCoV-19/Japan/DP0743/2020 | *Homo sapiens* | Asia / Japan / unknown | 2020-02-17 | Japanese Quarantine Stations | Pathogen Genomics Center, National Institute of Infectious Diseases | Tsuyoshi Sekizuka, Kentaro Itokawa, Rina Tanaka, Masanori Hashino, Tsutomu Kageyama, Shinji Saito, Ikuyo Takayama, Hideki Hasegawa, Takuri Takahashi, Hajime Kamiya, Takuya Yamagishi, Motoi Suzuki, Takaji Wakita, Makoto Kuroda |

| | | | | | | | | |
|---|---|---|---|---|---|---|---|---|
| 471 | EPI_ISL_416622 | hCoV-19/Japan/DP0752/2020 | *Homo sapiens* | Asia / Japan / unknown | 2020-02-17 | Japanese Quarantine Stations | Pathogen Genomics Center, National Institute of Infectious Diseases | Tsuyoshi Sekizuka, Kentaro Itokawa, Rina Tanaka, Masanori Hashino, Tsutomu Kageyama, Shinji Saito, Ikuyo Takayama, Hideki Hasegawa, Takuri Takahashi, Hajime Kamiya, Takuya Yamagishi, Motoi Suzuki, Takaji Wakita, Makoto Kuroda |
| 472 | EPI_ISL_416624 | hCoV-19/Japan/DP0764/2020 | *Homo sapiens* | Asia / Japan / unknown | 2020-02-17 | Japanese Quarantine Stations | Pathogen Genomics Center, National Institute of Infectious Diseases | Tsuyoshi Sekizuka, Kentaro Itokawa, Rina Tanaka, Masanori Hashino, Tsutomu Kageyama, Shinji Saito, Ikuyo Takayama, Hideki Hasegawa, Takuri Takahashi, Hajime Kamiya, Takuya Yamagishi, Motoi Suzuki, Takaji Wakita, Makoto Kuroda |
| 473 | EPI_ISL_416625 | hCoV-19/Japan/DP0765/2020 | *Homo sapiens* | Asia / Japan / unknown | 2020-02-17 | Japanese Quarantine Stations | Pathogen Genomics Center, National Institute of Infectious Diseases | Tsuyoshi Sekizuka, Kentaro Itokawa, Rina Tanaka, Masanori Hashino, Tsutomu Kageyama, Shinji Saito, Ikuyo Takayama, Hideki Hasegawa, Takuri Takahashi, Hajime Kamiya, Takuya Yamagishi, Motoi Suzuki, Takaji Wakita, Makoto Kuroda |
| 474 | EPI_ISL_416628 | hCoV-19/Japan/DP0786/2020 | *Homo sapiens* | Asia / Japan / unknown | 2020-02-17 | Japanese Quarantine Stations | Pathogen Genomics Center, National Institute of Infectious Diseases | Tsuyoshi Sekizuka, Kentaro Itokawa, Rina Tanaka, Masanori Hashino, Tsutomu Kageyama, Shinji Saito, Ikuyo Takayama, Hideki Hasegawa, Takuri Takahashi, Hajime Kamiya, Takuya Yamagishi, Motoi Suzuki, Takaji Wakita, Makoto Kuroda |
| 475 | EPI_ISL_416631 | hCoV-19/Japan/DP0804/2020 | *Homo sapiens* | Asia / Japan / unknown | 2020-02-17 | Japanese Quarantine Stations | Pathogen Genomics Center, National Institute of Infectious Diseases | Tsuyoshi Sekizuka, Kentaro Itokawa, Rina Tanaka, Masanori Hashino, Tsutomu Kageyama, Shinji Saito, Ikuyo Takayama, Hideki Hasegawa, Takuri Takahashi, Hajime Kamiya, Takuya Yamagishi, Motoi Suzuki, Takaji Wakita, Makoto Kuroda |
| 476 | EPI_ISL_416632 | hCoV-19/Japan/DP0827/2020 | *Homo sapiens* | Asia / Japan / unknown | 2020-02-17 | Japanese Quarantine Stations | Pathogen Genomics Center, National Institute of Infectious Diseases | Tsuyoshi Sekizuka, Kentaro Itokawa, Rina Tanaka, Masanori Hashino, Tsutomu Kageyama, Shinji Saito, Ikuyo Takayama, Hideki Hasegawa, Takuri Takahashi, Hajime Kamiya, Takuya Yamagishi, Motoi Suzuki, Takaji Wakita, Makoto Kuroda |
| 477 | EPI_ISL_416633 | hCoV-19/Japan/DP0880/2020 | *Homo sapiens* | Asia / Japan / unknown | 2020-02-17 | Japanese Quarantine Stations | Pathogen Genomics Center, National Institute of Infectious Diseases | Tsuyoshi Sekizuka, Kentaro Itokawa, Rina Tanaka, Masanori Hashino, Tsutomu Kageyama, Shinji Saito, Ikuyo Takayama, Hideki Hasegawa, Takuri Takahashi, Hajime Kamiya, Takuya Yamagishi, Motoi Suzuki, Takaji Wakita, Makoto Kuroda |
| 478 | EPI_ISL_416635 | hCoV-19/USA/WA-UW97/2020 | *Homo sapiens* | North America / USA / Washington | 2020-03-12 | UW Virology Lab | UW Virology Lab | Pavitra Roychoudhury, Hong Xie, Keith Jerome, Alexander Greninger |
| 479 | EPI_ISL_416636 | hCoV-19/USA/WA-UW98/2020 | *Homo sapiens* | North America / USA / Washington | 2020-03-12 | UW Virology Lab | UW Virology Lab | Pavitra Roychoudhury, Hong Xie, Keith Jerome, Alexander Greninger |
| 480 | EPI_ISL_416637 | hCoV-19/USA/WA-UW99/2020 | *Homo sapiens* | North America / USA / Washington | 2020-03-12 | UW Virology Lab | UW Virology Lab | Pavitra Roychoudhury, Hong Xie, Keith Jerome, Alexander Greninger |
| 480 | EPI_ISL_416659 | hCoV-19/USA/WA-UW121/2020 | *Homo sapiens* | North America / USA / Washington | 2020-03-11 | UW Virology Lab | UW Virology Lab | Pavitra Roychoudhury, Hong Xie, Keith Jerome, Alexander Greninger |
| 480 | EPI_ISL_416685 | hCoV-19/USA/WA-UW147/2020 | *Homo sapiens* | North America / USA / Washington | 2020-03-15 | UW Virology Lab | UW Virology Lab | Pavitra Roychoudhury, Hong Xie, Keith Jerome, Alexander Greninger |
| 481 | EPI_ISL_416638 | hCoV-19/USA/WA-UW100/2020 | *Homo sapiens* | North America / USA / Washington | 2020-03-12 | UW Virology Lab | UW Virology Lab | Pavitra Roychoudhury, Hong Xie, Keith Jerome, Alexander Greninger |
| 482 | EPI_ISL_416641 | hCoV-19/USA/WA-UW103/2020 | *Homo sapiens* | North America / USA / Washington | 2020-03-11 | UW Virology Lab | UW Virology Lab | Pavitra Roychoudhury, Hong Xie, Keith Jerome, Alexander Greninger |
| 483 | EPI_ISL_416642 | hCoV-19/USA/WA-UW104/2020 | *Homo sapiens* | North America / USA / Washington | 2020-03-11 | UW Virology Lab | UW Virology Lab | Pavitra Roychoudhury, Hong Xie, Keith Jerome, Alexander Greninger |
| 484 | EPI_ISL_416644 | hCoV-19/USA/WA-UW106/2020 | *Homo sapiens* | North America / USA / Washington | 2020-03-11 | UW Virology Lab | UW Virology Lab | Pavitra Roychoudhury, Hong Xie, Keith Jerome, Alexander Greninger |
| 485 | EPI_ISL_416646 | hCoV-19/USA/WA-UW108/2020 | *Homo sapiens* | North America / USA / Washington | 2020-03-11 | UW Virology Lab | UW Virology Lab | Pavitra Roychoudhury, Hong Xie, Keith Jerome, Alexander Greninger |

| | | | | | | | | |
|---|---|---|---|---|---|---|---|---|
| 486 | EPI_ISL_416648 | hCoV-19/USA/WA-UW110/2020 | *Homo sapiens* | North America / USA / Washington | 2020-03-11 | UW Virology Lab | UW Virology Lab | Pavitra Roychoudhury, Hong Xie, Keith Jerome, Alexander Greninger |
| 487 | EPI_ISL_416649 | hCoV-19/USA/WA-UW111/2020 | *Homo sapiens* | North America / USA / Washington | 2020-03-11 | UW Virology Lab | UW Virology Lab | Pavitra Roychoudhury, Hong Xie, Keith Jerome, Alexander Greninger |
| 488 | EPI_ISL_416650 | hCoV-19/USA/WA-UW112/2020 | *Homo sapiens* | North America / USA | 2020-03-10 | UW Virology Lab | UW Virology Lab | Pavitra Roychoudhury, Hong Xie, Keith Jerome, Alexander Greninger |
| 489 | EPI_ISL_416651 | hCoV-19/USA/WA-UW113/2020 | *Homo sapiens* | North America / USA | 2020-03-11 | UW Virology Lab | UW Virology Lab | Pavitra Roychoudhury, Hong Xie, Keith Jerome, Alexander Greninger |
| 490 | EPI_ISL_416652 | hCoV-19/USA/WA-UW114/2020 | *Homo sapiens* | North America / USA / Washington | 2020-03-11 | UW Virology Lab | UW Virology Lab | Pavitra Roychoudhury, Hong Xie, Keith Jerome, Alexander Greninger |
| 491 | EPI_ISL_416653 | hCoV-19/USA/WA-UW115/2020 | *Homo sapiens* | North America / USA / Washington | 2020-03-11 | UW Virology Lab | UW Virology Lab | Pavitra Roychoudhury, Hong Xie, Keith Jerome, Alexander Greninger |
| 492 | EPI_ISL_416655 | hCoV-19/USA/WA-UW117/2020 | *Homo sapiens* | North America / USA / Washington | 2020-03-11 | UW Virology Lab | UW Virology Lab | Pavitra Roychoudhury, Hong Xie, Keith Jerome, Alexander Greninger |
| 493 | EPI_ISL_416656 | hCoV-19/USA/WA-UW118/2020 | *Homo sapiens* | North America / USA / Washington | 2020-03-11 | UW Virology Lab | UW Virology Lab | Pavitra Roychoudhury, Hong Xie, Keith Jerome, Alexander Greninger |
| 494 | EPI_ISL_416657 | hCoV-19/USA/WA-UW119/2020 | *Homo sapiens* | North America / USA / Washington | 2020-03-11 | UW Virology Lab | UW Virology Lab | Pavitra Roychoudhury, Hong Xie, Keith Jerome, Alexander Greninger |
| 495 | EPI_ISL_416660 | hCoV-19/USA/WA-UW122/2020 | *Homo sapiens* | North America / USA | 2020-03-10 | UW Virology Lab | UW Virology Lab | Pavitra Roychoudhury, Hong Xie, Keith Jerome, Alexander Greninger |
| 496 | EPI_ISL_416661 | hCoV-19/USA/WA-UW123/2020 | *Homo sapiens* | North America / USA / Washington | 2020-03-12 | UW Virology Lab | UW Virology Lab | Pavitra Roychoudhury, Hong Xie, Keith Jerome, Alexander Greninger |
| 497 | EPI_ISL_416662 | hCoV-19/USA/WA-UW124/2020 | *Homo sapiens* | North America / USA / Washington | 2020-03-12 | UW Virology Lab | UW Virology Lab | Pavitra Roychoudhury, Hong Xie, Keith Jerome, Alexander Greninger |
| 498 | EPI_ISL_416663 | hCoV-19/USA/WA-UW125/2020 | *Homo sapiens* | North America / USA / Washington | 2020-03-12 | UW Virology Lab | UW Virology Lab | Pavitra Roychoudhury, Hong Xie, Keith Jerome, Alexander Greninger |
| 499 | EPI_ISL_416664 | hCoV-19/USA/WA-UW126/2020 | *Homo sapiens* | North America / USA / Washington | 2020-03-12 | UW Virology Lab | UW Virology Lab | Pavitra Roychoudhury, Hong Xie, Keith Jerome, Alexander Greninger |
| 500 | EPI_ISL_416665 | hCoV-19/USA/WA-UW127/2020 | *Homo sapiens* | North America / USA / Washington | 2020-03-12 | UW Virology Lab | UW Virology Lab | Pavitra Roychoudhury, Hong Xie, Keith Jerome, Alexander Greninger |
| 501 | EPI_ISL_416666 | hCoV-19/USA/WA-UW128/2020 | *Homo sapiens* | North America / USA / Washington | 2020-03-12 | UW Virology Lab | UW Virology Lab | Pavitra Roychoudhury, Hong Xie, Keith Jerome, Alexander Greninger |
| 502 | EPI_ISL_416667 | hCoV-19/USA/WA-UW129/2020 | *Homo sapiens* | North America / USA / Washington | 2020-03-12 | UW Virology Lab | UW Virology Lab | Pavitra Roychoudhury, Hong Xie, Keith Jerome, Alexander Greninger |
| 503 | EPI_ISL_416668 | hCoV-19/USA/WA-UW130/2020 | *Homo sapiens* | North America / USA / Washington | 2020-03-11 | UW Virology Lab | UW Virology Lab | Pavitra Roychoudhury, Hong Xie, Keith Jerome, Alexander Greninger |
| 504 | EPI_ISL_416669 | hCoV-19/USA/WA-UW131/2020 | *Homo sapiens* | North America / USA / Washington | 2020-03-11 | UW Virology Lab | UW Virology Lab | Pavitra Roychoudhury, Hong Xie, Keith Jerome, Alexander Greninger |
| 505 | EPI_ISL_416672 | hCoV-19/USA/WA-UW134/2020 | *Homo sapiens* | North America / USA | 2020-03-10 | UW Virology Lab | UW Virology Lab | Pavitra Roychoudhury, Hong Xie, Keith Jerome, Alexander Greninger |
| 506 | EPI_ISL_416674 | hCoV-19/USA/WA-UW136/2020 | *Homo sapiens* | North America / USA | 2020-03-11 | UW Virology Lab | UW Virology Lab | Pavitra Roychoudhury, Hong Xie, Keith Jerome, Alexander Greninger |

| | | | | | | | | |
|---|---|---|---|---|---|---|---|---|
| 507 | EPI_ISL_416675 | hCoV-19/USA/WA-UW137/2020 | *Homo sapiens* | North America / USA | 2020-03-11 | UW Virology Lab | UW Virology Lab | Pavitra Roychoudhury, Hong Xie, Keith Jerome, Alexander Greninger |
| 508 | EPI_ISL_416676 | hCoV-19/USA/WA-UW138/2020 | *Homo sapiens* | North America / USA | 2020-03-11 | UW Virology Lab | UW Virology Lab | Pavitra Roychoudhury, Hong Xie, Keith Jerome, Alexander Greninger |
| 509 | EPI_ISL_416677 | hCoV-19/USA/WA-UW139/2020 | *Homo sapiens* | North America / USA / Washington | 2020-03-12 | UW Virology Lab | UW Virology Lab | Pavitra Roychoudhury, Hong Xie, Keith Jerome, Alexander Greninger |
| 510 | EPI_ISL_416678 | hCoV-19/USA/WA-UW140/2020 | *Homo sapiens* | North America / USA / Washington | 2020-03-11 | UW Virology Lab | UW Virology Lab | Pavitra Roychoudhury, Hong Xie, Keith Jerome, Alexander Greninger |
| 511 | EPI_ISL_416679 | hCoV-19/USA/WA-UW141/2020 | *Homo sapiens* | North America / USA / Washington | 2020-03-11 | UW Virology Lab | UW Virology Lab | Pavitra Roychoudhury, Hong Xie, Keith Jerome, Alexander Greninger |
| 512 | EPI_ISL_416680 | hCoV-19/USA/WA-UW142/2020 | *Homo sapiens* | North America / USA / Washington | 2020-03-11 | UW Virology Lab | UW Virology Lab | Pavitra Roychoudhury, Hong Xie, Keith Jerome, Alexander Greninger |
| 513 | EPI_ISL_416683 | hCoV-19/USA/WA-UW145/2020 | *Homo sapiens* | North America / USA / Washington | 2020-03-15 | UW Virology Lab | UW Virology Lab | Pavitra Roychoudhury, Hong Xie, Keith Jerome, Alexander Greninger |
| 514 | EPI_ISL_416686 | hCoV-19/USA/WA-UW148/2020 | *Homo sapiens* | North America / USA / Washington | 2020-03-14 | UW Virology Lab | UW Virology Lab | Pavitra Roychoudhury, Hong Xie, Keith Jerome, Alexander Greninger |
| 515 | EPI_ISL_416687 | hCoV-19/USA/WA-UW149/2020 | *Homo sapiens* | North America / USA / Washington | 2020-03-14 | UW Virology Lab | UW Virology Lab | Pavitra Roychoudhury, Hong Xie, Keith Jerome, Alexander Greninger |
| 516 | EPI_ISL_416688 | hCoV-19/USA/WA-UW150/2020 | *Homo sapiens* | North America / USA / Washington | 2020-03-14 | UW Virology Lab | UW Virology Lab | Pavitra Roychoudhury, Hong Xie, Keith Jerome, Alexander Greninger |
| 516 | EPI_ISL_416705 | hCoV-19/USA/WA-UW167/2020 | *Homo sapiens* | North America / USA / Washington | 2020-03-13 | UW Virology Lab | UW Virology Lab | Pavitra Roychoudhury, Hong Xie, Keith Jerome, Alexander Greninger |
| 517 | EPI_ISL_416690 | hCoV-19/USA/WA-UW152/2020 | *Homo sapiens* | North America / USA / Washington | 2020-03-13 | UW Virology Lab | UW Virology Lab | Pavitra Roychoudhury, Hong Xie, Keith Jerome, Alexander Greninger |
| 518 | EPI_ISL_416691 | hCoV-19/USA/WA-UW153/2020 | *Homo sapiens* | North America / USA / Washington | 2020-03-13 | UW Virology Lab | UW Virology Lab | Pavitra Roychoudhury, Hong Xie, Keith Jerome, Alexander Greninger |
| 519 | EPI_ISL_416693 | hCoV-19/USA/WA-UW155/2020 | *Homo sapiens* | North America / USA / Washington | 2020-03-12 | UW Virology Lab | UW Virology Lab | Pavitra Roychoudhury, Hong Xie, Keith Jerome, Alexander Greninger |
| 520 | EPI_ISL_416694 | hCoV-19/USA/WA-UW156/2020 | *Homo sapiens* | North America / USA / Washington | 2020-03-13 | UW Virology Lab | UW Virology Lab | Pavitra Roychoudhury, Hong Xie, Keith Jerome, Alexander Greninger |
| 521 | EPI_ISL_416696 | hCoV-19/USA/WA-UW158/2020 | *Homo sapiens* | North America / USA / Washington | 2020-03-13 | UW Virology Lab | UW Virology Lab | Pavitra Roychoudhury, Hong Xie, Keith Jerome, Alexander Greninger |
| 522 | EPI_ISL_416697 | hCoV-19/USA/WA-UW159/2020 | *Homo sapiens* | North America / USA / Washington | 2020-03-13 | UW Virology Lab | UW Virology Lab | Pavitra Roychoudhury, Hong Xie, Keith Jerome, Alexander Greninger |
| 523 | EPI_ISL_416698 | hCoV-19/USA/WA-UW160/2020 | *Homo sapiens* | North America / USA / Washington | 2020-03-13 | UW Virology Lab | UW Virology Lab | Pavitra Roychoudhury, Hong Xie, Keith Jerome, Alexander Greninger |
| 524 | EPI_ISL_416700 | hCoV-19/USA/WA-UW162/2020 | *Homo sapiens* | North America / USA / Washington | 2020-03-13 | UW Virology Lab | UW Virology Lab | Pavitra Roychoudhury, Hong Xie, Keith Jerome, Alexander Greninger |
| 525 | EPI_ISL_416703 | hCoV-19/USA/WA-UW165/2020 | *Homo sapiens* | North America / USA / Washington | 2020-03-13 | UW Virology Lab | UW Virology Lab | Pavitra Roychoudhury, Hong Xie, Keith Jerome, Alexander Greninger |

| | | | | | | | | |
|---|---|---|---|---|---|---|---|---|
| 526 | EPI_ISL_416706 | hCoV-19/USA/WA-UW168/2020 | *Homo sapiens* | North America / USA / Washington | 2020-03-13 | UW Virology Lab | UW Virology Lab | Pavitra Roychoudhury, Hong Xie, Keith Jerome, Alexander Greninger |
| 527 | EPI_ISL_416707 | hCoV-19/USA/WA-UW169/2020 | *Homo sapiens* | North America / USA / Washington | 2020-03-14 | UW Virology Lab | UW Virology Lab | Pavitra Roychoudhury, Hong Xie, Keith Jerome, Alexander Greninger |
| 528 | EPI_ISL_416709 | hCoV-19/USA/WA-UW171/2020 | *Homo sapiens* | North America / USA / Washington | 2020-03-13 | UW Virology Lab | UW Virology Lab | Pavitra Roychoudhury, Hong Xie, Keith Jerome, Alexander Greninger |
| 529 | EPI_ISL_416710 | hCoV-19/USA/WA-UW172/2020 | *Homo sapiens* | North America / USA / Washington | 2020-03-13 | UW Virology Lab | UW Virology Lab | Pavitra Roychoudhury, Hong Xie, Keith Jerome, Alexander Greninger |
| 530 | EPI_ISL_416711 | hCoV-19/USA/WA-UW173/2020 | *Homo sapiens* | North America / USA / Washington | 2020-03-15 | UW Virology Lab | UW Virology Lab | Pavitra Roychoudhury, Hong Xie, Keith Jerome, Alexander Greninger |
| 531 | EPI_ISL_416713 | hCoV-19/USA/WA-UW175/2020 | *Homo sapiens* | North America / USA / Washington | 2020-03-15 | UW Virology Lab | UW Virology Lab | Pavitra Roychoudhury, Hong Xie, Keith Jerome, Alexander Greninger |
| 532 | EPI_ISL_416714 | hCoV-19/USA/WA-UW176/2020 | *Homo sapiens* | North America / USA / Washington | 2020-03-14 | UW Virology Lab | UW Virology Lab | Pavitra Roychoudhury, Hong Xie, Keith Jerome, Alexander Greninger |
| 533 | EPI_ISL_416715 | hCoV-19/USA/WA-UW177/2020 | *Homo sapiens* | North America / USA / Washington | 2020-03-15 | UW Virology Lab | UW Virology Lab | Pavitra Roychoudhury, Hong Xie, Keith Jerome, Alexander Greninger |
| 534 | EPI_ISL_416716 | hCoV-19/USA/WA-UW178/2020 | *Homo sapiens* | North America / USA / Washington | 2020-03-13 | UW Virology Lab | UW Virology Lab | Pavitra Roychoudhury, Hong Xie, Keith Jerome, Alexander Greninger |
| 535 | EPI_ISL_416717 | hCoV-19/USA/WA-UW179/2020 | *Homo sapiens* | North America / USA / Washington | 2020-03-15 | UW Virology Lab | UW Virology Lab | Pavitra Roychoudhury, Hong Xie, Keith Jerome, Alexander Greninger |
| 536 | EPI_ISL_416718 | hCoV-19/USA/WA-UW180/2020 | *Homo sapiens* | North America / USA / Washington | 2020-03-14 | UW Virology Lab | UW Virology Lab | Pavitra Roychoudhury, Hong Xie, Keith Jerome, Alexander Greninger |
| 537 | EPI_ISL_416721 | hCoV-19/USA/WA-UW183/2020 | *Homo sapiens* | North America / USA / Washington | 2020-03-13 | UW Virology Lab | UW Virology Lab | Pavitra Roychoudhury, Hong Xie, Keith Jerome, Alexander Greninger |
| 538 | EPI_ISL_416723 | hCoV-19/USA/WA-UW185/2020 | *Homo sapiens* | North America / USA / Washington | 2020-03-14 | UW Virology Lab | UW Virology Lab | Pavitra Roychoudhury, Hong Xie, Keith Jerome, Alexander Greninger |
| 539 | EPI_ISL_416724 | hCoV-19/USA/WA-UW186/2020 | *Homo sapiens* | North America / USA / Washington | 2020-03-13 | UW Virology Lab | UW Virology Lab | Pavitra Roychoudhury, Hong Xie, Keith Jerome, Alexander Greninger |
| 540 | EPI_ISL_416725 | hCoV-19/USA/WA-UW187/2020 | *Homo sapiens* | North America / USA / Washington | 2020-03-13 | UW Virology Lab | UW Virology Lab | Pavitra Roychoudhury, Hong Xie, Keith Jerome, Alexander Greninger |
| 541 | EPI_ISL_416726 | hCoV-19/USA/WA-UW188/2020 | *Homo sapiens* | North America / USA / Washington | 2020-03-13 | UW Virology Lab | UW Virology Lab | Pavitra Roychoudhury, Hong Xie, Keith Jerome, Alexander Greninger |
| 542 | EPI_ISL_416727 | hCoV-19/USA/WA-UW189/2020 | *Homo sapiens* | North America / USA / Washington | 2020-03-13 | UW Virology Lab | UW Virology Lab | Pavitra Roychoudhury, Hong Xie, Keith Jerome, Alexander Greninger |
| 543 | EPI_ISL_416728 | hCoV-19/USA/WA-UW190/2020 | *Homo sapiens* | North America / USA / Washington | 2020-03-13 | UW Virology Lab | UW Virology Lab | Pavitra Roychoudhury, Hong Xie, Keith Jerome, Alexander Greninger |
| 544 | EPI_ISL_416729 | hCoV-19/USA/WA-UW191/2020 | *Homo sapiens* | North America / USA / Washington | 2020-03-13 | UW Virology Lab | UW Virology Lab | Pavitra Roychoudhury, Hong Xie, Keith Jerome, Alexander Greninger |

| | | | | | | | | |
|---|---|---|---|---|---|---|---|---|
| 545 | EPI_ISL_416730 | hCoV-19/England/SHEF-BFCB1/2020 | *Homo sapiens* | Europe / England / South Yorkshire | 2020-03-03 | Virology Department, Sheffield Teaching Hospitals NHS Foundation Trust | Department of Infection, Immunity and Cardiovascular Disease, The Florey Institute, The Medical School, University of Sheffield | Thushan de Silva, Matthew Parker, Adri Angyal, Rebecca Brown, Matthew Wyles, Mehmet Yavuz, Mohammad Raza, Cariad Evans |
| 546 | EPI_ISL_416731 | hCoV-19/England/SHEF-BFCC0/2020 | *Homo sapiens* | Europe / England / Northamtonshire | 2020-03-03 | Virology Department, Sheffield Teaching Hospitals NHS Foundation Trust | Department of Infection, Immunity and Cardiovascular Disease, The Florey Institute, The Medical School, University of Sheffield | Thushan de Silva, Matthew Parker, Adri Angyal, Rebecca Brown, Matthew Wyles, Mehmet Yavuz, Mohammad Raza, Cariad Evans |
| 547 | EPI_ISL_416733 | hCoV-19/England/SHEF-BFCEE/2020 | *Homo sapiens* | Europe / England / South Yorkshire | 2020-03-07 | Virology Department, Sheffield Teaching Hospitals NHS Foundation Trust | Department of Infection, Immunity and Cardiovascular Disease, The Florey Institute, The Medical School, University of Sheffield | Thushan de Silva, Matthew Parker, Adri Angyal, Rebecca Brown, Matthew Wyles, Mehmet Yavuz, Mohammad Raza, Cariad Evans |
| 548 | EPI_ISL_416740 | hCoV-19/England/SHEF-BFD54/2020 | *Homo sapiens* | Europe / England / South Yorkshire | 2020-03-03 | Virology Department, Sheffield Teaching Hospitals NHS Foundation Trust | Department of Infection, Immunity and Cardiovascular Disease, The Florey Institute, The Medical School, University of Sheffield | Thushan de Silva, Matthew Parker, Adri Angyal, Rebecca Brown, Matthew Wyles, Mehmet Yavuz, Mohammad Raza, Cariad Evans |
| 549 | EPI_ISL_416741 | hCoV-19/Lithuania/ChVir1632/2020 | *Homo sapiens* | Europe / Lithuania / Vilnius | 2020-02 | National Public Health Surveillance Laboratory, Vilnius, Lithuania | Charite Universitaetsmedizin Berlin, Institute of Virology | Victor M Corman, Julia Schneider, Jörn Beheim-Schwarzbach, Talitha Veith, Barbara Muehlemann, Terry Jones, Ana Steponkiene, Christian Drosten |
| 550 | EPI_ISL_416744 | hCoV-19/Hungary/49/2020 | *Homo sapiens* | Europe / Hungary / Baranya | 2020-03-20 | Virological Research Group, Szentágothai Research Centre | Bioinformatics Research Group, Szentágothai Research Centre | Péter Urbán, Endre Gábor Tóth, Gábor Kemenesi, Róbert Herczeg, Attila Gyenesei, Ferenc Jakab |
| 551 | EPI_ISL_416746 | hCoV-19/France/Valence_425/2020 | *Homo sapiens* | Europe / France / ARA | 2020-03-03 | CNR Virus des Infections Respiratoires - France SUD | CNR Virus des Infections Respiratoires - France SUD | Bal, Antonin; Destras, Gregory; Gaymard, Alexandre; Bouscambert-Duchamp, Maude; Cheynet, Valérie; Brengel-Pesce, Karen; Morfin-Sherpa, Florence; Valette, Martine; Josset, Laurence; Lina, Bruno. |
| 552 | EPI_ISL_416749 | hCoV-19/France/Valence_532/2020 | *Homo sapiens* | Europe / France / ARA | 2020-03-04 | Centre Hospitalier de Valence | CNR Virus des Infections Respiratoires - France SUD | Bal, Antonin; Destras, Gregory; Gaymard, Alexandre; Bouscambert-Duchamp, Maude; Cheynet, Valérie; Brengel-Pesce, Karen; Morfin-Sherpa, Florence; Valette, Martine; Josset, Laurence; Lina, Bruno. |
| 553 | EPI_ISL_416751 | hCoV-19/France/Clermont-Ferrand_651/2020 | *Homo sapiens* | Europe / France / ARA | 2020-03-05 | CHU Gabriel Montpied | CNR Virus des Infections Respiratoires - France SUD | Bal, Antonin; Destras, Gregory; Gaymard, Alexandre; Bouscambert-Duchamp, Maude; Cheynet, Valérie; Brengel-Pesce, Karen; Morfin-Sherpa, Florence; Valette, Martine; Josset, Laurence; Lina, Bruno. |
| 554 | EPI_ISL_416829 | hCoV-19/Malaysia/MKAK-CL-2020-5045/2020 | *Homo sapiens* | Asia / Malaysia / Selangor | 2020-02-20 | National Public Health Laboratory | Malaysia Genome Institute | Mohd Noor Mat Isa, Irni Suhayu Sapian, Yusuf Muhammd Noor, Hezreen Iqbal, Mohd Faizal Abu Bakar, Enizza Kassim, Shamsidar Sopie, Azrin Ahmad, Norazfa Johari, Norazimah Tajudin, Selvanesan Sengol, Yu Kie Chem, Hani Mat Hussin, Shahrul Hisham Zainal Ariffin |
| 555 | EPI_ISL_416830 | hCoV-19/USA/NY-NYUMC2/2020 | *Homo sapiens* | North America / USA / New York City | 2020-03-16 | NYU Langone Health | Department of Pathology and Medicine, New York University School of Medicine | John Chen, Dacia Dimartino, Xiaojun Feng, Adriana Heguy, Megan Hogan, Emily Huang, George Jour, Christian Marier, Matt Maurano, Mark Mulligan, Peter Meyn, Marie Samanovic-Golden, Amy Rapkiewicz, Guomiao Shen, Matija Snuderl, Gael Westby, Paul Zappile |
| 556 | EPI_ISL_416831 | hCoV-19/USA/NY-NYUMC3/2020 | *Homo sapiens* | North America / USA / New York City | 2020-03-16 | NYU Langone Health | Department of Pathology and Medicine, New York University School of Medicine | John Chen, Dacia Dimartino, Xiaojun Feng, Adriana Heguy, Megan Hogan, Emily Huang, George Jour, Christian Marier, Matt Maurano, Mark Mulligan, Peter Meyn, Marie Samanovic-Golden, Amy Rapkiewicz, Guomiao Shen, Matija Snuderl, Gael Westby, Paul Zappile |
| 557 | EPI_ISL_416866 | hCoV-19/Malaysia/MKAK-CL-2020-5047/2020 | *Homo sapiens* | Asia/ Malaysia/Selangor | 2020-02-20 | National Public Health Laboratory | Malaysia Genome Institute | Mohd Noor Mat Isa, Irni Suhayu Sapian, Yusuf Muhammd Noor, Hezreen Iqbal, Mohd Faizal Abu Bakar, Enizza Kassim, Shamsidar Sopie, Azrin Ahmad, Norazfa Johari, Norazimah Tajudin, Selvanesan Sengol, Yu Kie Chem, Hani Mat Hussin, Shahrul Hisham Zainal Ariffin |
| 558 | EPI_ISL_417004 | hCoV-19/Belgium/ULG-3000/2020 | *Homo sapiens* | Europe / Belgium / Liège | 2020-03-05 | Department of Clinical Microbiology | GIGA Medical Genomics | Durkin Keith, Artesi Maria, Bontems Sébastien, Boreux Raphaël, Meex Cécile, Melin Pierrette, Hayette Marie-Pierre, Bours Vincent. |
| 559 | EPI_ISL_417014 | hCoV-19/Belgium/ULG-6216/2020 | *Homo sapiens* | Europe / Belgium / Liège | 2020-03-13 | Department of Clinical Microbiology | GIGA Medical Genomics | Durkin Keith, Artesi Maria, Bontems Sébastien, Boreux Raphaël, Meex Cécile, Melin Pierrette, Hayette Marie-Pierre, Bours Vincent. |

| | | | | | | | | |
|---|---|---|---|---|---|---|---|---|
| 560 | EPI_ISL_417015 | hCoV-19/Belgium/ULG-6457/2020 | *Homo sapiens* | Europe / Belgium / Liège | 2020-03-13 | Department of Clinical Microbiology | GIGA Medical Genomics | Durkin Keith, Artesi Maria, Bontems Sébastien, Boreux Raphaël, Meex Cécile, Melin Pierrette, Hayette Marie-Pierre, Bours Vincent. |
| 561 | EPI_ISL_417017 | hCoV-19/Belgium/ULG-6638/2020 | *Homo sapiens* | Europe / Belgium / Liège | 2020-03-14 | Department of Clinical Microbiology | GIGA Medical Genomics | Durkin Keith, Artesi Maria, Bontems Sébastien, Boreux Raphaël, Meex Cécile, Melin Pierrette, Hayette Marie-Pierre, Bours Vincent. |
| 562 | EPI_ISL_417018 | hCoV-19/Belgium/ULG-6670/2020 | *Homo sapiens* | Europe / Belgium / Liège | 2020-03-14 | Department of Clinical Microbiology | GIGA Medical Genomics | Durkin Keith, Artesi Maria, Bontems Sébastien, Boreux Raphaël, Meex Cécile, Melin Pierrette, Hayette Marie-Pierre, Bours Vincent. |
| 563 | EPI_ISL_417019 | hCoV-19/Belgium/ULG-6754/2020 | *Homo sapiens* | Europe / Belgium / Liège | 2020-03-14 | Department of Clinical Microbiology | GIGA Medical Genomics | Durkin Keith, Artesi Maria, Bontems Sébastien, Boreux Raphaël, Meex Cécile, Melin Pierrette, Hayette Marie-Pierre, Bours Vincent. |
| 564 | EPI_ISL_417020 | hCoV-19/Belgium/ULG-6939/2020 | *Homo sapiens* | Europe / Belgium / Liège | 2020-03-15 | Department of Clinical Microbiology | GIGA Medical Genomics | Durkin Keith, Artesi Maria, Bontems Sébastien, Boreux Raphaël, Meex Cécile, Melin Pierrette, Hayette Marie-Pierre, Bours Vincent. |
| 565 | EPI_ISL_417021 | hCoV-19/Belgium/ULG-6942/2020 | *Homo sapiens* | Europe / Belgium / Liège | 2020-03-15 | Department of Clinical Microbiology | GIGA Medical Genomics | Durkin Keith, Artesi Maria, Bontems Sébastien, Boreux Raphaël, Meex Cécile, Melin Pierrette, Hayette Marie-Pierre, Bours Vincent. |
| 565 | EPI_ISL_417022 | hCoV-19/Belgium/ULG-6948/2020 | *Homo sapiens* | Europe / Belgium / Liège | 2020-03-15 | Department of Clinical Microbiology | GIGA Medical Genomics | Durkin Keith, Artesi Maria, Bontems Sébastien, Boreux Raphaël, Meex Cécile, Melin Pierrette, Hayette Marie-Pierre, Bours Vincent. |
| 565 | EPI_ISL_417023 | hCoV-19/Belgium/ULG-6950/2020 | *Homo sapiens* | Europe / Belgium / Liège | 2020-03-15 | Department of Clinical Microbiology | GIGA Medical Genomics | Durkin Keith, Artesi Maria, Bontems Sébastien, Boreux Raphaël, Meex Cécile, Melin Pierrette, Hayette Marie-Pierre, Bours Vincent. |
| 566 | EPI_ISL_417025 | hCoV-19/Belgium/ULG-7019/2020 | *Homo sapiens* | Europe / Belgium / Liège | 2020-03-15 | Department of Clinical Microbiology | GIGA Medical Genomics | Durkin Keith, Artesi Maria, Bontems Sébastien, Boreux Raphaël, Meex Cécile, Melin Pierrette, Hayette Marie-Pierre, Bours Vincent. |
| 567 | EPI_ISL_417028 | hCoV-19/USA/UT-00020/2020 | *Homo sapiens* | North America / USA / Utah | 2020-03-20 | Utah Public Health Laboratory | Utah Public Health Laboratory | Erin Young, Kelly Oakeson |
| 568 | EPI_ISL_417031 | hCoV-19/Australia/QLDID919/2020 | *Homo sapiens* | Oceania / Australia / Queensland / Gold Coast | 2020-03-11 | Pathology Queensland | Public Health Virology Laboratory | Bixing Huang, Alyssa Pyke, Amanda De Jong, Andrew Van Den Hurk, Carmel Taylor, David Warrilow, Doris Genge, Elisabeth Gamez, Glen Hewitson, Ian Maxwell Mackay, Inga Sultana, Jamie McMahon, Jean Barcelon, Judy Northill, Mitchell Finger, Natalie Simpson, Neelima Nair, Peter Burtonclay, Peter Moore, Sarah Wheatley, Sean Moody, Sonja Hall-Mendelin, Timothy Gardam, and Frederick Moore |
| 569 | EPI_ISL_417034 | hCoV-19/Brazil/AMBR-02/2020 | *Homo sapiens* | South America / Brazil / Amazonas State / Manaus | 2020-03-16 | Laboratorio de Ecologia de Doencas Transmissiveis na Amazonia, Instituto Leonidas e Maria Deane - Fiocruz Amazonia | Laboratorio de Ecologia de Doencas Transmissiveis na Amazonia, Instituto Leonidas e Maria Deane - Fiocruz Amazonia | Valdinete Nascimento, André Corado, Fernanda Nascimento, Ágatha Costa, Debora Duarte, Luciana Gonçalves, Michele Jesus, Sérgio Luz, Felipe Naveca |
| 570 | EPI_ISL_417064 | hCoV-19/Hong Kong/case2_VB20017970/2020 | *Homo sapiens* | Asia / Hong Kong | 2020-01-21 | Prince of Wales Hospital | Hong Kong Department of Health | Alan K.L. Tsang, Peter C.W. Yip, Edman T.K. Lam, Rickjason C.W. Chan, Dominic N.C. Tsang |
| 571 | EPI_ISL_417075 | hCoV-19/USA/WA-S22/2020 | *Homo sapiens* | North America / USA / Washington / Snohomish County | 2020-03-02 | Washington State Department of Health | Seattle Flu Study | Chu etl al |
| 572 | EPI_ISL_417082 | hCoV-19/USA/WA-S29/2020 | *Homo sapiens* | North America / USA / Washington / Grant County | 2020-03-02 | Washington State Department of Health | Seattle Flu Study | Chu etl al |
| 573 | EPI_ISL_417093 | hCoV-19/USA/WA-S40/2020 | *Homo sapiens* | North America / USA / Washington / Snohomish County | 2020-02-28 | Washington State Department of Health | Seattle Flu Study | Chu etl al |
| 574 | EPI_ISL_417096 | hCoV-19/USA/WA-S43/2020 | *Homo sapiens* | North America / USA / Washington | 2020-02-27 | Washington State Department of Health | Seattle Flu Study | Chu etl al |
| 575 | EPI_ISL_417117 | hCoV-19/USA/WA-S64/2020 | *Homo sapiens* | North America / USA / Washington / King County | 2020-03-03 | Washington State Department of Health | Seattle Flu Study | Chu etl al |

| | | | | | | | | |
|---|---|---|---|---|---|---|---|---|
| 576 | EPI_ISL_417121 | hCoV-19/USA/WA-S68/2020 | *Homo sapiens* | North America / USA / Washington / King County | 2020-03-04 | Washington State Department of Health | Seattle Flu Study | Chu etl al |
| 577 | EPI_ISL_417123 | hCoV-19/USA/WA-S70/2020 | *Homo sapiens* | North America / USA / Washington / King County | 2020-03-05 | Washington State Department of Health | Seattle Flu Study | Chu etl al |
| 578 | EPI_ISL_417142 | hCoV-19/USA/WA-S89/2020 | *Homo sapiens* | North America / USA / Washington / Umatilla County | 2020-02-29 | Washington State Department of Health | Seattle Flu Study | Chu etl al |
| 579 | EPI_ISL_417148 | hCoV-19/USA/WA-S95/2020 | *Homo sapiens* | North America / USA / Washington / King County | 2020-02-28 | Washington State Department of Health | Seattle Flu Study | Chu etl al |
| 580 | MT226610 | SARS-CoV-2/KMS1/Homo sapiens/2020/CHN | *Homo sapiens* | China | 2020-01-20 | not specified | not specified | Xu,X., Liao,Y., Wang,L., Zhou,X., Xie,Z., Chen,H., Fan,S., Liu,L., Zheng,H., Jiang,G. and Li,Q. |
| 581 | MT020781 | nCoV-FIN-29-Jan-2020 | *Homo sapiens* | Finland | 2020-01-29 | not specified | not specified | Smura,T., Kuivanen,S., Broas,M., Peltola,J., Kallio-Kokko,H. and Vapalahti,O. |
| 582 | MT192773 | SARS-CoV-2/nCoV-19-02S/Homo sapiens/2020/VNM | *Homo sapiens* | Viet Nam: Ho Chi Minh city | 2020-01-22 | not specified | not specified | Nguyen,H.T., Cao,T.M., Pham,H.T.T., Vu,N.P.H., Dao,M.H., Huynh,L.T.K., Nguyen,L.T., Nguyen,N.T., Nguyen,T.T.N., Nguyen,A.H., Luong,Q.C., Nguyen,T.V., Tran,K.C., Pham,Q.D., Tran,T., Hoang,C.Q., Nguyen,T.T., Le,H.Q., Phung,T.M., Vo,T.N.A., Nguyen,S.N., Pham,D.T., Nguyen,T.V. and Phan,L.T. |
| 583 | MT192772 | SARS-CoV-2/nCoV-19-01S/Homo sapiens/2020/VNM | *Homo sapiens* | Viet Nam: Ho Chi Minh city | 2020-01-22 | not specified | not specified | Cao,T.M., Nguyen,H.T., Pham,H.T.T., Vu,N.P.H., Dao,M.H., Huynh,L.T.K., Nguyen,L.T., Nguyen,N.T., Nguyen,T.T.N., Nguyen,A.H., Luong,Q.C., Nguyen,T.V., Tran,K.C., Pham,Q.D., Tran,T., Hoang,C.Q., Nguyen,T.T., Le,H.Q., Phung,T.M., Vo,T.N.A., Nguyen,S.N., Pham,D.T., Phan,L.T. and Nguyen,T.V. |
| 584 | MT192765 | SARS-CoV-2/PC00101P/Homo sapiens/2020/USA | *Homo sapiens* | USA: CA, San Diego County | 2020-03-11 | not specified | not specified | Zeller,M., Anderson,C., Spencer,E., Klitting,R., Robles-Sikisaka,R., Gangavarapu,K., Nicholson,L. and Andersen,K. |
| 585 | MT188341 | USA/MN1-MDH1/2020 | *Homo sapiens* | USA: MN | 2020-03-05 | not specified | not specified | Plumb,M., Garfin,J. and Wang,X. |
| 586 | MT123293 | SARS-CoV-2/IQTC03/Homo sapiens/2020/CHN | *Homo sapiens* | China: Guangzhou | 2020-01-29 | not specified | not specified | Zheng,K., Shi,Y., Sun,J., Huang,J., Zhu,A., Sun,F., Zhuang,Z., Dai,J., Chen,Z., Huang,S., Zhang,Z., Li,X. and Wang,Y. |
| 587 | MT123292 | SARS-CoV-2/IQTC04/Homo sapiens/2020/CHN | *Homo sapiens* | China: Guangzhou | 2020-01-27 | not specified | not specified | Huang,J., Shi,Y., Sun,J., Zheng,K., Zhu,A., Sun,F., Zhuang,Z., Dai,J., Zhang,Z., Huang,S., Wang,Y. and Li,X. |
| 588 | MT159716 | 2019-nCoV/USA-CruiseA-18/2020 | *Homo sapiens* | USA | 2020-02-24 | not specified | not specified | Paden,C.R., Tao,Y., Queen,K., Uehara,A., Zhang,J., Li,Y., Wang,H., Kamili,S., Lu,X., Lynch,B., Sakthivel,S.K.K., Whitaker,B.L., Wang,L., Murray,J.R., Padilla,J., Lee,J., Gerber,S.I., Lindstrom,S. and Tong,S. |
| 589 | MT012098 | SARS-CoV-2/29/Homo sapiens/2020/IND | *Homo sapiens* | India: Kerala State | 2020-01-27 | not specified | not specified | Yadav,P.D., Potdar,V. and Abraham,P. |
| 590 | MT126808 | SARS-CoV-2/SP02/Homo sapiens/2020/BRA | *Homo sapiens* | Brazil | 2020-02-28 | not specified | not specified | Amgarten,D., Malta,F., Ruiz,R.M., Petroni,R., Santana,R.A.F., de Menezes,F.G., Mangueira,C.L.P. and Pinho,J.R.R. |
| 591 | MT049951 | SARS-CoV-2/Yunnan-01/Homo sapiens/2020/CHN | *Homo sapiens* | China: Yunnan | 2020-01-17 | not specified | not specified | Fu,X., Li,D. and Sun,Y. |
| 592 | MT044258 | 2019-nCoV/USA-CA6/2020 | *Homo sapiens* | USA: CA | 2020-01-27 | not specified | not specified | Zhang,J., Queen,K., Li,Y., Tao,Y., Uehara,A., Paden,C.R., Lu,X., Lynch,B., Sakthivel,S.K.K., Whitaker,B.L., Kamili,S., Wang,L., Murray,J.R., Gerber,S.I., Lindstrom,S. and Tong,S. |
| 593 | MT039887 | 2019-nCoV/USA-WI1/2020 | *Homo sapiens* | USA: WI | 2020-01-31 | not specified | not specified | Zhang,J., Uehara,A., Queen,K., Li,Y., Tao,Y., Paden,C.R., Lu,X., Lynch,B., Sakthivel,S.K.K., Whitaker,B.L., Kamili,S., Wang,L., Murray,J.R., Gerber,S.I., Lindstrom,S. and Tong,S. |

| GENOME ID | WSP POSITION IN THE CISTRON/GENE | | | | | | | | | | | |
|---|---|---|---|---|---|---|---|---|---|---|---|---|
| | *nsp3* | *nsp4* | *nsp6* | *nsp12* | *nsp13* | | *nsp14* | *S* | *ORF8* | *N* | | |
| | #318 | #228 | #111 | #967 | #1511 | #1622 | #21 | #1,841 | #251 | #608 | #609 | #610 |
| hCoV 19 Brazil SPBR 12 2020 EPI ISL 416034 | U | C | U | U | C | A | C | G | U | A | A | C |
| hCoV 19 Switzerland SZ1417 2020 EPI ISL 415702 | U | C | U | U | C | A | C | G | U | A | A | C |
| hCoV 19 Belgium BC 03016 2020 EPI ISL 415157 | U | C | G | U | C | A | C | G | U | A | A | C |
| hCoV 19 Belgium DBA 03032 2020 EPI ISL 416475 | U | C | G | U | C | A | C | G | U | A | A | C |
| hCoV 19 Belgium DBD 03024 2020 EPI ISL 416471 | U | C | G | U | C | A | C | G | U | A | A | C |
| hCoV 19 Belgium GMH 03022 2020 EPI ISL 416468 | U | C | G | U | C | A | C | G | U | A | A | C |
| hCoV 19 Belgium MTR 03021 2020 EPI ISL 416467 | U | C | G | U | C | A | C | G | U | A | A | C |
| hCoV 19 Belgium MTR 03026 2020 EPI ISL 416476 | U | C | G | U | C | A | C | G | U | A | A | C |
| hCoV 19 Belgium QKJ 03015 2020 EPI ISL 415158 | U | C | G | U | C | A | C | G | U | A | A | C |
| hCoV 19 Belgium ULG 3000 2020 EPI ISL 417004 | U | C | G | U | C | A | C | G | U | A | A | C |
| hCoV 19 Belgium ULG 6216 2020 EPI ISL 417014 | U | C | G | U | C | A | C | G | U | A | A | C |
| hCoV 19 Belgium ULG 6457 2020 EPI ISL 417015 | U | C | G | U | C | A | C | G | U | A | A | C |
| hCoV 19 Belgium ULG 6942 2020 EPI ISL 417021 | U | C | G | U | C | A | C | G | U | A | A | C |
| hCoV 19 Belgium UMF 03025 2020 EPI ISL 416472 | U | C | G | U | C | A | C | G | U | A | A | C |
| hCoV 19 Brazil SPBR 03 2020 EPI ISL 414014 | U | C | G | U | C | A | C | G | U | A | A | C |
| hCoV 19 Brazil SPBR 08 2020 EPI ISL 416029 | U | C | G | U | C | A | C | G | U | A | A | C |
| hCoV 19 Brazil SPBR 14 2020 EPI ISL 416036 | U | C | G | U | C | A | C | G | U | A | A | C |
| hCoV 19 Chile Santiago 2 2020 EPI ISL 414580 | U | C | G | U | C | A | C | G | U | A | A | C |
| hCoV 19 Denmark SSI 01 2020 EPI ISL 416142 | U | C | G | U | C | A | C | G | U | A | A | C |
| hCoV 19 Denmark SSI 05 2020 EPI ISL 416140 | U | C | G | U | C | A | C | G | U | A | A | C |
| hCoV 19 England 200990723 2020 EPI ISL 414012 | U | C | G | U | C | A | C | G | U | A | A | C |
| hCoV 19 England SHEF BFCB1 2020 EPI ISL 416730 | U | C | G | U | C | A | C | G | U | A | A | C |
| hCoV 19 Finland FIN 313 2020 EPI ISL 414641 | U | C | G | U | C | A | C | G | U | A | A | C |
| hCoV 19 Finland FIN 508 2020 EPI ISL 414643 | U | C | G | U | C | A | C | G | U | A | A | C |
| hCoV 19 Finland FIN03032020C 2020 EPI ISL 413604 | U | C | G | U | C | A | C | G | U | A | A | C |
| hCoV 19 France Valence 425 2020 EPI ISL 416746 | U | C | G | U | C | A | C | G | U | A | A | C |
| hCoV 19 Hungary 49 2020 EPI ISL 416744 | U | C | G | U | C | A | C | G | U | A | A | C |
| hCoV 19 Mexico CDMX InDRE 01 2020 EPI ISL 412972 | U | C | G | U | C | A | C | G | U | A | A | C |
| hCoV 19 Netherlands NA 11 2020 EPI ISL 415467 | U | C | G | U | C | A | C | G | U | A | A | C |
| hCoV 19 Netherlands NA 12 2020 EPI ISL 415468 | U | C | G | U | C | A | C | G | U | A | A | C |
| hCoV 19 Netherlands NA 15 2020 EPI ISL 415471 | U | C | G | U | C | A | C | G | U | A | A | C |
| hCoV 19 Netherlands NA 19 2020 EPI ISL 415475 | U | C | G | U | C | A | C | G | U | A | A | C |
| hCoV 19 Netherlands NA 2 2020 EPI ISL 415476 | U | C | G | U | C | A | C | G | U | A | A | C |
| hCoV 19 Netherlands NA 25 2020 EPI ISL 415482 | U | C | G | U | C | A | C | G | U | A | A | C |
| hCoV 19 Netherlands NoordBrabant 3 2020 EPI ISL 414429 | U | C | G | U | C | A | C | G | U | A | A | C |
| hCoV 19 Netherlands NoordBrabant 36 2020 EPI ISL 414545 | U | C | G | U | C | A | C | G | U | A | A | C |
| hCoV 19 Netherlands NoordHolland 2 2020 EPI ISL 414549 | U | C | G | U | C | A | C | G | U | A | A | C |
| hCoV 19 Netherlands Nootdorp 1364222 2020 EPI ISL 413579 | U | C | G | U | C | A | C | G | U | A | A | C |
| hCoV 19 Netherlands Rotterdam 1364740 2020 EPI ISL 413584 | U | C | G | U | C | A | C | G | U | A | A | C |
| hCoV 19 Netherlands Utrecht 1 2020 EPI ISL 414435 | U | C | G | U | C | A | C | G | U | A | A | C |
| hCoV 19 Netherlands Utrecht 13 2020 EPI ISL 414552 | U | C | G | U | C | A | C | G | U | A | A | C |
| hCoV 19 Netherlands Utrecht 15 2020 EPI ISL 414554 | U | C | G | U | C | A | C | G | U | A | A | C |
| hCoV 19 Netherlands ZuidHolland 13 2020 EPI ISL 414470 | U | C | G | U | C | A | C | G | U | A | A | C |
| hCoV 19 Netherlands ZuidHolland 20 2020 EPI ISL 414562 | U | C | G | U | C | A | C | G | U | A | A | C |
| hCoV 19 Netherlands ZuidHolland 28 2020 EPI ISL 415532 | U | C | G | U | C | A | C | G | U | A | A | C |
| hCoV 19 New Zealand 20VR0189 2020 EPI ISL 416519 | U | C | G | U | C | A | C | G | U | A | A | C |
| hCoV 19 Peru 010 2020 EPI ISL 415787 | U | C | G | U | C | A | C | G | U | A | A | C |
| hCoV 19 Portugal CV62 2020 EPI ISL 413647 | U | C | G | U | C | A | C | G | U | A | A | C |
| hCoV 19 Switzerland 1000477797 2020 EPI ISL 413023 | U | C | G | U | C | A | C | G | U | A | A | C |
| hCoV 19 Switzerland AG7120 2020 EPI ISL 415457 | U | C | G | U | C | A | C | G | U | A | A | C |
| hCoV 19 Switzerland BE6651 2020 EPI ISL 415456 | U | C | G | U | C | A | C | G | U | A | A | C |
| hCoV 19 Switzerland GE1402 2020 EPI ISL 415700 | U | C | G | U | C | A | C | G | U | A | A | C |
| hCoV 19 Switzerland GE1422 2020 EPI ISL 415454 | U | C | G | U | C | A | C | G | U | A | A | C |
| hCoV 19 Switzerland GE3895 2020 EPI ISL 413997 | U | C | G | U | C | A | C | G | U | A | A | C |
| hCoV 19 Switzerland GE8102 2020 EPI ISL 415458 | U | C | G | U | C | A | C | G | U | A | A | C |
| hCoV 19 Switzerland GR2988 2020 EPI ISL 415698 | U | C | G | U | C | A | C | G | U | A | A | C |
| hCoV 19 Switzerland TI2045 2020 EPI ISL 415703 | U | C | G | U | C | A | C | G | U | A | A | C |
| hCoV 19 Switzerland VD5615 2020 EPI ISL 414023 | U | C | G | U | C | A | C | G | U | A | A | C |
| hCoV 19 USA WA UW138 2020 EPI ISL 416676 | U | C | G | U | C | A | C | G | U | A | A | C |
| hCoV 19 USA WA UW159 2020 EPI ISL 416697 | U | C | G | U | C | A | C | G | U | A | A | C |
| hCoV 19 Netherlands Utrecht 1363628 2020 EPI ISL 413589 | U | C | U | U | C | A | C | G | U | G | G | G |
| hCoV 19 Germany BavPat1 2020 EPI ISL 406862 | U | C | G | C | C | A | C | G | U | G | G | G |
| hCoV 19 Belgium BA 02291 2020 EPI ISL 415159 | U | C | G | U | C | A | C | G | U | G | G | G |
| hCoV 19 Belgium BM 03012 2020 EPI ISL 415154 | U | C | G | U | C | A | C | G | U | G | G | G |
| hCoV 19 Belgium DB 03023 2020 EPI ISL 416470 | U | C | G | U | C | A | C | G | U | G | G | G |
| hCoV 19 Belgium SH 03014 2020 EPI ISL 415156 | U | C | G | U | C | A | C | G | U | G | G | G |
| hCoV 19 Belgium SN 03031 2020 EPI ISL 416469 | U | C | G | U | C | A | C | G | U | G | G | G |
| hCoV 19 Belgium ULG 6670 2020 EPI ISL 417018 | U | C | G | U | C | A | C | G | U | G | G | G |
| hCoV 19 Belgium ULG 6754 2020 EPI ISL 417019 | U | C | G | U | C | A | C | G | U | G | G | G |
| hCoV 19 Belgium ULG 6939 2020 EPI ISL 417020 | U | C | G | U | C | A | C | G | U | G | G | G |
| hCoV 19 Belgium VAG 03013 2020 EPI ISL 415155 | U | C | G | U | C | A | C | G | U | G | G | G |
| hCoV 19 Belgium VLM 03011 2020 EPI ISL 415153 | U | C | G | U | C | A | C | G | U | G | G | G |
| hCoV 19 Brazil SPBR 05 2020 EPI ISL 414016 | U | C | G | U | C | A | C | G | U | G | G | G |
| hCoV 19 Brazil SPBR 06 2020 EPI ISL 414015 | U | C | G | U | C | A | C | G | U | G | G | G |
| hCoV 19 Denmark SSI 02 2020 EPI ISL 416143 | U | C | G | U | C | A | C | G | U | G | G | G |
| hCoV 19 Denmark SSI 03 2020 EPI ISL 416144 | U | C | G | U | C | A | C | G | U | G | G | G |
| hCoV 19 Denmark SSI 04 2020 EPI ISL 416153 | U | C | G | U | C | A | C | G | U | G | G | G |
| hCoV 19 Denmark SSI 09 2020 EPI ISL 416141 | U | C | G | U | C | A | C | G | U | G | G | G |
| hCoV 19 Denmark SSI 101 2020 EPI ISL 415646 | U | C | G | U | C | A | C | G | U | G | G | G |
| hCoV 19 Denmark SSI 104 2020 EPI ISL 415648 | U | C | G | U | C | A | C | G | U | G | G | G |
| hCoV 19 England 200990660 2020 EPI ISL 414523 | U | C | G | U | C | A | C | G | U | G | G | G |

| | | | | | | | | | | | | |
|---|---|---|---|---|---|---|---|---|---|---|---|---|
| hCoV 19 England 201040081 2020 EPI ISL 414525 | U | C | G | U | C | A | C | G | U | G | G | G |
| hCoV 19 England SHEF BFCC0 2020 EPI ISL 416731 | U | C | G | U | C | A | C | G | U | G | G | G |
| hCoV 19 Finland FIN 266 2020 EPI ISL 414646 | U | C | G | U | C | A | C | G | U | G | G | G |
| hCoV 19 Finland FIN 455 2020 EPI ISL 414642 | U | C | G | U | C | A | C | G | U | G | G | G |
| hCoV 19 Finland FIN03032020A 2020 EPI ISL 413602 | U | C | G | U | C | A | C | G | U | G | G | G |
| hCoV 19 France B2330 2020 EPI ISL 416502 | U | C | G | U | C | A | C | G | U | G | G | G |
| hCoV 19 France B2336 2020 EPI ISL 416505 | U | C | G | U | C | A | C | G | U | G | G | G |
| hCoV 19 France B2337 2020 EPI ISL 416506 | U | C | G | U | C | A | C | G | U | G | G | G |
| hCoV 19 France B2344 2020 EPI ISL 416509 | U | C | G | U | C | A | C | G | U | G | G | G |
| hCoV 19 France B2346 2020 EPI ISL 416510 | U | C | G | U | C | A | C | G | U | G | G | G |
| hCoV 19 France B2351 2020 EPI ISL 416513 | U | C | G | U | C | A | C | G | U | G | G | G |
| hCoV 19 France BFC2094 2020 EPI ISL 415651 | U | C | G | U | C | A | C | G | U | G | G | G |
| hCoV 19 France BFC2147 2020 EPI ISL 415652 | U | C | G | U | C | A | C | G | U | G | G | G |
| hCoV 19 France Clermont Ferrand 651 2020 EPI ISL 416751 | U | C | G | U | C | A | C | G | U | G | G | G |
| hCoV 19 France GE1973 2020 EPI ISL 414631 | U | C | G | U | C | A | C | G | U | G | G | G |
| hCoV 19 France GE1977 2020 EPI ISL 414632 | U | C | G | U | C | A | C | G | U | G | G | G |
| hCoV 19 France HF1684 2020 EPI ISL 414626 | U | C | G | U | C | A | C | G | U | G | G | G |
| hCoV 19 France HF1795 2020 EPI ISL 414627 | U | C | G | U | C | A | C | G | U | G | G | G |
| hCoV 19 France HF1870 2020 EPI ISL 414629 | U | C | G | U | C | A | C | G | U | G | G | G |
| hCoV 19 France HF1871 2020 EPI ISL 414630 | U | C | G | U | C | A | C | G | U | G | G | G |
| hCoV 19 France HF1988 2020 EPI ISL 414635 | U | C | G | U | C | A | C | G | U | G | G | G |
| hCoV 19 France HF1993 2020 EPI ISL 414637 | U | C | G | U | C | A | C | G | U | G | G | G |
| hCoV 19 France HF1995 2020 EPI ISL 414638 | U | C | G | U | C | A | C | G | U | G | G | G |
| hCoV 19 France HF2039 2020 EPI ISL 415649 | U | C | G | U | C | A | C | G | U | G | G | G |
| hCoV 19 France HF2234 2020 EPI ISL 416495 | U | C | G | U | C | A | C | G | U | G | G | G |
| hCoV 19 France HF2237 2020 EPI ISL 416496 | U | C | G | U | C | A | C | G | U | G | G | G |
| hCoV 19 France HF2239 2020 EPI ISL 416497 | U | C | G | U | C | A | C | G | U | G | G | G |
| hCoV 19 France IDF1980 2020 EPI ISL 414633 | U | C | G | U | C | A | C | G | U | G | G | G |
| hCoV 19 France IDF2256 2020 EPI ISL 416498 | U | C | G | U | C | A | C | G | U | G | G | G |
| hCoV 19 France IDF2278 2020 EPI ISL 416499 | U | C | G | U | C | A | C | G | U | G | G | G |
| hCoV 19 France N1620 2020 EPI ISL 414601 | U | C | G | U | C | A | C | G | U | G | G | G |
| hCoV 19 France N1620 2020 EPI ISL 414624 | U | C | G | U | C | A | C | G | U | G | G | G |
| hCoV 19 France N2223 2020 EPI ISL 416494 | U | C | G | U | C | A | C | G | U | G | G | G |
| hCoV 19 France PL1643 2020 EPI ISL 414625 | U | C | G | U | C | A | C | G | U | G | G | G |
| hCoV 19 France Valence 532 2020 EPI ISL 416749 | U | C | G | U | C | A | C | G | U | G | G | G |
| hCoV 19 Georgia Tb 2020 EPI ISL 416482 | U | C | G | U | C | A | C | G | U | G | G | G |
| hCoV 19 Georgia Tb 273 2020 EPI ISL 416479 | U | C | G | U | C | A | C | G | U | G | G | G |
| hCoV 19 Georgia Tb 477 2020 EPI ISL 415642 | U | C | G | U | C | A | C | G | U | G | G | G |
| hCoV 19 Georgia Tb 673 2020 EPI ISL 416478 | U | C | G | U | C | A | C | G | U | G | G | G |
| hCoV 19 Georgia Tb 712 2020 EPI ISL 416481 | U | C | G | U | C | A | C | G | U | G | G | G |
| hCoV 19 Hungary mbl1 2020 EPI ISL 416426 | U | C | G | U | C | A | C | G | U | G | G | G |
| hCoV 19 Italy CDG1 2020 EPI ISL 412973 | U | C | G | U | C | A | C | G | U | G | G | G |
| hCoV 19 Italy UniSR1 2020 EPI ISL 413489 | U | C | G | U | C | A | C | G | U | G | G | G |
| hCoV 19 Japan SMU 0311S2 2020 EPI ISL 416524 | U | C | G | U | C | A | C | G | U | G | G | G |
| hCoV 19 Japan SMU 0311S3 2020 EPI ISL 416525 | U | C | G | U | C | A | C | G | U | G | G | G |
| hCoV 19 Lithuania ChVir1632 2020 EPI ISL 416741 | U | C | G | U | C | A | C | G | U | G | G | G |
| hCoV 19 Luxembourg Lux1 2020 EPI ISL 413593 | U | C | G | U | C | A | C | G | U | G | G | G |
| hCoV 19 Netherlands Blaricum 1364780 2020 EPI ISL 413566 | U | C | G | U | C | A | C | G | U | G | G | G |
| hCoV 19 Netherlands Flevoland 1 2020 EPI ISL 415460 | U | C | G | U | C | A | C | G | U | G | G | G |
| hCoV 19 Netherlands Gelderland 2 2020 EPI ISL 415462 | U | C | G | U | C | A | C | G | U | G | G | G |
| hCoV 19 Netherlands Haarlem 1363688 2020 EPI ISL 413572 | U | C | G | U | C | A | C | G | U | G | G | G |
| hCoV 19 Netherlands NA 16 2020 EPI ISL 415472 | U | C | G | U | C | A | C | G | U | G | G | G |
| hCoV 19 Netherlands NA 17 2020 EPI ISL 415473 | U | C | G | U | C | A | C | G | U | G | G | G |
| hCoV 19 Netherlands NA 24 2020 EPI ISL 415481 | U | C | G | U | C | A | C | G | U | G | G | G |
| hCoV 19 Netherlands NA 28 2020 EPI ISL 415485 | U | C | G | U | C | A | C | G | U | G | G | G |
| hCoV 19 Netherlands NA 30 2020 EPI ISL 415487 | U | C | G | U | C | A | C | G | U | G | G | G |
| hCoV 19 Netherlands NA 31 2020 EPI ISL 415488 | U | C | G | U | C | A | C | G | U | G | G | G |
| hCoV 19 Netherlands NoordBrabant 1 2020 EPI ISL 414428 | U | C | G | U | C | A | C | G | U | G | G | G |
| hCoV 19 Netherlands NoordBrabant 17 2020 EPI ISL 414457 | U | C | G | U | C | A | C | G | U | G | G | G |
| hCoV 19 Netherlands NoordHolland 3 2020 EPI ISL 415525 | U | C | G | U | C | A | C | G | U | G | G | G |
| hCoV 19 Netherlands ZuidHolland 17 2020 EPI ISL 414559 | U | C | G | U | C | A | C | G | U | G | G | G |
| hCoV 19 Netherlands ZuidHolland 31 2020 EPI ISL 415535 | U | C | G | U | C | A | C | G | U | G | G | G |
| hCoV 19 Netherlands ZuidHolland 9 2020 EPI ISL 414445 | U | C | G | U | C | A | C | G | U | G | G | G |
| hCoV 19 Portugal CV63 2020 EPI ISL 413648 | U | C | G | U | C | A | C | G | U | G | G | G |
| hCoV 19 Saudi Arabia SCDC 3321 2020 EPI ISL 416521 | U | C | G | U | C | A | C | G | U | G | G | G |
| hCoV 19 Scotland CVR05 2020 EPI ISL 414027 | U | C | G | U | C | A | C | G | U | G | G | G |
| hCoV 19 Scotland EDB003 2020 EPI ISL 415640 | U | C | G | U | C | A | C | G | U | G | G | G |
| hCoV 19 Scotland EDB004 2020 EPI ISL 415629 | U | C | G | U | C | A | C | G | U | G | G | G |
| hCoV 19 Switzerland AG0361 2020 EPI ISL 413999 | U | C | G | U | C | A | C | G | U | G | G | G |
| hCoV 19 Switzerland BE2536 2020 EPI ISL 415704 | U | C | G | U | C | A | C | G | U | G | G | G |
| hCoV 19 Switzerland GE3121 2020 EPI ISL 414019 | U | C | G | U | C | A | C | G | U | G | G | G |
| hCoV 19 Switzerland GE4984 2020 EPI ISL 415708 | U | C | G | U | C | A | C | G | U | G | G | G |
| hCoV 19 Switzerland TI9486 2020 EPI ISL 413996 | U | C | G | U | C | A | C | G | U | G | G | G |
| hCoV 19 USA CA MG0987 2020 EPI ISL 416457 | U | C | G | U | C | A | C | G | U | G | G | G |
| hCoV 19 USA CA PC101P 2020 EPI ISL 414648 | U | C | G | U | C | A | C | G | U | G | G | G |
| hCoV 19 USA NY NYUMC2 2020 EPI ISL 416830 | U | C | G | U | C | A | C | G | U | G | G | G |
| hCoV 19 USA NY NYUMC3 2020 EPI ISL 416831 | U | C | G | U | C | A | C | G | U | G | G | G |
| hCoV 19 USA UPHL 05 2020 EPI ISL 415543 | U | C | G | U | C | A | C | G | U | G | G | G |
| hCoV 19 USA WA UW104 2020 EPI ISL 416642 | U | C | G | U | C | A | C | G | U | G | G | G |
| hCoV 19 USA WA UW110 2020 EPI ISL 416648 | U | C | G | U | C | A | C | G | U | G | G | G |
| hCoV 19 USA WA UW115 2020 EPI ISL 416653 | U | C | G | U | C | A | C | G | U | G | G | G |
| hCoV 19 USA WA UW117 2020 EPI ISL 416655 | U | C | G | U | C | A | C | G | U | G | G | G |
| hCoV 19 USA WA UW122 2020 EPI ISL 416660 | U | C | G | U | C | A | C | G | U | G | G | G |
| hCoV 19 USA WA UW123 2020 EPI ISL 416661 | U | C | G | U | C | A | C | G | U | G | G | G |

| | | | | | | | | | | | | |
|---|---|---|---|---|---|---|---|---|---|---|---|---|
| hCoV 19 USA WA UW134 2020 EPI ISL 416672 | U | C | G | U | C | A | C | G | U | G | G | G |
| hCoV 19 USA WA UW136 2020 EPI ISL 416674 | U | C | G | U | C | A | C | G | U | G | G | G |
| hCoV 19 USA WA UW140 2020 EPI ISL 416678 | U | C | G | U | C | A | C | G | U | G | G | G |
| hCoV 19 USA WA UW153 2020 EPI ISL 416691 | U | C | G | U | C | A | C | G | U | G | G | G |
| hCoV 19 USA WA UW155 2020 EPI ISL 416693 | U | C | G | U | C | A | C | G | U | G | G | G |
| hCoV 19 USA WA UW160 2020 EPI ISL 416698 | U | C | G | U | C | A | C | G | U | G | G | G |
| hCoV 19 USA WA UW169 2020 EPI ISL 416707 | U | C | G | U | C | A | C | G | U | G | G | G |
| hCoV 19 USA WA UW178 2020 EPI ISL 416716 | U | C | G | U | C | A | C | G | U | G | G | G |
| hCoV 19 USA WA UW186 2020 EPI ISL 416724 | U | C | G | U | C | A | C | G | U | G | G | G |
| hCoV 19 USA WA UW187 2020 EPI ISL 416725 | U | C | G | U | C | A | C | G | U | G | G | G |
| hCoV 19 USA WA UW191 2020 EPI ISL 416729 | U | C | G | U | C | A | C | G | U | G | G | G |
| hCoV 19 USA WA UW60 2020 EPI ISL 415625 | U | C | G | U | C | A | C | G | U | G | G | G |
| hCoV 19 USA WA UW69 2020 EPI ISL 415597 | U | C | G | U | C | A | C | G | U | G | G | G |
| hCoV 19 USA WA UW73 2020 EPI ISL 415601 | U | C | G | U | C | A | C | G | U | G | G | G |
| hCoV 19 USA WA UW78 2020 EPI ISL 416434 | U | C | G | U | C | A | C | G | U | G | G | G |
| hCoV 19 USA WA UW82 2020 EPI ISL 416438 | U | C | G | U | C | A | C | G | U | G | G | G |
| hCoV 19 USA WA UW96 2020 EPI ISL 416452 | U | C | G | U | C | A | C | G | U | G | G | G |
| hCoV 19 Wales PHW1 2020 EPI ISL 413555 | U | C | G | U | C | A | C | G | U | G | G | G |
| MT192765.1 Severe acute respiratory syndrome coronavirus 2 | U | C | G | U | C | A | C | G | U | G | G | G |
| hCoV 19 Australia QLDID919 2020 EPI ISL 417031 | C | U | G | C | U | G | U | A | C | G | G | G |
| hCoV 19 USA CA CDPH UC9 2020 EPI ISL 413928 | C | U | G | C | U | G | U | A | C | G | G | G |
| hCoV 19 USA MN3 MDH3 2020 EPI ISL 414590 | C | U | G | C | U | G | U | A | C | G | G | G |
| hCoV 19 USA UC CDPH UC11 2020 EPI ISL 413931 | C | U | G | C | U | G | U | A | C | G | G | G |
| hCoV 19 USA UT 00020 2020 EPI ISL 417028 | C | U | G | C | U | G | U | A | C | G | G | G |
| hCoV 19 USA WA S11 2020 EPI ISL 416466 | C | U | G | C | U | G | U | A | C | G | G | G |
| hCoV 19 USA WA S2 2020 EPI ISL 413456 | C | U | G | C | U | G | U | A | C | G | G | G |
| hCoV 19 USA WA S22 2020 EPI ISL 417075 | C | U | G | C | U | G | U | A | C | G | G | G |
| hCoV 19 USA WA S29 2020 EPI ISL 417082 | C | U | G | C | U | G | U | A | C | G | G | G |
| hCoV 19 USA WA S40 2020 EPI ISL 417093 | C | U | G | C | U | G | U | A | C | G | G | G |
| hCoV 19 USA WA S43 2020 EPI ISL 417096 | C | U | G | C | U | G | U | A | C | G | G | G |
| hCoV 19 USA WA S64 2020 EPI ISL 417117 | C | U | G | C | U | G | U | A | C | G | G | G |
| hCoV 19 USA WA S68 2020 EPI ISL 417121 | C | U | G | C | U | G | U | A | C | G | G | G |
| hCoV 19 USA WA S70 2020 EPI ISL 417123 | C | U | G | C | U | G | U | A | C | G | G | G |
| hCoV 19 USA WA S89 2020 EPI ISL 417142 | C | U | G | C | U | G | U | A | C | G | G | G |
| hCoV 19 USA WA S95 2020 EPI ISL 417148 | C | U | G | C | U | G | U | A | C | G | G | G |
| hCoV 19 USA WA UW100 2020 EPI ISL 416638 | C | U | G | C | U | G | U | A | C | G | G | G |
| hCoV 19 USA WA UW103 2020 EPI ISL 416641 | C | U | G | C | U | G | U | A | C | G | G | G |
| hCoV 19 USA WA UW106 2020 EPI ISL 416644 | C | U | G | C | U | G | U | A | C | G | G | G |
| hCoV 19 USA WA UW111 2020 EPI ISL 416649 | C | U | G | C | U | G | U | A | C | G | G | G |
| hCoV 19 USA WA UW112 2020 EPI ISL 416650 | C | U | G | C | U | G | U | A | C | G | G | G |
| hCoV 19 USA WA UW113 2020 EPI ISL 416651 | C | U | G | C | U | G | U | A | C | G | G | G |
| hCoV 19 USA WA UW114 2020 EPI ISL 416652 | C | U | G | C | U | G | U | A | C | G | G | G |
| hCoV 19 USA WA UW118 2020 EPI ISL 416656 | C | U | G | C | U | G | U | A | C | G | G | G |
| hCoV 19 USA WA UW119 2020 EPI ISL 416657 | C | U | G | C | U | G | U | A | C | G | G | G |
| hCoV 19 USA WA UW124 2020 EPI ISL 416662 | C | U | G | C | U | G | U | A | C | G | G | G |
| hCoV 19 USA WA UW125 2020 EPI ISL 416663 | C | U | G | C | U | G | U | A | C | G | G | G |
| hCoV 19 USA WA UW126 2020 EPI ISL 416664 | C | U | G | C | U | G | U | A | C | G | G | G |
| hCoV 19 USA WA UW127 2020 EPI ISL 416665 | C | U | G | C | U | G | U | A | C | G | G | G |
| hCoV 19 USA WA UW128 2020 EPI ISL 416666 | C | U | G | C | U | G | U | A | C | G | G | G |
| hCoV 19 USA WA UW129 2020 EPI ISL 416667 | C | U | G | C | U | G | U | A | C | G | G | G |
| hCoV 19 USA WA UW130 2020 EPI ISL 416668 | C | U | G | C | U | G | U | A | C | G | G | G |
| hCoV 19 USA WA UW131 2020 EPI ISL 416669 | C | U | G | C | U | G | U | A | C | G | G | G |
| hCoV 19 USA WA UW139 2020 EPI ISL 416677 | C | U | G | C | U | G | U | A | C | G | G | G |
| hCoV 19 USA WA UW141 2020 EPI ISL 416679 | C | U | G | C | U | G | U | A | C | G | G | G |
| hCoV 19 USA WA UW142 2020 EPI ISL 416680 | C | U | G | C | U | G | U | A | C | G | G | G |
| hCoV 19 USA WA UW145 2020 EPI ISL 416683 | C | U | G | C | U | G | U | A | C | G | G | G |
| hCoV 19 USA WA UW148 2020 EPI ISL 416686 | C | U | G | C | U | G | U | A | C | G | G | G |
| hCoV 19 USA WA UW149 2020 EPI ISL 416687 | C | U | G | C | U | G | U | A | C | G | G | G |
| hCoV 19 USA WA UW15 2020 EPI ISL 414363 | C | U | G | C | U | G | U | A | C | G | G | G |
| hCoV 19 USA WA UW150 2020 EPI ISL 416688 | C | U | G | C | U | G | U | A | C | G | G | G |
| hCoV 19 USA WA UW152 2020 EPI ISL 416690 | C | U | G | C | U | G | U | A | C | G | G | G |
| hCoV 19 USA WA UW156 2020 EPI ISL 416694 | C | U | G | C | U | G | U | A | C | G | G | G |
| hCoV 19 USA WA UW162 2020 EPI ISL 416700 | C | U | G | C | U | G | U | A | C | G | G | G |
| hCoV 19 USA WA UW165 2020 EPI ISL 416703 | C | U | G | C | U | G | U | A | C | G | G | G |
| hCoV 19 USA WA UW171 2020 EPI ISL 416709 | C | U | G | C | U | G | U | A | C | G | G | G |
| hCoV 19 USA WA UW172 2020 EPI ISL 416710 | C | U | G | C | U | G | U | A | C | G | G | G |
| hCoV 19 USA WA UW173 2020 EPI ISL 416711 | C | U | G | C | U | G | U | A | C | G | G | G |
| hCoV 19 USA WA UW175 2020 EPI ISL 416713 | C | U | G | C | U | G | U | A | C | G | G | G |
| hCoV 19 USA WA UW176 2020 EPI ISL 416714 | C | U | G | C | U | G | U | A | C | G | G | G |
| hCoV 19 USA WA UW177 2020 EPI ISL 416715 | C | U | G | C | U | G | U | A | C | G | G | G |
| hCoV 19 USA WA UW179 2020 EPI ISL 416717 | C | U | G | C | U | G | U | A | C | G | G | G |
| hCoV 19 USA WA UW180 2020 EPI ISL 416718 | C | U | G | C | U | G | U | A | C | G | G | G |
| hCoV 19 USA WA UW183 2020 EPI ISL 416721 | C | U | G | C | U | G | U | A | C | G | G | G |
| hCoV 19 USA WA UW188 2020 EPI ISL 416726 | C | U | G | C | U | G | U | A | C | G | G | G |
| hCoV 19 USA WA UW189 2020 EPI ISL 416727 | C | U | G | C | U | G | U | A | C | G | G | G |
| hCoV 19 USA WA UW190 2020 EPI ISL 416728 | C | U | G | C | U | G | U | A | C | G | G | G |
| hCoV 19 USA WA UW20 2020 EPI ISL 414368 | C | U | G | C | U | G | U | A | C | G | G | G |
| hCoV 19 USA WA UW23 2020 EPI ISL 414592 | C | U | G | C | U | G | U | A | C | G | G | G |
| hCoV 19 USA WA UW26 2020 EPI ISL 414595 | C | U | G | C | U | G | U | A | C | G | G | G |
| hCoV 19 USA WA UW31 2020 EPI ISL 414618 | C | U | G | C | U | G | U | A | C | G | G | G |
| hCoV 19 USA WA UW41 2020 EPI ISL 415606 | C | U | G | C | U | G | U | A | C | G | G | G |
| hCoV 19 USA WA UW43 2020 EPI ISL 415608 | C | U | G | C | U | G | U | A | C | G | G | G |
| hCoV 19 USA WA UW44 2020 EPI ISL 415609 | C | U | G | C | U | G | U | A | C | G | G | G |
| hCoV 19 USA WA UW46 2020 EPI ISL 415611 | C | U | G | C | U | G | U | A | C | G | G | G |

| | | | | | | | | | | | | |
|---|---|---|---|---|---|---|---|---|---|---|---|---|
| hCoV 19 USA WA UW50 2020 EPI ISL 415615 | C | U | G | C | U | G | U | A | C | G | G | G |
| hCoV 19 USA WA UW52 2020 EPI ISL 415617 | C | U | G | C | U | G | U | A | C | G | G | G |
| hCoV 19 USA WA UW54 2020 EPI ISL 415619 | C | U | G | C | U | G | U | A | C | G | G | G |
| hCoV 19 USA WA UW55 2020 EPI ISL 415620 | C | U | G | C | U | G | U | A | C | G | G | G |
| hCoV 19 USA WA UW56 2020 EPI ISL 415621 | C | U | G | C | U | G | U | A | C | G | G | G |
| hCoV 19 USA WA UW57 2020 EPI ISL 415622 | C | U | G | C | U | G | U | A | C | G | G | G |
| hCoV 19 USA WA UW59 2020 EPI ISL 415624 | C | U | G | C | U | G | U | A | C | G | G | G |
| hCoV 19 USA WA UW62 2020 EPI ISL 415627 | C | U | G | C | U | G | U | A | C | G | G | G |
| hCoV 19 USA WA UW63 2020 EPI ISL 415591 | C | U | G | C | U | G | U | A | C | G | G | G |
| hCoV 19 USA WA UW66 2020 EPI ISL 415594 | C | U | G | C | U | G | U | A | C | G | G | G |
| hCoV 19 USA WA UW67 2020 EPI ISL 415595 | C | U | G | C | U | G | U | A | C | G | G | G |
| hCoV 19 USA WA UW68 2020 EPI ISL 415596 | C | U | G | C | U | G | U | A | C | G | G | G |
| hCoV 19 USA WA UW70 2020 EPI ISL 415598 | C | U | G | C | U | G | U | A | C | G | G | G |
| hCoV 19 USA WA UW74 2020 EPI ISL 415602 | C | U | G | C | U | G | U | A | C | G | G | G |
| hCoV 19 USA WA UW75 2020 EPI ISL 415603 | C | U | G | C | U | G | U | A | C | G | G | G |
| hCoV 19 USA WA UW76 2020 EPI ISL 415604 | C | U | G | C | U | G | U | A | C | G | G | G |
| hCoV 19 USA WA UW77 2020 EPI ISL 416433 | C | U | G | C | U | G | U | A | C | G | G | G |
| hCoV 19 USA WA UW79 2020 EPI ISL 416435 | C | U | G | C | U | G | U | A | C | G | G | G |
| hCoV 19 USA WA UW80 2020 EPI ISL 416436 | C | U | G | C | U | G | U | A | C | G | G | G |
| hCoV 19 USA WA UW83 2020 EPI ISL 416439 | C | U | G | C | U | G | U | A | C | G | G | G |
| hCoV 19 USA WA UW85 2020 EPI ISL 416441 | C | U | G | C | U | G | U | A | C | G | G | G |
| hCoV 19 USA WA UW86 2020 EPI ISL 416442 | C | U | G | C | U | G | U | A | C | G | G | G |
| hCoV 19 USA WA UW87 2020 EPI ISL 416443 | C | U | G | C | U | G | U | A | C | G | G | G |
| hCoV 19 USA WA UW88 2020 EPI ISL 416444 | C | U | G | C | U | G | U | A | C | G | G | G |
| hCoV 19 USA WA UW89 2020 EPI ISL 416445 | C | U | G | C | U | G | U | A | C | G | G | G |
| hCoV 19 USA WA UW91 2020 EPI ISL 416447 | C | U | G | C | U | G | U | A | C | G | G | G |
| hCoV 19 USA WA UW92 2020 EPI ISL 416448 | C | U | G | C | U | G | U | A | C | G | G | G |
| hCoV 19 USA WA UW95 2020 EPI ISL 416451 | C | U | G | C | U | G | U | A | C | G | G | G |
| hCoV 19 USA WA UW97 2020 EPI ISL 416635 | C | U | G | C | U | G | U | A | C | G | G | G |
| hCoV 19 USA WA UW98 2020 EPI ISL 416636 | C | U | G | C | U | G | U | A | C | G | G | G |
| hCoV 19 USA WA UW99 2020 EPI ISL 416637 | C | U | G | C | U | G | U | A | C | G | G | G |
| hCoV 19 USA WA12 UW8 2020 EPI ISL 413563 | C | U | G | C | U | G | U | A | C | G | G | G |
| hCoV 19 USA WA2 2020 EPI ISL 412970 | C | U | G | C | U | G | U | A | C | G | G | G |
| hCoV 19 USA WA4 UW2 2020 EPI ISL 413455 | C | U | G | C | U | G | U | A | C | G | G | G |
| hCoV 19 USA WA7 UW4 2020 EPI ISL 413458 | C | U | G | C | U | G | U | A | C | G | G | G |
| MT188341.1 Severe acute respiratory syndrome coronavirus 2 | C | U | G | C | U | G | U | A | C | G | G | G |
| hCoV 19 Australia NSW03 2020 EPI ISL 408977 | C | C | G | C | C | A | C | A | U | G | G | G |
| hCoV 19 Australia VIC01 2020 EPI ISL 406844 | C | C | G | C | C | A | C | A | U | G | G | G |
| hCoV 19 Australia VIC03 2020 EPI ISL 416411 | C | C | G | C | C | A | C | A | U | G | G | G |
| hCoV 19 Belgium ULG 7019 2020 EPI ISL 417025 | C | C | G | C | C | A | C | A | U | G | G | G |
| hCoV 19 Brazil SPBR 10 2020 EPI ISL 416032 | C | C | G | C | C | A | C | A | U | G | G | G |
| hCoV 19 Brazil SPBR 11 2020 EPI ISL 416033 | C | C | G | C | C | A | C | A | U | G | G | G |
| hCoV 19 Cambodia 0012 2020 EPI ISL 411902 | C | C | G | C | C | A | C | A | U | G | G | G |
| hCoV 19 Canada BC 40860 2020 EPI ISL 415583 | C | C | G | C | C | A | C | A | U | G | G | G |
| hCoV 19 China IQTC01 2020 EPI ISL 412966 | C | C | G | C | C | A | C | A | U | G | G | G |
| hCoV 19 China IQTC02 2020 EPI ISL 412967 | C | C | G | C | C | A | C | A | U | G | G | G |
| hCoV 19 China WF0002 2020 EPI ISL 413692 | C | C | G | C | C | A | C | A | U | G | G | G |
| hCoV 19 China WF0003 2020 EPI ISL 413693 | C | C | G | C | C | A | C | A | U | G | G | G |
| hCoV 19 China WH 09 2020 EPI ISL 411957 | C | C | G | C | C | A | C | A | U | G | G | G |
| hCoV 19 England SHEF BFD54 2020 EPI ISL 416740 | C | C | G | C | C | A | C | A | U | G | G | G |
| hCoV 19 Foshan 20SF207 2020 EPI ISL 406534 | C | C | G | C | C | A | C | A | U | G | G | G |
| hCoV 19 Foshan 20SF210 2020 EPI ISL 406535 | C | C | G | C | C | A | C | A | U | G | G | G |
| hCoV 19 Foshan 20SF211 2020 EPI ISL 406536 | C | C | G | C | C | A | C | A | U | G | G | G |
| hCoV 19 France IDF0372 2020 EPI ISL 406596 | C | C | G | C | C | A | C | A | U | G | G | G |
| hCoV 19 France IDF0372 isl 2020 EPI ISL 410720 | C | C | G | C | C | A | C | A | U | G | G | G |
| hCoV 19 France IDF0373 2020 EPI ISL 406597 | C | C | G | C | C | A | C | A | U | G | G | G |
| hCoV 19 France IDF0626 2020 EPI ISL 408431 | C | C | G | C | C | A | C | A | U | G | G | G |
| hCoV 19 Fujian 13 2020 EPI ISL 411066 | C | C | G | C | C | A | C | A | U | G | G | G |
| hCoV 19 Germany NRW 09 2020 EPI ISL 414509 | C | C | G | C | C | A | C | A | U | G | G | G |
| hCoV 19 Guangdong 2020XN4448 P0002 2020 EPI ISL 413857 | C | C | G | C | C | A | C | A | U | G | G | G |
| hCoV 19 Guangdong 20SF014 2020 EPI ISL 403934 | C | C | G | C | C | A | C | A | U | G | G | G |
| hCoV 19 Guangdong 20SF028 2020 EPI ISL 403936 | C | C | G | C | C | A | C | A | U | G | G | G |
| hCoV 19 Guangdong 20SF174 2020 EPI ISL 406531 | C | C | G | C | C | A | C | A | U | G | G | G |
| hCoV 19 Guangdong 20SF201 2020 EPI ISL 406538 | C | C | G | C | C | A | C | A | U | G | G | G |
| hCoV 19 Guangdong GD2020080 P0010 2020 EPI ISL 413861 | C | C | G | C | C | A | C | A | U | G | G | G |
| hCoV 19 Guangdong GD2020087 P0008 2020 EPI ISL 413863 | C | C | G | C | C | A | C | A | U | G | G | G |
| hCoV 19 Guangzhou 20SF206 2020 EPI ISL 406533 | C | C | G | C | C | A | C | A | U | G | G | G |
| hCoV 19 Guangzhou GZMU0014 2020 EPI ISL 414692 | C | C | G | C | C | A | C | A | U | G | G | G |
| hCoV 19 Hangzhou HZ 1 2020 EPI ISL 406970 | C | C | G | C | C | A | C | A | U | G | G | G |
| hCoV 19 Hangzhou HZCDC0001 2020 EPI ISL 407313 | C | C | G | C | C | A | C | A | U | G | G | G |
| hCoV 19 Hangzhou ZJU 03 2020 EPI ISL 416044 | C | C | G | C | C | A | C | A | U | G | G | G |
| hCoV 19 Hangzhou ZJU 06 2020 EPI ISL 416047 | C | C | G | C | C | A | C | A | U | G | G | G |
| hCoV 19 Hangzhou ZJU 07 2020 EPI ISL 416425 | C | C | G | C | C | A | C | A | U | G | G | G |
| hCoV 19 Hangzhou ZJU 09 2020 EPI ISL 416474 | C | C | G | C | C | A | C | A | U | G | G | G |
| hCoV 19 Hong Kong case2 VB20017970 2020 EPI ISL 417064 | C | C | G | C | C | A | C | A | U | G | G | G |
| hCoV 19 Hong Kong VB20024950 2020 EPI ISL 412029 | C | C | G | C | C | A | C | A | U | G | G | G |
| hCoV 19 Ireland COR 20134 2020 EPI ISL 414487 | C | C | G | C | C | A | C | A | U | G | G | G |
| hCoV 19 Japan KY V 029 2020 EPI ISL 408669 | C | C | G | C | C | A | C | A | U | G | G | G |
| hCoV 19 Japan TKYE6182 2020 EPI ISL 414511 | C | C | G | C | C | A | C | A | U | G | G | G |
| hCoV 19 Jiangsu IVDC JS 001 2020 EPI ISL 408488 | C | C | G | C | C | A | C | A | U | G | G | G |
| hCoV 19 Jiangsu JS01 2020 EPI ISL 411950 | C | C | G | C | C | A | C | A | U | G | G | G |
| hCoV 19 Jiangsu JS02 2020 EPI ISL 411952 | C | C | G | C | C | A | C | A | U | G | G | G |
| hCoV 19 Jiangxi IVDC JX 002 2020 EPI ISL 408486 | C | C | G | C | C | A | C | A | U | G | G | G |
| hCoV 19 Jingzhou HBCDC HB 01 2020 EPI ISL 412459 | C | C | G | C | C | A | C | A | U | G | G | G |

| | | | | | | | | | | | | |
|---|---|---|---|---|---|---|---|---|---|---|---|---|
| hCoV 19 Malaysia MKAK CL 2020 5045 2020 EPI ISL 416829 | C | C | G | C | C | A | C | A | U | G | G | G |
| hCoV 19 Malaysia MKAK CL 2020 5047 2020 EPI ISL 416866 | C | C | G | C | C | A | C | A | U | G | G | G |
| hCoV 19 Nepal 61 2020 EPI ISL 410301 | C | C | G | C | C | A | C | A | U | G | G | G |
| hCoV 19 Netherlands Limburg 7 2020 EPI ISL 415464 | C | C | G | C | C | A | C | A | U | G | G | G |
| hCoV 19 Netherlands NA 6 2020 EPI ISL 415495 | C | C | G | C | C | A | C | A | U | G | G | G |
| hCoV 19 Netherlands ZuidHolland 27 2020 EPI ISL 415531 | C | C | G | C | C | A | C | A | U | G | G | G |
| hCoV 19 Nonthaburi 61 2020 EPI ISL 403962 | C | C | G | C | C | A | C | A | U | G | G | G |
| hCoV 19 Poland PL P1 2020 EPI ISL 416488 | C | C | G | C | C | A | C | A | U | G | G | G |
| hCoV 19 Shanghai SH0007 2020 EPI ISL 416320 | C | C | G | C | C | A | C | A | U | G | G | G |
| hCoV 19 Shanghai SH0008 2020 EPI ISL 416321 | C | C | G | C | C | A | C | A | U | G | G | G |
| hCoV 19 Shanghai SH0012 2020 EPI ISL 416325 | C | C | G | C | C | A | C | A | U | G | G | G |
| hCoV 19 Shanghai SH0027 2020 EPI ISL 416336 | C | C | G | C | C | A | C | A | U | G | G | G |
| hCoV 19 Shanghai SH0044 2020 EPI ISL 416353 | C | C | G | C | C | A | C | A | U | G | G | G |
| hCoV 19 Shanghai SH0053 2020 EPI ISL 416361 | C | C | G | C | C | A | C | A | U | G | G | G |
| hCoV 19 Shanghai SH0054 2020 EPI ISL 416362 | C | C | G | C | C | A | C | A | U | G | G | G |
| hCoV 19 Shanghai SH0055 2020 EPI ISL 416363 | C | C | G | C | C | A | C | A | U | G | G | G |
| hCoV 19 Shanghai SH0060 2020 EPI ISL 416367 | C | C | G | C | C | A | C | A | U | G | G | G |
| hCoV 19 Shanghai SH0073 2020 EPI ISL 416376 | C | C | G | C | C | A | C | A | U | G | G | G |
| hCoV 19 Shanghai SH0094 2020 EPI ISL 416390 | C | C | G | C | C | A | C | A | U | G | G | G |
| hCoV 19 Shanghai SH01 2020 EPI ISL 414510 | C | C | G | C | C | A | C | A | U | G | G | G |
| hCoV 19 Shanghai SH0107 2020 EPI ISL 416397 | C | C | G | C | C | A | C | A | U | G | G | G |
| hCoV 19 Shanghai SH0109 2020 EPI ISL 416398 | C | C | G | C | C | A | C | A | U | G | G | G |
| hCoV 19 Shanghai SH0110 2020 EPI ISL 416399 | C | C | G | C | C | A | C | A | U | G | G | G |
| hCoV 19 Shanghai SH0114 2020 EPI ISL 416401 | C | C | G | C | C | A | C | A | U | G | G | G |
| hCoV 19 Shanghai SH0121 2020 EPI ISL 416405 | C | C | G | C | C | A | C | A | U | G | G | G |
| hCoV 19 Shenzhen SZTH 003 2020 EPI ISL 406594 | C | C | G | C | C | A | C | A | U | G | G | G |
| hCoV 19 Shenzhen SZTH 004 2020 EPI ISL 406595 | C | C | G | C | C | A | C | A | U | G | G | G |
| hCoV 19 Singapore 1 2020 EPI ISL 406973 | C | C | G | C | C | A | C | A | U | G | G | G |
| hCoV 19 Singapore 11 2020 EPI ISL 410719 | C | C | G | C | C | A | C | A | U | G | G | G |
| hCoV 19 Singapore 2 2020 EPI ISL 407987 | C | C | G | C | C | A | C | A | U | G | G | G |
| hCoV 19 Singapore 5 2020 EPI ISL 410536 | C | C | G | C | C | A | C | A | U | G | G | G |
| hCoV 19 Singapore 7 2020 EPI ISL 410713 | C | C | G | C | C | A | C | A | U | G | G | G |
| hCoV 19 Singapore 9 2020 EPI ISL 410715 | C | C | G | C | C | A | C | A | U | G | G | G |
| hCoV 19 South Korea KUMC02 2020 EPI ISL 413018 | C | C | G | C | C | A | C | A | U | G | G | G |
| hCoV 19 South Korea SNU01 2020 EPI ISL 411929 | C | C | G | C | C | A | C | A | U | G | G | G |
| hCoV 19 Sweden 01 2020 EPI ISL 411951 | C | C | G | C | C | A | C | A | U | G | G | G |
| hCoV 19 Taiwan 2 2020 EPI ISL 406031 | C | C | G | C | C | A | C | A | U | G | G | G |
| hCoV 19 Taiwan NTU02 2020 EPI ISL 410218 | C | C | G | C | C | A | C | A | U | G | G | G |
| hCoV 19 USA CA CDPH UC2 2020 EPI ISL 413558 | C | C | G | C | C | A | C | A | U | G | G | G |
| hCoV 19 USA CA CDPH UC3 2020 EPI ISL 413559 | C | C | G | C | C | A | C | A | U | G | G | G |
| hCoV 19 USA CA2 2020 EPI ISL 406036 | C | C | G | C | C | A | C | A | U | G | G | G |
| hCoV 19 USA CA3 2020 EPI ISL 408008 | C | C | G | C | C | A | C | A | U | G | G | G |
| hCoV 19 USA CA5 2020 EPI ISL 408010 | C | C | G | C | C | A | C | A | U | G | G | G |
| hCoV 19 USA CA8 2020 EPI ISL 411955 | C | C | G | C | C | A | C | A | U | G | G | G |
| hCoV 19 USA CA9 2020 EPI ISL 412862 | C | C | G | C | C | A | C | A | U | G | G | G |
| hCoV 19 USA CruiseA 1 2020 EPI ISL 413606 | C | C | G | C | C | A | C | A | U | G | G | G |
| hCoV 19 USA CruiseA 10 2020 EPI ISL 413615 | C | C | G | C | C | A | C | A | U | G | G | G |
| hCoV 19 USA CruiseA 11 2020 EPI ISL 413616 | C | C | G | C | C | A | C | A | U | G | G | G |
| hCoV 19 USA CruiseA 12 2020 EPI ISL 413617 | C | C | G | C | C | A | C | A | U | G | G | G |
| hCoV 19 USA CruiseA 14 2020 EPI ISL 413619 | C | C | G | C | C | A | C | A | U | G | G | G |
| hCoV 19 USA CruiseA 17 2020 EPI ISL 413622 | C | C | G | C | C | A | C | A | U | G | G | G |
| hCoV 19 USA CruiseA 2 2020 EPI ISL 413607 | C | C | G | C | C | A | C | A | U | G | G | G |
| hCoV 19 USA CruiseA 4 2020 EPI ISL 413609 | C | C | G | C | C | A | C | A | U | G | G | G |
| hCoV 19 USA CruiseA 6 2020 EPI ISL 413611 | C | C | G | C | C | A | C | A | U | G | G | G |
| hCoV 19 USA CruiseA 7 2020 EPI ISL 413612 | C | C | G | C | C | A | C | A | U | G | G | G |
| hCoV 19 USA CruiseA 8 2020 EPI ISL 413613 | C | C | G | C | C | A | C | A | U | G | G | G |
| hCoV 19 USA MN2 MDH2 2020 EPI ISL 414589 | C | C | G | C | C | A | C | A | U | G | G | G |
| hCoV 19 Wales PHW32 2020 EPI ISL 415920 | C | C | G | C | C | A | C | A | U | G | G | G |
| hCoV 19 Wales PHW35 2020 EPI ISL 416024 | C | C | G | C | C | A | C | A | U | G | G | G |
| hCoV 19 Wuhan HBCDC HB 01 2019 EPI ISL 402132 | C | C | G | C | C | A | C | A | U | G | G | G |
| hCoV 19 Wuhan HBCDC HB 02 2019 EPI ISL 412898 | C | C | G | C | C | A | C | A | U | G | G | G |
| hCoV 19 Wuhan IPBCAMS WH 01 2019 EPI ISL 402123 | C | C | G | C | C | A | C | A | U | G | G | G |
| hCoV 19 Wuhan IPBCAMS WH 02 2019 EPI ISL 403931 | C | C | G | C | C | A | C | A | U | G | G | G |
| hCoV 19 Wuhan IPBCAMS WH 03 2019 EPI ISL 403930 | C | C | G | C | C | A | C | A | U | G | G | G |
| hCoV 19 Wuhan IPBCAMS WH 05 2020 EPI ISL 403928 | C | C | G | C | C | A | C | A | U | G | G | G |
| hCoV 19 Wuhan IVDC HB 04 2020 EPI ISL 402120 | C | C | G | C | C | A | C | A | U | G | G | G |
| hCoV 19 Wuhan IVDC HB 05 2019 EPI ISL 402121 | C | C | G | C | C | A | C | A | U | G | G | G |
| hCoV 19 Wuhan IVDC HB envF13 20 2020 EPI ISL 408514 | C | C | G | C | C | A | C | A | U | G | G | G |
| hCoV 19 Wuhan IVDC HB envF13 21 2020 EPI ISL 408515 | C | C | G | C | C | A | C | A | U | G | G | G |
| hCoV 19 Wuhan WH01 2019 EPI ISL 406798 | C | C | G | C | C | A | C | A | U | G | G | G |
| hCoV 19 Wuhan WIV02 2019 EPI ISL 402127 | C | C | G | C | C | A | C | A | U | G | G | G |
| hCoV 19 Wuhan WIV04 2019 EPI ISL 402124 | C | C | G | C | C | A | C | A | U | G | G | G |
| hCoV 19 Wuhan WIV05 2019 EPI ISL 402128 | C | C | G | C | C | A | C | A | U | G | G | G |
| hCoV 19 Wuhan WIV06 2019 EPI ISL 402129 | C | C | G | C | C | A | C | A | U | G | G | G |
| hCoV 19 Wuhan WIV07 2019 EPI ISL 402130 | C | C | G | C | C | A | C | A | U | G | G | G |
| hCoV 19 Zhejiang WZ 01 2020 EPI ISL 404227 | C | C | G | C | C | A | C | A | U | G | G | G |
| MT012098.1 Severe acute respiratory syndrome coronavirus 2 | C | C | G | C | C | A | C | A | U | G | G | G |
| MT020781.2 Severe acute respiratory syndrome coronavirus 2 | C | C | G | C | C | A | C | A | U | G | G | G |
| MT039887.1 Severe acute respiratory syndrome coronavirus 2 | C | C | G | C | C | A | C | A | U | G | G | G |
| MT044258.1 Severe acute respiratory syndrome coronavirus 2 | C | C | G | C | C | A | C | A | U | G | G | G |
| MT123293.2 Severe acute respiratory syndrome coronavirus 2 | C | C | G | C | C | A | C | A | U | G | G | G |
| MT159716.1 Severe acute respiratory syndrome coronavirus 2 | C | C | G | C | C | A | C | A | U | G | G | G |
| MT192772.1 Severe acute respiratory syndrome coronavirus 2 | C | C | G | C | C | A | C | A | U | G | G | G |
| MT192773.1 Severe acute respiratory syndrome coronavirus 2 | C | C | G | C | C | A | C | A | U | G | G | G |

| | | | | | | | | | | | | |
|---|---|---|---|---|---|---|---|---|---|---|---|---|
| MT049951.1 Severe acute respiratory syndrome coronavirus 2 | C | U | C | C | C | A | C | A | C | G | G | G |
| hCoV 19 USA AZ1 2020 EPI ISL 406223 | C | U | U | C | C | A | C | A | C | G | G | G |
| hCoV 19 Yunnan IVDC YN 003 2020 EPI ISL 408480 | C | U | U | C | C | A | C | A | C | G | G | G |
| MT226610.1 | C | U | U | C | C | A | C | A | C | G | G | G |
| hCoV 19 South Korea KUMC01 2020 EPI ISL 413017 | C | C | G | C | C | A | C | A | C | G | G | G |
| hCoV 19 Wuhan HBCDC HB 06 2020 EPI ISL 412982 | C | U | G | C | C | A | C | G | C | G | G | G |
| hCoV 19 Chongqing YC01 2020 EPI ISL 408478 | C | U | G | C | C | A | U | A | C | G | G | G |
| hCoV 19 Hangzhou ZJU 08 2020 EPI ISL 416473 | C | U | G | C | C | A | U | A | C | G | G | G |
| hCoV 19 USA WA1 2020 EPI ISL 404895 | C | U | G | C | C | A | U | A | C | G | G | G |
| hCoV 19 Australia NSW01 2020 EPI ISL 407893 | C | U | G | C | C | A | C | A | C | G | G | G |
| hCoV 19 Australia QLD01 2020 EPI ISL 407894 | C | U | G | C | C | A | C | A | C | G | G | G |
| hCoV 19 Australia QLD02 2020 EPI ISL 407896 | C | U | G | C | C | A | C | A | C | G | G | G |
| hCoV 19 Australia QLD03 2020 EPI ISL 410717 | C | U | G | C | C | A | C | A | C | G | G | G |
| hCoV 19 Australia QLD04 2020 EPI ISL 410718 | C | U | G | C | C | A | C | A | C | G | G | G |
| hCoV 19 Australia VIC07 2020 EPI ISL 416415 | C | U | G | C | C | A | C | A | C | G | G | G |
| hCoV 19 Beijing 233 2020 EPI ISL 413520 | C | U | G | C | C | A | C | A | C | G | G | G |
| hCoV 19 Beijing IVDC BJ 005 2020 EPI ISL 408485 | C | U | G | C | C | A | C | A | C | G | G | G |
| hCoV 19 Belgium GHB 03021 2020 EPI ISL 407976 | C | U | G | C | C | A | C | A | C | G | G | G |
| hCoV 19 Brazil AMBR 02 2020 EPI ISL 417034 | C | U | G | C | C | A | C | A | C | G | G | G |
| hCoV 19 Chile Santiago 1 2020 EPI ISL 414579 | C | U | G | C | C | A | C | A | C | G | G | G |
| hCoV 19 Chile Talca 1 2020 EPI ISL 414577 | C | U | G | C | C | A | C | A | C | G | G | G |
| hCoV 19 China WF0001 2020 EPI ISL 413691 | C | U | G | C | C | A | C | A | C | G | G | G |
| hCoV 19 China WF0012 2020 EPI ISL 413697 | C | U | G | C | C | A | C | A | C | G | G | G |
| hCoV 19 China WF0014 2020 EPI ISL 413711 | C | U | G | C | C | A | C | A | C | G | G | G |
| hCoV 19 China WF0019 2020 EPI ISL 413749 | C | U | G | C | C | A | C | A | C | G | G | G |
| hCoV 19 China WF0028 2020 EPI ISL 413791 | C | U | G | C | C | A | C | A | C | G | G | G |
| hCoV 19 China WF0029 2020 EPI ISL 413809 | C | U | G | C | C | A | C | A | C | G | G | G |
| hCoV 19 England 02 2020 EPI ISL 407073 | C | U | G | C | C | A | C | A | C | G | G | G |
| hCoV 19 France GE1583 2020 EPI ISL 414600 | C | U | G | C | C | A | C | A | C | G | G | G |
| hCoV 19 France GE1583 2020 EPI ISL 414623 | C | U | G | C | C | A | C | A | C | G | G | G |
| hCoV 19 Georgia Tb 390 2020 EPI ISL 416477 | C | U | G | C | C | A | C | A | C | G | G | G |
| hCoV 19 Germany BavPat3 2020 EPI ISL 414521 | C | U | G | C | C | A | C | A | C | G | G | G |
| hCoV 19 Guangdong 2020XN4459 P0041 2020 EPI ISL 413858 | C | U | G | C | C | A | C | A | C | G | G | G |
| hCoV 19 Guangdong 2020XN4475 P0042 2020 EPI ISL 413854 | C | U | G | C | C | A | C | A | C | G | G | G |
| hCoV 19 Guangdong 20SF012 2020 EPI ISL 403932 | C | U | G | C | C | A | C | A | C | G | G | G |
| hCoV 19 Guangzhou GZMU0016 2020 EPI ISL 414663 | C | U | G | C | C | A | C | A | C | G | G | G |
| hCoV 19 Guangzhou GZMU0042 2020 EPI ISL 414688 | C | U | G | C | C | A | C | A | C | G | G | G |
| hCoV 19 India 1 31 2020 EPI ISL 413523 | C | U | G | C | C | A | C | A | C | G | G | G |
| hCoV 19 Japan TY WK 012 2020 EPI ISL 408665 | C | U | G | C | C | A | C | A | C | G | G | G |
| hCoV 19 Japan TY WK 501 2020 EPI ISL 408666 | C | U | G | C | C | A | C | A | C | G | G | G |
| hCoV 19 Japan TY WK 521 2020 EPI ISL 408667 | C | U | G | C | C | A | C | A | C | G | G | G |
| hCoV 19 Netherlands Gelderland 1 2020 EPI ISL 415461 | C | U | G | C | C | A | C | A | C | G | G | G |
| hCoV 19 Netherlands NoordBrabant 46 2020 EPI ISL 415503 | C | U | G | C | C | A | C | A | C | G | G | G |
| hCoV 19 Netherlands Utrecht 16 2020 EPI ISL 414555 | C | U | G | C | C | A | C | A | C | G | G | G |
| hCoV 19 New Zealand 20VR0275 2020 EPI ISL 416538 | C | U | G | C | C | A | C | A | C | G | G | G |
| hCoV 19 New Zealand 20VR0276 2020 EPI ISL 416539 | C | U | G | C | C | A | C | A | C | G | G | G |
| hCoV 19 Shandong LY005 2020 EPI ISL 414938 | C | U | G | C | C | A | C | A | C | G | G | G |
| hCoV 19 Shandong LY007 2020 EPI ISL 414940 | C | U | G | C | C | A | C | A | C | G | G | G |
| hCoV 19 Shandong LY008 2020 EPI ISL 414941 | C | U | G | C | C | A | C | A | C | G | G | G |
| hCoV 19 Shanghai SH0002 2020 EPI ISL 416316 | C | U | G | C | C | A | C | A | C | G | G | G |
| hCoV 19 Shanghai SH0003 2020 EPI ISL 416317 | C | U | G | C | C | A | C | A | C | G | G | G |
| hCoV 19 Shanghai SH0004 2020 EPI ISL 416318 | C | U | G | C | C | A | C | A | C | G | G | G |
| hCoV 19 Shanghai SH0009 2020 EPI ISL 416322 | C | U | G | C | C | A | C | A | C | G | G | G |
| hCoV 19 Shanghai SH0010 2020 EPI ISL 416323 | C | U | G | C | C | A | C | A | C | G | G | G |
| hCoV 19 Shanghai SH0013 2020 EPI ISL 416326 | C | U | G | C | C | A | C | A | C | G | G | G |
| hCoV 19 Shanghai SH0021 2020 EPI ISL 416330 | C | U | G | C | C | A | C | A | C | G | G | G |
| hCoV 19 Shanghai SH0026 2020 EPI ISL 416335 | C | U | G | C | C | A | C | A | C | G | G | G |
| hCoV 19 Shanghai SH0029 2020 EPI ISL 416338 | C | U | G | C | C | A | C | A | C | G | G | G |
| hCoV 19 Shanghai SH0030 2020 EPI ISL 416339 | C | U | G | C | C | A | C | A | C | G | G | G |
| hCoV 19 Shanghai SH0032 2020 EPI ISL 416341 | C | U | G | C | C | A | C | A | C | G | G | G |
| hCoV 19 Shanghai SH0059 2020 EPI ISL 416366 | C | U | G | C | C | A | C | A | C | G | G | G |
| hCoV 19 Shanghai SH0074 2020 EPI ISL 416377 | C | U | G | C | C | A | C | A | C | G | G | G |
| hCoV 19 Shanghai SH0115 2020 EPI ISL 416402 | C | U | G | C | C | A | C | A | C | G | G | G |
| hCoV 19 Shanghai SH0117 2020 EPI ISL 416403 | C | U | G | C | C | A | C | A | C | G | G | G |
| hCoV 19 Shanghai SH0128 2020 EPI ISL 416409 | C | U | G | C | C | A | C | A | C | G | G | G |
| hCoV 19 Shenzhen HKU SZ 002 2020 EPI ISL 406030 | C | U | G | C | C | A | C | A | C | G | G | G |
| hCoV 19 Shenzhen HKU SZ 005 2020 EPI ISL 405839 | C | U | G | C | C | A | C | A | C | G | G | G |
| hCoV 19 Shenzhen SZTH 002 2020 EPI ISL 406593 | C | U | G | C | C | A | C | A | C | G | G | G |
| hCoV 19 Sichuan IVDC SC 001 2020 EPI ISL 408484 | C | U | G | C | C | A | C | A | C | G | G | G |
| hCoV 19 South Korea KCDC03 2020 EPI ISL 407193 | C | U | G | C | C | A | C | A | C | G | G | G |
| hCoV 19 South Korea KUMC03 2020 EPI ISL 413513 | C | U | G | C | C | A | C | A | C | G | G | G |
| hCoV 19 Spain Valencia5 2020 EPI ISL 416484 | C | U | G | C | C | A | C | A | C | G | G | G |
| hCoV 19 USA CA1 2020 EPI ISL 406034 | C | U | G | C | C | A | C | A | C | G | G | G |
| hCoV 19 USA IL2 2020 EPI ISL 410045 | C | U | G | C | C | A | C | A | C | G | G | G |
| hCoV 19 USA TX1 2020 EPI ISL 411956 | C | U | G | C | C | A | C | A | C | G | G | G |
| hCoV 19 USA WA UW158 2020 EPI ISL 416696 | C | U | G | C | C | A | C | A | C | G | G | G |
| hCoV 19 Vietnam CM99 2020 EPI ISL 416429 | C | U | G | C | C | A | C | A | C | G | G | G |
| hCoV 19 Vietnam VR03 38142 2020 EPI ISL 408668 | C | U | G | C | C | A | C | A | C | G | G | G |
| hCoV 19 Wuhan HBCDC HB 02 2020 EPI ISL 412978 | C | U | G | C | C | A | C | A | C | G | G | G |
| hCoV 19 Wuhan HBCDC HB 03 2020 EPI ISL 412979 | C | U | G | C | C | A | C | A | C | G | G | G |
| hCoV 19 Wuhan HBCDC HB 04 2020 EPI ISL 412980 | C | U | G | C | C | A | C | A | C | G | G | G |
| hCoV 19 Wuhan WH04 2020 EPI ISL 406801 | C | U | G | C | C | A | C | A | C | G | G | G |
| MT123292.2 Severe acute respiratory syndrome coronavirus 2 | C | U | G | C | C | A | C | A | C | G | G | G |
| hCoV 19 Netherlands NA 35 2020 EPI ISL 415492 | C | C | U | U | C | A | C | A | U | G | G | G |

| | | | | | | | | | | | | |
|---|---|---|---|---|---|---|---|---|---|---|---|---|
| hCoV 19 USA UPHL 01 2020 EPI ISL 415539 | C | C | U | C | C | A | U | A | U | G | G | G |
| hCoV 19 Australia NSW05 2020 EPI ISL 412975 | C | C | U | C | C | A | C | A | U | G | G | G |
| hCoV 19 Australia NSW06 2020 EPI ISL 413213 | C | C | U | C | C | A | C | A | U | G | G | G |
| hCoV 19 Australia NSW07 2020 EPI ISL 413214 | C | C | U | C | C | A | C | A | U | G | G | G |
| hCoV 19 Australia QLD09 2020 EPI ISL 414414 | C | C | U | C | C | A | C | A | U | G | G | G |
| hCoV 19 Australia VIC04 2020 EPI ISL 416412 | C | C | U | C | C | A | C | A | U | G | G | G |
| hCoV 19 Belgium ULG 6638 2020 EPI ISL 417017 | C | C | U | C | C | A | C | A | U | G | G | G |
| hCoV 19 Brazil SPBR 02 2020 EPI ISL 413016 | C | C | U | C | C | A | C | A | U | G | G | G |
| hCoV 19 Canada BC 02421 2020 EPI ISL 415581 | C | C | U | C | C | A | C | A | U | G | G | G |
| hCoV 19 Chongqing IVDC CQ 001 2020 EPI ISL 408481 | C | C | U | C | C | A | C | A | U | G | G | G |
| hCoV 19 England SHEF BFCEE 2020 EPI ISL 416733 | C | C | U | C | C | A | C | A | U | G | G | G |
| hCoV 19 Finland FIN03032020B 2020 EPI ISL 413603 | C | C | U | C | C | A | C | A | U | G | G | G |
| hCoV 19 France B2334 2020 EPI ISL 416503 | C | C | U | C | C | A | C | A | U | G | G | G |
| hCoV 19 France B2340 2020 EPI ISL 416507 | C | C | U | C | C | A | C | A | U | G | G | G |
| hCoV 19 France IDF0515 2020 EPI ISL 408430 | C | C | U | C | C | A | C | A | U | G | G | G |
| hCoV 19 France IDF0515 isl 2020 EPI ISL 410984 | C | C | U | C | C | A | C | A | U | G | G | G |
| hCoV 19 Georgia Tb 468 2020 EPI ISL 415643 | C | C | U | C | C | A | C | A | U | G | G | G |
| hCoV 19 Georgia Tb 537 2020 EPI ISL 416480 | C | C | U | C | C | A | C | A | U | G | G | G |
| hCoV 19 Georgia Tb 54 2020 EPI ISL 415641 | C | C | U | C | C | A | C | A | U | G | G | G |
| hCoV 19 Germany BavPat2 2020 EPI ISL 414520 | C | C | U | C | C | A | C | A | U | G | G | G |
| hCoV 19 Hangzhou ZJU 01 2020 EPI ISL 415709 | C | C | U | C | C | A | C | A | U | G | G | G |
| hCoV 19 Hong Kong VB20026565 2020 EPI ISL 412030 | C | C | U | C | C | A | C | A | U | G | G | G |
| hCoV 19 Italy SPL1 2020 EPI ISL 412974 | C | C | U | C | C | A | C | A | U | G | G | G |
| hCoV 19 Japan DP0027 2020 EPI ISL 416566 | C | C | U | C | C | A | C | A | U | G | G | G |
| hCoV 19 Japan DP0037 2020 EPI ISL 416567 | C | C | U | C | C | A | C | A | U | G | G | G |
| hCoV 19 Japan DP0059 2020 EPI ISL 416569 | C | C | U | C | C | A | C | A | U | G | G | G |
| hCoV 19 Japan DP0065 2020 EPI ISL 416570 | C | C | U | C | C | A | C | A | U | G | G | G |
| hCoV 19 Japan DP0077 2020 EPI ISL 416571 | C | C | U | C | C | A | C | A | U | G | G | G |
| hCoV 19 Japan DP0078 2020 EPI ISL 416572 | C | C | U | C | C | A | C | A | U | G | G | G |
| hCoV 19 Japan DP0152 2020 EPI ISL 416578 | C | C | U | C | C | A | C | A | U | G | G | G |
| hCoV 19 Japan DP0158 2020 EPI ISL 416579 | C | C | U | C | C | A | C | A | U | G | G | G |
| hCoV 19 Japan DP0190 2020 EPI ISL 416581 | C | C | U | C | C | A | C | A | U | G | G | G |
| hCoV 19 Japan DP0191 2020 EPI ISL 416582 | C | C | U | C | C | A | C | A | U | G | G | G |
| hCoV 19 Japan DP0196 2020 EPI ISL 416583 | C | C | U | C | C | A | C | A | U | G | G | G |
| hCoV 19 Japan DP0200 2020 EPI ISL 416584 | C | C | U | C | C | A | C | A | U | G | G | G |
| hCoV 19 Japan DP0236 2020 EPI ISL 416585 | C | C | U | C | C | A | C | A | U | G | G | G |
| hCoV 19 Japan DP0274 2020 EPI ISL 416586 | C | C | U | C | C | A | C | A | U | G | G | G |
| hCoV 19 Japan DP0278 2020 EPI ISL 416587 | C | C | U | C | C | A | C | A | U | G | G | G |
| hCoV 19 Japan DP0287 2020 EPI ISL 416589 | C | C | U | C | C | A | C | A | U | G | G | G |
| hCoV 19 Japan DP0290 2020 EPI ISL 416591 | C | C | U | C | C | A | C | A | U | G | G | G |
| hCoV 19 Japan DP0294 2020 EPI ISL 416592 | C | C | U | C | C | A | C | A | U | G | G | G |
| hCoV 19 Japan DP0311 2020 EPI ISL 416593 | C | C | U | C | C | A | C | A | U | G | G | G |
| hCoV 19 Japan DP0319 2020 EPI ISL 416594 | C | C | U | C | C | A | C | A | U | G | G | G |
| hCoV 19 Japan DP0346 2020 EPI ISL 416597 | C | C | U | C | C | A | C | A | U | G | G | G |
| hCoV 19 Japan DP0357 2020 EPI ISL 416598 | C | C | U | C | C | A | C | A | U | G | G | G |
| hCoV 19 Japan DP0361 2020 EPI ISL 416599 | C | C | U | C | C | A | C | A | U | G | G | G |
| hCoV 19 Japan DP0457 2020 EPI ISL 416601 | C | C | U | C | C | A | C | A | U | G | G | G |
| hCoV 19 Japan DP0464 2020 EPI ISL 416603 | C | C | U | C | C | A | C | A | U | G | G | G |
| hCoV 19 Japan DP0543 2020 EPI ISL 416607 | C | C | U | C | C | A | C | A | U | G | G | G |
| hCoV 19 Japan DP0568 2020 EPI ISL 416609 | C | C | U | C | C | A | C | A | U | G | G | G |
| hCoV 19 Japan DP0588 2020 EPI ISL 416610 | C | C | U | C | C | A | C | A | U | G | G | G |
| hCoV 19 Japan DP0645 2020 EPI ISL 416612 | C | C | U | C | C | A | C | A | U | G | G | G |
| hCoV 19 Japan DP0687 2020 EPI ISL 416614 | C | C | U | C | C | A | C | A | U | G | G | G |
| hCoV 19 Japan DP0699 2020 EPI ISL 416617 | C | C | U | C | C | A | C | A | U | G | G | G |
| hCoV 19 Japan DP0724 2020 EPI ISL 416620 | C | C | U | C | C | A | C | A | U | G | G | G |
| hCoV 19 Japan DP0743 2020 EPI ISL 416621 | C | C | U | C | C | A | C | A | U | G | G | G |
| hCoV 19 Japan DP0752 2020 EPI ISL 416622 | C | C | U | C | C | A | C | A | U | G | G | G |
| hCoV 19 Japan DP0764 2020 EPI ISL 416624 | C | C | U | C | C | A | C | A | U | G | G | G |
| hCoV 19 Japan DP0765 2020 EPI ISL 416625 | C | C | U | C | C | A | C | A | U | G | G | G |
| hCoV 19 Japan DP0786 2020 EPI ISL 416628 | C | C | U | C | C | A | C | A | U | G | G | G |
| hCoV 19 Japan DP0804 2020 EPI ISL 416631 | C | C | U | C | C | A | C | A | U | G | G | G |
| hCoV 19 Japan DP0827 2020 EPI ISL 416632 | C | C | U | C | C | A | C | A | U | G | G | G |
| hCoV 19 Japan DP0880 2020 EPI ISL 416633 | C | C | U | C | C | A | C | A | U | G | G | G |
| hCoV 19 Japan Hu DP Kng 19 027 2020 EPI ISL 412969 | C | C | U | C | C | A | C | A | U | G | G | G |
| hCoV 19 Kuwait KU09 2020 EPI ISL 416541 | C | C | U | C | C | A | C | A | U | G | G | G |
| hCoV 19 Kuwait KU17 2020 EPI ISL 416542 | C | C | U | C | C | A | C | A | U | G | G | G |
| hCoV 19 Netherlands NA 21 2020 EPI ISL 415478 | C | C | U | C | C | A | C | A | U | G | G | G |
| hCoV 19 Netherlands NA 29 2020 EPI ISL 415486 | C | C | U | C | C | A | C | A | U | G | G | G |
| hCoV 19 Netherlands NA 34 2020 EPI ISL 415491 | C | C | U | C | C | A | C | A | U | G | G | G |
| hCoV 19 Netherlands NoordBrabant 56 2020 EPI ISL 415512 | C | C | U | C | C | A | C | A | U | G | G | G |
| hCoV 19 Netherlands NoordBrabant 6 2020 EPI ISL 414451 | C | C | U | C | C | A | C | A | U | G | G | G |
| hCoV 19 Netherlands Utrecht 18 2020 EPI ISL 415527 | C | C | U | C | C | A | C | A | U | G | G | G |
| hCoV 19 Netherlands ZuidHolland 16 2020 EPI ISL 414558 | C | C | U | C | C | A | C | A | U | G | G | G |
| hCoV 19 Netherlands ZuidHolland 8 2020 EPI ISL 414468 | C | C | U | C | C | A | C | A | U | G | G | G |
| hCoV 19 Saudi Arabia KAIMRC Alghoribi 2020 EPI ISL 416432 | C | C | U | C | C | A | C | A | U | G | G | G |
| hCoV 19 Scotland CVR07 2020 EPI ISL 415630 | C | C | U | C | C | A | C | A | U | G | G | G |
| hCoV 19 Scotland CVR10 2020 EPI ISL 415631 | C | C | U | C | C | A | C | A | U | G | G | G |
| hCoV 19 Shandong IVDC SD 001 2020 EPI ISL 408482 | C | C | U | C | C | A | C | A | U | G | G | G |
| hCoV 19 Shanghai SH0022 2020 EPI ISL 416331 | C | C | U | C | C | A | C | A | U | G | G | G |
| hCoV 19 Shanghai SH0070 2020 EPI ISL 416373 | C | C | U | C | C | A | C | A | U | G | G | G |
| hCoV 19 Shanghai SH0080 2020 EPI ISL 416382 | C | C | U | C | C | A | C | A | U | G | G | G |
| hCoV 19 Shanghai SH0126 2020 EPI ISL 416407 | C | C | U | C | C | A | C | A | U | G | G | G |
| hCoV 19 Singapore 3 2020 EPI ISL 407988 | C | C | U | C | C | A | C | A | U | G | G | G |
| hCoV 19 Taiwan CGMH CGU 03 2020 EPI ISL 415741 | C | C | U | C | C | A | C | A | U | G | G | G |

| | | | | | | | | | | | | |
|---|---|---|---|---|---|---|---|---|---|---|---|---|
| hCoV 19 USA CruiseA 23 2020 EPI ISL 414482 | C | C | U | C | C | A | C | A | U | G | G | G |
| hCoV 19 USA WA UW108 2020 EPI ISL 416646 | C | C | U | C | C | A | C | A | U | G | G | G |
| hCoV 19 USA WA UW137 2020 EPI ISL 416675 | C | C | U | C | C | A | C | A | U | G | G | G |
| hCoV 19 USA WA UW168 2020 EPI ISL 416706 | C | C | U | C | C | A | C | A | U | G | G | G |
| hCoV 19 USA WA UW185 2020 EPI ISL 416723 | C | C | U | C | C | A | C | A | U | G | G | G |
| hCoV 19 USA WA UW22 2020 EPI ISL 414591 | C | C | U | C | C | A | C | A | U | G | G | G |
| hCoV 19 USA WA3 UW1 2020 EPI ISL 413025 | C | C | U | C | C | A | C | A | U | G | G | G |
| MT126808.1 Severe acute respiratory syndrome coronavirus 2 | C | C | U | C | C | A | C | A | U | G | G | G |
